# *N*-[(Thiophen-3-yl)methyl]benzamides as Influenza Virus Fusion Inhibitors Acting on H1 and H5 Hemagglutinins

**DOI:** 10.1101/2025.07.28.667118

**Authors:** Silke Rimaux, Aitor Valdivia, Juan Martín-López, Valeria Francesconi, Celia Escriche, Cato Mestdagh, Ria Van Berwaer, Lieselotte Schurmans, Kaat Verleye, Samuel Noppen, Óscar Lozano, Annelies Stevaert, F. Javier Luque, Lieve Naesens, Santiago Vázquez

## Abstract

Novel antiviral drugs are needed to prepare against infections from influenza A virus (IAV). Here a series of *N*-[(thiophen-3-yl)methylbenzamides which target the hemagglutinin (HA)-mediated fusion process is reported. The most active compound, **VF-57a**, displays a 50% effective concentration (EC_50_) of ∼0.8 μM and antiviral selectivity index >130, in Madin-Darby canine kidney (MDCK) cells infected with A/H1N1 virus. **VF-57a** proved to be a strong inhibitor of A/H1N1- and A/H5N1-pseudovirus entry (EC_50_ values of 0.3 and 0.8 µM, respectively). Cell-cell fusion assays in HA-expressing cells, surface plasmon resonance-based assessment of HA protein refolding, and resistance studies suggested that **VF-57a** prevents the conformational change of HA at acidic pH. Molecular modelling highlighted the role of the dimethylthiophene moiety and the amide-based tether in the anchoring to the binding cavity of HA. Our findings support further development of this class of IAV fusion inhibitors against A/H1N1 and A/H5N1 viruses.

## INTRODUCTION

According to the World Health Organization, the annual epidemics caused by influenza A and B viruses are globally responsible for ∼470,000 respiratory deaths per year.^1^ In addition, antigenically distinct influenza A viruses (IAVs), emerging from zoonotic sources, can cause pandemics with potentially grave consequences,^2^ as evident from the estimated toll of ∼200,000 respiratory deaths and ∼83,000 cardiovascular deaths, during the last IAV pandemic that occurred in 2009.^3^ Nowadays, a highly pathogenic avian influenza A/H5N1 virus is causing great concern, after recent outbreaks in dairy cattle in the USA and fur farms in Europe, with regular spillover to human individuals.^4-6^ Although annual influenza vaccination represents the main prophylaxis for people at risk, such as the elderly or persons suffering from comorbidities, the overall effectiveness of the current vaccines is only between 30 and 60%.^7^ Therefore, antiviral drugs are crucial to treat influenza-infected persons who are seriously ill or at high risk for severe complications.^8, 9^ Oseltamivir and other neuraminidase inhibitors are the standard-of-care since many years, while the polymerase inhibitors baloxavir, marboxil and favipiravir are also available in certain countries.^10, 11^ For both pharmacological classes, close monitoring of potential resistance is needed.^12^ Other classes of easily accessible and cost-effective antiviral molecules remain urgently needed to prepare against the threat of influenza pandemics. This includes exploration of alternative druggable targets, such as the viral hemagglutinin (HA).^13^ The small molecule arbidol (**1**) (Chart 1) acts on HA as well as other targets,^14, 15^ and is currently approved in two countries (Russia and China).

HA is a homotrimeric protein embedded in the viral envelope. As the key player for viral entry into the host cells, HA forms the initial binding interaction with sialylated glycans on the host cell surface, which results in uptake of the virus by endocytosis.^2, 16^ As the endosomes mature to reach a pH of ∼5.5, HA is triggered to undergo a drastic conformational change in its stem region,^13, 17-19^ which promotes the release of the hydrophobic fusion peptide and, ultimately, fusion of the endosomal membrane and viral envelope. Diverse strategies are being developed to interfere with the viral entry process,^20-23^ yet many of these are challenged by the high variability of HA. The 19 currently known IAV HA subtypes fall into two phylogenetic groups. The H1, H2 and H5 HAs belong to group 1, whereas the H3 and H7 HAs fall in group 2.^21^ Broadly neutralizing anti-HA antibodies with group 1-, group 2-, or pan-IAV activity have sparked major interest from the pharmaceutical industry, with a handful having progressed towards clinical evaluation.^24^

Besides, there is significant interest in small molecule inhibitors of the HA-mediated membrane fusion process. Several fusion inhibitors have been reported in the literature, with extensive variety in their scaffold structures (Chart 1). In almost all cases, the activity proved HA subtype-dependent, covering either H1 or H3 HA, but not both, although several molecules showed inhibition of more than one HA subtype of the same group (Chart 1). A few fusion inhibitors were also validated in influenza mouse models.^25-27^ So far, arbidol (**1**),^14, 15^ M090 (**2**),^28^ OA-10 (**3**),^29^ and **4**^30^ are exceptional in having activity against both H1 and H3 HAs, albeit at potencies that are quite low compared to the subtype- or group-restricted inhibitors.

Up to now, structural studies have revealed that fusion inhibitors bind to two sites in the HA stem region (Figure 1). Site A corresponds to the pocket occupied by *t*-butylhydroquinone (**5**) (TBHQ; PDB IDs 3EYK and 3EYM)^31^ and arbidol (PDB IDs 5T6N and 5T6S),^14^ and consists of residues located at helices A and C from protomer 1 and C’ from protomer 2. Site B is the groove filled by JNJ4796 (**9**) (PDB IDs 6CFG and 6CF7),^25^ (*S*)-F0045 (**10**) (PDB ID 6WCR)^32^ and CBS1117, (**11a**) (PDB ID 6VMZ).^33^ Considering the substantial chemical diversity among the reported fusion inhibitors (Chart 1) and the structural plasticity of HA, understanding the binding mode is crucial to rationalize the structure-activity relationships (SAR) and conduct lead optimization.

**Figure 1.**
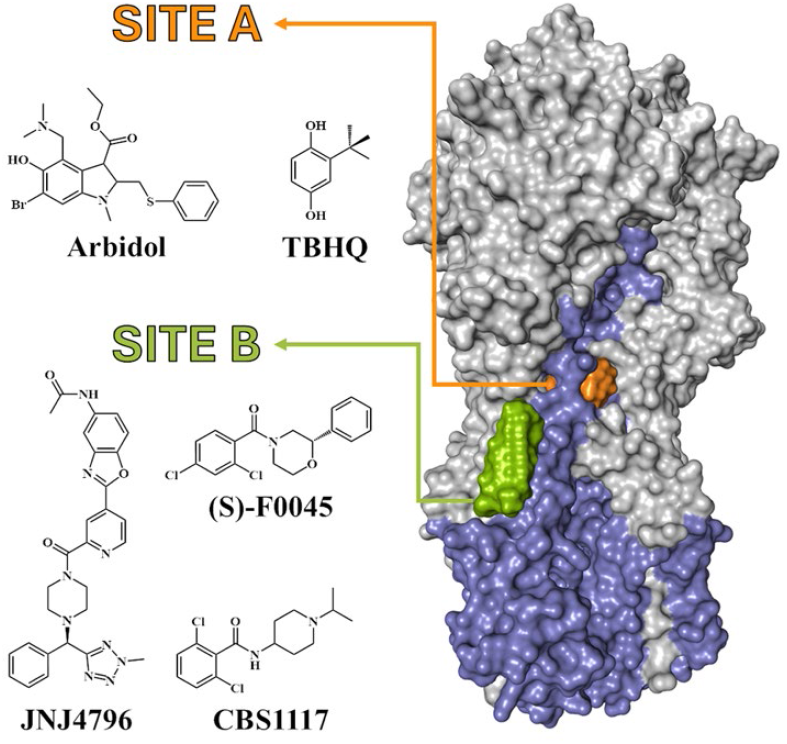
Surface representation of the HA protein (HA1: grey; HA2: lavender) and the ligand binding sites of TBHQ and arbidol (Site A; orange surface) and JNJ4796, (*S*)-F0045 and CBS1117 (green surface).

This study elaborates on the chemotype for which the first prototype, CL385319 (**19**, Chart 1), was published in 1999.^34^ The authors demonstrated antiviral activity against A/H1N1 and A/H2N2 viruses (group 1), with A/H3N2 (group 2) being much less sensitive. Several years later, another team found that CL385319 is also effective against A/H5N1 virus.^35^ Subsequent studies explored the effect of replacing the piperidine ring of CL385319 by a variety of heteroaromatic rings. This led to submicromolar potencies, as in the case of the 2-(thiophenyl-2-yl)ethyl derivative **20** (Chart 1), which exhibited an EC_50_ of 0.22 µM against A/H5N1 pseudovirus.^36^ These researchers also identified an oligothiophene compound having 58-fold higher potency than CL385319 in A/H1N1 virus-infected cell cultures and 360-fold higher activity in an A/H5N1 pseudovirus entry assay.^37^ Finally, in 2019, a systematic SAR investigation on a series of heteroaromatic-based benzenesulfonamides, leading to compound **21**, which exhibited an EC_50_ of 0.47 µM against A/H5N1 virus, was reported.^38^

Due to the promising anti-IAV activities of benzamides **19** and **20** and of compound **21**, we decided to further investigate the SAR, considering a series of hybrid molecules featuring, on the left-hand side, the benzamide core of CL385319 and its analog **20** and, on the right-hand side, the *N*-[(thiophen-3-yl)methyl]amino unit of **21** (Scheme 1). Starting from hybrid molecule **22**, chemical variations were designed to explore (i) the effects of substitutions attached to the phenyl ring (Charts 2-4); (ii) the optimal substitution for the thiophene ring (Charts 5-6); and (iii) the nature of the linker between the thiophene and the phenyl moieties (Chart 7).

The anti-IAV activity was evaluated in cell-based assays with different A/H1N1 and A/H5N1 (pseudo)viruses, and the fusion-inhibiting effect was determined in HA-expressing cells. To understand the HA binding mode, resistant virus was selected by serial passaging. Finally, molecular dynamics simulations and free energy (Thermodynamic Integration) calculations were conducted to identify the plausible binding site of the lead compound within the HA protein.

## RESULTS AND DISCUSSION

### Synthesis and SAR analysis

The benzamides were easily synthesized through the reaction of the required acyl chloride, either commercially available or freshly synthesized from its corresponding benzoic acid derivative and thionyl chloride, with the required amine, typically (2,5-dimethyl-thiophen-3-yl)methanamine. Some particular compounds, such as ester **63**, secondary amine **64** and inverse amide **65**, were synthesized using specific procedures as reported in the Experimental Section. All the new compounds were fully characterized through their spectroscopic data and elemental analyses or HPLC/MS (see Experimental Section and Supporting Information for further details).

Starting from hybrid molecule **22** (Scheme 1), the SAR analysis of the *N*-[(thiophen-3-yl)methylbenzamides aimed to gain insight into how the anti-IAV activity depends on the thiophene and phenyl rings, and the linker that joins these moieties.

**Chart 1.**
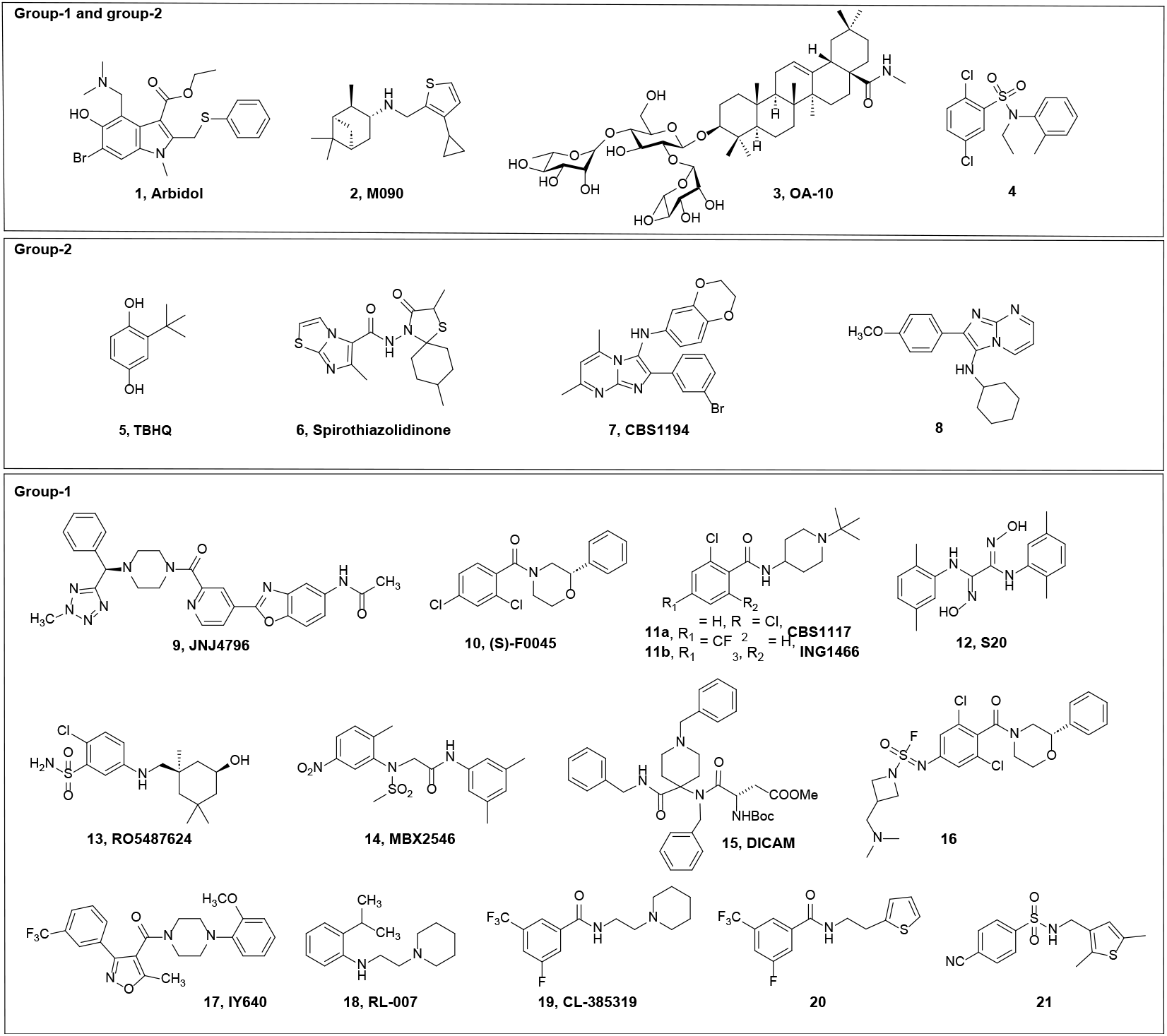
Structures of reported inhibitors of IAV HA-mediated fusion, specifically: arbidol,^14, 15^ M090,^28^ OA-10,^29^ 4,^30^ TBHQ,^31^ spirothiazolidinones,^39, 40^ CBS1194,^41^ 8,^42^ JNJ4796,^25^ (*S*)-F0045,^32^ CBS1117,^33^ ING1466,^26^ S20,^43^ RO5487624,^27^ MBX-2456,^44^ DICAM,^45^ 16,^46^ IY640,^47^ RL-007 (9d in the reference),^48^ CL-385319,^34^ **20**,^36^ and **21**.^38^

With regard to the benzene ring of **22**, besides the replacement of fluorine by bromine, chlorine or trifluoromethyl in *meta* position and some related combinations shown in Chart 2, we also explored the effect of attaching additional electron-withdrawing groups in position *ortho* and *para* (Chart 3). On the other hand, since previous works on CL-385319 and **20** did not perform a full SAR study of the benzene ring, a classical Topliss approach was adopted to consider a more diverse range of substituents attached to the benzene ring (Chart 4).^49,50^

Regarding the thiophene ring, attention was paid to the influence of the methyl groups present in positions 2 and 5, which were removed either separately or simultaneously (Chart 5). Furthermore, the replacement of the thiophene moiety by more polar rings (furan, thiazole, isoxazole, pyrazole, oxazole and pyridine) was also examined (Chart 6).

Finally, we envisaged the replacement of the methylamide linker by other chemical moieties, keeping the total length of the tether between both aromatic rings (Chart 7).

To determine the anti-influenza virus activity, we conducted cytopathic effect (CPE) reduction assays in MDCK cells, in which we used the colorimetric MTS cell viability method to measure the compounds’ protective effect against viral CPE, as well as their cytotoxicity in mock-infected cells. Parallel with the MTS readout, we scored the CPE and cytotoxicity by microscopic inspection; the two readout methods gave very similar results. The virus test panel included two A/H1N1 (PR8 and Virg09), two A/H3N2 (A/Victoria/361/11 and A/Hong Kong/7/87) and one influenza B strain (B/Ned/537/05; Yamagata lineage). This evaluation showed that several members of the series exhibited strong activity against A/H1N1 virus (see Table 1, which only shows the results for the most active compounds). In line with the reported data on the congeners CL-385319,^34^ **20**^36^ and **21**,^38^ we saw no inhibitory effect on A/H3N2 and influenza B viruses (highest tested concentration: 100 μM, data not shown).

**Table 1.** Antiviral (A/H1N1) activity and cytotoxicity in MDCK cells.

| Cpd | R<sub>1</sub> | R<sub>2</sub> | R<sub>3</sub> | R<sub>4</sub> | R<sub>5</sub> | R<sub>6</sub> | R<sub>7</sub> | Antiviral EC<sub>50</sub><sup>a</sup> against A/H1N1 virus (μM) |  |  |  | Cytotoxicity (μM) |  |
| --- | --- | --- | --- | --- | --- | --- | --- | --- | --- | --- | --- | --- | --- |
|  |  |  |  |  |  |  |  | PR8 strain |  | Virg09 strain |  | CC<sub>50</sub><sup>b</sup> | MCC<sup>c</sup> |
|  |  |  |  |  |  |  |  | MTS | Microscopy | MTS | Microscopy |  |  |
| Series A |  |  |  |  |  |  |  |  |  |  |  |  |  |
| 22 | H | F | H | CF<sub>3</sub> | H | Me | Me | 1.1 ± 0.3 | 1.5 ± 0.6 | 0.44 ± 0.19 | 1.7 ± 1.1 | >100 | >100 |
| 23 | H | Cl | H | CF<sub>3</sub> | H | Me | Me | 0.92 ± 0.19 | 1.1 ± 0.2 | 0.31 ± 0.12 | 0.71 ± 0.45 | >100 | >100 |
| 35 | H | F | F | CF<sub>3</sub> | H | Me | Me | 0.22 ± 0.03 | 0.86 ± 0.41 | 0.20 ± 0.05 | 0.89 ± 0.13 | >100 | >100 |
| 37 | F | Cl | H | CF<sub>3</sub> | H | Me | Me | 2.9 | 1.6 ± 0.8 | >100 | >100 | >100 | 56 ± 22 |
| 38 | H | Cl | F | CF<sub>3</sub> | H | Me | Me | 0.27 ± 0.12 | 1.6 ± 0.8 | 0.18 ± 0.04 | 3.1 ± 0.5 | >100 | 40 |
| 39 | H | Cl | H | CF<sub>3</sub> | Cl | Me | Me | 5.9 ± 3.2 | 11 ± 6 | 0.82 ± 0.61 | 8.1 ± 0.5 | >100 | 52 ± 25 |
| 45 | H | F | H | CF<sub>3</sub> | H | H | Me | 0.68 ± 0.51 | 0.26 ± 0.02 | 0.39 ± 0.18 | 1.6 ± 0.5 | 67 ± 1 | 78 ± 22 |
| 46 | H | F | H | CF<sub>3</sub> | H | Me | H | 0.23 ± 0.08 | 0.17 ± 0.04 | 6.8 ± 6.5 | 8.3 ± 5.1 | >100 | 67 ± 33 |
| 47 | H | F | H | CF<sub>3</sub> | H | H | H | 0.42 ± 0.10 | 0.44 ± 0.16 | 7.2 ± 4.1 | 20 ± 8 | >100 | >100 |
| 48 | H | Cl | H | CF<sub>3</sub> | H | H | Me | 0.34 ± 0.18 | 0.46 ± 0.14 | 0.15 ± 0.12 | 1.9 ± 1.1 | >100 | >100 |
| 49 | H | Cl | H | CF<sub>3</sub> | H | Me | H | 0.14 ± 0.04 | 0.14 ± 0.05 | 0.37 ± 0.16 | 0.48 ± 0.07 | >100 | 78 ± 22 |
| 50 | H | Cl | H | CF<sub>3</sub> | H | H | H | 2.0 ± 1.0 | 1.0 ± 0.6 | 1.1 ± 0.5 | 2.1 ± 0.0 | 70 ± 2 | 100 ± 0 |
| 51 | H | F | F | CF<sub>3</sub> | H | H | Me | 2.9 ± 1.1 | 3.5 ± 1.2 | 5.3 ± 0.8 | >100 | 24 ± 2 | 33 ± 0 |
| 52 | H | Cl | F | CF<sub>3</sub> | H | H | Me | 8.5 ± 5.6 | 13 ± 8 | >100 | >100 | >100 | >100 |
| 53 | H | Cl | H | CF<sub>3</sub> | Cl | H | Me | 17 ± 8 | >100 | 31 ± 3 | >100 | >100 | >100 |
| Series B |  |  |  |  |  |  |  |  |  |  |  |  |  |
| 54 | H | F | H | CF<sub>3</sub> | H | Me | Me | 0.47 ± 0.19 | 0.54 ± 0.17 | 0.54 ± 0.06 | 3.6 ± 2.8 | 79 ± 5 | 100 ± 0 |
| 55 | H | Cl | H | CF<sub>3</sub> | H | Me | Me | 0.24 ± 0.08 | 0.54 ± 0.37 | 1.1 ± 0.3 | >100 | 29 ± 0 | 33 ± 0 |
| Reference compounds |  |  |  |  |  |  |  |  |  |  |  |  |  |
| RL-007 |  |  |  |  |  |  |  | 1.4 ± 0.0 | 1.5 ± 0.1 | 0.61 ± 0.06 | 0.69 ± 0.09 | 33 ± 5 | >40 |
| Ribavirin |  |  |  |  |  |  |  | 38 ± 3 | 27 ± 1 | 20 ± 2 | 25 ± 0 | >250 | 80 ± 20 |
| BXA (nM)<sup>d</sup> |  |  |  |  |  |  |  | 6.1 ± 1.7 | 6.1 ± 2.0 | 3.2 ± 0.4 | 3.5 ± 0.0 | >100 | >100 |

<sup>a</sup>EC<sub>50</sub>: half-maximal effective concentration for protection against virus-induced CPE, based on MTS cell viability assay or microscopic scoring. A/H1N1 virus strains: A/PR/8/34 (PR8) and A/Virginia/ATCC3/2009 (Virg09). <sup>b</sup>CC<sub>50</sub>: 50% cytotoxic concentration based on the MTS assay. <sup>c</sup>MCC: minimum cytotoxic concentration, i.e. compound concentration causing minimal changes in cell morphology. <sup>d</sup>BXA: baloxavir acid; concentrations expressed in nM. Compounds **24–34**, **36**, **40–44**, **56–65** were inactive (EC<sub>50</sub> > 100 μM) against the PR8 and Virg09 strains. All the compounds were inactive against A/H3N2 and influenza B virus. Data for **VF-57a** (compound **23**) are highlighted in bold.

**Scheme 1.**
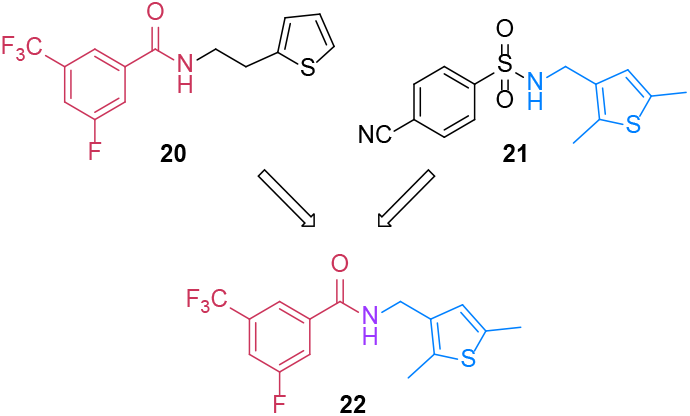
Structures of anti-IAV compounds **20** and **21** and newly designed hybrid **22**.

**Chart 2.**
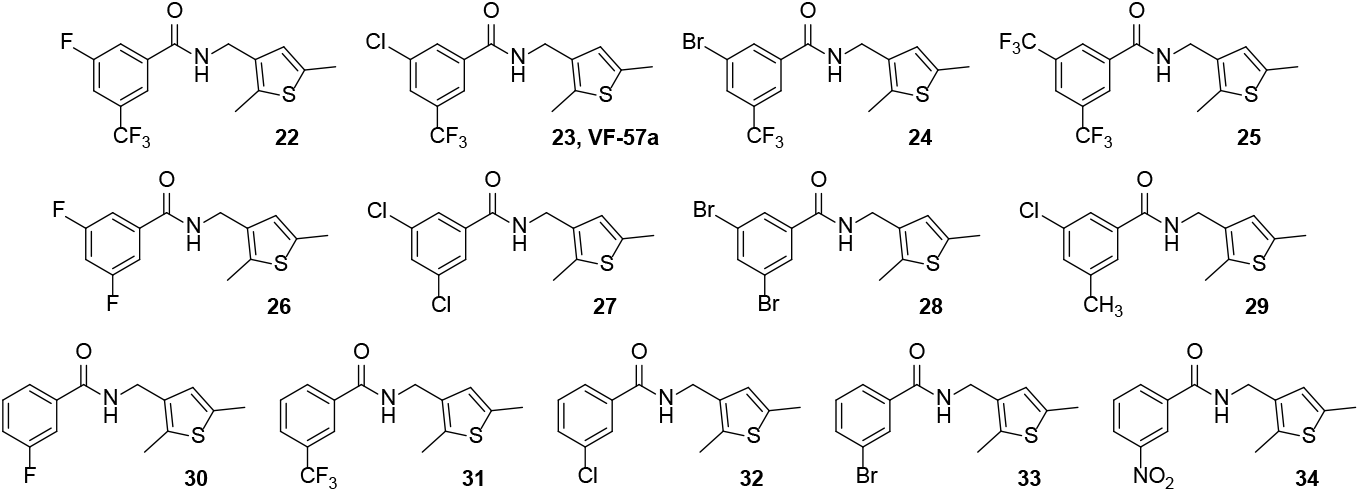
Structures of benzamides with a mono- or di-substituted phenyl ring.

**Chart 3.**
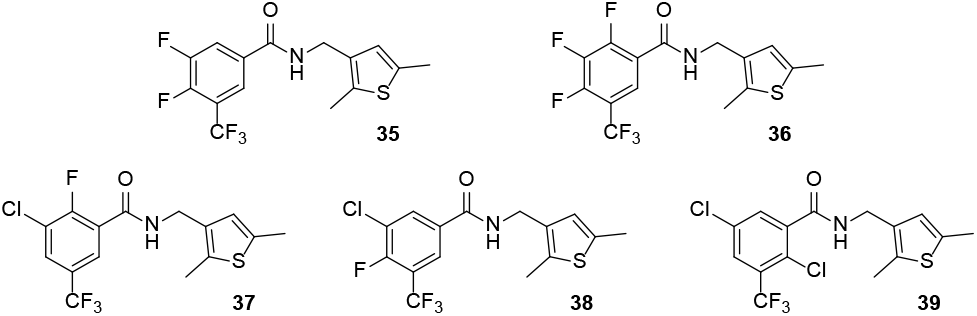
Structures of benzamides with tri- and tetra-substituted phenyl ring.

**Chart 4.**
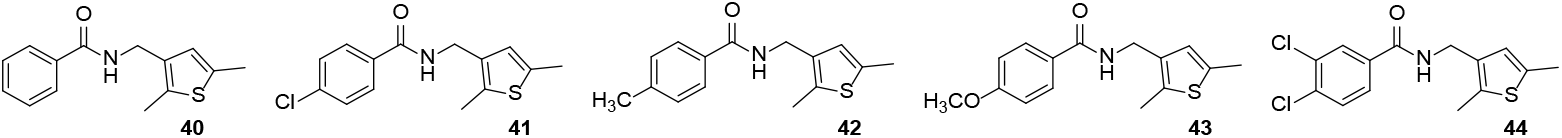
Structures of benzamides explored for a Topliss approach of the benzene ring.

**Chart 5.**
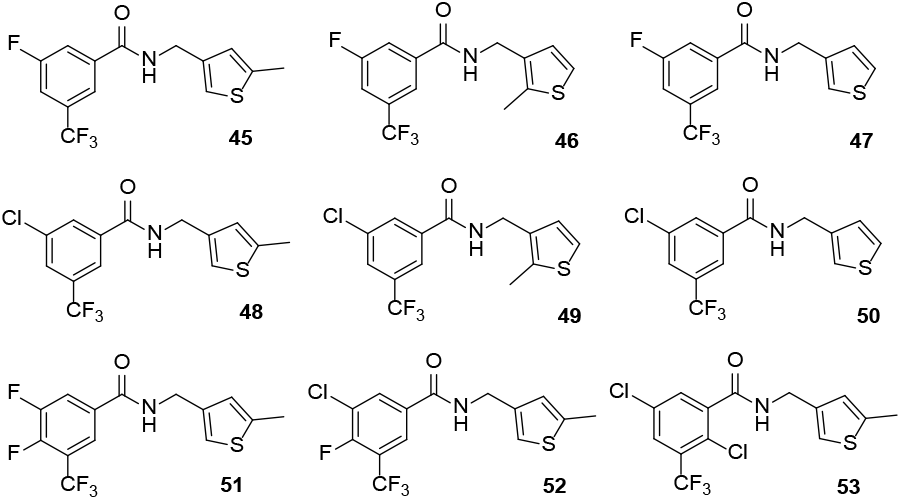
Structures of benzamides synthesized for exploring the influence of the thiophene methyl groups.

**Chart 6.**
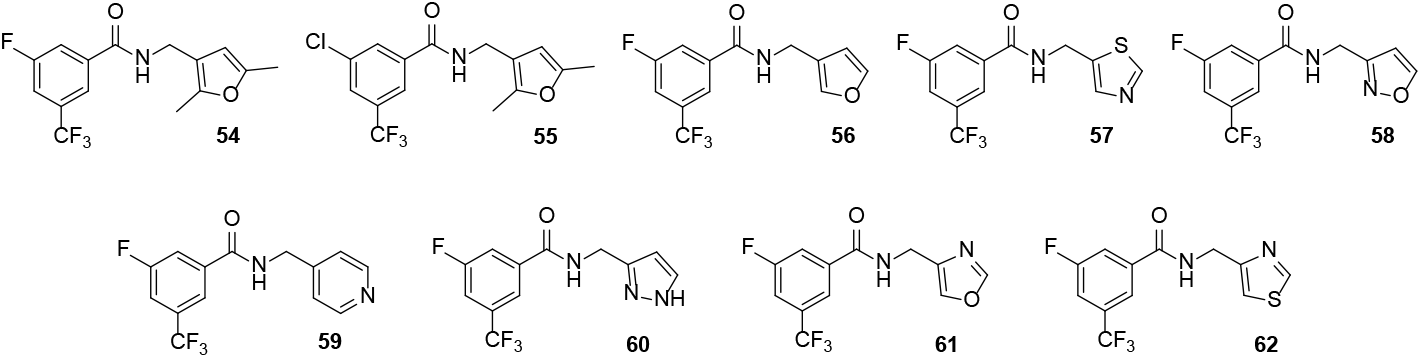
Structures of benzamides featuring different heterocycles on the right-hand side of the molecule.

**Chart 7.**
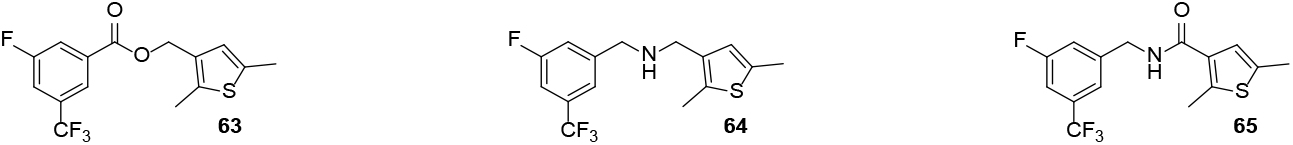
Structures of benzamides with alternative bridges between the phenyl and thiophene rings.

The starting benzamide, **22**, had antiviral EC_50_ values of ∼1 µM for the PR8 and Virg09 virus, without showing any cytotoxicity at 100 µM (the highest concentration tested; Table 1), thus yielding a selectivity index (= ratio of CC_50_ to EC_50_, SI) of >100. Interestingly, while substituting the fluorine atom of **22** by chlorine, as in **23**, led to a slightly more potent compound (Table 1), its replacement by bromine (**24**) or trifluoromethyl (**25**) abolished the antiviral activity. Three analogs featuring the same halogen atom in both meta positions, **26**-**28**, were also inactive. No activity was also noted for compound **29**, that bears a methyl group instead of the trifluoromethyl group of **23**. Finally, five monosubstituted benzamides, **30**−**34**, were inactive, showing that the antiviral activity requires the presence of two electron-withdrawing groups in the *meta* positions (i.e., R_2_ and R_4_ in Table 1; see also Chart 2).

Next, from active benzamides **22** and **23**, the incorporation of one or two additional halogen atoms on the benzene ring was explored. The introduction of a fluorine atom at the *para* position of **22**, leading to **35**, slightly increased the activity. However, the introduction of a further fluorine atom at the *ortho* position, as in **36**, abolished the antiviral effect. Regarding **23**, the effect of introducing additional halogen atoms paralleled those seen with the derivatives of **22**, with compound **38** showing good antiviral activity and the *ortho*-substituted derivatives **37** and **39** being less active.

Relying on the Topliss batchwise scheme (TBS)^49, 50^ to optimize the substitution pattern of a phenyl ring, compounds **40**−**44** were synthesized and tested, but they were inactive, thus preventing further TBS-guided design (Chart 4).

Regarding the right-hand side of the molecule, a few analogues of the more potent compounds, **22, 23, 35** and **38**, were synthesized to explore the effect of the methyl groups (Chart 5). Compounds obtained upon deletion of one or both methyl groups (i.e., **45**-**53**) generally showed antiviral activity, although deletion of the 2-methyl group (i.e., R_6_ in Table 1) seemed to be more relevant relative to the 5-methyl group (i.e., R_7_) for the antiviral activity, particularly against the Virg09 virus. Furthermore, except for the dimethylated furan derivatives **54** and **55**, replacements of the thiophene ring by more polar heterocycles led to loss of activity (Chart 6 and Table 1), suggesting that the polarity of the ring is detrimental for the antiviral activity.

Finally, the importance of the benzamide bridge of **22** was assessed. Neither the ester **63**, nor the secondary amine **64** showed antiviral activity. Finally, also the inverse amide **65** was fully inactive (Chart 7).

### Inhibition of H1 and H5 HA-mediated membrane fusion

Based on the EC_50_ and SI values, compound **23**, dubbed **VF-57a**, was selected for mechanistic investigations. This lead compound had an EC_50_ value of 1 µM and 0.5 µM for the PR8 and Virg09 virus, respectively, and SI > 100 (Table 1 and Figure 2A). For comparison, RL-007 (Chart 1), an aniline-based IAV fusion inhibitor included as a reference compound,^48^ had an EC_50_ of 0.6−1.5 µM and CC_50_ of 33 µM (Table 1), with an SI of ∼31. Besides this activity in MDCK cells, **VF-57a** and RL-007 showed strong anti-A/H1N1 activity in human lung tissue-derived Calu-3 cells, the EC_50_ values being 0.84 µM, 0.53 µM and 1.1 nM for **VF-57a**, RL-007 and BXA, respectively (Fig. 2B).

**Figure 2.**
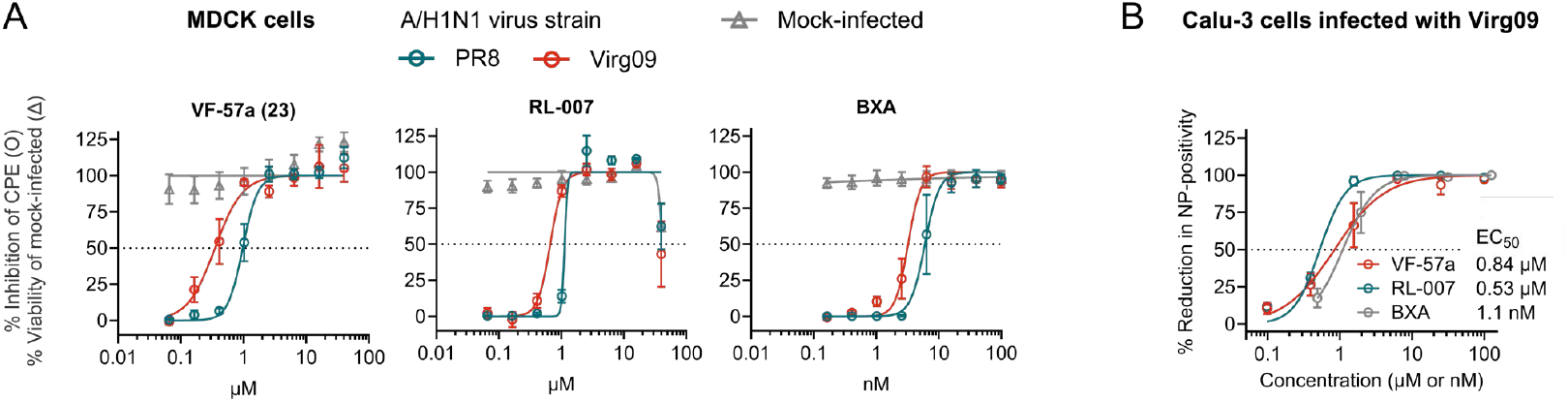
Dose-response curves for antiviral (A/H1N1) activity of VF-57a (23) and RL-007. (A) Assay in MDCK cells: inhibition of CPE (o) and viability of mock-infected cells (Δ), both based on the MTS assay. (B) Calu-3 cells infected with strain Virg09 and stained for viral nucleoprotein (NP). Data points are the mean values ± SEM (n =3); curve fitting with GraphPad Prism 10.2.2 software.

Since the subtype-dependent antiviral activity aligned with HA being the target of the *N*-[(thiophene-3-yl)methyl]benzamides, we determined the inhibitory effect of **VF-57a** in a luciferase-based pseudovirus entry assay in MDCK cells. This method enabled us to include, at BSL2 level, MLV-based pseudovirus bearing the HA and NA of highly pathogenic A/H5N1 virus, specifically strain FL22 (A/bald eagle/FL/W22-114/2022),^51^ which belongs to clade 2.3.4.4b alike the A/H5N1 viruses that are causing the current outbreaks in cattle in the USA.^4^ **VF-57a** proved to be a strong inhibitor of A/H1N1- and A/H5N1-pseudo-virus entry (Figure 3A), with EC_50_ values of 0.30 and 0.81 µM, respectively. On the other hand, RL-007 had almost 127-fold higher activity against A/H1N1 than A/H5N1 (EC_50_: 0.079 and 10 µM, respectively). Arbidol, included as reference compound, had a comparable EC_50_ for A/H1N1 and A/H5N1, but ∼20-fold lower potency than **VF-57a**.

**Figure 3.**
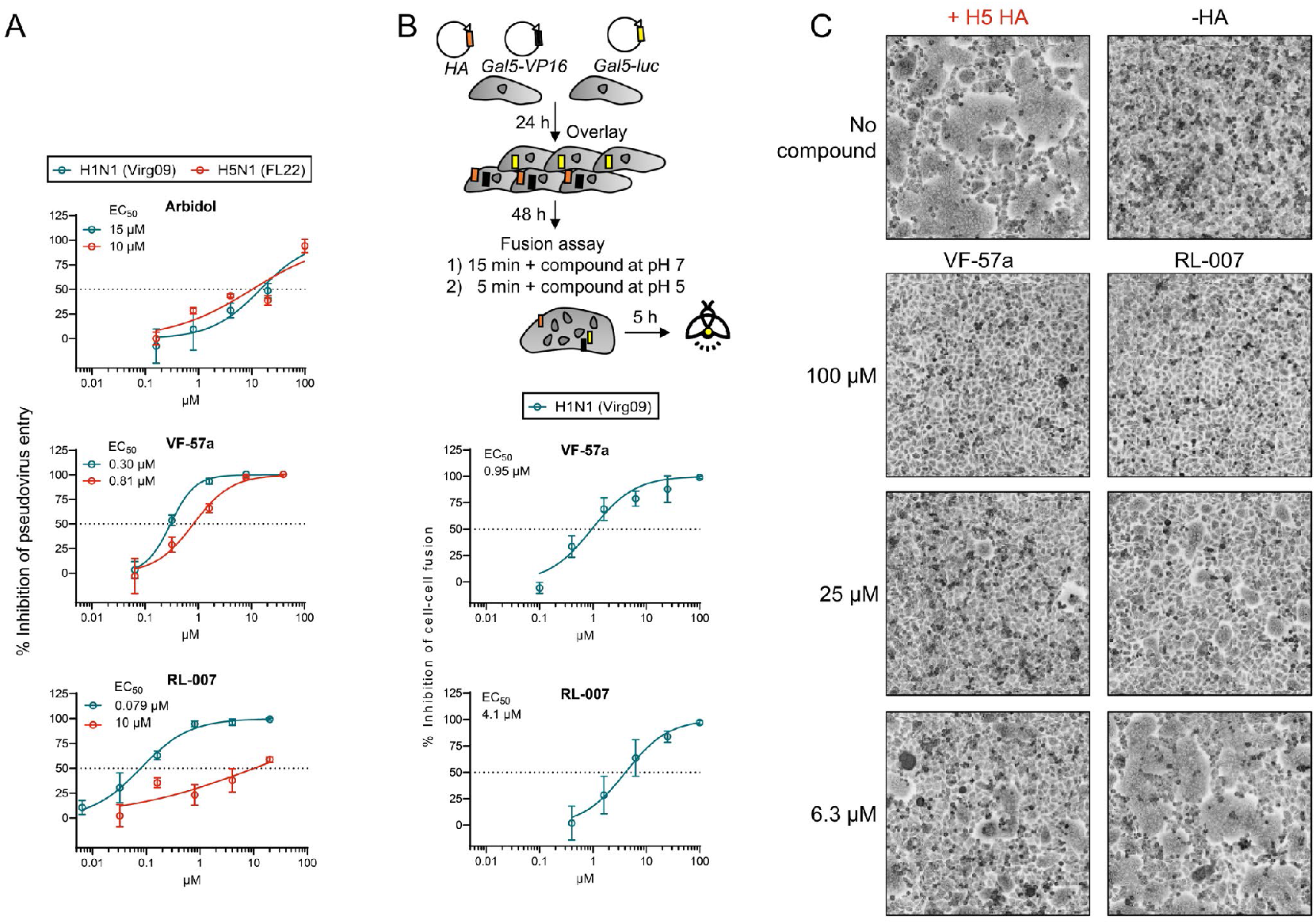
Dose-response curves for inhibition of HA-mediated pseudovirus entry and cell-cell fusion. (A) MDCK cells were transduced with luciferase-expressing A/H1N1- and A/H5N1-pseudoviruses, in the presence of the compounds. Three days later, luminescence was measured to assess the compounds’ inhibitory effect on pseudovirus entry. (B) Cell-cell fusion assay in H1 HA-expressing HeLa cells. The scheme shows the assay setup. The compounds were present during the 15-min preincubation and 5-min acidic stage. Luminescence was measured 5 h after inducing cell-cell fusion. Data points are the mean ± SEM of three independent experiments. (C) Microscopic images of HeLa cells transfected with H5 HA and briefly exposed to pH 5.3. The compounds were present during the 15-min preincubation and 5-min acidic stage. In panels A and B, curve fitting was done with GraphPad Prism 10.2.2 software.

More direct evidence that **VF-57a** acts on membrane fusion was obtained via cell-cell fusion assays in HA-expressing HeLa cells exposed to pH∼5. **VF-57a** and RL-007 were pre-incubated with the HeLa cells for 15 min, then further present during the 5-min acidic stage. Using a luciferase-based readout (see Figure 3B for an assay scheme), we determined an EC_50_ value for H1 HA of 0.95 µM for **VF-57a** and 4.1 µM for RL-007. Both compounds proved also effective against H5 HA, giving complete inhibition of cell-cell fusion at 100 µM (see microscopic pictures in Figure 3C). At lower concentrations, **VF-57a** still gave almost complete (25 µM) or partial (6.3 µM) inhibition, while RL-007 was partially effective at 25 µM and inactive at 6.3 µM.

Overall, the results obtained from pseudovirus entry and cell-cell fusion assays validate **VF-57a** as a strong fusion inhibitor of H1 and H5 HAs. In contrast, RL-007 showed stronger inhibition towards H1 than H5 HA.

Finally, we conducted SPR analysis with two anti-HA antibodies to evaluate the compounds’ direct effect at preventing the conformational change of HA at low pH. Whereas the anti-HA stem antibody C179 only binds when the protein is in its prefusion conformation,^52,53^ the anti-head antibody 7B2-32 binds to both pre- and post-fusion HA. Indeed, recombinant H1 HA protein lost the ability to bind to the C179-coated sensor chip, after incubation in an acidic buffer of pH 5.2, which induces the post-fusion structure, (Figure 4). Its binding to the 7B2-32 antibody was not affected by the acidic treatment. When **VF-57a** or RL-007 was present during the acidic incubation stage, the binding interaction between HA and antibody C179 remained significantly higher (*p*<0.001 for comparison to the DMSO control). This proves that the two molecules stabilize the H1 HA protein in its prefusion structure. Arbidol, on the other hand, only slightly protected against the loss of C179 binding, and its effect was not significant. This is consistent with our pseudovirus entry data, where arbidol was much less effective than **VF-57a** and RL-007.

**Figure 4.**
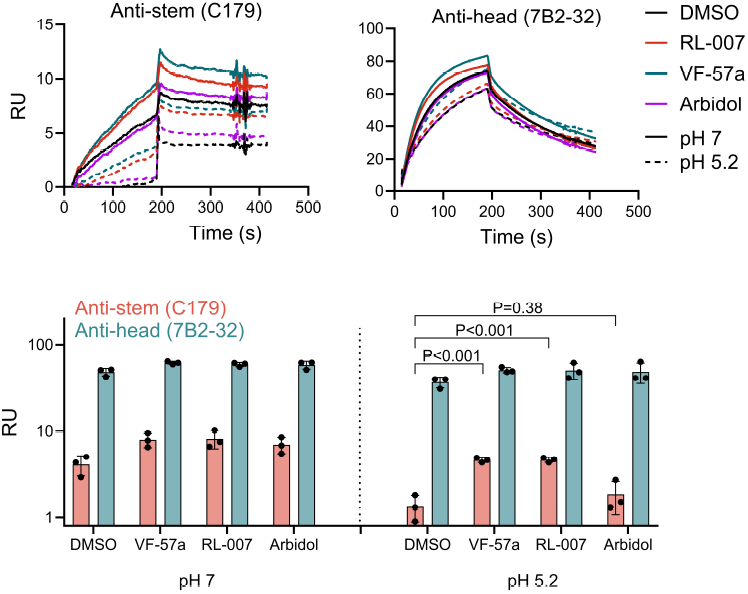
VF-57a and RL-007 inhibit HA refolding at pH 5.2. SPR was used to measure the binding of H1 HA to stem-directed antibody C179 and head-directed antibody 7B2-32. H1 HA was pre-incubated with DMSO or compound, then exposed to neutral or low pH, prior to SPR analysis. (A) Sensorgrams of a representative experiment. (B) Average binding response of three biological replicates. Statistical significance was analysed by a two-sided unpaired t-test. Isotype controls yielded RU values lower than 1 and are therefore not shown. RU: resonance units.

### Phenotypic characterization of VF-57a-resistance markers in H1 HA protein

To understand the mechanism of **VF-57a** vis-à-vis RL-007, we selected resistant viruses via serial passaging of Virg09 virus under increasing concentrations of these inhibitors. After three passages, breakthrough viruses were plaque-purified. In CPE reduction assays, all the virus clones, selected under either **VF-57a** or RL-007, proved at least 60-fold resistant to both compounds (EC_50_ >40 µM; Table 2), indicating cross-resistance between these two inhibitors. Baloxavir, included as control, was equally active against all viruses tested.

**Table 2.** Impact of the HA mutations on viral sensitivity to VF-57a or RL-007 and on acid-lability of HA.

| Selecting compound<sup>a</sup> and HA mutation<sup>b</sup> vs parent virus | Antiviral EC<sub>50</sub> by MTS assay<sup>c</sup> | | | Hemolysis pH ($p$ vs control)<sup>d</sup> |
| --- | --- | --- | --- | --- |
| | VF-57a (μM) | RL-007 (μM) | BXA (nM) | |
| VF-57a: F1181L | >40 | >40 | 2.5 ± 0.5 | 5.25 ($p=0.81$) |
| VF-57a: T3011I | >40 | >40 | 1.9 ± 0.1 | 5.48 ($p<0.0001$) |
| VF-57a: S542P | >40 | >40 | 1.7 ± 0.8 | 5.30 ($p=0.030$) |
| RL-007: A81/821V | >40 | >40 | 3.0 ± 0.7 | 5.10 ($p<0.0001$) |
| RL-007: D902G | >40 | >40 | 1.7 ± 0.2 | 5.41 ($p=0.030$) |
| No compound control: none | 2.5 ± 1.0 | 0.94 ± 0.4 | 2.6 ± 0.7 | 5.24 |

<sup>a</sup>Virg09 virus was passed three times under **VF-57a** and RL-007 (at successively 1.6, 4.0 and 10 μM for **VF-57a** and at 0.64, 1.6 and 4.0 μM for RL-007); a control was included that was passed in the absence of a compound. <sup>b</sup>Residues numbered according to the H3 numbering system (see Supporting Information Figure S1 for an alignment of HA subtypes used in our study). <sup>c</sup>Antiviral EC<sub>50</sub> value in MDCK cells, using the MTS-based CPE assay. Mean ± SEM of three independent experiments. <sup>d</sup>The hemolysis pH was defined as the pH where 50% of virus-induced hemolysis occurred relative to pH 5.0, as based on the pooled data from four independent experiments. Statistical significance of differences between the hemolysis pH of different mutants *versus* the no compound control, were determined with an extra-sum-of-squares F test (GraphPad Prism 10.2.2).

As shown in Figure 5, the Virg09 clones carried amino acid changes at different parts of HA. In a previous resistance study with an H3 HA-specific fusion inhibitor, we found that this class of agents selects two mechanistically distinct types of amino acid substitutions.^39^ While some are directly located at the compound’s binding pocket, other changes are lying at remote sites and associated with higher acid-lability of the HA protein. Such mutant viruses escape from the fusion inhibitor by undergoing fusion in less acidic endosomes. Hence, we used a hemolysis assay to determine which of the mutations, selected under **VF-57a** or RL-007, rendered Virg09 HA more acid-labile. This was the case for mutants T301_1_I and D90_2_G, which showed a hemolysis pH of 5.48 and 5.41, compared to 5.24 for the virus that was passaged without compound and did not acquire any HA mutations during this process (Table 2). Mutant S54_2_P showed a hemolysis pH of 5.30, which is a slight increase compared to the control (*p*=0.030). The F118_1_L virus had the same hemolysis pH as the control, while mutation A81/82_1_V reduced the acid-lability (pH 5.10; *p*<0.0001). This suggests that these sites may lie close to the binding pocket of both **VF-57a** and RL-007.

**Figure 5.**
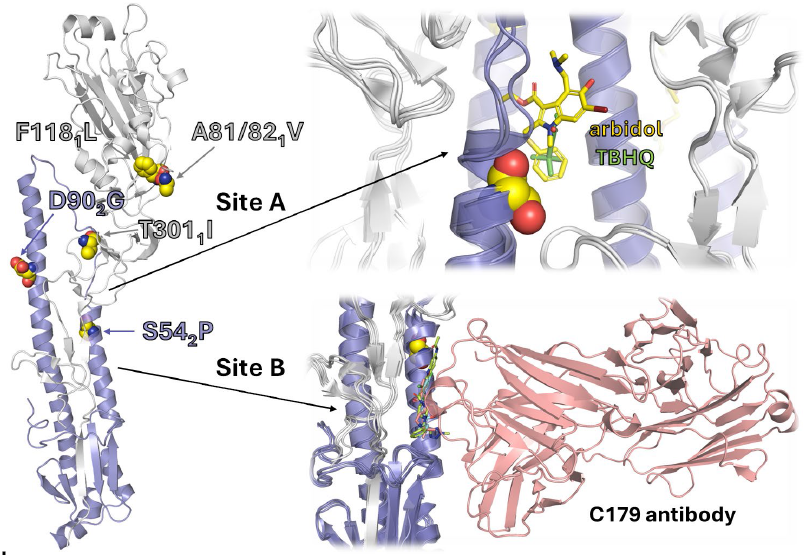
Location of the HA mutations that Virg09 virus acquired when passaged under VF-57a or RL-007. The left panel shows the X-ray structure of a Virg09-related H1 HA (PDB ID 3M6S);^52^ the suffix refers to the HA1 (grey) or HA2 (lavender) chains. Zoom in the top panel: view of the binding site of TBHQ (PDB ID 3EYM)^31^ and arbidol (PDB ID 5T6N)^14^ in group 2 (i.e. H3) HA. Zoom in the bottom panel: binding site of antibody C179 (PDB ID 5C0R)^53^ and the group 1-specific inhibitors JNJ4796, (*S*)-F0045 and CBS1117 (PDB IDs: 6CF7,^25^ 6WCR^32^ and 6WMZ,^33^ respectively).

Molecular Dynamics simulations to explore the binding site of VF-57a in the HA stem

#### Binding of (S)-F0045 and VF-57a to H1 HA Site B

The reported X-ray structures of HA in complex with a fusion inhibitor show the existence of two binding sites (Figure 1), here denoted A (targeted by TBHQ^31^ and arbidol^14^) and B (targeted by JNJ4796,^25^ (*S*)-F0045^32^ and CBS1117)^33^. Accordingly, MD simulations were used to explore the potential binding of **VF-57a** to these sites (for details of the system setup and Molecular Dynamics simulations, see Experimental Section. Molecular Modeling). To this end, the computational procedure was validated by performing additional MD simulations for (*S*)-F0045 and arbidol as reference compounds (see below). Taking advantage of the trimeric nature of HA, the ligands (**VF-57a**, (*S*)-F0045 and arbidol) were positioned in the three pockets of HA, thus enabling to assess the stability of the bound ligand in triplicate in a single trajectory. Note that we use the H3 amino acid numbering system (see Supporting Information Figure S1 for an alignment of the relevant HA subtypes).

Inspection of the X-ray structure of (*S*)-F0045 in complex with H1 HA (PDB ID 6WCR)^322^ shows that the ligand fills a groove (Site B), shaped by residues H18_1_, W21_1_, H38_1_, V40_1_, T318_1_, D19_2_, T41_2_, I45_2_, T49_2_ and V52_2_. Our MD simulations confirmed the stability of the binding mode of (*S*)-F0045, as noted in root-mean square deviation (RMSD) values of 1.3 ± 0.3, 1.3 ± 0.4 and 1.3 ± 0.3 Å for the three bound ligands (Figure 6). Remarkably, the hydrogen bond between the amide carbonyl oxygen of (*S*)-F0045 and the hydroxyl group of T318_1_ was maintained along the trajectory in all cases (average distance of 2.8 ± 0.1 Å for the three B sites, see Supporting Information Figure S2 for the plot of distance against simulation time).

**Figure 6.**
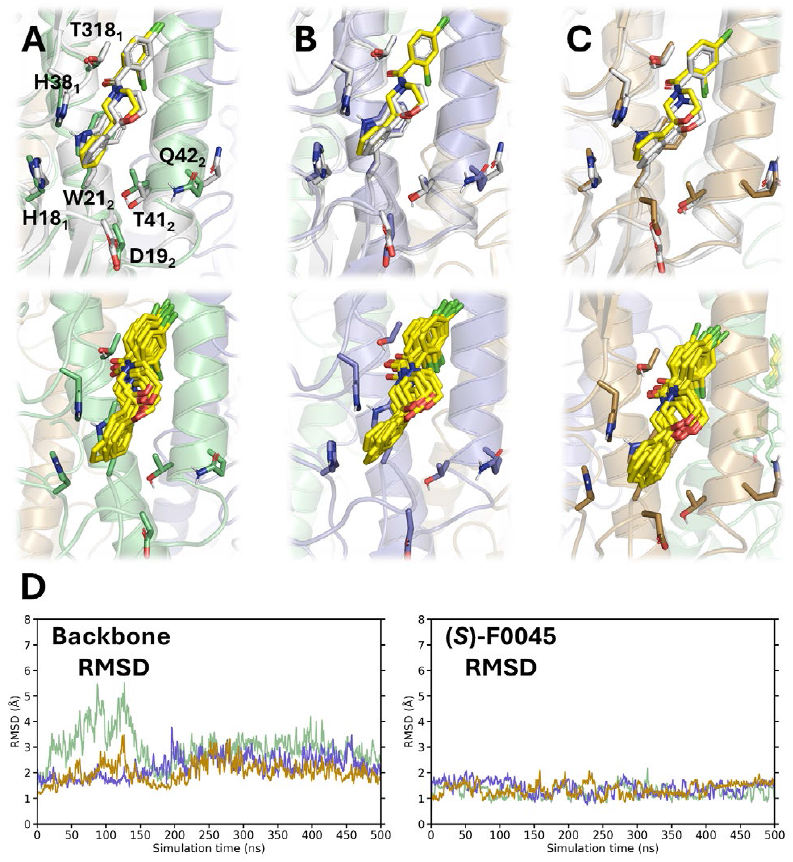
Binding mode of (*S*)-F0045 to Site B of H1 HA. (A–C) Top panels: Overlay of (*S*)-F0045 at the end of the MD simulation (yellow sticks) and its crystallographic pose (grey sticks; PDB ID 6WCR)^32^ in each of the three HA (Virg09) protomers (shown in green, lavender and beige). Bottom panels: Superposition of 10 snapshots taken along the last 100 ns of the trajectory. (D) RMSD plots for each HA protomer’s backbone and for each of the three ligands (shown in green, lavender and beige).

The potential binding of **VF-57a** to Site B was examined for two distinct binding modes. These maintain the hydrogen bond between the amide carbonyl oxygen of the ligand and the hydroxyl group of T318_1_ but differ in the relative arrangement of the phenyl and thiophene rings along the groove (see Supporting Information Figure S3). This leads to two distinct overlaps of the molecular skeleton of **VF-57a** and (*S*)-F0045, where either the phenyl or thiophene rings overlap the dichlorobenzene unit of (*S*)-F0045.

In contrast to the behaviour of (*S*)-F0045, neither of the two **VF-57a** binding modes to H1 HA Site B were stable (Figure 7). The average RMSD values ranged from 5.5 ± 2.6 Å to 9.0 ± 3.1 Å for the six ligands (= two poses × three sites). The unstable binding was also reflected in the loss of the hydrogen bond with T318_1_, since the average distance ranged from 5.2 ± 1.8 to 7.0 ± 3.8 Å (see Supporting Information Figure S4 for the plot of distance against simulation time). The instability of **VF-57a** vis-à-vis (*S*)-F0045 may be related to the increased flexibility caused by the methylene unit between the amide and thiophene moieties. In addition, due to the presence of the trifluoromethyl and methyl groups in the phenyl and thiophene rings, respectively, **VF-57a** cannot engage in the CH-π interactions which (*S*)-F0045 forms with H18_1_, H38_1_ and W21_2_.

**Figure 7.**
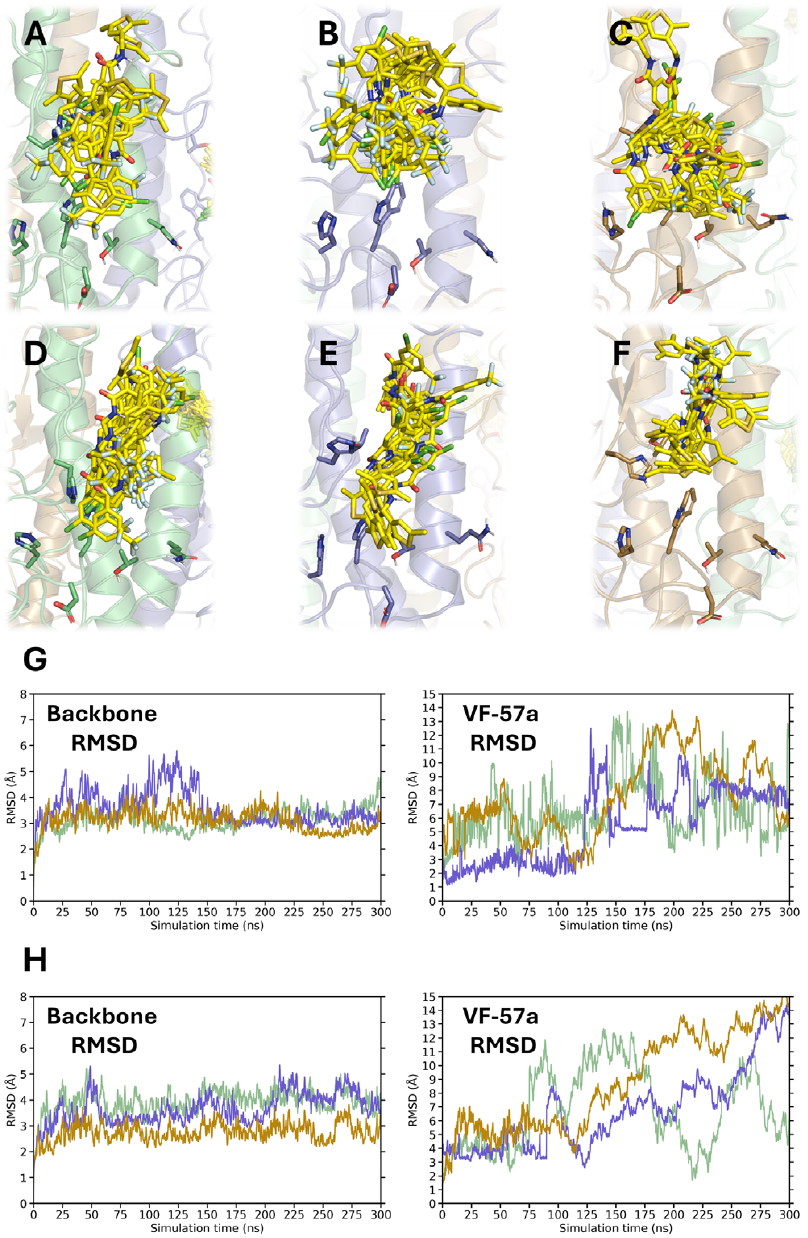
Binding mode of VF-57a to Site B of H1 HA. (A– F) Superposition of 20 snapshots (**VF-57a** shown as yellow sticks) taken every 5 ns along the last 100 ns of the trajectory in each of the three HA (Virg09) protomers (shown in green, lavender and beige) for the two distinct alignments of **VF-57a** (A–C and D–F) along the groove. (G, H) RMSD plots for each HA protomer’s backbone and for each of the three ligands (shown in green, lavender and beige) in the orientation shown in panels (G) A–C and (H) D–F.

Overall, these results suggest that Site B is not the binding pocket of **VF-57a** in H1 HA. This conclusion agrees with our SPR results (see above), since the binding of **VF-57a** to HA protein (exposed to pH 7) had no effect on the subsequent binding of antibody C179 which, alike (*S*)-F0045, interacts with Site B (Figure 5).^53^

#### Binding of arbidol and VF-57a to HA site A

In the X-ray structure of H3 HA in complex with arbidol, the binding pocket (Site A) is shaped by residues of two HA monomers. Monomer 1 contributes with P293_1_, F294_1_, K307_1_ (R in A/Hong Kong/7/87) and R54_2_, E57_2_ (G in A/Hong Kong/7/87), K58_2_, N60_2_, W92_2_ and E103_2_, whereas monomer 2 contributes with D90_1_, A101_2_, K310_1_ and I29_1_.^14^

MD simulations for the H3 HA complex with arbidol revealed a stable behavior of the protein backbone during the entire simulation, with average RMSD values of 2.2 ± 0.3, 2.5 ± 0.4 and 2.3 ± 0.3 Å for each protomer (Figure 8). The crystallographic pose of arbidol is maintained in two pockets, as noted in average RMSD values of 2.4 ± 0.6 Å and 3.0 ± 0.5 Å. The tertiary amine moiety of arbidol forms transient hydrogen bonds with E57_2_ and the backbone carbonyl oxygen of K58_2_. In the third pocket, the RMSD profile (green line in Figure 8D) increased after 80-90 ns and achieved a stable value of 5.3 ± 1.8 Å after 210 ns. This reflects a rearrangement of the molecule within its binding pocket, where the indole ring flips by 180° upside down and the thiophenyl group ends more buried in the cavity (panel A in Figure 8).

**Figure 8.**
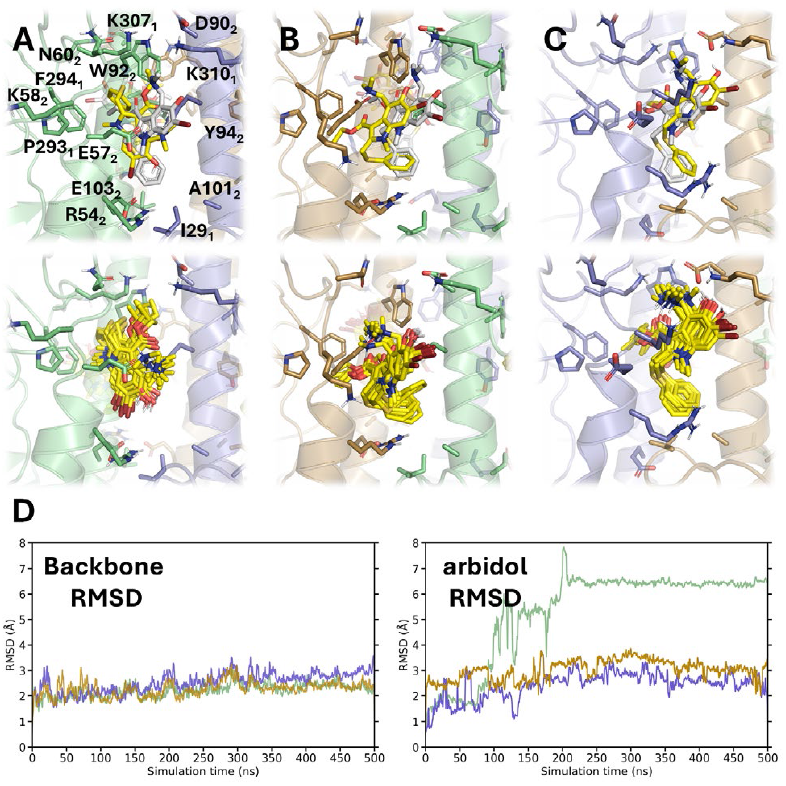
Binding mode of arbidol to Site A of H3 HA. (A– C) Top panels: Overlay of arbidol at the end of the MD simulation (yellow sticks) and its crystallographic pose (grey sticks) in Site A of H3 HA (PDB ID 5T6N).^14^ Bottom panels: Superposition of 10 snapshots taken along the last 100 ns of the trajectory. (D) RMSD plots for each HA protomer’s backbone atoms and for each of the three ligands (shown in green, lavender and beige).

Our results align with the conclusion that arbidol acts as a ‘molecular glue’ for trimeric HA, by engaging in hydrophobic interactions and inducing the formation of salt bridges, such as the contact between the carboxylate group of D90_2_ and the amine nitrogen of K310_1_ (average N^…^O distance of 4.1 ± 1.4 Å), and the interactions of the guanidinium group of R54_2_ with either E97_2_ or E103_2_. This underscores the structural plasticity of Site A to accommodate arbidol and presumably other ligands.

Considering that **VF-57a** is active against A/H1N1 but not A/H3N2 virus (see Table 1), we explored the binding of **VF-57a** to Site A in H1 HA (Virg09). To this end, the last helical turn of the short α-helix in HA2 was unfolded to enable the binding of a ligand, thus simulating the local structure observed in the H3 HA-arbidol complex (PDB ID 5T6N),^14^ (see Experimental Section. Molecular modeling: Homology modelling and setup of simulated systems and Supporting Information Figure S5). MD simulations were run for the complex between HA and **VF-57a** (stoichiometric ratio 1:3), thus enabling the comparison of the binding pose attained by three ligands.

The modelled HA trimer was stable with average RMSD values ranging from 2.8 ± 0.6 to 3.0 ± 0.6 Å for the three HA protomers (Figure 9). The overall structural stability of the HA trimer was confirmed by the small RMSD values determined for the stem helices for each protomer, as noted in values ranging from 1.2 ± 0.2 to 1.5 ± 0.2 Å (see Supporting Information Figure S6). The initial pose obtained from docking of **VF-57a** in two pockets was fully preserved along the MD trajectory, as noted in RMSD values of 1.1 ± 0.4 and 1.4 ± 0.5 Å. Regarding the binding of **VF-57a** to the third pocket, the initial docked pose was slightly different due to the adoption of a distinct arrangement of the thiophene ring (see Supporting Information Figure S7). The increase in the RMSD profile observed for the third ligand after the first 100 ns reflects a fast rearrangement of the thiophene ring relative to the original docked pose, which enabled the three ligands to adopt a similar pose at the end of the MD simulation. Compared to arbidol, the binding pose places VF-57a more deeply in the pocket (Figure 9, panel E).

**Figure 9.**
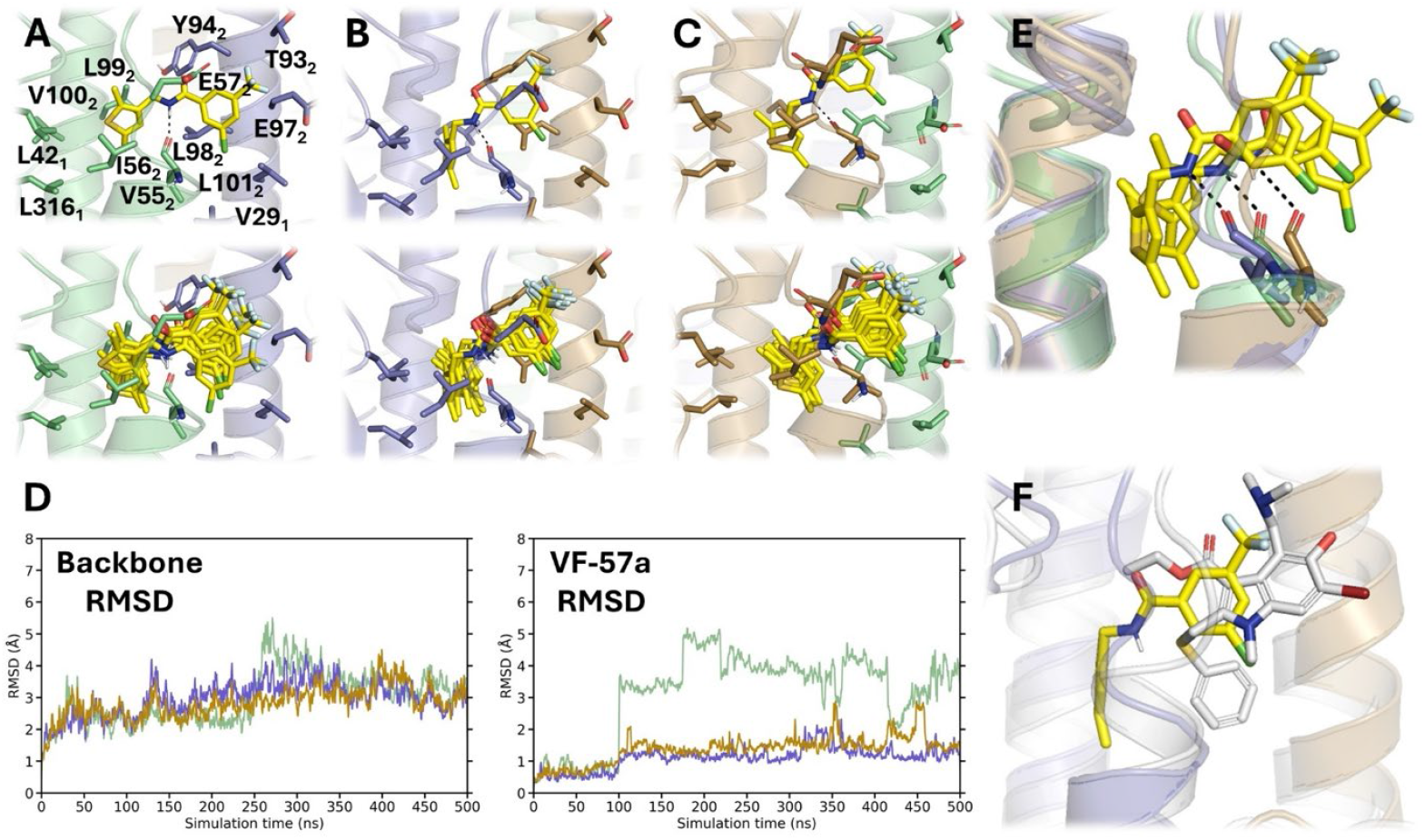
Binding mode of VF-57a to Site A of H1 HA. (A–C) Top panels: Final position of **VF-57a** (yellow sticks) in Site A of the H1 HA (Virg09)-modified homology model. Bottom panels: Superposition of 10 snapshots taken along the last 100 ns of the trajectory. (D) RMSD plots for each HA protomer’s backbone and for each of the three ligands (shown in green, lavender and beige). (E) Superposition of the three protomers at the end of the MD simulation, showing the adoption of a common pose for the three **VF-57a** ligands. The dashed line reflects the hydrogen bond interaction formed between the ligand’s amide NH group and the carbonyl oxygen of V55_2_. (F) Detailed view of the final pose of **VF-57a** with arbidol (grey sticks; PDB ID 5T6N)^14^.

In all cases, the thiophene ring is surrounded by hydrophobic residues in the lower part of the pocket, such as L42_1_, L316_1_, I56_2_, L99_2_, and V100_2_ from protomer 1, and L98_2_ from protomer 2 (Figure 9). This suggest that the thiophenyl ring exerts a hydrophobic anchoring in site A. Indeed, the occupancy of this subpocket explains the loss of antiviral activity observed when the thiophenyl unit is replaced by more polar heterocycles (see above), reflecting the penalty due to the desolvation cost upon burial of this chemical moiety in the pocket. This is reflected in the comparison of the dipole moment and octanol/water partition coefficient determined (Table 3) for the heterocyclic rings in compounds **56**-**62** relative to dimethyl thiophene (μ=0.80 D, logP=3.02; **VF-57a**) and dimethyl furan (μ=0.42 D, logP=2.63; compounds **54** and **55**).

**Table 3.** Dipole moment (μ; Debye)^a^ and octanol/water partition coeficient (logP)^b^ determined for the hetero-cyclic moieties in compounds 23 and 54-62.

| Model ring (compound) | $\mu$ | $\log P$ |
| --- | --- | --- |
| 2,3,5-trimethyl thiophene (**23**, **VF-57a**) | 0.80 | 3.02 |
| 2,3,5-trimethyl furan (**54**, **55**) | 0.42 | 2.63 |
| 3-methyl furan (**56**) | 0.91 | 1.70 |
| 2-methyl thiazole (**57**) | 1.85 | 1.24 |
| 3-methyl isoxazole (**58**) | 2.98 | 0.61 |
| 4-methyl pyridine (**59**) | 2.65 | 1.23 |
| 3-methyl pyrazole (**60**) | 1.97 | -0.11 |
| 4-methyl oxazole (**61**) | 1.34 | 0.67 |
| 4-methyl thiazole (**62**) | 1.14 | 1.26 |

<sup>a</sup>Determined in the gas phase at the B3LYP/6-31G(d) level.
<sup>b</sup>Estimated from IEFPCM/MST calculations at the B3LYP/6-31G(d) level.

Besides, the alchemical transformation from **VF-57a** to **50**, which involves the removal of the two methyl groups on the thiophene moiety, predicts that binding of **50** is destabilized by 1.6 kcal/mol relative to **VF-57a** (Table 4), reflecting a decrease in the enthalpic component to the binding. This can be attributed to the loss of stabilizing van der Waals interactions upon removal of the methyl groups and is consistent with the reduced antiviral activity against A/H1N1 Virg09 virus for **50** (EC_50_=1.1 µM) relative to **VF-57a** (EC_50_=0.31 µM; Table 1). Moreover, this trend is reinforced from the results obtained for the alchemical transformations from **VF-57a** to **46** and **47**, which imply the removal of one or both methyl groups from the thiophene moiety as well as the replacement of the Cl atom by F in the trifluorotoluene ring of both compounds. Thus, compounds **46** and **47** are predicted to be destabilized by 1.8 and 1.9 kcal/mol relative to **VF-57a** (Table 4), which are in qualitative agreement with the free energy difference estimated from the EC_50_ values determined experimentally for these compounds (Table 1).

**Table 4.** Relative free energy difference (kcal/mol) determined for selected alchemical transformations between pairs of compounds (see Experimental Section. Relative Binding Free Energy (RBFE) Simulations).

| Change | $\Delta\Delta G$ (TI) | $\Delta\Delta G$ (exp)<sup>a</sup> |
| --- | --- | --- |
| **VF-57a** → **50** | $1.6 \pm 0.1$ | 0.8 |
| **50** → **VF-57a** | $-1.6 \pm 0.1$ | -0.8 |
| **VF-57a** → **22** | $0.7 \pm 0.1$ | 0.2 |
| **22** → **VF-57a** | $-0.7 \pm 0.1$ | -0.2 |
| **VF-57a** → **38** | $-0.1 \pm 0.1$ | -0.3 |
| **38** → **VF-57a** | $0.1 \pm 0.1$ | 0.3 |
| **VF-57a** → **46** | $3.4 \pm 0.1$ | 1.8 |
| **46** → **VF-57a** | $-3.4 \pm 0.1$ | -1.8 |
| **VF-57a** → **47** | $4.3 \pm 0.1$ | 1.9 |
| **47** → **VF-57a** | $-4.3 \pm 0.1$ | -1.9 |

<sup>a</sup>Estimated from the EC<sub>50</sub> values determined from experimental measurements (see Table 1).

Also, the amide NH group of the linker in **VF-57a** forms a hydrogen bond with the backbone carbonyl oxygen of V55_2_, as noted in averaged distances ranging from 3.2 ± 0.5 to 3.9 ± 0.6 Å for the three **VF-57a** molecules bound to HA. This feature may explain the sensitivity of the antiviral activity to the nature of the linker, since replacement of the amide unit by ester (**63**), secondary amine (**64**) and inverse amide (**65**) led to inactive compounds (see above).

Finally, the phenyl ring containing the trifluoromethyl moiety is oriented toward the mouth of the cavity, being partially overlapped with the indole ring of arbidol (Figure 9). Accordingly, the substituted phenyl ring forms van der Waals contacts with the side chains of V55_2_, E56_2_, T93_2_, Y94_2_, E97_2_ and L101_2_. This arrangement justifies the small changes in binding affinity associated with the change of chlorine in **VF-57a** to fluorine in **22**, which is destabilized by 0.7 kcal/mol, and the subsequent attachment of fluorine in *para* position to yield **38**, which would imply a free energy change of 0.1 kcal/mol (Table 4). These values agree with the slight difference in EC_50_ values determined against A/H1N1 Virg09 (values of 0.3 µM for **VF-57a**, 0.4 µM for **22**, and 0.1 µM for **38**; Table 1).

#### Interpretation of the VF-57a binding mode in relation to its resistance profile and group 1-specificity

As explained above, the **VF-57a**-resistance mutations were located at different parts of the HA protein, including the head domain. This heterogenous mutation profile was also seen with other fusion inhibitors, such as arbidol,^14^ M090,^54^ MBX2456,^55^ and RL-007,^48^ and complicates the interpretation of which mutations have direct impact on inhibitor binding, while others may act by modifying the acid-stability of HA.

In the case of **VF-57a**, mutation S54_2_P, which had no significant impact on acid-stability, is located close to the entrance of the Site A binding pocket (Figure 10). Exchanging S54_2_ by P is expected to promote a local structural destabilization, which would affect the conformational flexibility of the unfolded loop that shapes the mouth of Site A. Indeed, this may affect the stabilization exerted by the hydrogen bond between the ligand with V55_2_ (see above and panel A in Figure 10). In this regard, it is worth noting that the S54_2_P mutant also proved resistant against RL-007, confirming our MD-based prediction on the binding of RL-007 to Site A.^48^

**Figure 10.**
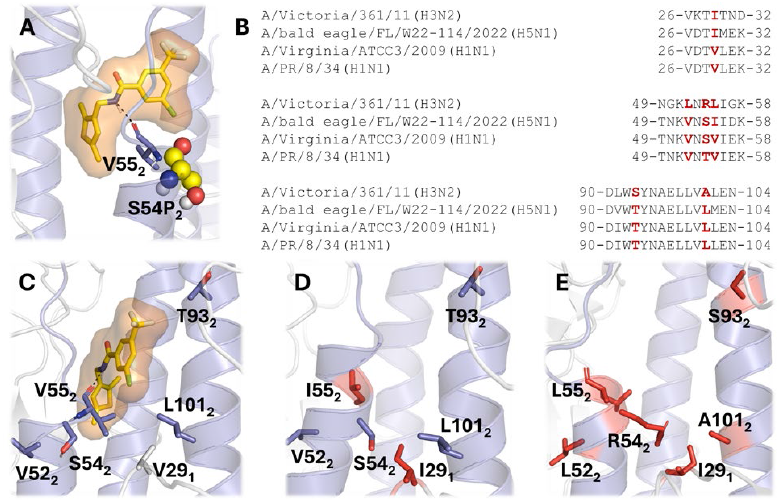
Structural basis for the viral resistance profile and group 1-specificity of VF-57a. (A) Representation of the binding mode of VF-57a together with the hydrogen bond formed with V55_2_ (shown as sticks) and the resistance mutation S54P_2_ (shown as spheres). (B) Comparison of selected sequence regions that define the walls of the binding pocket. (C-D) Representation of the binding pose of **VF-57a** and the location of residue differences (highlighted as red sticks) between H1 HA (C: Virg09); H5 HA (D: FL22); and H3 HA (E: A/Victoria/361/11).

The MD-based model also provides a basis to rationalize why **VF-57a** shows strong inhibitory activity against A/H1N1 and A/H5N1 (pseudo)viruses, but not against A/H3N2 virus. Specifically, comparison of the residues that shape the binding pocket in H1 HA shows only one difference between the PR8 and Virg09 strains, namely the replacement of T54_2_ in PR8 by S54_2_ in Virg09 (Figure 10B and C). The binding pocket is also highly conserved between the H1 (Virg09) and H5 (FL22) HAs, as the only differences concern the conservative substitutions of V29_1_ by I29_1_ and of V55_2_ by I55_2_(Figure 10B and D). In contrast, up to six differences are noticeable between H1 HA (Virg09) and H3 HA (from A/Victoria/361/11), with residues V29_1_, V52_2_, S54_2_, V55_2_, T93_2_ and L101_2_ in the H1 protein being replaced by I, L, R, L, S and A in its H3 counterpart. In particular, the changes S54_2_R, which implies the incorporation of a positive charge, and L101_2_A, which enlarges the size of the pocket, may deteriorate the binding of **VF-57a**, by increasing the structural flexibility due to enhancement of interactions with water molecules (R54_2_) and weakening van der Waals interactions (A101_2_) with the ligand.

## CONCLUSIONS

This study reports the synthesis, antiviral evaluation and mechanism of action of an original series of *N*-[(thiophen-3-yl)methylbenzamides acting as strong inhibitors of group 1 HA-mediated fusion. The lead compound, **VF-57a**, exhibits sub-micromolar activity against authentic A/H1N1 virus and a similarly high potency for inhibiting the cell entry of A/H1N1 and A/H5N1 (i.e. highly pathogenic IAV-derived) pseudoviruses.

The strong inhibitory activity of **VF-57a** is proposedly mediated through its binding to the pocket targeted by the broad IAV HA inhibitor arbidol. Within this pocket, the dimethylthiophene moiety of **VF-57a** anchors the ligand to a hydrophobic pocket, and the amide unit of the linker forms a hydrogen bond with V55_2_. Finally, the substituted phenyl is located at the entrance of the pocket, partly overlapping the indole ring of arbidol. Being validated by our resistance data and thorough SAR of this compound series, our study offers a solid basis for further structure-based development of highly active IAV fusion inhibitors. In addition, further follow-up of **VF-57a** for anti-influenza drug development is envisaged, including its validation in *in vivo* models.

## EXPERIMENTAL SECTION

### Chemical synthesis

Commercially available reagents and solvents were used without further purification unless stated otherwise. Preparative normal phase chromatography was performed on a CombiFlash Rf 150 (Teledyne Isco) with pre-packed RediSep Rf silica gel cartridges. Thin-layer chromatography was performed with aluminum-backed sheets with silica gel 60 F254 (Merck, ref 1.05554), and spots were visualized with UV light and 1% aqueous solution of KMnO_4_. All compounds showed a sharp melting point and a single spot on TLC. Melting points were determined in open capillary tubes with a MFB 595010M Gallenkamp. 400 MHz ^1^H and 100.6 MHz ^13^C NMR spectra were recorded on a Varian Mercury 400 or on a Bruker 400 Avance III spectrometers. The chemical shifts are reported in ppm (δ scale) relative to internal tetramethylsilane, and coupling constants are reported in Hertz (Hz). Assignments given for the NMR spectra of selected new compounds have been carried out on the basis of DEPT, COSY ^1^H/^1^H (standard procedures), and COSY ^1^H/^13^C (gHSQC and gHMBC sequences) experiments. IR spectra were run on Perkin-Elmer Spectrum Two FT-IR spectrophotometer. Absorption values are expressed as wave-numbers (cm^−1^); only significant absorption bands are given. High-resolution mass spectrometry (HRMS) analyses were performed with an LC/MSD TOF Agilent Technologies spectrometer. The elemental analyses were carried out in a Flash 1112 series Thermofinnigan elemental microanalyzator (A5) to determine C, H and N. The structure of all new compounds was confirmed by elemental analysis and/or accurate mass measurement, IR, ^1^H NMR and ^13^C NMR. The analytical samples of all the new compounds, which were subjected to pharmacological evaluation, possessed purity ≥95% as evidenced by their elemental analyses (Table S1) or HPLC/UV. HPLC/UV were determined with a HPLC Agilent 1260 Infinity II LC/MSD coupled to a photodiode array. 5 µL of sample 0.5 mg/mL in methanol:acetonitrile were injected, using an Agilent Poroshell 120 EC-C18, 2.7 µm, 50 mm x 4.6 mm column at 40 °C. The mobile phase was a mixture of A = water with 0.05% formic acid and B = acetonitrile with 0.05% formic acid, with the method described as follows: flow 0.6 mL/min, 5% B-95% A 3 min, 100% B 4 min, 95% B-5% A 1 min. Purity is given as % of absorbance at 220 nm.

### General Procedure A

To a solution of (2,5-dimethylthiophen-3-yl)methanamine (1 equiv.) in anh. dichlorometane (3 mL), triethylamine was added (1.2 equiv.). Then, the solution was cooled to 0 °C and a solution of the required commercial acyl chloride (1.1 equiv.) in anh. dichloromethane (2 mL) was slowly added dropwise. The mixture was kept under stirring at room temperature for 5 h. Then the reaction mixture was concentrated *in vacuo*. Column chromatography (hexane/ethyl acetate mixtures from 0 to 10%) gave the expected product.

### General Procedure B

To a solution of the commercially available carboxylic acid (1 equiv.) in dry acetonitrile (5 mL), thionyl chloride (5 equiv.) was added dropwise. The mixture was heated under reflux for 2h. Then, the solvent was evaporated *in vacuo* to yield the crude product, which was used directly in the next step.

To a solution of (2,5-dimethylthiophen-3-yl)methanamine (1 equiv.) in anh. dichlorometane (3 mL), triethylamine was added (1.2 equiv.). Then, the solution was cooled to 0 °C and a solution of the required commercial acyl chloride (1.1 equiv.) in anh. dichloromethane (2 mL) was slowly added dropwise. The mixture was kept under stirring at room temperature for 5 h. Then the reaction mixture was concentrated *in vacuo*. Column chromatography in silica gel (using as eluent mixtures of EtOAc in hexane from 0 to 10%) gave the expected product.

### General Procedure C

To a solution of the commercially available carboxylic acid in anh. toluene, thionyl chloride (5 equiv.) followed by some drops of DMF were added dropwise. The mixture was heated under reflux for 2h. Then, the solvent was evaporated *in vacuo* to yield the crude product, which was used directly in the next step.

To a suspension of the required amine and triethylamine in anh. DCM, a solution of the proper acyl chloride in anh. DCM was slowly added. The mixture was kept under stirring at room temperature for 5 h. Then, water (15 mL) followed by EtOAc (15 mL) were added and the mixture was extracted. The organic layer was washed with brine (15 mL), dried over anh. Na_2_SO_4_ and filtered. Solvents were concentrated *in vacuo* and the resulting crude was purified by column chromatography in silica gel (using as eluent mixtures of EtOAc in hexane from 0% to 10%) to obtain the desired product.

### General Procedure D

To a solution of the commercially available carboxylic acid in anh. toluene, thionyl chloride (5 equiv.) followed by some drops of DMF were added dropwise. The mixture was heated under reflux for 2h. Then, the solvent was evaporated *in vacuo* to yield the crude product, which was used directly in the next step.

To a suspension of the required amine and triethylamine in anh. DCM, a solution of the proper acyl chloride in anh. DCM was slowly added. The mixture was kept under stirring at room temperature for 16 h. Then, water (15 mL) followed by EtOAc (15 mL) were added and the mixture was extracted. The organic layer was washed with brine (15 mL), dried over anh. Na_2_SO_4_ and filtered. Solvents were concentrated *in vacuo* and the resulting crude was purified by column chromatography in silica gel (using as eluent mixtures of EtOAc in hexane from 0% to 10%) to obtain the desired product.

### *N*-[(2,5-dimethylthiophen-3-yl)methyl]-3-fluoro-5-(trifluoromethyl)benzamide, 22

Following general procedure A, 3-fluoro-5-(trifluoromethyl)benzoyl chloride (88 mg, 0.39 mmol) was reacted with (2,5-dimethyl-thiophen-3-yl)methanamine (50 mg, 0.35 mmol) giving the product as a white solid (101 mg, 86% yield). m.p. 115-116 °C. ν: 3232, 3086, 2922, 1767, 1643, 1602, 1552, 1468, 1446, 1427, 1367, 1346, 1293, 1252, 1220, 1174, 1134, 1125, 1093, 1042, 998, 932, 909, 882, 837, 757, 733, 692 cm^-1^. ^1^H NMR (400 MHz, CDCl_3_) δ: 2.39 (s, 6H, —C*<u>H</u>*_*<u>3</u>*_), 4.45 (s, 2H, —NH—C*<u>H</u>*_*<u>2</u>*_), 6.30 (broad s, 1H, NH), 6.59 (s, 1H, C*<u>H</u>* thiophene), 7.45 (d, *J* = 8.0 Hz, 1H, H(4)arom.), 7.68 (d, *J* = 8.6 Hz, 1H, H(2)arom.), 7.79 (pseudo s, 1H, H(6)arom.). ^13^C NMR (101 MHz, CDCl_3_) δ: 12.8, 15.0, 37.5, 115.7 (dq, J_CF_ = 24.4, 3.7 Hz), 117.9 (d, J_CF_ = 22.8 Hz), 119.4 (p, J_CF_ = 3.5 Hz), 122.8 (d, J = 271.7 Hz), 126.2, 132.4, 132.9 (dd, J_CF_ = 34.1, 7.9 Hz), 134.3, 136.7, 137.7 (d, J_CF_ = 6.9 Hz), 162.5 (d, J_CF_ = 251.1 Hz), 164.4 (d, J_CF_= 2.1 Hz).

### 3-chloro-*N*-[(2,5-dimethylthiophen-3-yl)methyl]-5-(trifluoromethyl)benzamide, 23 (VF-57a)

Following general procedure B, 3-chloro-5-(trifluoromethyl)benzoyl chloride (106 mg, 0.44 mmol) was reacted with (2,5-dimethylthiophen-3-yl)methanamine (56 mg, 0.40 mmol) giving the product as a white solid (62 mg, 45% yield). m.p. 121-122 °C. ν: 3290, 3086, 2920, 1638, 1613, 1591, 1581, 1549, 1454, 1433, 1357, 1324, 1288, 1261, 1250, 1177, 1129, 1111, 1039, 980, 921, 889, 839, 830, 739, 715, 699, 689, 665 cm^-1^. ^1^H NMR (400 MHz, CDCl_3_) δ: 2.39 (s, 6H, — C*<u>H</u>*_*<u>3</u>*_), 4.46 (d, *J* = 5.2 Hz, 2H, —NH—C*<u>H</u>*_*<u>2</u>*_), 6.30 (broad s, 1H, NH), 6.59 (s, 1H, C*<u>H</u>* thiophene), 7.72 (s, 1H, H(4)arom.), 7.89 (s, 1H, H(6)arom.), 7.92 (s, 1H, H(2)arom.). ^13^C NMR (101 MHz, CDCl_3_) δ: 12.8, 15.1, 37.5, 122.1 (d, *J*_*CF*_ = 3.5 Hz), 126.2, 128.3 (d, *J*_*-F*_ = 3.5 Hz, 2C), 130.6, 132.4, 132.5, 134.3, 135.6 (d, *J*_CF_ = 35.3 Hz), 136.7, 136.9, 164.4.

### 3-bromo-*N*-[(2,5-dimethylthiophen-3-yl)methyl]-5-(trifluoromethyl)benzamide, 24

Following general procedure B, 3-bromo-5-(trifluoromethyl)benzoyl chloride (150 mg, 0.39 mmol) was reacted with (2,5-dimethylthiophen-3-yl)methanamine (50mg, 0.35mmol) giving the product as a white solid (113 mg, 82% yield). m.p. 124-125 °C. ν: 3289, 3082, 2919, 1636, 1614, 1581, 1547, 1448, 1432, 1356, 1321, 1285, 1263, 1250, 1174, 1130, 1110, 1038, 977, 924, 888, 839, 807, 723, 698, 687, 665 cm^-1^. ^1^H NMR (400 MHz, CDCl_3_) δ: 2.39 (s, 6H, —C*<u>H</u>*_*<u>3</u>*_), 4.46 (d, *J* = 5.2 Hz, 2H, —NH—C*<u>H</u>*_*<u>2</u>*_), 6.28 (broad s, 1H, NH), 6.59 (s, 1H, C*<u>H</u>* thiophene), 7.87 (s, 1H, H(4)arom.), 7.94 (s, 1H, H(6)arom.), 8.07 (s, 1H, H(2)arom.). ^13^C NMR (101 MHz, CDCl_3_) δ: 12.9, 15.1, 37.6, 122.8 (d, *J*_*CF*_ = 274.7 Hz), 122.7 (d, *J*_*CF*_ = 4.2 Hz), 123.2, 126.3, 131.2 (d, *J*_*CF*_ = 4.2 Hz), 132.4, 132.8 (d, *J*_*CF*_ = 31.8 Hz), 133.5, 134.3, 136.7, 137.1, 164.3.

### *N*-[(2,5-dimethylthiophen-3-yl)methyl]-3,5-bis(trifluoromethyl)benzamide, 25

Following general procedure B, 3,5-*bis(trifluoromethyl*)benzoyl chloride (130 mg, 0.47 mmol) was reacted with (2,5-dimethylthiophen-3-yl)methanamine (60 mg, 0.43 mmol) giving the product as a white solid (100 mg, 70% yield). m.p. 163-165 °C. ν: 3233, 3090, 2930, 1644, 1617, 1552, 1429, 1385, 1350, 1329, 1286, 1276, 1250, 1183, 1168, 1158, 1131, 1040, 987, 909, 871, 837, 756, 734, 699, 682 cm^-1^. ^1^H NMR (400 MHz, CDCl_3_) δ: 2.39 (s superimposed, 3H, —C*<u>H</u>*_*<u>3</u>*_), 2.40 (s superimposed, 3H, —C*<u>H</u>*_*<u>3</u>*_), 4.49 (d, *J* = 5.2 Hz, 2H, —NH— C*<u>H</u>*_*<u>2</u>*_), 6.38 (broad s, 1H, NH), 6.60 (s, 1H, C*<u>H</u>* thiophene), 8.00 (s, 1H, H(4)arom.), 8.21 (s, 2H, H(2,6)arom.). ^13^C NMR (101 MHz, CDCl_3_) δ: 12.8, 15.1, 37.6, 122.9 (d, 2C, *J*_*CF*_ = 272.6 Hz), 125.0 (t, *J*_*CF*_ = 3.6 Hz), 126.2, 127.3, 127.3 (d, 2C, *J*_*CF*_ = 2.5 Hz), 132.2 (q, *J*_*CF*_ = 34.3 Hz), 132.3, 134.4, 136.4, 136.8, 164.2.

### 3,5-difluoro-*N*-[(2,5-dimethylthiophen-3-yl)methyl]benzamide, 26

Following general procedure B, 3,5-difluorobenzoyl chloride (118 mg, 0.47 mmol) was reacted with (2,5-dimethylthiophen-3-yl)methanamine (60 mg, 0.43 mmol) giving the product as a white solid (35 mg, 29% yield). m.p. 120-121 °C. ν: 3249, 3092, 3068, 2924, 2868, 1635, 1591, 1542, 1465, 1435, 1354, 1323, 1310, 1257, 1226, 1214, 1146, 1125, 1111, 1057, 1042, 984, 899, 878, 868, 853, 836, 820, 783, 761, 717, 662, 643 cm^-1^. ^1^H NMR (400 MHz, CDCl_3_) δ: 2.38 (s, 6H, —C*<u>H</u>*_*<u>3</u>*_), 4.43 (d, *J* = 5.2 Hz, 2H, —NH—C*<u>H</u>*_*<u>2</u>*_), 6.25 (broad s, 1H, NH), 6.57 (s, 1H, C*<u>H</u>* thiophene), 6.93 (t, *J* = 8.2 Hz, 1H, H(4)arom.), 7.28 (s, 2H, H(2,6)arom.). _13_C NMR (101 MHz, CDCl_3_) δ: 12.8, 15.1, 37.5, 106.8 (t, *J*_*CF*_ = 27.4 Hz), 110.2 (d, 2C, *J*_*CF*_ = 26.8 Hz), 126.2, 132.6, 134.1, 136.6, 137.6, 161.7 (d, *J*_*CF*_ = 252.1 Hz), 162.9 (d, *J*_*CF*_ = 251.2 Hz), 164.7.

### 3,5-dichloro-*N*-[(2,5-dimethylthiophen-3-yl)methyl]benzamide, 27

By following general procedure A, 3,5-dichlorobenzoyl chloride (81 mg, 0.39 mmol) was reacted with with (2,5-dimethylthiophen-3-yl)methanamine (50 mg, 0.35 mmol) giving the product as a white solid (56 mg, 46% yield). m.p. 136-137 °C. ν: 3233, 3058, 2916, 2865, 1630, 1592, 1566, 1538, 1458, 1433, 1351, 1304, 1283, 1216, 1146, 1115, 1096, 1053, 935, 889, 864, 820, 801, 762, 745, 700, 685, 660, 638 cm^-1^. ^1^H NMR (400 MHz, CDCl_3_) δ: 2.39 (s, 3H, —C*<u>H</u>*_*<u>3</u>*_), 2.38 (s, 3H, —C*<u>H</u>*_*<u>3</u>*_), 4.44 (d, J = 5.2 Hz, 2H, —NH—C*<u>H</u>*_*<u>2</u>*_), 6.15 (broad s, 1H, NH), 6.58 (s, 1H, C*<u>H</u>* thiophene), 7.47 (t, *J* = 1.8 Hz, 1H, H(4)arom.), 7.66-7.60 (m, 2H, H(2,6)arom.). ^13^C NMR (101 MHz, CDCl_3_) δ: 12.8, 15.1, 37.5, 125.6 (2C), 126.2, 131.4, 132.5, 134.2, 135.5 (2C), 136.7, 137.3, 164.6.

### 3,5-dibromo-*N*-[(2,5-dimethylthiophen-3-yl)methyl]benzamide, 28

Following general procedure B, 3,5-dibromobenzoyl chloride (178 mg, 0.60 mmol) was reacted with (2,5-dimethylthiophen-3-yl)methanamine (70 mg, 0.50 mmol) giving the product as a white solid (101 mg, 46% yield). m.p. 134-135 °C. ν: 3233, 3058, 2916, 2865, 1630, 1592, 1566, 1538, 1458, 1433, 1351, 1304, 1283, 1216, 1146, 1115, 1096, 1053, 935, 889, 864, 820, 801, 762, 745, 700, 685, 660, 638 cm_^-1^. 1_H NMR (400 MHz, CDCl^3^) δ: 2.39 (s, 3H, —C*<u>H</u>*_*<u>3</u>*_), 2.40 (s, 3H, —C*<u>H</u>*_*<u>3</u>*_), 4.44 (d, *J* = 5.2 Hz, 2H, —NH—C*<u>H</u>*_*<u>2</u>*_), 6.09 (broad s, 1H, NH), 6.60 – 6.52 (m, 1H, C*<u>H</u>* thiophene), 7.78 (t, *J* = 1.7 Hz, 1H, H(4)arom.), 7.81 (d, *J* = 1.7 Hz, 2H, H(2,6)arom.).

### 3-chloro-*N*-[(2,5-dimethylthiophen-3-yl)methyl]-5-methylbenzamide, 29

Following general procedure B, 3-chloro-5-methylbenzoyl chloride (61 mg, 0.32 mmol) was reacted with (2,5-dimethylthiophen-3-yl)methanamine (35 mg, 0.25 mmol) giving the product as a white solid (65 mg, 90% yield). m.p. 102-103 °C. ν: 3282, 2918, 2856, 1628, 1600, 1578, 1527, 1455, 1436, 1371, 1346, 1316, 1289, 1223, 1146, 1056, 1035, 994, 864, 770, 748, 677, 644 cm^-1^. ^1^H NMR (400 MHz, CDCl_3_) δ: 2.36 (s, 3H, C*H*_*3*_— C(5)arom.), 2.38 (s superimposed, 3H, —C*<u>H</u>*_*<u>3</u>*_), 2.39 (s superimposed, 3H, —C*<u>H</u>*_*<u>3</u>*_), 4.43 (d, *J* = 5.3 Hz, 2H, —NH— C*<u>H</u>*_*<u>2</u>*_), 6.19 (broad s, 1H, NH), 6.56 (s, 1H, C*<u>H</u>* thiophene), 7.28 (s, 1H, H(4)arom.), 7.45 (s, 1H, H(6)arom.), 7.51 (s, 1H, H(2)arom.). ^13^C NMR (101 MHz, CDCl_3_) δ: 12.8, 15.2, 21.1, 37.4, 124.2, 125.9, 126.3, 132.0, 132.9, 133.9, 134.4, 135.9, 136.5, 140.3, 166.0.

### *N*-[(2,5-dimethylthiophen-3-yl)methyl]-3-fluorobenzamide, 30

Following general procedure A, 3-fluorobenzoyl chloride (63 mg, 0.39 mmol) was reacted with (2,5-dimethylthiophen-3-yl)methanamine (50 mg, 0.35 mmol) giving the product as a white solid (93 mg, 94% yield). m.p. 93-94 °C. ν: 3245, 3072, 2920, 1632, 1587, 1548, 1485, 1445, 1426, 1358, 1316, 1306, 1274, 1256, 1225, 1144, 1117, 1039, 980, 904, 891, 832, 808, 787, 726, 708, 674, 666 cm^-1^. ^1^H NMR (400 MHz, CDCl_3_) δ: 2.38 (s, 6H, — C*<u>H</u>*_*<u>3</u>*_), 4.45 (d, J = 5.2 Hz, 2H, —NH—C*<u>H</u>*_*<u>2</u>*_), 6.26 (broad s, 1H, NH), 6.59 (s, 1H, C*<u>H</u>* thiophene), 7.23-7.13 (m, *J* = 8.2, 0.8 Hz, 1H, H(4)arom.), 7.43-7.33 (m, *J* = 8.1, 2.1 Hz, 1H, H(5)arom.), 7.53–7.46 (m, 2H, H(2,6)arom.). ^13^C NMR (101 MHz, CDCl_3_) δ: 12.8, 15.1, 37.4, 114.4 (d, J_CF_ = 23.4 Hz), 118.5 (d, J_CF_ = 21.1 Hz), 122.4, 126.3, 130.2 (d, J_CF_ = 8.1 Hz), 132.9, 133.9, 136.5, 136.7, 162.7 (d, J_CF_ = 248.4 Hz), 165.9.

### *N*-[(2,5-dimethylthiophen-3-yl)methyl]-3-(trifluoromethyl)benzamide, 31

Following general procedure A, 3-(trifluoromethyl) benzoyl chloride (81 mg, 0.39 mmol) was reacted with (2,5-dimethylthiophen-3-yl)methanamine (50 mg, 0.35 mmol) giving the product as a white solid (63 mg, 54% yield). m.p. 111-112 °C. ν: 3225, 3071, 2924, 1636, 1542, 1487, 1443, 1427, 1360, 1336, 1319, 1301, 1281, 1256, 1213, 1166, 1115, 1071, 1040, 975, 921, 837, 817, 785, 735, 690, 653 cm^-1. 1^H NMR (400 MHz, CDCl_3_) δ: 2.39 (s, 6H, —CH_3_), 4.47 (d, J = 5.2 Hz, 2H, — NH—CH_2_), 6.28 (broad s, 1H, NH), 6.60 (s, 1H, CH thiophene), 7.56 (t, J = 7.8 Hz, 1H, H(5)arom.), 7.75 (dt, J = 7.9, 0.7 Hz, 1H, H(6)arom.), 7.95 (d, *J* = 7.8, 1H, H(4)arom.), 8.03 (s, 1H, H(2)arom.). ^13^C NMR (101 MHz, CDCl_3_) δ: 12.81, 15.07, 37.47, 123.68 (d, *J*_*CF*_ = 273 Hz), 123.95 (q, *J*_*CF*_ = 3.8 Hz), 126.29, 128.09 (q, *J*_*CF*_ = 3.8 Hz), 129.19, 130.23, 131.13 (d, *J*_*CF*_ = 33.0 Hz), 132.7, 134.1, 135.2, 136.6, 165.7.

### 3-chloro-*N*-[(2,5-dimethylthiophen-3-yl)methyl]benzamide, 32

Following general procedure A, 3-chlorobenzoyl chloride (62 mg, 0.39 mmol) was reacted with (2,5-dimethylthiophen-3-yl)methanamine (50 mg, 0.35 mmol) giving the product as a white solid (74 mg, 79% yield). m.p. 89-90 °C. ν: 3240, 3071, 2916, 1631, 1549, 1474, 1430, 1357, 1297, 1255, 1212, 1162, 1145, 1077, 1040, 974, 911, 832, 808, 728, 707, 660 cm^-1. 1^H NMR (400 MHz, CDCl_3_) δ: 2.38 (s, 6H, —C*<u>H</u>*_*<u>3</u>*_), 4.44 (d, *J* = 5.2 Hz, 2H, — NH—C*<u>H</u>*_*<u>2</u>*_), 6.26 (broad s, 1H, NH), 6.58 (s, 1H, C*<u>H</u>* thiophene), 7.34 (t, *J* = 7.9 Hz, 1H, H(5)arom.), 7.45 (ddd, *J* = 7.9, 2.0, 1.0 Hz, 1H, H(4)arom.), 7.63 (dt, *J* = 7.8, 1.3 Hz, 1H, H(6)arom.), 7.75 (t, *J* = 1.7 Hz, 1H, H(2)arom.) ^13^C NMR (101 MHz, CDCl_3_) δ: 12.8, 15.1, 37.4, 125.0, 126.3, 127.3, 129.9, 131.5, 132.8, 133.9, 134.7, 136.2, 136.5, 165.8.

### 3-bromo-*N*-[(2,5-dimethylthiophen-3-yl)methyl]benzamide, 33

Following general procedure A, 3-fluorobenzoyl chloride (68 mg, 0.31mmol) was reacted with (2,5-dimethylthiophen-3-yl)methanamine (40 mg, 0.28 mmol) giving the product as a white solid (55 mg, 60% yield). m.p. 91-92 °C. ν: 3233, 3067, 2948, 1631, 1549, 1471, 1428, 1357, 1314, 1295, 1255, 1211, 1146, 1072, 1039, 971, 912, 833, 808, 731, 709 cm^-1. 1^H NMR (400 MHz, CDCl_3_) δ: 2.38 (s, 6H, —C*<u>H</u>*_*<u>3</u>*_), 4.44 (d, J = 5.2 Hz, 2H, — NH—C*<u>H</u>*_*<u>2</u>*_), 6.21 (broad s, 1H, NH), 6.58 (s, 1H, C*<u>H</u>* thiophene), 7.29 (t, *J* = 7.8 Hz, 1H, H(5)arom.), 7.61 (dd, *J* = 8.0, 2.0 Hz, 1H, H(4)arom.), 7.68 (dd, *J* = 7.8, 1.6 Hz, 1H, H(6)arom.), 7.90 (t, *J* = 1.7 Hz, 1H, H(2)arom.). ^13^C NMR (101 MHz, CDCl_3_) δ: 12.8, 15.1, 37.4, 122.7, 125.5, 126.3, 130.1 (2C), 132.8, 133.9, 134.4, 136.4, 136.5, 165.7.

### *N*-[(2,5-dimethylthiophen-3-yl)methyl]-3-nitrobenzamide, 34

Following general procedure A, 3-nitrobenzoyl chloride (72 mg, 0.39 mmol) was reacted with (2,5-dimethylthiophen-3-yl)methanamine (50 mg, 0.35 mmol) giving the product as a white solid (86 mg, 84% yield). m.p. 156-157 °C. ν: 3233, 3086, 2911, 1635, 1618, 1580, 1552, 1523, 1474, 1426, 1349, 1322, 1307, 1275, 1256, 1214, 1145, 1086, 1042, 977, 921, 885, 842, 816, 750, 723, 712, 702 cm^-1. 1^H NMR (400 MHz, CDCl_3_) δ: 2.38 (s, 3H, — C*<u>H</u>*_*<u>3</u>*_), 2.40 (s, 3H, —C*<u>H</u>*_*<u>3</u>*_), 4.49 (d, *J* = 5.4 Hz, 2H, —NH— C*<u>H</u>*_*<u>2</u>*_), 6.44 (broad s, 1H, NH), 6.60 (s, 1H, C*<u>H</u>* thiophene), 7.63 (t, *J* = 8.0 Hz, 1H, H(5)arom.), 8.16 (dt, *J* = 7.7, 1.2 Hz, 1H, H(6)arom.), 8.34 (ddd, *J* = 8.2, 2.0, 0.9 Hz, 1H, H(4)arom.), 8.57 (t, *J* = 1.8 Hz, 1H, H(2)arom.). _13_C NMR (101 MHz, CDCl^3^) δ: 12.8, 15.1, 37.6, 121.74, 126.1, 126.3, 129.8, 132.5, 133.3, 134.2, 135.9, 136.7, 148.2, 164.7.

### *N*-((2,5-dimethylthiophen-3-yl)methyl)-3,4-difluoro-5-(trifluoromethyl)benzamide, 35

Following general procedure C, 3,4-difluoro-5-(trifluoromethyl)benzoic acid (106 mg, 0.47 mmol) in anh. toluene (2.0 mL) and drops of DMF was reacted with thionyl chloride (170 µL, 279 mg, 2.35 mmol). Then, (2,5-dimethylthiophen-3-yl)methanamine hydrochloride (100 mg, 0.56 mmol) and triethylamine (234 µL, 170 mg, 1.68 mmol) in anh. DCM (1.0 mL), were reacted with the crude acyl chloride (115 mg, 0.47 mmol) in anh. DCM (1.0 mL) giving the product as a beige solid (38 mg, 23% yield). m.p. 130-131 °C. *v*: 3239, 1647, 1627, 1552, 1504, 1436, 1371, 1349, 1295, 1276, 1192, 1139, 1042, 1001, 942, 885, 838, 731, 674 cm^−1. 1^H NMR (400 MHz, CDCl_3_) δ: 2.38 (m, 6H, 2’-C<u>H</u>_3,_ 5’-C<u>H</u>_3_), 4.44 (d, *J* = 5.2 Hz, 2H, C<u>H</u>_2_), 6.34 (broad s, 1H, N<u>H</u>), 6.57 (s, 1H, 4’-H), 7.78 (m, 1H, 6-H), 7.85 (ddd, *J* = 9.6 Hz, *J’* = 6.9 Hz, *J’’* = 2.2 Hz, 1H, 2-H). ^13^C NMR (101 MHz, CDCl_3_) δ: 12.9 (C2’-<u>C</u>H_3_), 15.2 (C5’-<u>C</u>H_3_), 37.8 (CH_2_, <u>C</u>H_2_), 120.5 (^*2*^*J*_*CF*_ = 18.8 Hz, CH, C2), 120.7 (m, CH, C6), 120.8 (qd, ^*2*^*J*_*CF*_ = 33.9 Hz, ^*2*^*J*_*CF*_ = 9.6 Hz, C, C5), 121.7 (qd, ^*1*^*J*_*CF*_ = 272.9 Hz, ^*4*^*J*_*CF*_ = 3.5 Hz, C, <u>C</u>F_3_), 126.3 (CH, C4’), 131.4 (t, ^*3*^*J*_*CF*_ = 4.8 Hz, C, C1), 132.5 (C, C3’), 134.4 (C, C2’), 136.9 (C, C5’), 150.2 (dd, ^*1*^*J*_*CF*_ = 263.7 Hz, ^*2*^*J*_*CF*_ = 13.7 Hz, C, C3), 150.8 (dd, ^*1*^*J*_*CF*_ = 251.1 Hz, _*2*_*J*_*CF*_ = 9.6 Hz, C, C4), 163.8 (C, <u>C</u>O). HRMS: Calcd for [C_15_H_12_F_5_NOS+H]_+_: 350.0633, found: 350.0636.

### *N*-((2,5-dimethylthiophen-3-yl)methyl)-2,3,4-trifluoro-5-(trifluoromethyl)benzamide, 36

Following general procedure C, 2,3,4-trifluoro-5-(trifluoromethyl)benzoic acid (150 mg, 0.61 mmol) in anh. toluene (4.0 mL) and drops of DMF was reacted with thionyl chloride (445 µL, 726 mg, 6.14 mmol). Then, (2,5-dimethylthiophen-3-yl)methanamine hydrochloride (131 mg, 0.74 mmol) and triethylamine (411 µL, 298 mg, 2.95 mmol) in anh. DCM (2.0 mL) were reacted with the crude acyl chloride (161 mg, 0.61 mmol) in anh. DCM (1.0 mL) giving the product as a white solid (110 mg, 49% yield). m.p. 109-110 °C. *v*: 3282, 2925, 1644, 1553, 1487, 1369, 1280, 1205, 1168, 1133, 1054, 994, 905, 721, 709, 672, 632, 574 cm_−1. 1_H NMR (400 MHz, CDCl_3_) δ: 2.39 (m, 6H, 2’-C<u>H</u>_3,_ 5’-C<u>H</u>_3_), 4.49 (d, *J* = 5.3 Hz, 2H, C<u>H</u>_2_), 6.58 (s, 1H, 4’-H), 6.60 (s, 1H, N<u>H</u>), 8.23 (td, *J* = 7.5 Hz, *J’* = 1.7 Hz, 1H, 6-H). ^13^C NMR (101 MHz, CDCl_3_) δ: 13.0 (C2’-<u>C</u>H_3_), 15.2 (C5’-<u>C</u>H_3_), 37.8 (CH_2_, <u>C</u>H_2_), 116.8 (m, C, C5), 118.7 (d, ^*2*^*J*_*CF*_ = 10.1 Hz, C, C1), 121.4 (q, ^*1*^*J*_*CF*_ = 272.0 Hz, C, <u>C</u>F_3_), 123.9 (CH, C6), 126.2 (CH, C4’), 132.3 (C, C3’), 134.3 (C, C2’), 136.9 (C, C5’), 140.5 (ddd, ^*1*^*J*_*CF*_ = 256.5 Hz, ^*2*^*J*_*CF*_ = 17.5 Hz, ^*3*^*J*_*CF*_ = 2.6 Hz, C, C2), 150.9 (dd, ^*1*^*J*_*CF*_ = 265.7 Hz, ^*2*^*J*_*CF*_ = 12.0 Hz, C, C3), 151.85 (ddd, ^*1*^*J*_*CF*_ = 259.1 Hz, ^*2*^*J*_*CF*_ = 11.0 Hz, ^*3*^*J*_*CF*_ = 5.3 Hz, C, C4), 159.9 (C, <u>C</u>O). HRMS: Calcd for [C_15_H_11_F_6_NOS+H]^+^: 368.0538, found: 368.0532.

### 3-chloro-*N*-((2,5-dimethylthiophen-3-yl)methyl)-2-fluoro-5-(trifluoromethyl)benzamide, 37

Following general procedure C, 3-chloro-2-fluoro-5-(trifluoromethyl)benzoic acid (68 mg, 0.28 mmol) in anh. toluene (2.0 mL) and drops of DMF was reacted with thionyl chloride (101 µL, 166 mg, 1.40 mmol). Then, (2,5-dimethylthiophen-3-yl)methanamine hydrochloride (50 mg, 0.28 mmol) and triethylamine (117 µL, 85 mg, 0.84 mmol) in anh. DCM (0.5 mL), were reacted with the crude acyl chloride (73 mg, 0.28 mmol) in anh. DCM (0.5 mL) giving the product as a white solid (98 mg, 95% yield). m.p. 118-119 °C. *v*: 3285, 2920, 1633, 1544, 1345, 1328, 1277, 1161, 1144, 1122, 1055, 896, 867, 767, 723, 661, 648, 578 cm^−1. 1^H NMR (400 MHz, CDCl_3_) δ: 2.40 (m, 6H, 2’-C<u>H</u>_3,_ 5’-C<u>H</u>_3_), 4.50 (d, *J* = 5.2 Hz, 2H, C<u>H</u>_2_), 6.59 (s, 1H, 4’-H), 6.68 (broad s, 1H, N<u>H</u>), 7.80 (dd, *J* = 6.5 Hz, *J’* = 2.4 Hz, 1H, 4-H), 8.30 (dd, *J* = 6.2 Hz, *J’* = 2.5 Hz, 1H, 6-H). ^13^C NMR (101 MHz, CDCl_3_) δ: 13.0 (C2’-<u>C</u>H_3_), 15.2 (C5’-<u>C</u>H_3_), 37.9 (CH_2_, <u>C</u>H_2_), 122.8 (q, ^*1*^*J*_*CF*_ = 272.6 Hz, C, <u>C</u>F_3_), 123.1 (d, ^*2*^*J*_*CF*_ = 21.4 Hz, C, C1), 123.6 (d, ^*2*^*J*_*CF*_ = 13.7 Hz, C, C3), 126.2 (CH, C4’), 128.1 (m, C, C5), 128.2 (p, ^*3*^*J*_*CF*_ = 3.6 Hz, CH, C6), 130.8 (CH, C4), 132.4 (C, C3’), 134.3 (C, C2’), 136.8 (C, C5’), 157.8 (d, ^*1*^*J*_*CF*_ = 253.7 Hz, C, C2), 160.8 (d, ^*3*^*J*_*CF*_ = 3.5 Hz, C, <u>C</u>O). HRMS: Calcd for [C_15_H_12_ClF_4_NOS+H]^+^: 366.0337, found: 366.0339.

### 3-chloro-*N*-((2,5-dimethylthiophen-3-yl)methyl)-4-fluoro-5-(trifluoromethyl)benzamide, 38

Following general procedure C, 3-chloro-4-fluoro-5-(trifluoromethyl)benzoic acid (68 mg, 0.28 mmol) in anh. toluene (1.0 mL) and drops of DMF was reacted with thionyl chloride (101 µL, 166 mg, 1.40 mmol). Then, (2,5-dimethylthiophen-3-yl)methanamine hydrochloride (50 mg, 0.28 mmol) and triethylamine (117 µL, 85 mg, 0.84 mmol) in anh. DCM (0.5 mL), were reacted with the crude acyl chloride (73 mg, 0.28 mmol) in anh. DCM (0.5 mL) giving the product as a white solid (43 mg, 42% yield). m.p. 133-134 °C. *v*: 3273, 1636, 1552, 1481, 1417, 1327, 1317, 1262, 1221, 1140, 1037, 919, 904, 833, 744, 716, 672, 626, 573 cm^−1. 1^H NMR (400 MHz, CDCl_3_) δ: 2.38 (m, 6H, 2’-C<u>H</u>_3,_ 5’-C<u>H</u>_3_), 4.44 (dd, *J* = 5.3 Hz, *J’* = 2.0 Hz, 2H, C<u>H</u>_2_), 6.29 (broad s, 1H, N<u>H</u>), 6.57 (s, 1H, 4’-H), 7.92 (dd, *J* = 5.9 Hz, *J’* = 2.2 Hz, 1H, 6-H), 8.04 (dd, *J* = 6.3 Hz, *J’* = 2.2 Hz, 1H, 2-H). ^13^C NMR (101 MHz, CDCl_3_) δ: 13.0 (C2’-<u>C</u>H_3_), 15.2 (C5’-<u>C</u>H_3_), 37.8 (CH_2_, <u>C</u>H_2_), 120.2 (qd, ^*2*^*J*_*CF*_ = 34.0 Hz, ^*2*^*J*_*CF*_ = 13.0 Hz, C, C5), 121.8 (q, ^*1*^*J*_*CF*_ = 273.3 Hz, C, <u>C</u>F_3_), 123.6 (d, ^*2*^*J*_*CF*_ = 17.6 Hz, C, C3), 124.5 (m, CH, C6), 126.3 (CH, C4’), 131.4 (d, ^*4*^*J*_*CF*_ = 4.6 Hz, C, C1), 132.5 (C, C3’), 133.5 (CH, C2), 134.4 (C, C2’), 136.9 (C, C5’), 157.3 (d, ^*1*^*J*_*CF*_ = 264.0 Hz, C, C4), 163.80 (C, <u>C</u>O). HRMS: Calcd for [C_15_H_12_ClF_4_NOS+H]^+^: 366.0337, found: 366.0335.

### 2,5-dichloro-*N*-((2,5-dimethylthiophen-3-yl)methyl)-3-(trifluoromethyl)benzamide, 39

Following general procedure C, 2,5-dichloro-3-(trifluoromethyl)benzoic acid (122 mg, 0.47 mmol) in anh. toluene (2.0 mL) and drops of DMF was reacted with thionyl chloride (170 µL, 279 mg, 2.35 mmol). Then, (2,5-dimethylthiophen-3-yl)methanamine hydrochloride (100 mg, 0.56 mmol) and triethylamine (234 µL, 170 mg, 1.68 mmol) in anh. DCM (1.0 mL) were reacted with the crude acyl chloride (130 mg, 0.47 mmol) in anh. DCM (1.0 mL) giving the product as a white solid (110 mg, 49% yield). m.p. 157-158 °C. *v*: 3268, 1646, 1544, 1482, 1429, 1316, 1289, 1258, 1172, 1142, 1051, 886, 830, 742, 711, 689, 830, 576 cm^−1. 1^H NMR (400 MHz, CDCl_3_) δ: 2.39 (m, 6H, 2’-C<u>H</u>_3,_ 5’-C<u>H</u>_3_), 4.46 (d, *J* = 5.2 Hz, 2H, C<u>H</u>_2_), 6.03 (s, broad, 1H, N<u>H</u>), 6.59 (s, 1H, 4’-H), 7.67 (d, *J* = 2.5 Hz, 1H, 2-H), 7.72 (d, *J* = 2.5 Hz, 1H, 4-H). ^13^C NMR (101 MHz, CDCl_3_) δ: 13.0 (C2’-<u>C</u>H_3_), 15.2 (C5’-<u>C</u>H_3_), 37.7 (CH_2_, <u>C</u>H_2_), 121.9 (q, ^*1*^*J*_*CF*_ = 274.2 Hz, C, <u>C</u>F_3_), 126.3 (CH, C4’), 127.5 (C, C6), 129.2 (q, ^*3*^*J*_*CF*_ = 5.6 Hz, CH, C4), 130.9 (q, ^*2*^*J*_*CF*_ = 32.0 Hz, C, C5), 132.1 (C, C3’), 132.6 (CH, C2), 133.5 (C, C3), 134.5 (C, C2’), 136.8 (C, C5’), 139.8 (C, C1), 164.3 (C, <u>C</u>O). HRMS: Calcd for [C_15_H_12_Cl_2_F_3_NOS+H]^+^: 382.0042, found: 382.0047.

### *N*-((2,5-dimethylthiophen-3-yl)methyl)benzamide, 40

Following general procedure D, benzoic acid (41 mg, 0.34 mmol) in anh. toluene (1.0 mL) and drops of DMF was reacted with thionyl chloride (245 µL, 402 mg, 3.38 mmol). Then, (2,5-dimethylthiophen-3-yl)methanamine hydrochloride (50 mg, 0.28 mmol) and triethylamine (118 µL, 85 mg, 0.84 mmol) in anh. DCM (1.0 mL) reacted with the crude acyl chloride in anh. DCM (1.0 mL) giving the product as a white solid (26 mg, 39% yield). m.p. 98-99 °C. *v*: 3237, 1627, 1536, 1490, 1356, 1307, 1253, 1210, 1141, 1038, 961, 931, 830, 807, 718, 694, 568 cm^−1. 1^H NMR (400 MHz, CDCl_3_) δ: 2.38 (s, 6H, 2’-C<u>H</u>_3_, 5’-C<u>H</u>_3_), 4.46 (d, *J* = 5.2 Hz, 2H, C<u>H</u>_2_), 6.24 (broad s, 1H, N<u>H</u>), 6.60 (m, 1H, 4’-H), 7.42 [m, 2H, 3(5)-H], 7.49 (m, 1H, 4-H), 7.77 [m, 2H, 2(6)-H]. ^13^C NMR (101 MHz, CDCl_3_) δ: 12.9 (C2’-<u>C</u>H_3_), 15.2 (C5’-<u>C</u>H_3_), 37.5 (CH_2_, <u>C</u>H_2_), 126.5 (CH, C4’), 127.1 [CH, C2(6)], 128.7 [CH, C3(5)], 131.6 (CH, C4), 133.3 (C, C3’), 133.9 (C, C1), 134.5 (C, C2’), 136.5 (C, C5’), 167.3 (C, <u>C</u>O). HRMS: Calcd for [C_14_H_15_NOS+H]^+^: 246.0947, found: 246.0944.

### 4-chloro-*N*-((2,5-dimethylthiophen-3-yl)methyl)benzamide, 41

Following general procedure D, 4-chlorobenzoic acid (53 mg, 0.34 mmol) in anh. toluene (1.0 mL) and drops of DMF was reacted with thionyl chloride (245 µL, 402 mg, 3.38 mmol). Then, (2,5-dimethylthiophen-3-yl)methanamine hydrochloride (50 mg, 0.28 mmol) and triethylamine (118 µL, 85 mg, 0.84 mmol) in anh. DCM (1.0 mL) reacted with the crude acyl chloride in anh. DCM (1.0 mL) giving the product as an off-white solid (59 mg, 75% yield). m.p. 133-134 °C. *v*: 3313, 2915, 1715, 1632, 1544, 1486, 1447, 1357, 1318, 1214, 1090, 1039, 1013, 851, 514, 759, 708, 635, 570 cm^−1. 1^H NMR (400 MHz, CDCl_3_) δ: 2.37-2.39 (complex signal, 6H, 2’-C<u>H</u>_3_, 5’-C<u>H</u>_3_), 4.44 (d, *J* = 5.2 Hz, 2H, C<u>H</u>_2_), 6.19 (broad s, 1H, N<u>H</u>), 6.58 (q, *J* = 1.1 Hz, 1H, 4’-H), 7.38 [d, *J* = 8.8 Hz, 2H, 3(5)-H], 7.70 [d, *J* = 8.8 Hz, 2H, 2(6)-H]. ^13^C NMR (101 MHz, CDCl_3_) δ: 13.0 (C2’-<u>C</u>H_3_), 15.2 (C5’-<u>C</u>H_3_), 37.5 (CH_2_, <u>C</u>H_2_), 126.4 (CH, C4’), 128.5 [CH, C2(6)], 128.9 [CH, C3(5)], 132.9 (C, C1), 133.1 (C, C3’), 134.1 (C, C2’), 136.6 (C, C5’), 137.9 (C, C4), 166.2 (C, <u>C</u>O). HRMS: Calcd for [C_14_H_14_ClNOS+H]^+^: 280.0557, found: 280.0558.

### *N-*((2,5-dimethylthiophen-3-yl)methyl)-4-methylbenzamide, 42

Following general procedure D, 4-methylbenzoic acid (46 mg, 0.34 mmol) in anh. toluene (1.0 mL) and drops of DMF was reacted with thionyl chloride (245 µL, 402 mg, 3.38 mmol). Then, (2,5-dimethylthiophen-3-yl)methanamine hydrochloride (50 mg, 0.28 mmol) and triethylamine (118 µL, 85 mg, 0.84 mmol) in anh. DCM (1.0 mL) reacted with the crude acyl chloride in anh. DCM (1.0 mL) giving the product as a beige solid (39 mg, 53% yield). m.p. 132-133 °C. *v*: 277, 2917, 1631, 1612, 1532, 1501, 1348, 1292, 1279, 1211, 1187, 1140, 1118, 1062, 1021, 966, 907, 835, 752, 655, 632, 607, 571 cm^−1. 1^H NMR (400 MHz, CDCl_3_) δ: 2.38-2.38 (complex signal, 9H, 4-C<u>H</u>_3_, 2’-C<u>H</u>_3_, 5’-C<u>H</u>_3_), 4.44 (d, *J* = 5.2 Hz, 2H, C<u>H</u>_2_), 6.25 (broad s, 1H, N<u>H</u>), 6.59 (q, *J* = 1.2 Hz, 1H, 4’-H), 7.21 [dd, *J* = 8.6, 0.7 Hz, 2H, 3(5)-H], 7.66 [d, *J* = 8.2 Hz, 2H, 2(6)-H]. ^13^C NMR (101 MHz, CDCl_3_) δ: 12.9 (C2’-<u>C</u>H_3_), 15.2 (C5’-<u>C</u>H_3_), 21.5 (C4-<u>C</u>H_3_), 37.4 (CH_2_, <u>C</u>H_2_), 126.5 (CH, C4’), 127.0 [CH, C2(6)], 129.3 [CH, C3(5)], 131.6 (C, C1), 133.5 (C, C3’), 133.8 (C, C2’), 136.4 (C, C5’), 142.0 (C, C4), 167.2 (C, <u>C</u>O). HRMS: Calcd for [C_15_H_17_NOS+H]^+^: 260.1104, found: 260.1109.

### *N*-((2,5-dimethylthiophen-3-yl)methyl)-4-methoxybenzamide, 43

Following general procedure D, 4-methoxybenzoic acid (83 mg, 0.54 mmol) in anh. toluene (1.0 mL) and drops of DMF was reacted with thionyl chloride (391 µL, 641 mg, 5.39 mmol). Then, (2,5-dimethylthiophen-3-yl)methanamine hydrochloride (80 mg, 0.45 mmol) and triethylamine (211 µL, 152 mg, 1.50 mmol) in anh. DCM (1.0 mL) reacted with the crude acyl chloride in anh. DCM (1.0 mL) giving the product as a brownish solid (46 mg, 38% yield). m.p. 117-118 °C. *v*: 3309, 2914, 2838, 1633, 1606, 1549, 1504, 1462, 1317, 1252, 1214, 1174, 1112, 1031, 974, 844, 825, 770, 722, 661, 632, 606 cm^−1. 1^H NMR (400 MHz, CDCl_3_) δ: 2.36-2.39 (complex signal, 6H, 2’-C<u>H</u>_3_, 5’-C<u>H</u>_3_), 3.83 (s, OC<u>H</u>_3_), 4.43 (d, *J* = 5.2 Hz, 2H, C<u>H</u>_2_), 6.17 (broad s, 1H, N<u>H</u>), 6.60 (m, 1H, 4’-H), 6.90 [d, *J* = 8.8 Hz, 2H, 3(5)-H], 7.73 [d, *J* = 8.9 Hz, 2H, 2(6)-H]. ^13^C NMR (101 MHz, CDCl_3_) δ: 12.9 (C2’-<u>C</u>H_3_), 15.2 (C5’-<u>C</u>H_3_), 37.4 CH_2_, <u>C</u>H_2_), 55.5 (CH_3_, O<u>C</u>H_3_), 113.8 [CH, C3(5)], 126.6 (CH, C4’), 126.8 (C, C1), 128.9 [CH, C2(6)], 133.5 (C, C3’), 133.8 (C, C2’), 136.4 (C, C5’), 162.3 (C, C4), 166.8 (C, <u>C</u>O). HRMS: Calcd for [C_15_H_17_NO_2_S+H]^+^: 279.1053, found: 279.1059.

### 3,4-dichloro-*N*-((2,5-dimethylthiophen-3-yl)methyl)benzamide, 44

Following general procedure D, 3,4-dichlorobenzoic acid (90 mg, 0.47 mmol) in anh. toluene (1.0 mL) and drops of DMF was reacted with thionyl chloride (339 µL, 556 mg, 4.67 mmol). Then, (2,5-dimethylthiophen-3-yl)methenamine hydrochloride (70 mg, 0.39 mmol) and triethylamine (164 µL, 118 mg, 1.17 mmol) in anh. DCM (1.0 mL) reacted with the crude acyl chloride in anh. DCM (1.0 mL) giving the product as a brownish solid (43 mg, 35% yield). m.p. 102-103 °C. *v*: 3241, 2917, 1634, 1593, 1538, 1469, 1425, 1375, 1354, 1309, 1251, 1213, 1142, 1131, 1029, 971, 886, 858, 833, 760, 724, 683, 670, 567 cm^−1. 1^H NMR (400 MHz, CDCl_3_) δ: 2.38 (s, 3H, 2’-C<u>H</u>_3_), 2.39 (m, 3H, 5’-C<u>H</u>_3_), 4.43 (d, *J* = 5.2 Hz, 2H, C<u>H</u>_2_), 6.22 (broad s, 1H, N<u>H</u>), 6.58 (d, *J* = 1.2 Hz, 1H, 4’-H), 7.48 (d, *J* = 8.3 Hz, 1H, 5-H), 7.58 (dd, *J* = 8.3 Hz, *J’* = 2.1 Hz, 1H, 6-H), 7.85 (d, *J* = 2.1 Hz, 1H, 2-H). _13_C NMR (101 MHz, CDCl_3_) δ: 13.0 (C2’-<u>C</u>H_3_), 15.2 (C5’-<u>C</u>H_3_), 37.6 (CH_2_, <u>C</u>H_2_), 126.3 (CH, C6), 126.4 (CH, C4’), 129.3 (CH, C2), 130.8 (CH, C5), 132.8 (C, C1), 133.2 (C, C3’), 134.2 (C, C2’), 134.3 (C, C3), 136.1 (C, C4), 136.7 (C, C5’), 165.1 (C, <u>C</u>O). HRMS: Calcd for [C_14_H_13_Cl_2_NOS+H]^+^: 314.0168, found: 314.0165.

### 3-fluoro-*N*-((5-methylthiophen-3-yl)methyl)-5-(trifluoromethyl)benzamide, 45

Following general procedure C, 3-fluoro-5-(trifluoromethyl)benzoic acid (100 mg, 0.48 mmol) in anh. toluene (2.0 mL) and drops of DMF was reacted with thionyl chloride (174 µL, 285 mg, 2.40 mmol). Then, (5-methylthiophen-3-yl)methanamine (73 mg, 0.58 mmol) and triethylamine (162 µL, 118 mg, 1.16 mmol) in anh. DCM (1.0 mL), were reacted with the crude acyl chloride (109 mg, 0.48 mmol) in anh. DCM (1.0 mL) giving the product as a yellow solid (30 mg, 20% yield). m.p. 86-87 °C. *v*: 3274, 1639, 1603, 1549, 1446, 1357, 1279, 1219, 1181, 1164, 1124, 1094, 1068, 1006, 952, 885, 835, 779, 710, 964, 626, 584 cm^−1. 1^H NMR (400 MHz, CDCl_3_) δ: 2.45 (s, 3H, 5’-C<u>H</u>_3_), 4.53 (d, *J* = 5.5 Hz, 2H, C<u>H</u>_2_), 6.57 (broad s, 1H, N<u>H</u>), 6.73 (s, 1H, 4’-H), 6.94 (s, 1H, 2’-H), 7.45 (dt, *J* = 8.3 Hz, *J’* = 1.9 Hz, 1H, 4-H), 7.70 (dt, *J* = 8.7 Hz, *J’* = 2.0 Hz, 1H, 2-H), 7.81 (s, 1H, 6-H). ^13^C NMR (101 MHz, CDCl_3_) δ: 15.4 (C5’-<u>C</u>H_3_), 39.9 (CH_2_, <u>C</u>H_2_), 115.9 (dq, ^*2*^*J*_*CF*_ = 24.6 Hz, ^*3*^*J*_*CF*_ = 3.7 Hz, CH, C4), 118.1 (d, ^*2*^*J*_*CF*_ = 22.8 Hz, CH, C2), 119.6 (p, ^*3*^*J*_*CF*_ = 3.7 Hz, CH, C6), 120.8 (CH, C2’), 123.0 (qd, ^*1*^*J*_*CF*_ = 272.8 Hz, ^*4*^*J*_*CF*_ = 2.9 Hz, C, <u>C</u>F_3_), 125.7 (CH, C4’), 133.1 (qd, ^*2*^*J*_*CF*_ = 33.8 Hz, ^*3*^*J*_*CF*_ = 7.7 Hz, C, C5), 137.9 (d, ^*3*^*J*_*CF*_ = 6.9 Hz, C1), 138.0 (C, C3’), 141.5 (C, C5’), 162.6 (d, ^*1*^*J*_*CF*_ = 251.1 Hz, C, C3), 164.7 (d, ^*4*^*J*_*CF*_ = 2.3 Hz, C, <u>C</u>O). HRMS: Calcd for [C_14_H_11_F_4_NOS+H]^+^: 318.0570, found: 318.0576.

### 3-fluoro-*N*-((2-methylthiophen-3-yl)methyl)-5-(trifluoromethyl)benzamide, 46

Following general procedure C, 3-fluoro-5-(trifluoromethyl)benzoic acid (100 mg, 0.48 mmol) in anh. toluene (2.0 mL) and drops of DMF was reacted with thionyl chloride (349 µL, 572 mg, 4.81 mmol). Then, ((2-methylthiophen-3-yl)methanamine hydrochloride (95 mg, 0.58 mmol) and triethylamine (323 µL, 235 mg, 2.32 mmol) in anh. DCM (1.0 mL), were reacted with the crude acyl chloride (109 mg, 0.48 mmol) in anh. DCM (1.0 mL) giving the product as a beige solid (45 mg, 30% yield). m.p. 109-110 °C. *v*: 3325, 3227, 3066, 2923, 1635, 1604, 1539, 1466, 1443, 1364, 1335, 1283, 1220, 1170, 1122, 1093, 1052, 895, 875, 775, 722, 689, 654 cm^−1. 1^H NMR (400 MHz, CDCl_3_) δ: 2.47 (s, 3H, 2’-C<u>H</u>_3_), 4.54 (d, *J* = 5.3 Hz, 2H, C<u>H</u>_2_), 6.44 (broad s, 1H, N<u>H</u>), 6.94 (d, *J* = 5.3 Hz, 1H, 4’-H), 7.07 (d, *J* = 5.2 Hz, 1H, 5’-H), 7.45 (d, *J* = 8.1 Hz, 1H, 4-H), 7.68 (dt, *J* = 8.6 Hz, *J’* = 2.0 Hz, 1H, 2-H), 7.79 (s, 1H, 6-H). ^13^C NMR (101 MHz, CDCl_3_) δ: 13.1 (C2’-<u>C</u>H_3_), 37.6 (CH_2_, <u>C</u>H_2_), 115.9 (dq, ^*2*^*J*_*CF*_ = 24.6 Hz, ^*3*^*J*_*CF*_ = 3.7 Hz, CH, C4), 118.1 (d, ^*2*^*J*_*CF*_ = 22.8 Hz, CH, C2), 119.6 (p, ^*3*^*J*_*CF*_ = 3.7 Hz, CH, C6), 122.5 (CH, C5’), 123.0 (qd, ^*1*^*J*_*CF*_ = 272.7 Hz, ^*4*^*J*_*CF*_ = 2.9 Hz, C, <u>C</u>F_3_), 128.5 (CH, C4’), 133.0 (C, C3’), 133.1 (qd, ^*2*^*J*_*CF*_ = 33.9 Hz, ^*3*^*J*_*CF*_ = 7.7 Hz, C, C5), 136.8 (C, C2’), 137.8 (d, ^*3*^*J*_*CF*_ = 6.9 Hz, C1), 162.6 (d, ^*1*^*J*_*CF*_ = 251.1 Hz, C, C3), 164.6 (d, ^*4*^*J*_*CF*_ = 2.3 Hz, C, <u>C</u>O). HRMS: Calcd for [C_14_H_11_F_4_NOS-H]^-^: 316.0425, found: 316.0431.

### 3-fluoro-*N*-(thiophen-3-ylmethyl)-5-(trifluoromethyl)benzamide, 47

Following general procedure C, 3-fluoro-5-(trifluoromethyl)benzoic acid (100 mg, 0.48 mmol) in anh. toluene (2.0 mL) and drops of DMF was reacted with thionyl chloride (349 µL, 572 mg, 4.81 mmol). Then, thiophen-3-ylmethanamine (65 mg, 0.58 mmol) and triethylamine (323 µL, 235 mg, 2.32 mmol) in anh. DCM (1.0 mL), were reacted with the crude acyl chloride (109 mg, 0.48 mmol) in anh. DCM (1.0 mL) giving the product as an off-white solid (65 mg, 45% yield). m.p. 75-76 °C. *v*: 3231, 3103, 2938, 1642, 1603, 1550, 1469, 1447, 1368, 1348, 1291, 1251, 1219, 1169, 1130, 1096, 1046, 1010, 945, 925, 886, 784, 691, 635, 563 cm^−1. 1^H NMR (400 MHz, CDCl_3_) δ: 4.61 (d, *J* = 5.7 Hz, 2H, C<u>H</u>_2_), 6.78 (broad s, 1H, N<u>H</u>), 7.06 (dd, *J* = 4.9 Hz, *J’* = 1.3 Hz, 1H, 4’-H), 7.19 (m, 1H, 2’-H), 7.31 (dd, *J* = 5.0 Hz, *J’* = 3.0 Hz, 1H, 5’-H), 7.44 (d, *J* = 8.1 Hz, 1H, 4-H), 7.68 (dt, *J* = 8.7 Hz, *J’* = 2.0 Hz, 1H, 2-H), 7.81 (s, 1H, 6-H). ^13^C NMR (101 MHz, CDCl_3_) δ: 39.6 (CH_2_, <u>C</u>H_2_), 115.9 (dq, ^*2*^*J*_*CF*_ = 24.6 Hz, ^*3*^*J*_*CF*_ = 3.7 Hz, CH, C4), 118.1 (d, ^*2*^*J*_*CF*_ = 22.8 Hz, CH, C2), 119.7 (p, ^*3*^*J*_*CF*_ = 3.7 Hz, CH, C6), 123.0 (qd, ^*1*^*J*_*CF*_ = 272.8 Hz, ^*4*^*J*_*CF*_ = 2.9 Hz, C, <u>C</u>F_3_), 123.1 (CH, C2’), 126.9 (CH, C5’), 127.5 (CH, C4’), 133.1 (qd, ^*2*^*J*_*CF*_ = 33.8 Hz, ^*3*^*J*_*CF*_ = 7.7 Hz, C, C5), 137.8 (d, ^*3*^*J*_*CF*_ = 6.9 Hz, C1), 138.2 (C, C3’), 162.6 (d, ^*1*^*J*_*CF*_ = 251.1 Hz, C, C3), 164.8 (d, ^*4*^*J*_*CF*_ = 2.3 Hz, C, <u>C</u>O). HRMS: Calcd for [C_13_H_9_F_4_NOS-H]^-^: 302.0268, found: 302.0270.

### 3-chloro-*N*-((5-methylthiophen-3-yl)methyl)-5-(trifluoromethyl)benzamide, 48

Following general procedure C, 3-chloro-5-(trifluoromethyl)benzoic acid (100 mg, 0.45 mmol) in anh. toluene (2.0 mL) and drops of DMF was reacted with thionyl chloride (163 µL, 268 mg, 2.25 mmol). Then, (5-methylthiophen-3-yl)methanamine (68 mg, 0.53 mmol) and triethylamine (148 µL, 107 mg, 1.06 mmol) in anh. DCM (1.0 mL), were reacted with the crude acyl chloride (109 mg, 0.45 mmol) in anh. DCM (1.0 mL) giving the product as a pale-yellow solid (87 mg, 58% yield). m.p. 130-131 °C. *v*: 3268, 1632, 1583, 1538, 1437, 1362, 1322, 1275, 1171, 1136, 1101, 1056, 886, 828, 756, 691, 648, 577, 560 cm^−1. 1^H NMR (400 MHz, CDCl_3_) δ: 2.44 (d, *J* = 1.1 Hz, 3H, 5’-C<u>H</u>_3_), 4.51 (d, *J* = 5.6 Hz, 2H, C<u>H</u>_2_), 6.71 (s, 1H, 4’-H), 6.75 (broad s, 1H, N<u>H</u>), 6.92 (s, 1H, 2’-H), 7.71 (s, 1H, 4-H), 7.91 (s, 1H, 6-H), 7.93 (s, 1H, 2-H). ^13^C NMR (101 MHz, CDCl_3_) δ: 15.4 (C5’-<u>C</u>H_3_), 39.8 (CH_2_, <u>C</u>H_2_), 120.7 (CH, C2’), 122.3 (q, ^*3*^*J*_*CF*_ = 3.7 Hz, CH, C6), 123.0 (q, ^*1*^*J*_*CF*_ = 273.2 Hz, C, <u>C</u>F_3_), 125.6 (CH, C4’), 128.4 (q, ^*3*^*J*_*CF*_ = 3.8 Hz, CH, C4), 130.8 (CH, C2), 132.7 (q, ^*2*^*J*_*CF*_ = 33.6 Hz, C, C5), 135.7 (C, C3), 137.1 (C, C1), 138.0 (C, C3’), 141.4 (C, C5’), 164.7 (C, <u>C</u>O). HRMS: Calcd for [C_14_H_11_ClF_3_NOS+H]^+^: 334.0275, found: 334.0282.

### 3-chloro-*N*-((2-methylthiophen-3-yl)methyl)-5-(trifluoromethyl)benzamide, 49

Following general procedure C, 3-chloro-5-(trifluoromethyl)benzoic acid (100 mg, 0.45 mmol) in anh. toluene (2.0 mL) and drops of DMF was reacted with thionyl chloride (323 µL, 530 mg, 4.45 mmol). Then, (2-methylthiophen-3-yl)methanamine hydrochloride (87 mg, 0.53 mmol) and triethylamine (298 µL, 217 mg, 2.14 mmol) in anh. DCM (1.0 mL), were reacted with the crude acyl chloride (108 mg, 0.45 mmol) in anh. DCM (1.0 mL) giving the product as a beige solid (81 mg, 55% yield). m.p. 117-118 °C. *v*: 3299, 3082, 2935, 1639, 1540, 1440, 1322, 1279, 1166, 1124, 887, 827, 770, 745, 721, 690, 658, 611 cm^−1. 1^H NMR (400 MHz, CDCl_3_) δ: 2.46 (s, 3H, 2’-C<u>H</u>_3_), 4.53 (d, *J* = 5.3 Hz, 2H, C<u>H</u>_2_), 6.55 (broad s, 1H, N<u>H</u>), 6.93 (d, *J* = 5.3 Hz, 1H, 4’-H), 7.06 (d, *J* = 5.2 Hz, 1H, 5’-H), 7.71 (m, 1H, 4-H), 7.89 (m, 1H, 6-H), 7.91 (t, *J* = 1.8 Hz, 1H, 2-H). ^13^C NMR (101 MHz, CDCl_3_) δ: 13.1 (C2’-<u>C</u>H_3_), 37.6 (CH_2_, <u>C</u>H_2_), 122.3 (q, ^*3*^*J*_*CF*_ = 3.7 Hz, CH, C6), 122.5 (CH, C5’), 123.0 (q, ^*1*^*J*_*CF*_ = 273.1 Hz, C, <u>C</u>F_3_), 128.4 (q, ^*3*^*J*_*CF*_ = 3.8 Hz, CH, C4), 128.5 (CH, C4’), 130.8 (CH, C2), 132.7 (q, ^*2*^*J*_*CF*_ = 33.6 Hz, C, C5), 133.0 (C, C3’), 135.7 (C, C3), 136.8 (C, C2’), 137.0 (C, C1), 164.7 (C, <u>C</u>O). HRMS: Calcd for [C_14_H_11_ClF_3_NOS-H]^-^: 332.0129, found: 332.0134.

### 3-chloro-*N*-(thiophen-3-ylmethyl)-5-(trifluoromethyl)benzamide, 50

Following general procedure C, 3-chloro-5-(trifluoromethyl)benzoic acid (100 mg, 0.45 mmol) in anh. toluene (2.0 mL) and drops of DMF was reacted with thionyl chloride (323 µL, 530 mg, 4.45 mmol). Then, thiophen-3-ylmethanamine (60 mg, 0.53 mmol) and triethylamine (149 µL, 108 mg, 1.07 mmol) in anh. DCM (1.0 mL), were reacted with the crude acyl chloride (108 mg, 0.45 mmol) in anh. DCM (1.0 mL) giving the product as a white solid (72 mg, 51% yield). m.p. 91-92 °C. *v*: 3246, 3091, 2927, 1635, 1549, 1429, 1349, 1324, 1284, 1175, 1124, 1050, 1004, 889, 826, 788, 738, 689, 637, 621 cm^−1. 1^H NMR (400 MHz, CDCl_3_) δ: 4.60 (d, *J* = 5.6 Hz, 2H, C<u>H</u>_2_), 6.84 (broad s, 1H, N<u>H</u>), 7.05 (dd, *J* = 4.9 Hz, *J’* = 1.3 Hz, 1H, 4’-H), 7.18 (m, 1H, 2’-H), 7.30 (dd, *J* = 5.0 Hz, *J’* = 3.0 Hz, 1H, 5’-H), 7.71 (m, 1H, 4-H), 7.90 (m, 1H, 6-H), 7.93 (t, *J* = 1.8 Hz, 1H, 2-H). ^13^C NMR (101 MHz, CDCl_3_) δ: 39.6 (CH_2_, <u>C</u>H_2_), 122.3 (q, ^*3*^*J*_*CF*_ = 3.7 Hz, CH, C6), 123.0 (q, ^*1*^*J*_*CF*_ = 273.2 Hz, C, <u>C</u>F_3_), 123.0 (CH, C2’), 126.8 (CH, C5’), 127.4 (CH, C4’), 128.5 (q, ^*3*^*J*_*CF*_ = 3.7 Hz, CH, C4), 130.8 (CH, C2), 132.7 (q, ^*2*^*J*_*CF*_ = 33.6 Hz, C, C5), 135.7 (C, C3), 137.0 (C, C1), 138.2 (C, C3’), 164.8 (C, <u>C</u>O). HRMS: Calcd for [C_14_H_11_ClF_3_NOS-H]^-^: 317.9973, found: 317.9975.

### 3,4-difluoro-*N*-((5-methylthiophen-3-yl)methyl)-5-(trifluoromethyl)benzamide, 51

Following general procedure C, 3,4-difluoro-5-(trifluoromethyl)benzoic acid (85 mg, 0.38 mmol) in anh. toluene (2.0 mL) and drops of DMF was reacted with thionyl chloride (276 µL, 452 mg, 3.80 mmol). Then, (5-methylthiophen-3-yl)methanamine hydrochloride (40 mg, 0.24 mmol) and triethylamine (175 µL, 127 mg, 1.26 mmol) in anh. DCM (1.0 mL) were reacted with the crude acyl chloride in anh. DCM (1.0 mL) giving the product as a white solid (59 mg, 72% yield). m.p. 93-94 °C. *v*: 3291, 3095, 2931, 1625, 1605, 1545, 1501, 1435, 1373, 1337, 1273, 1186, 1156, 1134, 1056, 1000, 966, 904, 895, 838, 772, 746, 739, 672, 641, 573 cm^−1. 1^H NMR (400 MHz, CDCl_3_) δ: 2.45 (d, *J* = 1.1 Hz, 3H, 5’-C<u>H</u>_3_), 4.51 (d, *J* = 5.6 Hz, 2H, C<u>H</u>_2_), 6.61 (broad s, 1H, N<u>H</u>), 6.71 (p, *J* = 1.1, 1H, 4’-H), 6.93 (m, 1H, 2’-H), 7.80 (m, 1H, 6-H), 7.86 (ddd, *J* = 9.6 Hz, *J’* = 7.0 Hz, *J’’* = 2.2 Hz, 1H, 2-H). ^13^C NMR (101 MHz, CDCl_3_) δ: 15.4 (C5’-<u>C</u>H_3_), 39.9 (CH_2_, <u>C</u>H_2_), 120.6 (d, ^*2*^*J*_*CF*_ = 18.6 Hz, CH, C2), 120.7 (m, CH, C6), 120.7 (m, C, C5), 120.8 (CH, C2’), 121.7 (qd, ^*1*^*J*_*CF*_ = 273.3 Hz, ^*3*^*J*_*CF*_ = 3.5 Hz, C, <u>C</u>F_3_), 125.6 (CH, C4’), 131.4 (t, ^*3*^*J*_*CF*_ = 4.8 Hz, C, C1), 137.8 (C, C3’), 141.6 (C, C5’), 150.2 (dd, ^*1*^*J*_*CF*_ = 263.7 Hz, ^*2*^*J*_*CF*_ = 13.1 Hz, C, C3), 150.8 (d, ^*1*^*J*_*CF*_ = 241.5 Hz, C, C4), 163.9 (C, <u>C</u>O). HRMS: Calcd for [C_14_H_10_F_5_NOS-H]^-^: 334.0330, found: 334.0333.

### 3-chloro-4-fluoro-*N*-((5-methylthiophen-3-yl)methyl)-5-(trifluoromethyl)benzamide, 52

Following general procedure C, 3-chloro-4-fluoro-5-(trifluoromethyl)benzoic acid (92 mg, 0.38 mmol) in anh. toluene (2.0 mL) and drops of DMF was reacted with thionyl chloride (276 µL, 452 mg, 3.80 mmol). Then, (5-methylthiophen-3-yl)methanamine hydrochloride (40 mg, 0.24 mmol) and triethylamine (175 µL, 127 mg, 1.26 mmol) in anh. DCM (1.0 mL) were reacted with the crude acyl chloride in anh. DCM (1.0 mL) giving the product as a white solid (56 mg, 65% yield). m.p. 127-128 °C. *v*: 3273, 3097, 2928, 1631, 1543, 1477, 1374, 1337, 1318, 1274, 1218, 1177, 1153, 1055, 1000, 967, 894, 832, 744, 684, 668, 638, 569 cm^−1. 1^H NMR (400 MHz, CDCl_3_) δ: 2.45 (d, *J* = 1.1 Hz, 3H, 5’-C<u>H</u>_3_), 4.51 (d, *J* = 5.5 Hz, 2H, C<u>H</u>_2_), 6.57 (broad s, 1H, N<u>H</u>), 6.71 (p, *J* = 1.1 Hz, 1H, 4’-H), 6.93 (m, 1H, 2’-H), 7.93 (dd, *J* = 5.9 Hz, *J’* = 1.5 Hz, 1H, 6-H), 8.05 (dd, *J* = 6.4 Hz, *J’* = 2.2 Hz, 1H, 2-H). ^13^C NMR (101 MHz, CDCl_3_) δ: 15.4 (C5’-<u>C</u>H_3_), 39.9 (CH_2_, <u>C</u>H_2_), 120.1 (qd, ^*2*^*J*_*CF*_ = 34.1 Hz, ^*2*^*J*_*CF*_ *=* 13.0 Hz, C, C5), 120.9 (CH, C2’), 121.7 (^*1*^*J*_*CF*_ *=* 273.0 Hz, C, <u>C</u>F_3_), 123.6 (d, ^*2*^*J*_*CF*_ = 17.6 Hz, C, C3), 124.5 (m, C, C6), 125.6 (CH, C4’), 131.4 (d, ^*4*^*J*_*CF*_ = 4.6 Hz, C, C1), 133.5 (C, C2), 137.8 (C, C3’), 141.6 (C, C5’), 157.4 (d, ^*1*^*J*_*CF*_ = 264.3 Hz, C, C4), 163.9 (C, <u>C</u>O). HRMS: Calcd for [C_14_H_10_ClF_4_NOS-H]^-^: 350.0035, found: 350.0043.

### 2,5-dichloro-*N*-((5-methylthiophen-3-yl)methyl)-3-(trifluoromethyl)benzamide, 53

Following general procedure C, 2,5-dichloro-3-(trifluoromethyl)benzoic acid (98 mg, 0.38 mmol) in anh. toluene (2.0 mL) and drops of DMF was reacted with thionyl chloride (276 µL, 452 mg, 3.80 mmol). Then, (5-methylthiophen-3-yl)methanamine hydrochloride (40 mg, 0.24 mmol) and triethylamine (175 µL, 127 mg, 1.26 mmol) in anh. DCM (1.0 mL) were reacted with the crude acyl chloride in anh. DCM (1.0 mL) giving the product as a white solid (45 mg, 50% yield). m.p. 152-153 °C. *v*: 3326, 3091, 2924, 1651, 1537, 1426, 1331, 1294, 1259, 1220, 1172, 1138, 1043, 1008, 891, 824, 735, 697, 638, 601, 583 cm^−1. 1^H NMR (400 MHz, CDCl_3_) δ: 2.47 (d, *J* = 1.1 Hz, 3H, 5’-C<u>H</u>_3_), 4.55 (d, *J* = 5.5 Hz, 2H, C<u>H</u>_2_), 6.22 (broad s, 1H, N<u>H</u>), 6.75 (p, *J* = 1.2 Hz, 1H, 4’-H), 6.96 (m, 1H, 2’-H), 7.68 (d, *J* = 2.6 Hz, 1H, 2-H), 7.73 (d, *J* = 2.6 Hz, 1H, 4-H). ^13^C NMR (101 MHz, CDCl_3_) δ: 15.5 (C5’-<u>C</u>H_3_), 39.9 (CH_2_, <u>C</u>H_2_), 120.9 (CH, C2’), 121.9 (q, ^*1*^*J*_*CF*_ = 273.2 Hz, C, <u>C</u>F_3_), 125.6 (CH, C4’), 127.5 (C, C6), 129.2 (q, ^*3*^*J*_*CF*_ = 5.6 Hz, CH, C4), 130.9 (q, ^*2*^*J*_*CF*_ = 32.0 Hz, C, C5), 132.6 (CH, C2), 133.6 (C, C3), 137.5 (C, C3’), 139.8 (C, C1), 141.6 (C, C5’), 164.4 (C, <u>C</u>O). HRMS: Calcd for [C_14_H_10_Cl_2_F_3_NOS+H]^+^: 367.9885, found: 367.9879.

### *N*-((2,5-dimethylfuran-3-yl)methyl)-3-fluoro-5-(trifluoromethyl)benzamide, 54

Following general procedure C, 3-fluoro-5-(trifluoromethyl)benzoic acid (100 mg, 0.48 mmol) in anh. toluene (2.0 mL) and drops of DMF was reacted with thionyl chloride (349 µL, 572 mg, 4.81 mmol). Then, (2,5-dimethylfuran-3-yl)methanamine (72 mg, 0.58 mmol) and triethylamine (323 µL, 235 mg, 2.32 mmol) in anh. DCM (1.0 mL), were reacted with the crude acyl chloride (109 mg, 0.48 mmol) in anh. DCM (1.0 mL) giving the product as a white solid (46 mg, 30% yield). m.p. 71-72 °C. *v*: 3312, 3095, 2921, 1647, 1604, 1557, 1443, 1348, 1294, 1249, 1174, 1142, 1094, 1038, 926, 887, 803, 753, 691, 621 cm^−1. 1^H NMR (400 MHz, CDCl_3_) δ: 2.22 (s, 3H, 5’-C<u>H</u>_3_), 2.25 (s, 3H, 2’-C<u>H</u>_3_), 4.33 (d, *J* = 5.2 Hz, 2H, C<u>H</u>_2_), 5.89 (s, 1H, 4’-H), 6.35 (broad s, 1H, N<u>H</u>), 7.44 (m, 1H, 4-H), 7.67 (dt, *J* = 8.7 Hz, *J’* = 2.1 Hz, 1H, 2-H), 7.78 (s, 1H, 6-H). ^13^C NMR (101 MHz, CDCl_3_) δ: 11.6 (C2’-<u>C</u>H_3_), 13.5 (C5’-<u>C</u>H_3_), 35.7 (CH_2_, <u>C</u>H_2_), 106.9 (CH, C4’), 115.8 (C, C3’), 115.8 (dq, ^*2*^*J*_*CF*_ = 24.6 Hz, ^*3*^*J*_*CF*_ = 3.8 Hz, CH, C4), 118.0 (d, ^*2*^*J*_*CF*_ = 22.8 Hz, CH, C2), 119.6 (p, ^*3*^*J*_*CF*_ = 3.7 Hz, CH, C6), 123.0 (qd, ^*1*^*J*_*CF*_ = 272.8 Hz, ^*4*^*J*_*CF*_ = 2.9 Hz, C, <u>C</u>F_3_), 133.1 (qd, ^*2*^*J*_*CF*_ = 33.7 Hz, ^*3*^*J*_*CF*_ = 7.6 Hz, C, C5), 137.9 (d, ^*3*^*J*_*CF*_ = 6.8 Hz, C1), 147.7 (C, C2’), 150.5 (C, C5’), 162.6 (d, ^*1*^*J*_*CF*_ = 251.0 Hz, C, C3), 164.7 (^*4*^*J*_*CF*_ = 2.3 Hz, C, <u>C</u>O). HRMS: Calcd for [C_15_H_13_F_4_NO_2_-H]^-^: 314.0810, found: 314.0819.

### 3-chloro-*N*-((2,5-dimethylfuran-3-yl)methyl)-5-(trifluoromethyl)benzamide, 55

Following general procedure C, 3-chloro-5-(trifluoromethyl)benzoic acid (100 mg, 0.45 mmol) in anh. toluene (2.0 mL) and drops of DMF was reacted with thionyl chloride (323 µL, 530 mg, 4.45 mmol). Then, (2,5-dimethylfuran-3-yl)methanamine (67 mg, 0.53 mmol) and triethylamine (147 µL, 107 mg, 1.06 mmol) in anh. DCM (1.0 mL), were reacted with the crude acyl chloride (109 mg, 0.45 mmol) in anh. DCM (1.0 mL) giving the product as a pale-yellow solid (86 mg, 58% yield). m.p. 88-89 °C. *v*: 3304, 3079, 2922, 2441, 1636, 1542, 1428, 1321, 1289, 1174, 1129, 1031, 924, 889, 794, 691, 624 cm^−1. 1^H NMR (400 MHz, CD^3^OD) δ: 2.17 (s, 3H, 5’-C<u>H</u>_3_), 2.24 (s, 3H, 2’-C<u>H</u>_3_), 4.26 (s, 2H, C<u>H</u>_2_), 5.92 (s, 1H, 4’-H), 7.85 (m, 1H, 4-H), 8.06 (m, 1H, 6-H), 8.08 (m, 1H, 2-H). ^13^C NMR (101 MHz, CD_3_OD) δ: 11.4 (C2’-<u>C</u>H_3_), 13.3 (C5’-<u>C</u>H_3_), 35.9 (CH_2_, <u>C</u>H_2_), 108.1 (CH, C4’), 117.8 (C, C3’), 123.7 (q, ^*3*^*J*_*CF*_ = 3.9 Hz, CH, C6), 124.5 (q, ^*1*^*J*_*CF*_ = 272.0 Hz, C, <u>C</u>F_3_), 129.1 (q, ^*3*^*J*_*CF*_ = 3.8 Hz, CH, C4), 132.2 (CH, C2), 133.5 (q, ^*2*^*J*_*CF*_ = 33.3 Hz, C, C5), 136.5 (C, C3), 138.7 (C, C1), 148.3 (C, C2’), 150.9 (C, C5’), 166.5 (C, <u>C</u>O). HRMS: Calcd for [C_15_H_13_ClF_3_NO_2_-H]^-^: 330.0514, found: 330.0519.

### 3-fluoro-*N*-(furan-3-ylmethyl)-5-(trifluoromethyl)benzamide, 56

Following general procedure D, 3-fluoro-5-(trifluoromethyl)benzoic acid (100 mg, 0.48 mmol) in anh. toluene (2.0 mL) and drops of DMF was reacted with thionyl chloride (346 µL, 567 mg, 4.81 mmol). Then, furan-3-ylmethanamine (51 mg, 0.53 mmol) and triethylamine (268 µL, 194 mg, 1.92 mmol) in anh. DCM (1.0 mL), were reacted with the crude acyl chloride in anh. DCM (1.0 mL) giving the product as a reddish syrup (67 mg, 49% yield). *v*: 3295, 3087, 1645, 1604, 1544, 1467, 1445, 1341, 1285, 1249, 1219, 1171, 1128, 1093, 1021, 967, 925, 884, 874, 768, 692, 619, 599 cm^−1. 1^H NMR (400 MHz, CDCl_3_) δ: 4.50 (d, *J* = 5.5 Hz, 2H, C<u>H</u>_2_), 6.38 (broad s, 1H, N<u>H</u>), 6.43 (m, 1H, 4’-H), 7.42 (t, *J* = 1.7 Hz, 1H, 2’-H), 7.45-7.48 (complex signal, 2H, 4-H, 5’-H), 7.69 (dt, *J* = 8.6 Hz, *J’* = 2.0 Hz, 1H, 2-H), 7.79 (s, 1H, 6-H). ^13^C NMR (101 MHz, CDCl_3_) δ: 35.4 (CH_2_, <u>C</u>H_2_), 110.4 (CH, C4’), 116.0 (dq, ^*2*^*J*_*CF*_ = 24.4 Hz, ^*3*^*J*_*CF*_ = 3.7 Hz, CH, C4), 118.1 (d, ^*2*^*J*_*CF*_ = 22.9 Hz, CH, C2), 119.6 (p, ^*3*^*J*_*CF*_ = 3.8 Hz, CH, C6), 121.6 (C, C3’), 123.0 (q, ^*1*^*J*_*CF*_ = 271.5 Hz, C, <u>C</u>F_3_), 133.2 (qd, ^*2*^*J*_*CF*_ = 33.9 Hz, ^*3*^*J*_*CF*_ = 7.3 Hz, C, C5), 137.8 (d, ^*3*^*J*_*CF*_ = 6.8 Hz, C, C1), 140.7 (CH, C5’), 143.9 (CH, C2’), 162.7 (d, _*1*_*J*_*CF*_ = 251.2 Hz, C, C3), 164.8 (C, <u>C</u>O). HRMS: Calcd for [C_13_H_9_F_4_NO_2_-H]^-^: 286.0497, found: 286.0494.

### 3-fluoro-*N*-(thiazol-5-ylmethyl)-5-(trifluoromethyl)benzamide, 57

Following general procedure D, 3-fluoro-5-(trifluoromethyl)benzoic acid (100 mg, 0.48 mmol) in anh. toluene (2.0 mL) and drops of DMF was reacted with thionyl chloride (279 µL, 457 mg, 3.84 mmol). Then, thiazol-5-ylmethanamine (60 mg, 0.53 mmol) and triethylamine (268 µL, 194 mg, 1.92 mmol) in anh. DCM (1.0 mL), were reacted with the crude acyl chloride in anh. DCM (1.0 mL) giving the product as a reddish syrup (51 mg, 35% yield). *v*: 3269, 3080, 1646, 1604, 1543, 1520, 1467, 1446, 1408, 1348, 1280, 1220, 1169, 1126, 1093, 1039, 1003, 926, 881, 797, 752, 692, 633, 601 cm^−1. 1^H NMR (400 MHz, CDCl_3_) δ: 4.86 (dd, *J* = 5.9 Hz, *J’* = 0.9 Hz, 2H, C<u>H</u>_2_), 6.85 (broad m, 1H, N<u>H</u>), 7.48 (dt, *J* = 8.0 Hz, *J’* = 1.7 Hz, 1H, 4-H), 7.72 (dt, *J* = 8.5 Hz, *J’* = 2.1 Hz, 1H, 2-H), 7.81 (d, *J* = 0.8 Hz, 1H, 4’-H), 7.82 (s, 1H, 6-H), 8.76 (s, 1H, 2’-H). ^13^C NMR (101 MHz, CDCl_3_) δ: 36.3 (CH_2_, <u>C</u>H_2_), 116.3 (dq, ^*2*^*J*_*CF*_ = 24.4 Hz, ^*3*^*J*_*CF*_ = 3.7 Hz, CH, C4), 118.2 (d, ^*2*^*J*_*CF*_ = 23.0 Hz, CH, C2), 119.7 (p, ^*3*^*J*_*CF*_ = 3.7 Hz, CH, C6), 122.9 (q, ^*1*^*J*_*CF*_ = 272.7 Hz, C, <u>C</u>F_3_), 133.3 (qd, ^*2*^*J*_*CF*_ = 34.0 Hz, ^*3*^*J*_*CF*_ = 7.7 Hz, C, C5), 135.0 (C, C5’), 137.2 (d, ^*3*^*J*_*CF*_ = 6.9 Hz, C, C1), 142.5 (CH, C4’), 154.1 (CH, C2’), 162.7 (d, ^*1*^*J*_*CF*_ = 251.5 Hz, C, C3), 164.8 (C, <u>C</u>O). HRMS: Calcd for [C_12_H_8_F_4_N_2_OS+H]^+^: 285.0304, found: 285.0312.

### 3-fluoro-*N*-(isoxazol-3-ylmethyl)-5-(trifluoromethyl)benzamide, 58

Following general procedure D, 3-fluoro-5-(trifluoromethyl)benzoic acid (78 mg, 0.37 mmol) in anh. toluene (1.5 mL) and drops of DMF was reacted with thionyl chloride (272 µL, 446 mg, 3.75 mmol). Then, isoxazol-3-ylmethylamine (37 mg, 0.37 mmol) and triethylamine (209 µL, 152 mg, 1.50 mmol) in anh. DCM (1.0 mL), were reacted with the crude acyl chloride in anh. DCM (1.0 mL) giving the product as a brownish solid (34 mg, 32% yield). m.p. 110-111 °C. *v*: 267, 3114, 1670, 1630, 1606, 1573, 1550, 1501, 1467, 1438, 1422, 1354, 1293, 1245, 1222, 1162, 1128, 1093, 1059, 1044, 1020, 1003, 938, 888, 862, 795, 713, 692, 651, 592, 555 cm^−1. 1^H NMR (400 MHz, CDCl_3_) δ: 4.77 (d, *J* = 5.6 Hz, 2H, C<u>H</u>_2_), 6.43 (d, *J* = 1.7 Hz, 1H, 4’-H), 6.97 (broad s, 1H, N<u>H</u>), 7.49 (m, 1H, 4-H), 7.74 (dt, *J* = 8.5 Hz, *J’* = 1.8 Hz, 1H, 2-H), 7.86 (s, 1H, 6-H), 8.41 (d, *J* = 1.6 Hz, 1H, 5’-H). ^13^C NMR (101 MHz, CDCl_3_) δ: 36.3 (CH_2_, <u>C</u>H_2_), 104.1 (CH, C4’), 116.3 (dq, ^*2*^*J*_*CF*_ = 24.6 Hz, ^*3*^*J*_*CF*_ = 3.8 Hz, CH, C4), 118.2 (d, ^*2*^*J*_*CF*_ = 22.9 Hz, CH, C2), 119.8 (p, ^*3*^*J*_*CF*_ = 3.8 Hz, CH, C6), 122.9 (q, ^*1*^*J*_*CF*_ = 272.7 Hz, C, <u>C</u>F_3_), 133.3 (qd, ^*2*^*J*_*CF*_ = 34.0 Hz, ^*3*^*J*_*CF*_ = 7.7 Hz, C, C5), 137.2 (d, ^*3*^*J*_*CF*_ = 6.9 Hz, C, C1), 159.4 (CH, C5’), 159.7 (C, C3’), 162.7 (d, ^*1*^*J*_*CF*_ = 251.3 Hz, C, C3), 165.0 (C, <u>C</u>O). HRMS: Calcd for [C_12_H_8_F_4_N_2_O_2_-H]^-^: 287.0449, found: 287.0446.

### 3-fluoro-*N*-(pyridin-4-ylmethyl)-5-(trifluoromethyl)benzamide, 59

Following general procedure D, 3-fluoro-5-(trifluoromethyl)benzoic acid (100 mg, 0.48 mmol) in anh. toluene (2.0 mL) and drops of DMF was reacted with thionyl chloride (346 µL, 567 mg, 4.81 mmol). Then, pyridin-4-ylmethanamine (54 µL, 57 mg, 0.53 mmol) and triethylamine (268 µL, 194 mg, 1.92 mmol) in anh. DCM (1.0 mL), were reacted with the crude acyl chloride in anh. DCM (1.0 mL) giving the product as a pale-yellow syrup (81 mg, 57% yield). *v*: 3282, 3061, 1650, 1602, 1543, 1467, 1445, 1417, 1351, 1325, 1283, 1219, 1169, 1126, 1094, 1059, 1002, 928, 883, 799, 754, 691, 615 cm^−1. 1^H NMR (400 MHz, CDCl_3_) δ: 4.65 (d, *J* = 6.0 Hz, 2H, C<u>H</u>_2_), 7.12 (broad s, 1H, N<u>H</u>), 7.23 [d, *J* = 6.1 Hz, 2H, 2’(6’)-H], 7.49 (m, 1H, 4-H), 7.76 (dt, *J* = 8.6 Hz, *J’* = 2.1 Hz, 1H, 2-H), 7.86 (s, 1H, 6-H), 8.53 [d, *J* = 6.1 Hz, 2H, 3’(5’)-H]. ^13^C NMR (101 MHz, CDCl_3_) δ: 43.2 (CH_2_, <u>C</u>H_2_), 116.3 (dq, ^*2*^*J*_*CF*_ = 24.3 Hz, ^*3*^*J*_*CF*_ = 3.7 Hz, CH, C4), 118.3 (d, ^*2*^*J*_*CF*_ = 22.8 Hz, CH, C2), 119.7 (p, ^*3*^*J*_*CF*_ = 4.0 Hz, CH, C6), 122.6 [CH, C2’(6’)], 122.9 (qd, ^*1*^*J*_*CF*_ = 272.9 Hz, ^*3*^*J*_*CF*_ = 2.9 Hz, C, <u>C</u>F_3_), 133.3 (qd, ^*2*^*J*_*CF*_ = 33.7 Hz, ^*3*^*J*_*CF*_ = 7.6 Hz, C, C5), 137.3 (d, ^*3*^*J*_*CF*_ = 6.7 Hz, C, C1), 146.9 (C, C1’), 150.2 [CH, C3’(5’)], 162.7 (d, ^*1*^*J*_*CF*_ = 251.5 Hz, C, C3), 165.2 (d, ^*4*^*J*_*CF*_ = 2.4 Hz, C, <u>C</u>O). HRMS: Calcd for [C_14_H_10_F_4_N_2_O+H]_+_: 299.0802, found: 299.0805.

### *N*-((1*H*-pyrazol-3-yl)methyl)-3-fluoro-5-(trifluoromethyl)benzamide, 60

Following general procedure D, 3-fluoro-5-(trifluoromethyl)benzoic acid (100 mg, 0.48 mmol) in anh. toluene (2.0 mL) and drops of DMF was reacted with thionyl chloride (279 µL, 457 mg, 3.84 mmol). Then, (1*H*-pyrazol-3-yl)methanamine (51 mg, 0.53 mmol) and triethylamine (201 µL, 146 mg, 1.44 mmol) in anh. DCM (1.0 mL), were reacted with the crude acyl chloride in anh. DCM (1.0 mL) giving the product as a pale-yellow solid (75 mg, 54% yield). m.p. 130-131 °C. *v*: 3197, 3069, 2945, 1645, 1599, 1556, 1471, 1441, 1428, 1343, 1297, 1240, 1212, 1174, 1131, 1096, 1051, 1035, 1002, 921, 692, 859, 790, 765, 748, 716, 664, 615 cm_−1. 1_H NMR (400 MHz, CDCl_3_) δ: 4.70 (d, *J* = 5.6 Hz, 2H, C<u>H</u>_2_), 6.30 (d, *J* = 2.2 Hz, 1H, 4’-H), 7.31 (broad s, 1H, N<u>H</u>), 7.45 (m, 1H, 4-H), 7.54 (d, *J* = 2.2 Hz, 1H, 5’-H), 7.74 dt, *J* = 9.2 Hz, *J’* = 2.2 Hz, 1H, 2-H), 7.85 (td, *J* = 1.6 Hz, *J’* = 0.8 Hz, 1H, 6-H). ^13^C NMR (101 MHz, CDCl_3_) δ: 37.1 (CH_2_, <u>C</u>H_2_), 104.5 (CH, C4’), 115.8 (dq, ^*2*^*J*_*CF*_ = 24.5 Hz, ^*3*^*J*_*CF*_ = 4.0 Hz, CH, C4), 118.1 (d, ^*2*^*J*_*CF*_ = 22.9 Hz, CH, C2), 119.7 (m, CH, C6), 122.8 (qd, ^*1*^*J*_*CF*_ = 270.5 Hz, ^*3*^*J*_*CF*_ = 2.8 Hz, C, <u>C</u>F_3_), 132.3 (CH, C5’), 132.9 (qd, ^*2*^*J*_*CF*_ = 33.9 Hz, ^*3*^*J*_*CF*_ = 7.5 Hz, C, C5), 137.4 (d, ^*3*^*J*_*CF*_ = 6.9 Hz, C, C1), 146.3 (C, C3’), 162.4 (d, ^*1*^*J*_*CF*_ = 250.9 Hz, C, C3), 165.0 (d, *J* = 2.2 Hz, C, <u>C</u>O). HRMS: Calcd for [C_12_H_9_F_4_N_3_O-H]^-^: 286.0609, found: 286.0605.

### 3-fluoro-*N*-(oxazol-4-ylmethyl)-5-(trifluoromethyl)benzamide, 61

Following general procedure D, 3-fluoro-5-(trifluoromethyl)benzoic acid (100 mg, 0.48 mmol) in anh. toluene (2.0 mL) and drops of DMF was reacted with thionyl chloride (279 µL, 457 mg, 3.84 mmol). Then, oxazol-4-ylmethanamine hydrochloride (71 mg, 0.53 mmol) and triethylamine (268 µL, 194 mg, 1.92 mmol) in anh. DCM (1.0 mL), were reacted with the crude acyl chloride in anh. DCM (1.0 mL) giving the product as a yellowish solid (107 mg, 77% yield). m.p. 99-100 °C. *v*: 3272, 3117, 1659, 1608, 1547, 1506, 1470, 1442, 1427, 1348, 1328, 1295, 1280, 1247, 1232, 1210, 1171, 1122, 1095, 1068, 1045, 1004, 968, 923, 916, 881, 859, 796, 774, 748, 690, 664, 621 cm^−1. 1^H NMR (400 MHz, CDCl_3_) δ: 4.56 (d, *J* = 5.1 Hz, 2H, C<u>H</u>_2_), 7.28 (broad s, 1H, N<u>H</u>), 7.43 (m, 1H, 4-H), 7.68-7.72 (complex signal, 2H, 2-H, 2’-H), 7.82 (s, 1H, 6-H), 7.86 (s, 1H, 5’-H). ^13^C NMR (101 MHz, CDCl_3_) δ: 35.8 (CH_2_, <u>C</u>H_2_), 116.0 (dq, ^*2*^*J*_*CF*_ = 24.5 Hz, ^*3*^*J*_*CF*_ = 3.9 Hz, CH, C4), 118.2 (d, ^*2*^*J*_*CF*_ = 22.7 Hz, CH, C2), 119.8 (p, ^*3*^*J*_*CF*_ = 3.7 Hz, CH, C6), 122.9 (qd, ^*1*^*J*_*CF*_ = 273.0 Hz, ^*3*^*J*^*CF*^ = 3.1 Hz, C, <u>C</u>F_3_), 133.1 (qd, ^*2*^*J*_*CF*_ = 33.8 Hz, ^*3*^*J*_*CF*_ = 7.7 Hz, C, C5), 136.4 (CH, C2’), 136.4 (CH, C3’), 137.5 (d, ^*3*^*J*_*CF*_ = 6.9 Hz, C, C1), 151.6 (CH, C5’), 162.6 (d, ^*1*^*J*_*CF*_ = 251.0 Hz, C, C3), 164.9 (d, ^*4*^*J*_*CF*_ = 2.4 Hz, C, <u>C</u>O). HRMS: Calcd for [C_12_H_8_F_4_N_2_O_2_-H]^-^: 287.0449, found: 287.0445.

### 3-fluoro-*N*-(thiazol-4-ylmethyl)-5-(trifluoromethyl)benzamide, 62

Following general procedure D, 3-fluoro-5-(trifluoromethyl)benzoic acid (100 mg, 0.48 mmol) in anh. toluene (2.0 mL) and drops of DMF was reacted with thionyl chloride (279 µL, 457 mg, 3.84 mmol). Then, thiazol-4-ylmethanamine dihydrochloride (99 mg, 0.53 mmol) and triethylamine (268 µL, 194 mg, 1.92 mmol) in anh. DCM (1.0 mL), were reacted with the crude acyl chloride in anh. DCM (1.0 mL) giving the product as a white solid (61 mg, 42% yield). m.p, 98-99 °C. *v*: 3268, 3138, 3069, 1664, 1611, 1543, 1520, 1471, 1428, 1407, 1348, 1289, 1247, 1209, 1170, 1123, 1093, 1041, 1002, 943, 921, 882, 859, 825, 765, 750, 729, 689, 643, 584 cm^−1. 1^H NMR (400 MHz, CDCl_3_) δ: 4.77 (dd, *J* = 5.5 Hz, *J’* = 0.7 Hz, 2H, C<u>H</u>_2_), 7.31 (dt, *J* = 2.0 Hz, *J’* = 0.8 Hz, 1H, 2’-H), 7.40 (broad m, 1H, N<u>H</u>), 7.43 (m, 1H, 4-H), 7.72 (dt, *J* = 8.7 Hz, *J’* = 2.1 Hz, 1H, 2-H), 7.84 (td, *J* = 1.6 Hz, *J’* = 0.8 Hz, 1H, 6-H), 8.77 (d, *J* = 2.0 Hz, 1H, 5’-H). ^13^C NMR (101 MHz, CDCl_3_) δ: 40.1 (CH_2_, <u>C</u>H_2_), 115.9 (dq, ^*2*^*J*_*CF*_ = 24.8 Hz, ^*3*^*J*_*CF*_ = 3.7 Hz, CH, C4), 116.1 (CH, C2’), 118.2 (d, ^*2*^*J*_*CF*_ = 22.7 Hz, CH, C2), 119.8 (p, ^*3*^*J*_*CF*_ = 3.6 Hz, CH, C6), 123.0 (qd, ^*1*^*J*_*CF*_ = 272.7 Hz, ^*3*^*J*_*CF*_ = 2.8 Hz, C, <u>C</u>F_3_), 133.1 (qd, ^*2*^*J*_*CF*_ = 33.9 Hz, ^*3*^*J*_*CF*_ = 7.7 Hz, C, C5), 137.6 (d, ^*3*^*J*_*CF*_ = 6.8 Hz, C, C1), 153.0 (C, C3’), 153.7 (CH, C5’), 162.6 (d, ^*1*^*J*_*CF*_ = 251.0 Hz, C, C3), 164.8 (d, ^*4*^*J*_*CF*_ = 2.4 Hz, C, <u>C</u>O). Anal. calcd for C_12_H_8_F_4_N_2_OS: C 47.37, H 2.65, N 9.21. HRMS: Calcd for [C_12_H_8_F_4_N_2_OS-H]^-^: 303.0221, found: 303.0219.

### (2,5-dimethylthiophen-3-yl)methyl 3-fluoro-5-(trifluoromethyl)benzoate, 63

To a solution of (2,5-dimethylthiophen-3-yl)methanol (88 mg, 0.62 mmol), EDC·HCl (178 mg, 0.93 mmol) and DMAP (30 mg, 0.25 mmol) in anh. DCM (2.0 mL), 3-fluoro-5-(trifluoromethyl)benzoic acid (155 mg, 0.74 mmol) was added under Ar atmosphere and the mixture was stirred at RT for 16 h. Then, DCM (10 mL) was added and the mixture was washed with brine (2 × 20 mL). The organic layer was dried over anh. Na_2_SO_4_, filtered and concentrated *in vacuo*. The resulting crude was purified by column chromatography in silica gel (using as eluent mixtures of EtOAc in hexane from 0% to 10%) to obtain the product as a colorless oil (96 mg, 47% yield). *v*: 2923, 1727, 1606, 1453, 1363, 1347, 1249, 1238, 1202, 1171, 1131, 1103, 1091, 959, 906, 887, 828, 768, 692, 582 cm^−1. 1^H NMR (400 MHz, CDCl_3_) δ: 2.42 (m, 3H, 5’-C<u>H</u>_3_), 2.46 (s, 3H, 2’-C<u>H</u>_3_), 5.24 (s, 2H, C<u>H</u>_2_), 6.69 (d, *J* = 1.2 Hz, 1H, 4’-H), 7.51 (dddd, *J* = 8.1 Hz, *J’* = 2.5 Hz, *J’’* = 1.6 Hz, *J’’’* = 0.7 Hz, 1H, 4-H), 7.91 (ddd, *J* = 8.7 Hz, *J’* = 2.6 Hz, *J’’* = 1.4 Hz, 1H, 2-H), 8.11 (s, 1H, 6-H). ^13^C NMR (101 MHz, CDCl_3_) δ: 13.0 (C2’-<u>C</u>H_3_), 15.2 (C5’-<u>C</u>H_3_), 61.0 (CH_2_, <u>C</u>H_2_), 117.3 (dq, ^*2*^*J*_*CF*_ = 24.6 Hz, ^*3*^*J*_*CF*_ = 3.7 Hz, CH, C4), 120.2 (d, ^*2*^*J*_*CF*_ = 23.0 Hz, CH, C2), 122.5 (p, ^*3*^*J*_*CF*_ = 3.9 Hz, CH, C6), 123.0 (qd, ^*1*^*J*_*CF*_ = 272.8 Hz, ^*4*^*J*_*CF*_ = 2.9 Hz, C, <u>C</u>F_3_), 126.9 (CH, C4’), 131.0 (C, C3’), 133.0 (qd, ^*2*^*J*_*CF*_ = 33.9 Hz, ^*3*^*J*_*CF*_ = 7.5 Hz, C, C5), 133.7 (d, ^*3*^*J*_*CF*_ = 7.6 Hz, C1), 136.5 (C, C2’), 136.8 (C, C5’), 162.4 (d, ^*1*^*J*_*CF*_ = 250.4 Hz, C, C3), 164.2 (d, ^*4*^*J*_*CF*_ = 2.3 Hz, C, <u>C</u>O). HRMS: Calcd for [C_15_H_12_F_4_O_2_S+Na]^+^: 355.0386, found: 355.0389.

### 1-(2,5-dimethylthiophen-3-yl)-*N*-(3-fluoro-5-(trifluoromethyl)benzyl)methanamine, 64

To a solution 1- (bromomethyl)-3-fluoro-5-(trifluoromethyl)benzene (50 mg, 0.19 mmol) and triethylamine (81 µL, 59 mg, 0.58 mmol) in DMF (1.0 mL) was added (2,5-dimethylthiophen-3-yl)methanamine hydrochloride (35 mg, 0.19 mmol) and the mixture was kept under stirring at RT for 16 h. Water (15 mL) followed by EtOAc (10 mL) were added and the mixture was extracted. The organic layer was washed again with brine (15 mL) and then it was dried over anh. Na_2_SO_4_ and filtered. Solvents were concentrated *in vacuo* and the resulting crude was purified by column chromatography in silica gel (using as eluent mixtures of EtOAc in hexane from 0% to 15%) to afford the product as a colorless oil (27 mg, 45% yield). *v*: 2921, 2860, 1605, 1452, 1342, 1227, 1166, 1125, 1091, 975, 869, 830, 761, 721, 698, 712 cm_−1. 1_H NMR (400 MHz, CDCl_3_) δ: 2.30 (s, 3H, 2’-C<u>H</u>_3_), 2.40 (m, 3H, 5’-C<u>H</u>_3_), 3.62 (s, 2H, thio-C<u>H</u>_2_), 3.84 (s, 2H, aryl-C<u>H</u>_2_), 6.59 (q, *J* = 1.2 Hz, 1H, 4’-H), 7.20 (m, 1H, 4-H), 7.29 (dddd, *J* = 9.4 Hz, *J’* = 2.6 Hz, *J’’* = 1.4 Hz, *J’’’* = 0.7 Hz, 1H, 2-H), 7.41 (tq, *J* = 1.5 Hz, *J’* = 0.7 Hz, 1H, 6-H). ^13^C NMR (101 MHz, CDCl_3_) δ: 12.9 (C2’-<u>C</u>H_3_), 15.2 (C5’-<u>C</u>H_3_), 46.1 (CH_2_, thio-<u>C</u>H_2_), 52.3 (CH_2_, aryl-<u>C</u>H_2_), 111.4 (dq, ^*2*^*J*_*CF*_ = 24.6 Hz, ^*3*^*J*_*CF*_ = 3.8 Hz, CH, C4), 118.4 (d, ^*2*^*J*_*CF*_ = 21.3 Hz, CH, C2), 120.6 (p, ^*3*^*J*_*CF*_ = 3.6 Hz, CH, C6), 123.5 (qd, ^*1*^*J*_*CF*_ = 272.5 Hz, ^*3*^*J*_*CF*_ *=* 3.1 Hz, C, <u>C</u>F_3_), 126.7 (CH, C4’), 132.4 (qd, ^*2*^*J*_*CF*_ = 33.0 Hz, ^*3*^*J*_*CF*_ = 8.1 Hz, C, C5), 132.9 (C, C3’), 135.4 (C, C2’),, 136.0 (C, C5’), 144.8 (d, ^*3*^*J*_*CF*_ = 7.1 Hz, C, C1), 162.7 (d, ^*1*^*J*_*CF*_ = 248.2 Hz, C, C3). HRMS: Calcd for [C_15_H_15_F_4_NS+H]^+^: 318.0934, found: 318.0934.

### *N*-(3-fluoro-5-(trifluoromethyl)benzyl)-2,5-dimethylthiophene-3-carboxamide, 65

Following general procedure D, 2,5-dimethylthiophene-3-carboxylic acid (100 mg, 0.64 mmol) in anh. toluene (3.0 mL) and drops of DMF was reacted with thionyl chloride (279 µL, 457 mg, 3.84 mmol). Then, (3-fluoro-5-(trifluoromethyl)phenyl)methanamine (148 mg, 0.77 mmol) and triethylamine (268 µL, 194 mg, 1.92 mmol) in anh. DCM (2.0 mL) reacted with the crude acyl chloride in anh. DCM (1.0 mL) giving the product as a beige solid (85 mg, 40% yield). m.p. 105-106 °C. *v*: 3293, 2925, 1634, 1609, 1563, 1515, 1454, 1424, 1351, 1341, 1315, 1280, 1229, 1169, 1120, 1084, 1041, 994, 973, 875, 852, 834, 785, 768, 716, 701, 690, 650, 601 cm^−1. 1^H NMR (400 MHz, CDCl_3_) δ: 2.37 (s, 3H, 2’-C<u>H</u>_3_), 2.63 (s, 3H, 5’-C<u>H</u>_3_), 4.59 (d, *J* = 6.1 Hz, 2H, C<u>H</u>_2_), 6.35 (broad s, 1H, N<u>H</u>), 6.76 (d, *J* = 1.2 Hz, 1H, 4’-H), 7.22 (m, 2H, 2-H, 4-H), 7.35 (s, 1H, 6-H). ^13^C NMR (101 MHz, CDCl_3_) δ: 15.0 (C2’-<u>C</u>H_3_), 15.1 (C5’-<u>C</u>H_3_), 42.8 (d, ^*4*^*J*_*CF*_ = 1.7 Hz, CH_2_, C<u>H</u>_2_), 111.9 (dq, ^*2*^*J*_*CF*_ = 24.6 Hz, ^*3*^*J*_*CF*_ = 3.9 Hz, CH, C4), 118.1 (d, ^*2*^*J*_*CF*_ = 21.9 Hz, CH, C2), 120.1 (m, CH, C6), 123.3 (qd, ^*1*^*J*_*CF*_ = 272.5 Hz, ^*3*^*J*_*CF*_ *=* 3.3 Hz, C, <u>C</u>F_3_), 123.9 (CH, C4’), 130.6 (C, C3’), 132.9 (qd, ^*2*^*J*_*CF*_ = 33.3 Hz, ^*3*^*J*_*CF*_ = 8.2 Hz, C, C5), 136.5 (C, C5’), 142.9 (d, ^*3*^*J*_*CF*_ = 7.2 Hz, C, C1), 143.8 (C, C2’), 162.8 (d, ^*1*^*J*_*CF*_ = 249.4 Hz, C, C3), 164.7 (C, <u>C</u>O). HRMS: Calcd for [C_15_H_13_F_4_NOS-H]^-^: 330.0581, found: 330.0573.

### Virological experiments

#### Cells and viruses

Madin-Darby canine kidney (MDCK) cells were a kind gift from M. Matrosovich (Philipps University, Marburg). Human embryonic kidney (HEK293T) cells were purchased from Thermo Scientific and HeLa cells were from ATCC (CCL-2). The virus panel consisted of: A/PR/8/34 [A/H1N1; reverse-engineered from the plasmids which were kindly provided by M. Kim (Korea Research Institute of Chemical Technology); hereunder abbreviated as PR8]; A/Virginia/ATCC3/2009 [A/H1N1; ATCC VR-1738; hereunder abbreviated as Virg09]; A/Victoria/361/11 [A/H3N2; a kind gift from G. Rimmelzwaan (Erasmus Medical Center, Rotterdam)]; and B/Ned/537/05 (Yamagata lineage; a kind gift from R. Fouchier (Erasmus Medical Center, Rotterdam)]. The virus stocks were prepared in MDCK cells or embryonated hen eggs, and titrated by the 50% cell culture infective dose (CCID_50_) method.

#### Assessment of antiviral activity based on reduction of viral cytopathic effect (CPE)

The infection medium consisted of UltraMDCK medium (Lonza), supplemented with 225 mg/L sodium bicarbonate, 2 mM L-glutamine and 2 µg/mL TPCK (tosylphenylalanylchloromethyl-keton)-treated trypsin (Sigma-Aldrich). Our detailed method can be found elsewhere.^56^ In short, MDCK cells were seeded at 7,500 cells per well in 96-well plates. On the next day, the virus was added at an MOI of 50 CCID_50_ per well, immediately followed by serial dilutions of the test compounds.

After three days incubation at 35 °C, microscopy was performed to score virus-induced CPE and compound cytotoxicity. Next, the CellTiter 96^®^ AQ_ueous_ MTS Reagent (Promega) was added to the cells and, four hours later, the absorbance at 490 nm was measured in a plate reader. The compounds’ antiviral activity was expressed as the half-maximal effective concentration (EC_50_) in the MTS or microscopic scoring assay (see ref. 46 for calculation details]. Cytotoxicity was expressed as the CC_50_, i.e. 50% cytotoxic concentration by MTS assay and the MCC (minimum cytotoxic concentration), i.e. concentration producing minimal changes in cell morphology.

To assess antiviral activity in human lung epithelium-derived Calu-3 cells^57^, the cells were seeded at 30,000 cells per well in black 96-well plates. On the next day, 100 CCID_50_ of Virg09 virus was added together with compound. The inoculum was removed 2 h later, with addition of fresh compound. After three days incubation at 35 °C, the cells were fixed with 2% paraformaldehyde; permeabilized with 0.1% Triton X-100; and stained with anti-NP antibody (1:1,000 of #ab20343 from Abcam) followed by goat anti-mouse IgG Alexa Fluor 488 (1:500 of #A11001 from Invitrogen); and nuclear staining with Hoechst (Thermo Fisher Scientific). The % NP-positive cells was quantified via high-content imaging using a CellInsight CX5 instrument (Thermo Scientific).

#### Selection of resistant influenza viruses by serial passaging

MDCK cells were infected with Virg09 virus as above, and exposed to different concentrations of VF-57a or RL-007. Three days later, all wells were inspected to select the highest compound concentrations at which virus-induced CPE was visible, and freeze these supernatants at -80 °C. The harvests were further passaged under gradually higher compound concentrations, until resistance was reached (i.e. virus breakthrough at 40 µM of compound). A no compound control was passaged in parallel. Virus clones were obtained by plaque purification under 0.6 % agarose and 4 µM VF-57a or 10 µM RL-007. After expansion in MDCK cells, the clones underwent RNA extraction, reverse transcription and high-fidelity PCR, followed by Sanger sequencing (by Macrogen) of the HA gene. To determine the hemolysis pH of the wild-type (WT) and mutant Virg09 viruses, we first prepared allantoic stocks. The method reported in previous studies^48^ was adapted to round-bottom 96-well format. Briefly, virus was added to the wells together with an equal volume of 2% chicken red blood cell (RBC) suspension in PBS. After 10 min incubation at 37 °C, unbound virus was removed by centrifugation. The cell pellets were resuspended in acidic buffer, i.e. PBS that was acidified with acetic acid to a pH ranging from 5.0 to 6.0, with 0.1 increments. After 25 min incubation at 37 °C, the samples were neutralized with NaOH and the plates were centrifugated to pellet intact RBC. The supernatants were transferred to a fresh plate and the absorbance at 540 nm was measured in a plate reader. The hemolysis pH was defined as the pH at which 50% hemolysis occurred, relative to the value at pH 5.0.

#### Pseudovirus entry assay

The pCAGEN plasmids encoding the H1 HA and N1 NA of Virg09 virus were previously described.^58^ The codon-optimized DNAs encoding H5 HA and N1 NA of the highly pathogenic avian A/H5N1 virus (A/bald eagle/FL/W22-114/2022, hereunder abbreviated FL22) were purchased from Life Technologies and cloned into pCAGEN [provided by C. Cepko (Boston, MA) via Addgene (plasmid 11160)], using NEBuilder HiFi DNA Assembly mix. To prepare murine leukemia virus (MLV) pseudoviruses bearing these HA and NA proteins, HEK293T cells were transfected with a mixture of the pCAGEN and MLV backbone plasmids (kind gift from S. Pöhlmann, German Primate Center-Leibniz Institute for Primate Research, Göttingen). The details were published before.^58^ Three days after transfection, the Virg09 pseudo-viruses were incubated for 15 min with 80 µg/mL trypsin, followed by 80 µg/mL soybean inhibitor. Finally, the Virg09 and FL22 pseudoviruses were stored in aliquots at -80 °C.

To determine the compounds’ inhibitory effect on pseudo-virus entry, MDCK cells were seeded in white 96-well plates and, on the next day, preincubated with serial compound dilutions (in medium with 2% FCS) for 20 min. The pseudovirus was added and the plates were spinoculated at 37 °C for 45 min at 450 x *g*, followed by 60 min incubation at 37 °C. After replacing the supernatants by fresh medium without compound, the plates were incubated for three days at 37 °C. To quantify the expression of the firefly luciferase reporter, we used a luciferase assay system kit and Glomax Navigator instrument, both from Promega.

#### Cell-cell fusion assay

To quantify the inhibitory effect on H1 HA-mediated membrane fusion, we used the pGal5-luc and pGal4-VP16 plasmids (kind gift from S. Pöhlmann) and transactivation setup reported by his team,^59^ with several modifications. Briefly, suspensions of HeLa cells were prepared in two tubes in which either the pGAL5-luc or pGAL5-VP16 plus HA-pCAGEN plasmid were added (or pCAGEN-empty plasmid for the mock control), together with Fugene^®^ transfection reagent and growth medium (= MEM supplemented with non-essential amino acids, HEPES, L-glutamine and 10% FCS). The cells were seeded in 6- and 96-well plates, respectively, and incubated for one day. From the 6-well plate, the cells were detached with non-enzymatic cell dissociation solution (Sigma), then overlaid on the HA-expressing cells in the 96-well plate. On the next day, the cells were sequentially incubated at 37 °C with the following reagents (with removal after each step): (i) for 15 min: 5 µg/mL of TPCK-trypsin to activate HA; (ii) for 15 min: serial compound dilutions; (iii) for exactly 5 min: fresh compound diluted in PBS-CM (= PBS supplemented with Ca^2+^ and Mg^2+^) adjusted to pH 5.3; (iv) washing with growth medium; (v) for 5 h: growth medium to allow cell-cell fusion; and (vi) cell culture lysis reagent from Promega. After reading the luminescence with the luciferase assay system kit and Glomax Navigator instrument, the relative luminescence units (RLU) values were subtracted for the value seen in mock-transfected cells, and the % luminescence at each compound concentration was calculated, relative to the condition which received medium instead of compound.

For H5 HA, we used a microscopic readout as established in a previous study.^58^ HeLa cells were seeded in 96-well plates and transfected with H5 HA (strain A/duck/Hunan/795/2002). On the next day, the cells bearing surface-exposed HA (already activated by endogenous furin) were preincubated for 15 min with compound; exposed for 5 min to PBS-CM at pH 5.2 under continued presence of compound; and then washed with PBS-CM. Cell culture medium was added and after 3 h incubation, the cells were fixated and stained with Giemsa solution to allow microscopic assessment of polykaryon formation.

#### Surface plasmon resonance (SPR)-based assessment of HA refolding

H1 HA protein (strain A/California/06/09; ecto-domain 99.4% identical to that of Virg09-HA), produced in HEK293 cells, was purchased from eEnzyme (IA-H1-11SWt). The mouse monoclonal antibodies recognizing the HA head (7B2-32) or stem region (C179) were from Kerafast and Takara Bio, respectively. The HA protein was diluted in PBS to 50 µg/mL and pre-incubated with 100 µM VF-57a or RL-007 for 10 min at 37 °C. Next, a pre-determined amount of 1 M citric acid buffer of pH 4.7 was added to install a pH of 5.2. The protein was incubated for 1 h at 37 °C to induce the conformational change in HA, then re-neutralized using citric acid buffer pH 7.

To conduct SPR analysis, the protein sample was diluted to 5 µg/mL, in HEPES-buffered saline supplemented with 1 mg/mL BSA and 0.05% v/v Tween20. First, the antibodies were coupled onto a rabbit anti-mouse C1 sensor chip (Cytiva) at a density of around 100 resonance units (RU) with a flow rate of 10 µL/min for 120 s. Next, the sample was injected for 180 s, at a flow rate of 10 µL/min and with a dissociation time of 150 s. Isotype controls were used to assess non-specific binding and buffer injections for refractive index changes. After each sample injection, a regeneration was performed using three 10 mM Glycine.HCl pH 1.7 injections at a flow rate of 30 µL/min for 20 s and a 0.5% Triton X-100 injection at a flow rate of 30 µL/min for 15 s. All binding experiments were performed on a Biacore T200 instrument (Cytiva) and at 25 °C, using a binding buffer composed of 10 mM HEPES, 150 mM NaCl and 0.05% v/v Tween20. Data were analysed using Biacore T200 Evaluation Software 3.1 and a report point at 100 s after the injection stop was used as the binding response.

### Molecular modeling

#### Ligand parametrization with QM calculations

The 3D structure of VF-57a was prepared using GaussView 6.0 and optimized at the B3LYP/6-31G(d) level employing the Gaussian 16.0 software package.^60^ The minimum energy nature of the optimized compound was verified upon inspection of the vibrational frequencies, which were positive. The ligand was parametrized using the gaff2 force field.^61^ Partial atomic charges were derived following the RESP-charges protocol^62, 63^ at the B3LYP/6-31G(d) level. Arbidol and (*S*)-F0045 were parametrized using the same protocol. In this case, the geometry used in geometry optimization was taken from the crystallographic structures available in the Protein Data Bank (PDB ID 5T6N^14^ and 6WCR,^32^ respectively).

B3LYP/6-31G(d) calculations were also performed to estimate the dipole moment in the gas phase of heterocyclic moieties examined here as potential bioisoteres of the dimethylthiophene unit of **VF-57a**. Finally, additional calculations were performed to estimate the octanol/water partition coefficient of these heterocyclic moieties using the IEFPCM-MST continuum solvation model^64^ parametrized at the B3LYP/6-31G(d) level.^65, 66^

#### Molecular modelling: homology modelling and system setup

Two homology models were built up to examine the binding of **VF-57a** to the H1 HA Virg09 using SWISSMODEL.^67^ The first relied on the complex formed by (*S*)-F0045 with PR8 HA (PDB ID 6WCR),^32^ where modelling was used to unfold the last helical turn of the short α-helix in HA^2^. This change simulated the local structure observed in the complex between H3 HA and arbidol (PDB ID 5T6N),^14^ enabling the binding of a ligand at this pocket in PR8 HA. This model was used to simulate the binding of **VF-57a** to Site A. The second was the homology model obtained using 6WCR as structural template, which was used to explore the binding of **VF-57a** and (*S*)-F0045 to the Site B. Finally, the complex of H3 HA with arbidol was performed using the homology modeling obtained for H3N2 A/Hong Kong/7/1987 strain using the X-ray structure of arbidol bound to H3 HA (A/Hong Kong/1/1968) as structural template (PDB ID 5T6N).^14^

For Site A, the X-ray structure of the arbidol-bound H3 HA and modified H1 HA model (with the last turn of the α-helix in HA_2_ unfolded) were aligned. Then, **VF-57a** was placed in the binding pocket (see Supporting Information, Figure S5) through overlay of the thiophene ring of **VF-57a** onto the thiophenyl ring of arbidol, taking advantage of the hydrophobic nature of the residues surrounding this region of the binding site. In Site B, **VF-57a** was overlaid on the chemical skeleton of (*S*)-F0045 (see Supporting Information, Figure S3), retaining the hydrogen bond between the carbonyl oxygen and the hydroxyl group of T318_2_. However, two distinct binding modes were explored, depending on the overlay of the halogenated phenyl (binding mode A) or the thiophene unit (binding mode B) onto the 2,5-dichlorophenyl moiety of (*S*)-F0045.

To calibrate the structural stability of the complexes, the crystallographic complexes of (*S*)-F0045 in PR8 HA and arbidol in HK HA were used as control systems.

Finally, due to the trimeric nature of HA, the complexes formed by HA with either arbidol, (*S*)-F0045 and **VF-57a** were built up using a stoichiometric ratio of 1:3, that is, with three ligands bound to the trimeric HA.

#### Sequence alignment

Sequences of the studied IAV strains (see Supporting Information, Figure S1) were aligned using the multiple sequence alignment tool of CLUSTAL OMEGA (v. 1.2.4) in the EMBL’s European Bioinformatics Institute (EMBL-EBI) web server.^68^

#### Molecular Dynamics Simulations

The amberff14sb force field^69^ was used for the protein, and the gaff2 force field^61^ together with RESP chargesl^62, 63^ derived at the B3LYP/6-31G(d) level were adopted for the ligand (see *Ligand parametrization with QM calculations* above). Joung and Cheatham III parameters were used for the counterions^70^ and the TIP3P model for water.^71^ Counterions (K^+^ and Cl^-^) were added to maintain the neutrality of the simulated system and to maintain the ionic concentration at 0.15 M following the SPLIT method.^72^ All simulations were performed with *AMBER20* package.^73^

Each system was minimized using 10,000 steps of steepest descent in combination with 5,000 steps of conjugate gradient algorithm. Then, each system was equilibrated in 2 steps for a total simulation time of 5 ns. The systems were heated in the NVT ensemble from 5 K to 300 K in a 250 ps temperature ramp and kept at 300K for additional 50 ps. Subsequently, the density of the system was equilibrated for 4.7 ns in the NPT ensemble (pressure: 1 bar, T: 300 K).

Production runs were done for 500 ns. Temperature control was achieved using Langevin dynamics and pressure control was maintained using the Berendsen barostat. All bonds involving hydrogen atoms were constrained by the SHAKE algorithm^74^ in order to employ a timestep of 2 fs. During all the simulations a set of distance NMR restraints were applied to maintain the structural stability of the HA trimer, since the models (constructed from the mentioned X-ray structures above) do not include the transmembrane part of the protein. The restraints involved one residue of each HA monomer located at the end-chains. A force constant of 5 kcal/mol was applied gradually when a displacement higher than 2Å from the crystallographic distance occurs (the restraint becomes fixed at 5kcal/mol 1.5 Å after the first cutoff).

The analysis of the trajectories collected from MD simulations was performed using the tools included in the cpptraj^75^ software available in the Amber package.

#### Relative Binding Free Energy (RBFE) Simulations

The RBFE^76^ between derivatives was determined through alchemical transformations, wherein a ligand (L1) is converted into a structurally related analogue (L2) both in the protein-bound complex and in the unbound state in aqueous solution (see Eq. 1).

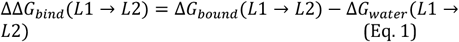

The transformation from L1 to L2 was divided into a series of windows, where λ = 0 and λ = 1 represent L1 and L2, respectively, and intermediate λ values denote a linear interpolation between the parameters of the initial and final systems.^77^ A dual-topology alchemical perturbation protocol was employed, utilizing 23 λ-windows evenly spaced following the trapezoidal rule. This choice, which was motivated by our own experience with similar alchemical transformations,^78^ enabled us to reduce the simulation time for each intermediate, since a smoother integration pathway allows for faster convergence and reduces fluctuations, although this involved a larger consumption of computational resources. To smoothly transition from L1 to L2, softcore potentials were applied to the atoms involved in a chemical change. Each λ simulation typically ran for 6 ns, adding extensions in the windows that were not converged when necessary. When simulations at a given window exceeded 6 ns, the last 6 ns of the trajectory ran were used to calculate the contribution of this window final free energy difference. The Thermodynamic Integration (TI) estimator, implemented in alchemlyb^79-81^ was utilized to estimate the free energy change for each λ simulation. All simulations were conducted using the GPU-accelerated TI implementation^76, 82^ of Amber20 (see Supporting Information Figures S8-S10 for representation of the free energy changes).

Initial coordinates for each binding mode were extracted from unbiased MD runs. All systems were meticulously equilibrated with the dual topology using the TI code in each λ before production runs. Each system underwent heating in the NVT ensemble from 5 K to 300 K (150 ps), followed by equilibration at 1 bar in the NPT ensemble (300 ps). Finally, the density equilibrated system was simulated for 100 ps more switching back to the NVT ensemble preparing the system to conduct the actual RBFE production. During the equilibration process, the heavy atoms of the protein and ligand were restrained with positional restraints (5 kcal·mol^−1^) additionally to the NMR restraints applied to the bottom of the HA model as in the unbiased MD simulations. Subsequently, each λ state was simulated in the NVT ensemble at 300 K using the Langevin thermostat and the SHAKE algorithm for bonds involving hydrogen atoms.

## ASSOCIATED CONTENT

### Supporting Information

Synthesis of some starting materials, ^1^H and ^13^C NMR spectra and elemental analysis data of the new compounds, Table S1 (elemental analysis data), Figures S1-S7 (PDF). Molecular formula string and data (CSV). The final frame obtained from the MD simulation of **VF-57a** bound to HA (PDB format). Authors will release the atomic coordinates and experimental data upon article publication. This material is available free of charge via the Internet at http://pubs.acs.org.

## AUTHOR INFORMATION

### Author Contributions

^‡^These authors (S.R. and A.V.) contributed equally.

^†^Deceased. Óscar Lozano passed away in 2021 while this work was ongoing.

L.N. and S.V. conceived the project. V.F., J.M., O.L., and C.E.M. synthesized and chemically characterized the compounds. A.V. and F.J.L. performed MD calculations. S.R., R.V.B., C.M., L.S., K.V., S.N., A.S. and L.N. performed the antiviral assays and the cytotoxicity studies. L.N., F.J.L., and S.V. analyzed the data. L.N., F.J.L., and S.V. wrote, edited and reviewed the manuscript with feedback from all the authors. All authors have given approval to the final version of the manuscript.

### Notes

None of the authors has any disclosures to declare.

## ACKNOWLEDGMENT

The authors would like to thank Manon Laporte for her contributions in setting up the luciferase-based cell-cell fusion assay.

This work was funded by the Spanish *Ministerio de Ciencia, Innovación y Universidades*, MICIU/AEI/10.13039/501100011033: Grants PID2023-147004OB-I00 (to S.V.), PID2020-117646RB-I00, PID2023-147942OB-I00 and Maria de Maetzu CEX2021-001202-M) (to F.J.L.). This study was supported by funding from *Fundació La Marató de TV3* (Nos. 201832 and 202135 to L.N. and S.V.), and Agència de Gestió d’Ajuts Universitaris i de Recerca (grant 2021SGR00671). S.R. is holder of an SB-PhD fellowship from the FWO Research Foundation Flanders (No. 1S92321N). C.E.M. was supported by the Research Personnel Training Program (FPI grant PRE2022-104952) of the Spanish *Ministerio de Ciencia, Innovación y Universidades*.

## ABBREVIATIONS

ATR: Attenuated Total Reflectance
BXA: baloxavir acid
CPE: Cytopathic Effect, CPE
HA: Hemagglutinin
MCC: Minimum Cytotoxic Concentration
MD: Molecular Dynamics
MDCK: Madin-Darby canine kidney cells
ND: Not Determined
RBFE: Relative Binding Free Energy
SAR: structure-activity relationships
SI: Selectivity Index
TBS: Topliss Batchwise Scheme

## Table of Contents Graphic

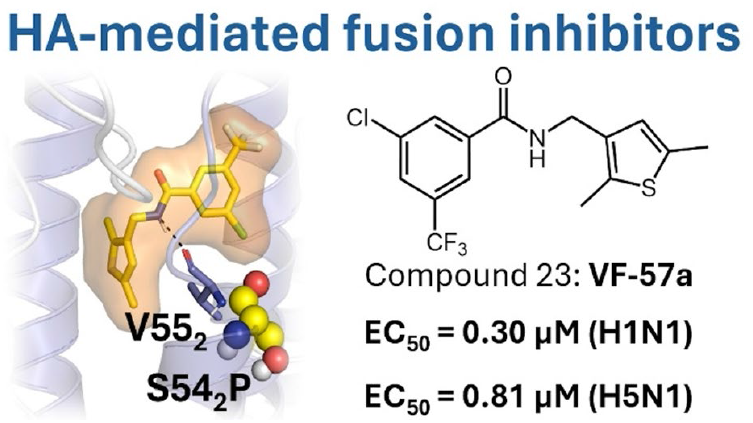

## REFERENCES

(1) Iuliano, A. D.; Roguski, K. M.; Chang, H. H.; Muscatello, D. J.; Palekar, R.; Tempia, S.; Cohen, C.; Gran, J. M.; Schanzer, D.; Cowling, B. J.; Wu, P.; Kyncl, J.; Ang, L. W.; Park, M.; Redlberger-Fritz, M.; Yu, H.; Espenhain, L.; Krishnan, A.; Emukule, G.; van Asten, L.; Pereira da Silva, S.; Aungkulanon, S.; Buchholz, U.; Widdowson, M. A.; Bresee, J. S., Estimates of global seasonal influenza-associated respiratory mortality: a modelling study. Lancet 2018, 391 (10127), 1285–1300. DOI: 10.1016/s0140-6736(17)33293-2.

(2) Krammer, F.; Smith, G. J. D.; Fouchier, R. A. M.; Peiris, M.; Kedzierska, K.; Doherty, P. C.; Palese, P.; Shaw, M. L.; Treanor, J.; Webster, R. G.; Garcia-Sastre, A., Influenza. Nat Rev Dis Primers 2018, 4 (1), 3. DOI: 10.1038/s41572-018-0002-y.

(3) Dawood, F. S.; Iuliano, A. D.; Reed, C.; Meltzer, M. I.; Shay, D. K.; Cheng, P.-Y.; Bandaranayake, D.; Breiman, R. F.; Brooks, W. A.; Buchy, P.; Feikin, D. R.; Fowler, K. B.; Gordon, A.; Hien, N. T.; Horby, P.; Huang, Q. S.; Katz, M. A.; Krishnan, A.; Lal, R.; Montgomery, J. M.; Mølbak, K.; Pebody, R.; Presanis, A. M.; Razuri, H.; Steens, A.; Tinoco, Y. O.; Wallinga, J.; Yu, H.; Vong, S.; Bresee, J.; Widdowson, M.-A., Estimated global mortality associated with the first 12 months of 2009 pandemic influenza A H1N1 virus circulation: a modelling study. Lancet Infect. Dis. 2012, 12 (9), 687–695. DOI: 10.1016/s1473-3099(12)70121-4.

(4) Nguyen, T. Q.; Hutter, C. R.; Markin, A.; Thomas, M.; Lantz, K.; Killian, M. L.; Janzen, G. M.; Vijendran, S.; Wagle, S.; Inderski, B.; Magstadt, D. R.; Li, G.; Diel, D. G.; Frye, E. A.; Dimitrov, K. M.; Swinford, A. K.; Thompson, A.; Snekvik, K. R.; Suarez, D. L.; Lakin, S. M.; Schwabenlander, S.; Ahola, S. C.; Johnson, K. R.; Baker, A. L.; Robbe-Austerman, S.; Torchetti, M. K.; Anderson, T. K., Emergence and interstate spread of highly pathogenic avian influenza A(H5N1) in dairy cattle in the United States. Science 2025, 388 (6745), eadq0900. DOI: 10.1126/science.adq0900.

(5) Peacock, T. P.; Moncla, L.; Dudas, G.; VanInsberghe, D.; Sukhova, K.; Lloyd-Smith, J. O.; Worobey, M.; Lowen, A. C.; Nelson, M. I., The global H5N1 influenza panzootic in mammals. Nature 2025, 637 (8045), 304–313. DOI: 10.1038/s41586-024-08054-z.

(6) Lin, T. H.; Zhu, X.; Wang, S.; Zhang, D.; McBride, R.; Yu, W.; Babarinde, S.; Paulson, J. C.; Wilson, I. A., A single mutation in bovine influenza H5N1 hemagglutinin switches specificity to human receptors. Science 2024, 386 (6726), 1128–1134. DOI: 10.1126/science.adt0180.

(7) CDC Seasonal Flu Vaccine Effectiveness Studies. https://www.cdc.gov/flu-vaccines-work/php/effectiveness-studies/?CDC_AAref_Val= https://www.cdc.gov/flu/vaccines-work/effectiveness-studies.htm (accessed 07/05/2025).

(8) Nguyen-Van-Tam, J. S.; Venkatesan, S.; Muthuri, S. G.; Myles, P. R., Neuraminidase inhibitors: who, when, where? Clin Microbiol Infect 2015, 21 (3), 222–5. DOI: 10.1016/j.cmi.2014.11.020.

(9) Uyeki, T. M.; Hui, D. S.; Zambon, M.; Wentworth, D. E.; Monto, A. S., Influenza. Lancet 2022, 400 (10353), 693–706. DOI: 10.1016/S0140-6736(22)00982-5.

(10) Beigel, J. H.; Hayden, F. G., Influenza therapeutics in clinical practice-challenges and recent advances. Cold Spring Harb Perspect Med 2021, 11 (4), a038463. DOI: 10.1101/cshperspect.a038463.

(11) Stevaert, A.; Naesens, L., The influenza virus polymerase complex: an update on its structure, functions, and significance for antiviral drug design. Med Res Rev 2016, 36 (6), 1127–1173. DOI: 10.1002/med.21401.

(12) Govorkova, E. A.; Takashita, E.; Daniels, R. S.; Fujisaki, S.; Presser, L. D.; Patel, M. C.; Huang, W.; Lackenby, A.; Nguyen, H. T.; Pereyaslov, D.; Rattigan, A.; Brown, S. K.; Samaan, M.; Subbarao, K.; Wong, S.; Wang, D.; Webby, R. J.; Yen, H. L.; Zhang, W.; Meijer, A.; Gubareva, L. V., Global update on the susceptibilities of human influenza viruses to neuraminidase inhibitors and the cap-dependent endonuclease inhibitor baloxavir, 2018-2020. Antiviral Res 2022, 200, 105281. DOI: 10.1016/j.antiviral.2022.105281.

(13) Gamblin, S. J.; Vachieri, S. G.; Xiong, X.; Zhang, J.; Martin, S. R.; Skehel, J. J., Hemagglutinin Structure and Activities. Cold Spring Harb Perspect Med 2020, 11 (10), :a038638. DOI: 10.1101/cshperspect.a038638.

(14) Kadam, R. U.; Wilson, I. A., Structural basis of influenza virus fusion inhibition by the antiviral drug Arbidol. PNAS 2017, 114 (2), 206–214. DOI: 10.1073/pnas.1617020114.

(15) Blaising, J.; Polyak, S. J.; Pecheur, E. I., Arbidol as a broad-spectrum antiviral: an update. Antiviral Res 2014, 107 (2014), 84–94. DOI: 10.1016/j.antiviral.2014.04.006.

(16) Huang, Q. J.; Kim, R.; Song, K.; Grigorieff, N.; Munro, J. B.; Schiffer, C. A.; Somasundaran, M., Virion-associated influenza hemagglutinin clusters upon sialic acid binding visualized by cryoelectron tomography. PNAS 2025, 122 (16), e2426427122. DOI: 10.1073/pnas.2426427122.

(17) Di Lella, S.; Herrmann, A.; Mair, C. M., Modulation of the pH stability of influenza virus hemagglutinin: a host cell adaptation strategy. Biophys J 2016, 110 (11), 2293–2301. DOI: 10.1016/j.bpj.2016.04.035.

(18) Benton, D. J.; Gamblin, S. J.; Rosenthal, P. B.; Skehel, J. J., Structural transitions in influenza haemagglutinin at membrane fusion pH. Nature 2020, 583 (7814), 150–153. DOI: 10.1038/s41586-020-2333-6.

(19) Garcia-Moro, E.; Zhang, J.; Calder, L. J.; Brown, N. R.; Gamblin, S. J.; Skehel, J. J.; Rosenthal, P. B., Reversible structural changes in the influenza hemagglutinin precursor at membrane fusion pH. PNAS 2022, 119 (33), e2208011119. DOI: 10.1073/pnas.2208011119.

(20) Vanderlinden, E.; Naesens, L., Emerging antiviral strategies to interfere with influenza virus entry. Med Res Rev 2014, 34 (2), 301–39. DOI: 10.1002/med.21289.

(21) Chen, Z.; Cui, Q.; Caffrey, M.; Rong, L.; Du, R., Small molecule inhibitors of influenza virus entry. Pharmaceuticals 2021, 14 (6), 587. DOI: 10.3390/ph14060587.

(22) Liu, H. Y.; Yang, P. L., Small-molecule inhibition of viral fusion glycoproteins. Annu Rev Virol 2021, 8 (1), 459–489. DOI: 10.1146/annurev-virology-022221-063725.

(23) Hermoso-Pinilla, F. J.; Valdivia, A.; Camarasa, M. J.; Ginex, T.; Luque, F. J., Influenza A virus hemagglutinin: from classical fusion inhibitors to proteolysis targeting chimera-based strategies in antiviral drug discovery. Explor Drug Sci. 2024, 2, 85–116. DOI: 10.37349/eds.2024.00037.

(24) Sun, X.; Ma, H.; Wang, X.; Bao, Z.; Tang, S.; Yi, C.; Sun, B., Broadly neutralizing antibodies to combat influenza virus infection. Antiviral Res 2024, 221, 105785. DOI: 10.1016/j.antiviral.2023.105785.

(25) van Dongen, M. J. P.; Kadam, R. U.; Juraszek, J.; Lawson, E.; Brandenburg, B.; Schmitz, F.; Schepens, W. B. G.; Stoops, B.; van Diepen, H. A.; Jongeneelen, M.; Tang, C.; Vermond, J.; van Eijgen-Obregoso Real, A.; Blokland, S.; Garg, D.; Yu, W.; Goutier, W.; Lanckacker, E.; Klap, J. M.; Peeters, D. C. G.; Wu, J.; Buyck, C.; Jonckers, T. H. M.; Roymans, D.; Roevens, P.; Vogels, R.; Koudstaal, W.; Friesen, R. H. E.; Raboisson, P.; Dhanak, D.; Goudsmit, J.; Wilson, I. A., A small-molecule fusion inhibitor of influenza virus is orally active in mice. Science 2019, 363 (6431), eaar6221. DOI: 10.1126/science.aar6221.

(26) Gaisina, I.; Li, P.; Du, R.; Cui, Q.; Dong, M.; Zhang, C.; Manicassamy, B.; Caffrey, M.; Moore, T.; Cooper, L.; Rong, L., An orally active entry inhibitor of influenza A viruses protects mice and synergizes with oseltamivir and baloxavir marboxil. Sci Adv 2024, 10 (8), eadk9004. DOI: 10.1126/sciadv.adk9004.

(27) Zhu, L.; Li, Y.; Li, S.; Li, H.; Qiu, Z.; Lee, C.; Lu, H.; Lin, X.; Zhao, R.; Chen, L.; Wu, J. Z.; Tang, G.; Yang, W., Inhibition of influenza A virus (H1N1) fusion by benzenesulfonamide derivatives targeting viral hemagglutinin. PLoS One 2011, 6 (12), e29120. DOI: 10.1371/journal.pone.0029120.

(28) Zhao, X.; Li, R.; Zhou, Y.; Xiao, M.; Ma, C.; Yang, Z.; Zeng, S.; Du, Q.; Yang, C.; Jiang, H.; Hu, Y.; Wang, K.; Mok, C. K. P.; Sun, P.; Dong, J.; Cui, W.; Wang, J.; Tu, Y.; Yang, Z.; Hu, W., Discovery of highly potent pinanamine-based inhibitors against amantadine- and oseltamivir-resistant influenza A viruses. J Med Chem 2018, 61 (12), 5187–5198. DOI: 10.1021/acs.jmedchem.8b00042.

(29) Ye, M.; Liao, Y.; Wu, L.; Qi, W.; Choudhry, N.; Liu, Y.; Chen, W.; Song, G.; Chen, J., An oleanolic acid derivative inhibits hemagglutinin-mediated entry of influenza A virus. Viruses 2020, 12 (2), 225. DOI: 10.3390/v12020225.

(30) White, K.; Esparza, M.; Liang, J.; Bhat, P.; Naidoo, J. McGovern, B.L.; Williams, M. A. P.; Alabi, B. R.; Shay J.; Niederstrasser, H.; Posner, B.; García-Sastre, A.; Ready, J.; Fontoura, B. M. A., Aryl sulfonamide inhibits entry and replication of diverse influenza viruses via the hemagglutinin protein. J Med Chem 2021, 64 (15), 10951–10966. DOI: 10.1021/acs.jmedchem.1c00304.

(31) Russell, R. J.; Kerry, P. S.; Stevens, D. J.; Steinhauer, D. A.; Martin, S. R.; Gamblin, S. J.; Skehel, J. J., Structure of influenza hemagglutinin in complex with an inhibitor of membrane fusion. PNAS 2008, 105 (46), 17736–41. DOI: 10.1073/pnas.0807142105.

(32) Yao, Y.; Kadam, R. U.; Lee, C. D.; Woehl, J. L.; Wu, N. C.; Zhu, X.; Kitamura, S.; Wilson, I. A.; Wolan, D. W., An influenza A hemagglutinin small-molecule fusion inhibitor identified by a new high-throughput fluorescence polarization screen. PNAS 2020, 117 (31), 18431–18438. DOI: 10.1073/pnas.2006893117.

(33) Antanasijevic, A.; Durst, M. A.; Cheng, H.; Gaisina, I. N.; Perez, J. T.; Manicassamy, B.; Rong, L.; Lavie, A.; Caffrey, M., Structure of avian influenza hemagglutinin in complex with a small molecule entry inhibitor. Life Sci Alliance 2020, 3 (8), :e202000724. DOI: 10.26508/lsa.202000724.

(34) Plotch, S. J.; O’Hara, B.; Morin, J.; Palant, O.; LaRocque, J.; Bloom, J. D.; Lang, S. A., Jr.; DiGrandi, M. J.; Bradley, M.; Nilakantan, R.; Gluzman, Y., Inhibition of influenza A virus replication by compounds interfering with the fusogenic function of the viral hemagglutinin. J Virol 1999, 73 (1), 140–51. DOI: 10.1128/JVI.73.1.140-151.1999.

(35) Liu, S.; Li, R.; Zhang, R.; Chan, C. C.; Xi, B.; Zhu, Z.; Yang, J.; Poon, V. K.; Zhou, J.; Chen, M.; Munch, J.; Kirchhoff, F.; Pleschka, S.; Haarmann, T.; Dietrich, U.; Pan, C.; Du, L.; Jiang, S.; Zheng, B., CL-385319 inhibits H5N1 avian influenza A virus infection by blocking viral entry. Eur J Pharmacol 2011, 660 (2-3), 460–7. DOI: 10.1016/j.ejphar.2011.04.013.

(36) Zhu, Z.; Li, R.; Xiao, G.; Chen, Z.; Yang, J.; Zhu, Q.; Liu, S., Design, synthesis and structure-activity relationship of novel inhibitors against H5N1 hemagglutinin-mediated membrane fusion. Eur J Med Chem 2012, 57, 211–6. DOI: 10.1016/j.ejmech.2012.08.041.

(37) Zhu, Z.; Yao, Z.; Shen, X.; Chen, Z.; Liu, X.; Parquette, J. R.; Liu, S., Oligothiophene compounds inhibit the membrane fusion between H5N1 avian influenza virus and the endosome of host cell. Eur J Med Chem 2017, 130, 185–194. DOI: 10.1016/j.ejmech.2017.02.040.

(38) Yu, Y.; Tazeem Xu, Z.; Du, L.; Jin, M.; Dong, C.; Zhou, H. B.; Wu, S., Design and synthesis of heteroaromatic-based benzenesulfonamide derivatives as potent inhibitors of H5N1 influenza A virus. Medchemcomm 2019, 10 (1), 89–100. DOI: 10.1039/c8md00474a.

(39) Vanderlinden, E.; Göktas, F.; Cesur, Z.; Froeyen, M.; Reed, M. L.; Russell, C. J.; Cesur, N.; Naesens, L., Novel inhibitors of influenza virus fusion: structure-activity relationship and interaction with the viral hemagglutinin. J Virol 2010, 84 (9), 4277–88. DOI: 10.1128/JVI.02325-09.

(40) Cihan-Ustundag, G.; Zopun, M.; Vanderlinden, E.; Ozkirimli, E.; Persoons, L.; Capan, G.; Naesens, L., Superior inhibition of influenza virus hemagglutinin-mediated fusion by indole-substituted spirothiazolidinones. Bioorg Med Chem 2020, 28 (1), 115130. DOI: 10.1016/j.bmc.2019.115130.

(41) Du, R.; Cheng, H.; Cui, Q.; Peet, N. P.; Gaisina, I. N.; Rong, L., Identification of a novel inhibitor targeting influenza A virus group 2 hemagglutinins. Antiviral Res 2021, 186, 105013. DOI: 10.1016/j.antiviral.2021.105013.

(42) Alqarni, S.; Cooper, L.; Galvan Achi, J.; Bott, R.; Sali, V. K.; Brown, A.; Santarsiero, B. D.; Krunic, A.; Manicassamy, B.; Peet, N. P.; Zhang, P.; Thatcher, G. R. J.; Gaisina, I. N.; Rong, L.; Moore, T. W., Synthesis, optimization, and structure-activity relationships of imidazo[1,2-a]pyrimidines as inhibitors of group 2 influenza A viruses. J Med Chem 2022, 65 (20), 14104–14120. DOI: 10.1021/acs.jmedchem.2c01329.

(43) White, K. M.; De Jesus, P.; Chen, Z.; Abreu, P., Jr.; Barile, E.; Mak, P. A.; Anderson, P.; Nguyen, Q. T.; Inoue, A.; Stertz, S.; Koenig, R.; Pellecchia, M.; Palese, P.; Kuhen, K.; Garcia-Sastre, A.; Chanda, S. K.; Shaw, M. L., A potent anti-influenza compound blocks fusion through stabilization of the prefusion conformation of the hemagglutinin protein. ACS Infect Dis 2015, 1 (2), 98–109. DOI: 10.1021/id500022h.

(44) Basu, A.; Antanasijevic, A.; Wang, M.; Li, B.; Mills, D. M.; Ames, J. A.; Nash, P. J.; Williams, J. D.; Peet, N. P.; Moir, D. T.; Prichard, M. N.; Keith, K. A.; Barnard, D. L.; Caffrey, M.; Rong, L.; Bowlin, T. L., New small molecule entry inhibitors targeting hemagglutinin-mediated influenza a virus fusion. J Virol 2014, 88 (3), 1447–60. DOI: 10.1128/JVI.01225-13.

(45) de Castro, S.; Ginex, T.; Vanderlinden, E.; Laporte, M.; Stevaert, A.; Cumella, J.; Gago, F.; Camarasa, M. J.; Luque, F. J.; Naesens, L.; Velazquez, S., N-benzyl 4,4-disubstituted piperidines as a potent class of influenza H1N1 virus inhibitors showing a novel mechanism of hemagglutinin fusion peptide interaction. Eur J Med Chem 2020, 194, 112223. DOI: 10.1016/j.ejmech.2020.112223.

(46) Kitamura, S.; Lin, T. H.; Lee, C. D.; Takamura, A.; Kadam, R. U.; Zhang, D.; Zhu, X.; Dada, L.; Nagai, E.; Yu, W.; Yao, Y.; Sharpless, K. B.; Wilson, I. A.; Wolan, D. W., Ultrapotent influenza hemagglutinin fusion inhibitors developed through SuFEx-enabled high-throughput medicinal chemistry. PNAS 2024, 121 (22), e2310677121. DOI: 10.1073/pnas.2310677121.

(47) Kim, J. I.; Lee, S.; Lee, G. Y.; Park, S.; Bae, J. Y.; Heo, J.; Kim, H. Y.; Woo, S. H.; Lee, H. U.; Ahn, C. A.; Bang, H. J.; Ju, H. S.; Ok, K.; Byun, Y.; Cho, D. J.; Shin, J. S.; Kim, D. Y.; Park, M. S.; Park, M. S., Novel small molecule targeting the hemagglutinin stalk of influenza viruses. J Virol 2019, 93 (17), e00878–19. DOI: 10.1128/JVI.00878-19.

(48) Leiva, R.; Barniol-Xicota, M.; Codony, S.; Ginex, T.; Vanderlinden, E.; Montes, M.; Caffrey, M.; Luque, F. J.; Naesens, L.; Vazquez, S., Aniline-based inhibitors of influenza H1N1 virus acting on hemagglutinin-mediated fusion. J Med Chem 2018, 61 (1), 98–118. DOI: 10.1021/acs.jmedchem.7b00908.

(49) Topliss, J. G., A manual method for applying the Hansch approach to drug design. J Med Chem 1977, 20 (4), 463–9. DOI: 10.1021/jm00214a001.

(50) Richter, L., Topliss batchwise schemes reviewed in the era of open data reveal significant differences between enzymes and membrane receptors. J Chem Inf Model 2017, 57 (10), 2575–2583. DOI: 10.1021/acs.jcim.7b00195.

(51) Kandeil, A.; Patton, C.; Jones, J. C.; Jeevan, T.; Harrington, W. N.; Trifkovic, S.; Seiler, J. P.; Fabrizio, T.; Woodard, K.; Turner, J. C.; Crumpton, J. C.; Miller, L.; Rubrum, A.; DeBeauchamp, J.; Russell, C. J.; Govorkova, E. A.; Vogel, P.; Kim-Torchetti, M.; Berhane, Y.; Stallknecht, D.; Poulson, R.; Kercher, L.; Webby, R. J., Rapid evolution of A(H5N1) influenza viruses after intercontinental spread to North America. Nat Commun 2023, 14 (1), 3082. DOI: 10.1038/s41467-023-38415-7.

(52) Yang, H.; Carney, P.; Stevens, J., Structure and Receptor binding properties of a pandemic H1N1 virus hemagglutinin. PLoS Curr 2010, 2, RRN1152. DOI: 10.1371/currents.RRN1152.

(53) Yassine, H. M.; Boyington, J. C.; McTamney, P. M.; Wei, C..; Kanekiyo, M.; Kong, W. P.; Gallagher, J. R.; Wang, L.; Zhang, Y.; Joyce, M. G.; Lingwood, D.; Moin, S. M.; Andersen, H.; Okuno, Y.; Rao, S. S.; Harris, A. K.; Kwong, P..; Mascola, J. R.; Nabel, G. J.; Graham, B. S., Hemagglutinin-stem nanoparticles generate heterosubtypic influenza protection. Nat Med 2015, 21 (9), 1065–70. DOI: 10.1038/nm.3927.

(54) Zhao, X.; Jie, Y.; Rosenberg, M. R.; Wan, J.; Zeng, S.; Cui, W.; Xiao, Y.; Li, Z.; Tu, Z.; Casarotto, M. G.; Hu, W., Design and synthesis of pinanamine derivatives as anti-influenza A M2 ion channel inhibitors. Antiviral Res 2012, 96 (2), 91–9. DOI: 10.1016/j.antiviral.2012.09.001.

(55) Basu, A.; Komazin-Meredith, G.; McCarthy, C.; Antanasijevic, A.; Cardinale, S. C.; Mishra, R. K.; Barnard, D..; Caffrey, M.; Rong, L.; Bowlin, T. L., Molecular mechanism underlying the action of the influenza A virus fusion inhibitor MBX2546. ACS Infect Dis 2017, 3 (5), 330–335. DOI: 10.1021/acsinfecdis.6b00194.

(56) Vrijens, P.; Noppen, S.; Boogaerts, T.; Vanstreels, E.; Ronca, R.; Chiodelli, P.; Laporte, M.; Vanderlinden, E.; Liekens, S.; Stevaert, A.; Naesens, L., Influenza virus entry via the GM3 ganglioside-mediated platelet-derived growth factor receptor beta signalling pathway. J Gen Virol 2019, 100 (4), 583–601. DOI: 10.1099/jgv.0.001235.

(57) Lawrenz, J.; Wettstein, L.; Rodriguez Alfonso, A.; Nchioua, R.; von Maltitz, P.; Albers, D. P. J.; Zech, F.; Vandeput, J.; Naesens, L.; Fois, G.; Neubauer, V.; Preising, N.; Schmierer, E.; Almeida-Hernandez, Y.; Petersen, M.; Standker, L.; Wiese, S.; Braubach, P.; Frick, M.; Barth, E.; Sauter, D.; Kirchhoff, F.; Sanchez-Garcia, E.; Stevaert, A.; Munch, J., Trypstatin as a Novel TMPRSS2 Inhibitor with Broad-Spectrum Efficacy against Corona and Influenza Viruses. Adv Sci (Weinh) 2025, 12 (25), e2506430. DOI: 10.1002/advs.202506430.

(58) Laporte, M.; Stevaert, A.; Raeymaekers, V.; Boogaerts, T.; Nehlmeier, I.; Chiu, W.; Benkheil, M.; Vanaudenaerde, B.; Pohlmann, S.; Naesens, L., Hemagglutinin cleavability, acid stability, and temperature dependence optimize influenza B virus for replication in human airways. J Virol 2019, 94 (1), e01430–19. DOI: 10.1128/JVI.01430-19.

(59) Glowacka, I.; Bertram, S.; Muller, M. A.; Allen, P.; Soilleux, E.; Pfefferle, S.; Steffen, I.; Tsegaye, T. S.; He, Y.; Gnirss, K.; Niemeyer, D.; Schneider, H.; Drosten, C.; Pohlmann, S., Evidence that TMPRSS2 activates the severe acute respiratory syndrome coronavirus spike protein for membrane fusion and reduces viral control by the humoral immune response. J Virol 2011, 85 (9), 4122–4134. DOI: 10.1128/jvi.02232-10.

(60) Frisch, M. J.; Trucks, G. W.; Schlegel, H. B.; Scuseria, G. E.; Robb, M. A.; Cheeseman, J. R.; Scalmani, G.; Barone, V.; Petersson, G. A.; Nakatsuji, H.; Li, X.; Caricato, M.; Marenich, A. V.; Bloino, J.; Janesko, B. G.; Gomperts, R.; Mennucci, B.; Hratchian, H. P.; Ortiz, J. V.; Izmaylov, A. F.; Sonnenberg, J. L.; Williams-Young, D.; Ding, F.; Lipparini, F.; Egidi, F.; Goings, J.; Peng, B.; Petrone, A.; Henderson, T.; Ranasinghe, D.; Zakrzewski, V. G.; Gao, J.; Rega, N.; Zheng, G.; Liang, W.; Hada, M.; Ehara, M.; Toyota, K.; Fukuda, R.; Hasegawa, J.; Ishida, M.; Nakajima, T.; Honda, Y.; Kitao, O.; Nakai, H.; Vreven, T.; Throssell, K.; Montgomery, J. A., Jr.; Peralta, J.E.; Ogliaro, F.; Bearpark, M.J.; Heyd, J. J.; Brothers, E. N.; Kudin, K. N.; Staroverov, V. N.; Keith, T. A.; Kobayashi, R.; Normand, J.; Raghavachari, K.; Rendell, A. P.; Burant, J. C.; Iyengar, S. S.; Tomasi, J.; Cossi, M.; Millam, J. M.; Klene, M.; Adamo, C.; Cammi, R.; Ochterski, J. W.; Martin, R. L.; Morokuma, K.; Farkas, O.; Foresman, J. B.; Fox, D. J. Gaussian 16, Revision B.01, Gaussian, Inc., Wallingford CT.: 2016.

(61) He, X.; Man, V. H.; Yang, W.; Lee, T. S.; Wang, J., A fast and high-quality charge model for the next generation general AMBER force field. J Chem Phys 2020, 153 (11), 114502. DOI: 10.1063/5.0019056.

(62) Cornell, W. D.; Cieplak, P.; Bayly, C. I.; Gould, I. R.; Merz, K. M.; Ferguson, D. M.; Spellmeyer, D. C.; Fox, T.; Caldwell, J. W.; Kollman, P. A., A Second Generation Force Field for the Simulation of Proteins, Nucleic Acids, and Organic Molecules. JACS 1995, 117 (19), 5179–5197. DOI: 10.1021/ja00124a002.

(63) Bayly, C. I.; Cieplak, P.; Cornell, W.; Kollman, P. A., A well-behaved electrostatic potential based method using charge restraints for deriving atomic charges: the RESP model. J Phys Chem 1993, 97 (40), 10269–10280. DOI: 10.1021/j100142a004.

(64) Javier Luque, F.; Curutchet, C.; Muñoz-Muriedas, J.; Bidon-Chanal, A.; Soteras, I.; Morreale, A.; Gelpí, J. L.; Orozco, M., Continuum solvation models: Dissecting the free energy of solvation. Phys Chem Chem Phys 2003, 5 (18), 3827–3836. DOI: 10.1039/B306954K.

(65) Curutchet, C.; Orozco, M.; Luque, F. J., Solvation in octanol: parametrization of the continuum MST model. J Comp Chem 2001, 22 (11), 1180–1193. DOI: 10.1002/jcc.1076.

(66) Soteras, I.; Curutchet, C.; Bidon-Chanal, A.; Orozco, M.; Luque, F. J., Extension of the MST model to the IEF formalism: HF and B3LYP parametrizations. J Mol Struct: THEOCHEM 2005, 727 (1), 29–40. DOI: 10.1016/j.theochem.2005.02.029.

(67) Waterhouse, A.; Bertoni, M.; Bienert, S.; Studer, G.; Tauriello, G.; Gumienny, R.; Heer, F. T.; de Beer, T. A. P.; Rempfer, C.; Bordoli, L.; Lepore, R.; Schwede, T., SWISS-MODEL: homology modelling of protein structures and complexes. Nucleic Acids Res 2018, 46 (W1), W296–W303. DOI: 10.1093/nar/gky427.

(68) Madeira, F.; Pearce, M.; Tivey, A. R. N.; Basutkar, P.; Lee, J.; Edbali, O.; Madhusoodanan, N.; Kolesnikov, A.; Lopez, R., Search and sequence analysis tools services from EMBL-EBI in 2022. Nucleic Acids Res 2022, 50 (W1), W276-W279. DOI: 10.1093/nar/gkac240.

(69) Teo, R. D.; Tieleman, D. P., Evaluation of all-atom force fields in viral capsid simulations and properties. RSC Adv 2022, 12 (1), 216–220. DOI: 10.1039/D1RA08431C.

(70) Joung, I. S.; Cheatham, T. E., III, Determination of alkali and halide monovalent ion parameters for use in explicitly solvated biomolecular simulations. J Phys Chem B 2008, 112 (30), 9020–9041. DOI: 10.1021/jp8001614.

(71) Mark, P.; Nilsson, L., Structure and dynamics of the TIP3P, SPC, and SPC/E water models at 298 K. J. Phys. Chem. A 2001, 105 (43), 9954–9960. DOI: 10.1021/jp003020w.

(72) Machado, M. R.; Pantano, S., Split the charge difference in two! A rule of thumb for adding proper amounts of ions in MD simulations. J Chem Theory Comput 2020, 16 (3), 1367–1372. DOI: 10.1021/acs.jctc.9b00953.

(73) Case, D. A. K. B., K.; Ben-Shalom, I. Y.; Brozell, S. R.; Cerutti, D. S.; Cheatham, T. E., III; Cruzeiro, V.W.D.; Darden, T. A.; Duke, R. E.; Giambasu, G.; Gilson, M. K.; Gohlke, H.; Goetz, A. W.; Harris, R.; Izadi, S.; Izmailov, S. A.; Kasavajhala, K.; Kovalen-ko, A.; Krasny, R.; Kurtzman, T.; Lee, T. S.; LeGrand, S.; Li, P.; Lin, C.; Liu, J.; Luchko, T.; Luo, R.; Man, V.; Merz, K. M.; Miao, Y.; Mikhailovskii, O.; Monard, G.; Nguyen, H.; O’Hearn, K. A.; Onufriev, A.; Pan, F.; Pantano, S.; Qi, R.; Rahnamoun, A.; Roe, D. R.; Roitberg, A.; Sagui, C.; Schott-Verdugo, S.; Shajan, A.; Shen, J.; Simmerling, C. L.; Skrynnikov, N. R.; Smith, J.; Swails, J.; Walk-er, R. C.; Wang, J.; Wang, J.; Wei, H.; Wu, X.; Wu, Y.; Xiong, Y.; Xue, Y.; York, D. M.; Zhao, S.; Zhu, Q.; Kollman, P. A., AMBER 2020. University of California: San Francisco., 2020.

(74) Ryckaert, J.-P.; Ciccotti, G.; Berendsen, H. J. C., Numerical integration of the cartesian equations of motion of a system with constraints: molecular dynamics of n-alkanes. J Comp Phys 1977, 23 (3), 327–341. DOI: 10.1016/0021-9991(77)90098-5.

(75) Roe, D. R.; Cheatham, T. E., 3rd, PTRAJ and CPPTRAJ: Software for Processing and Analysis of Molecular Dynamics Trajectory Data. J Chem Theory Comput 2013, 9 (7), 3084–95. DOI: 10.1021/ct400341p.

(76) Song, L. F.; Lee, T.-S.; Zhu, C.; York, D. M.; Merz, K. M., Jr., Using AMBER18 for relative free energy calculations. JCIM 2019, 59 (7), 3128–3135. DOI: 10.1021/acs.jcim.9b00105.

(77) Mey, A.; Allen, B. K.; Macdonald, H. E. B.; Chodera, J. D.; Hahn, D. F.; Kuhn, M.; Michel, J.; Mobley, D. L.; Naden, L..; Prasad, S.; Rizzi, A.; Scheen, J.; Shirts, M. R.; Tresadern, G.; Xu, H., Best practices for alchemical free energy calculations. Living J Comput Mol Sci 2020, 2 (1), 18378. DOI: 10.33011/livecoms.2.1.18378.

(78) Valdivia, A.; Rocha, M.; Luque, F. J., Mining Druggable Sites in Influenza A Hemagglutinin: Binding of the Pinanamine-Based Inhibitor M090. ACS Med Chem Lett 2025, 16 (1), 126–135. DOI: 10.1021/acsmedchemlett.4c00502.

(79) Zhiyi Wu, D. L. D. M. S. B. I. M. K., Irfan Alibay, Jérôme Hénin, Bryce K. Allen, Thomas T. Joseph, Hyungro Lee, Haoxi Li, Victoria Lim, Shuai Liu David Mobley, Domenico Marson, Pascal T. Merz, Michael R. Shirts, Alexander Schlaich, and Oliver Beckstein, alchemlyb: the simple alchemistry library. JOSS 2024, 9 (101), 6934. DOI: 10.21105/joss.06934.

(80) Shirts, M. R.; Chodera, J. D., Statistically optimal analysis of samples from multiple equilibrium states. J Chem Phys 2008, 129 (12), 124105. DOI: 10.1063/1.2978177.

(81) Chodera, J. D., A simple method for automated equilibration detection in molecular simulations. J Chem Theory Comput 2016, 12 (4), 1799–805. DOI: 10.1021/acs.jctc.5b00784.

(82) Hu, Y.; Muegge, I., In silico positional analogue scanning with amber GPU-TI. JCIM 2022, 62 (18), 4448–4459. DOI: 10.1021/acs.jcim.2c00860.

