## Supplement for "*N*-[(Thiophen-3-yl)methyl]benzamides as Influenza Virus Fusion Inhibitors Acting on H1 and H5 Hemagglutinins"

[†]Deceased.

### Table of contents


| | |
|---|---|
| Synthesis of starting materials | Page S4 |
| NMR spectra of key intermediates | Page S8 |
| $^1$H and $^{13}$C NMR spectra of compound **22** | Page S10 |
| $^1$H and $^{13}$C NMR spectra of compound **23** | Page S11 |
| $^1$H and $^{13}$C NMR spectra of compound **24** | Page S12 |
| $^1$H and $^{13}$C NMR spectra of compound **25** | Page S13 |
| $^1$H and $^{13}$C NMR spectra of compound **26** | Page S14 |
| $^1$H and $^{13}$C NMR spectra of compound **27** | Page S15 |
| $^1$H NMR spectrum of compound **28** | Page S16 |
| $^1$H and $^{13}$C NMR spectra of compound **29** | Page S17 |
| $^1$H and $^{13}$C NMR spectra of compound **30** | Page S18 |
| $^1$H and $^{13}$C NMR spectra of compound **31** | Page S19 |
| $^1$H and $^{13}$C NMR spectra of compound **32** | Page S20 |
| $^1$H and $^{13}$C NMR spectra of compound **33** | Page S21 |
| $^1$H and $^{13}$C NMR spectra of compound **34** | Page S22 |
| $^1$H and $^{13}$C NMR spectra of compound **35** | Page S23 |
| $^1$H and $^{13}$C NMR spectra of compound **36** | Page S24 |
| $^1$H and $^{13}$C NMR spectra of compound **37** | Page S25 |
| $^1$H and $^{13}$C NMR spectra of compound **38** | Page S26 |
| $^1$H and $^{13}$C NMR spectra of compound **39** | Page S27 |
| $^1$H and $^{13}$C NMR spectra of compound **40** | Page S28 |
| $^1$H and $^{13}$C NMR spectra of compound **41** | Page S29 |
| $^1$H and $^{13}$C NMR spectra of compound **42** | Page S30 |
| $^1$H and $^{13}$C NMR spectra of compound **43** | Page S31 |
| $^1$H and $^{13}$C NMR spectra of compound **44** | Page S32 |
| $^1$H and $^{13}$C NMR spectra of compound **45** | Page S33 |
| $^1$H and $^{13}$C NMR spectra of compound **46** | Page S34 |
| $^1$H and $^{13}$C NMR spectra of compound **47** | Page S35 |
| $^1$H and $^{13}$C NMR spectra of compound **48** | Page S36 |
| $^1$H and $^{13}$C NMR spectra of compound **49** | Page S37 |
| $^1$H and $^{13}$C NMR spectra of compound **50** | Page S38 |
| $^1$H and $^{13}$C NMR spectra of compound **51** | Page S39 |
| $^1$H and $^{13}$C NMR spectra of compound **52** | Page S40 |
| $^1$H and $^{13}$C NMR spectra of compound **53** | Page S41 |
| $^1$H and $^{13}$C NMR spectra of compound **54** | Page S42 |
| $^1$H and $^{13}$C NMR spectra of compound **55** | Page S43 |
| $^1$H and $^{13}$C NMR spectra of compound **56** | Page S44 |
| $^1$H and $^{13}$C NMR spectra of compound **57** | Page S45 |
| $^1$H and $^{13}$C NMR spectra of compound **58** | Page S46 |
| $^1$H and $^{13}$C NMR spectra of compound **59** | Page S47 |
| $^1$H and $^{13}$C NMR spectra of compound **60** | Page S48 |

$^{1}$H and $^{13}$C NMR spectra of compound **61** Page S49
$^{1}$H and $^{13}$C NMR spectra of compound **62** Page S50
$^{1}$H and $^{13}$C NMR spectra of compound **63** Page S51
$^{1}$H and $^{13}$C NMR spectra of compound **64** Page S52
$^{1}$H and $^{13}$C NMR spectra of compound **65** Page S53
Table S1. Elemental analysis data. Page S54
Figure S1. Sequence alignment for the studied Influenza A strains. Page S55
Figure S2. Distance between the carbonyl oxygen of (*S*)-F0045 and the hydroxyl oxygen of $T318_1$. Page S56
Figure S3. Representation of the two binding modes examined for VF-57a upon binding to site B in the H1 HA (A/Virginia/ATCC3/2009). Page S56
Figure S4. Distance between the carbonyl oxygen of **VF-57a** and the hydroxyl oxygen of $T318_1$. Page S57
Figure S5. Representation of the binding mode examined for VF-57a upon binding to site A. Page S57
Figure S6. RMSD plot for each HA-stem backbone atoms for the simulation of VF-57a in site A binding pocket of the A/Virginia/ATCC3/2009 Influenza A strain (H1N1). Page S58
Figure S7. Representation of the overlap between two monomers of HA. Page S58
Figure S8. Representation of the free energy change determined from the last 6ns of each λ divided in steps of 2 ns for **VF-57a**↔50 and TI analysis for the **VF-57a**↔50. Page S59
Figure S9. Representation of the free energy change determined from the last 6ns of each λ divided in steps of 2 ns for **VF-57a**↔22 and TI analysis for the **VF-57a**↔22. Page S60
Figure S10. Representation of the free energy change determined from the last 6ns of each λ divided in steps of 2 ns for **VF-57a**↔38 and TI analysis for the **VF-57a**↔38. Page S61
Figure S11. HPLC/UV (blank) Page S62
Figure S12. HPLC/UV (compound **VF-57a**) Page S63

#### SYNTHESIS OF STARTING MATERIALS

**(2,5-dimethylthiophen-3-yl)methanamine.** Thionyl chloride (2.32 mL, 3.81 g, 32.00 mmol) was added to a solution of 2,5-dimethylthiophen-3-carboxylic acid (1.0 g, 6.40 mmol) in anh. toluene (15.0 mL) and the complex mixture was kept under reflux for 2 h. After reaction completed, the solution was concentrated. Without further purification, the crude product 2,5-dimethylthiophen-3-carbonyl chloride was immediately used for the next step.

Ammonium hydroxide 25% aqueous solution (10.0 mL, 64.00 mmol) was added dropwise to a solution of 2,5-dimethylthiophen-3-carbonyl chloride in DCM (5.0 mL) at 0 °C and the mixture was allowed to warm up to room temperature and stirred for 1 h. Brine (30 mL) followed by DCM (20 mL) were added and the mixture was extracted. The aqueous layer was extracted again with DCM (2 x 20 mL). All the organic layers were joined, dried over anh. $Na_2SO_4$ and filtered to afford the crude product 2,5-dimethylthiophene-3-carboxamide that was used as such without further purification.

To a solution of 2,5-dimethylthiophene-3-carboxamide in anh. THF (10.0 mL) at 0 °C, $LiAlH_4$ 1 M in THF (32.0 mL, 32.00 mmol) was added dropwise. Then, the suspension was heated to reflux and stirred for 4 h. The reaction suspension was quenched adding water dropwise under ice-bath until no more bubbling was observed. Then, anh. $Na_2SO_4$ was added and the mixture was filtered through a pad with Celite® using MeOH (3 x 30 mL) as eluting agent. The solvent was concentrated *in vacuo* and the resulting crude was purified by column chromatography in silica gel (using as eluent mixtures of MeOH in DCM from 0% to 7%). Fractions containing the desired product were collected and concentrated *in vacuo*.

An excess of HCl 2 M in $Et_2O$ was added to the suspension of the amine in $Et_2O$ (5.0 mL) to form its hydrochloride salt, followed by filtration of the solid to afford the title compound as a brownish solid (620 mg, 55% yield). $^1$H NMR (400 MHz, $CD_3OD$) δ: 2.40 (m, 3H), 2.41 (s, 3H), 3.97 (s, 2H), 6.71 (s, 1H).

**(2-methylthiophen-3-yl)methanamine.** Thionyl chloride (6.96 mL, 11.42 g, 96.00 mmol) was added to a solution of 2-methylthiophen-3-carboxylic acid (1.36 g, 9.60 mmol) in anh. toluene (25.0 mL) followed by 20 drops of DMF and the mixture was kept under reflux for 2 h. After reaction completed, the solution was concentrated. Without further purification, the crude product 2-methylthiophen-3-carbonyl chloride was immediately used for the next step.

Ammonium hydroxide 25% aqueous solution (14.4 mL, 192.00 mmol) was added dropwise to a solution of 2-methylthiophen-3-carbonyl chloride in DCM (15 mL) at 0 °C and the mixture was allowed to warm up to room temperature and stirred for 1 h. Brine (40 mL) followed by DCM (30 mL) were added and the mixture was extracted. The aqueous layer was extracted again with DCM (2 x 30 mL). All the organic layers were joined, dried over anh. $Na_2SO_4$ and filtered to afford the crude product 2-methylthiophene-3-carboxamide that was used as such without further purification.

To a solution of 2-methylthiophene-3-carboxamide in anh. THF (40.0 mL) at 0 °C, $LiAlH_4$ (1.82 g, 48.00 mmol) was added portionwise. Then, the suspension was heated to reflux and stirred for 4 h. The reaction suspension was quenched adding water dropwise under ice-bath until no more bubbling was observed. Then, anh. $Na_2SO_4$ was added and the mixture was filtered through a pad with Celite® using MeOH (3 x 50 mL) as eluting agent. The solvent was concentrated *in vacuo* and the resulting crude was purified by column chromatography in silica gel (using as eluent mixtures of MeOH in DCM from 1% to 4%). Fractions containing the desired product were collected and concentrated *in vacuo*.

An excess of HCl 2 M in $Et_2O$ was added to the suspension of the amine in $Et_2O$ (10.0 mL) to form its hydrochloride, followed by filtration of the solid to afford the title compound as a beige solid (606 mg, 53% yield). $^1$H NMR (400 MHz, $CD_3OD$) δ: 2.50 (s, 3H), 4.07 (s, 2H), 7.05 (d, $J$ = 5.3 Hz, 1H), 7.25 (d, $J$ = 5.3 Hz, 1H).

**(5-methylthiophen-3-yl)methanamine.** Thionyl chloride (3.48 mL, 5.71 g, 48.00 mmol) was added to a solution of 5-methylthiophen-3-carboxylic acid (680 mg, 4.80 mmol) in anh. toluene (12.0 mL) followed by 10 drops of DMF and the mixture was kept under reflux for 2 h. After reaction completed, the solution was concentrated. Without further purification, the crude product 5-methylthiophen-3-carbonyl chloride was immediately used for the next step.

Ammonium hydroxide 25% aqueous solution (7.2 mL, 96.00 mmol) was added dropwise to a solution of 5-methylthiophen-3-carbonyl chloride in DCM (7 mL) at 0 °C and the mixture was allowed to warm up to room temperature and stirred for 1 h. Brine (20 mL) followed by DCM (15 mL) were added and the mixture was extracted. The aqueous layer was extracted again with DCM (2 x 15 mL). All the organic layers were joined, dried over anh. $Na_2SO_4$ and filtered to afford the crude product 5-methylthiophene-3-carboxamide that was used as such without further purification.

To a solution of 5-methylthiophene-3-carboxamide in anh. THF (20.0 mL) at 0 °C, $LiAlH_4$ (910 mg, 24.00 mmol) was added portionwise. Then, the suspension was heated to reflux and stirred for 4 h. The reaction suspension was quenched adding water dropwise under ice-bath until no more bubbling was observed. Then, anh. $Na_2SO_4$ was added and the mixture was filtered through a pad with Celite® using MeOH (3 x 25 mL) as eluting agent. Solvent was concentrated *in vacuo* and the resulting crude was purified by column chromatography in silica gel (using as eluent mixtures of MeOH in DCM from 1% to 5%). Fractions containing the desired product were collected and concentrated *in vacuo*.

An excess of HCl 2 M in $Et_2O$ was added to the suspension of the amine in $Et_2O$ (5.0 mL) to form its hydrochloride, followed by filtration of the solid to afford the title compound as a beige solid (127 mg, 16% yield). $^1H$ NMR (400 MHz, $CD_3OD$) δ: 2.48 (d, $J$ = 1.0 Hz, 3H), 4.05 (s, 2H), 6.87 (m, 1H), 7.26 (m, 1H).

**(2,5-dimethylthiophen-3-yl)methanol.** To a solution of ethyl 2,5-dimethylthiophene-3-carboxylate (100 mg, 0.64 mmol) in anh. THF (3.0 mL), $LiAlH_4$ (73 mg, 1.92 mmol) was added portinonwise under ice-bath. The mixture was stirred at RT for 16 h. Under an ice-bath, HCl 2 M (2.0 mL) was added and the mixture was stirred for 10 more min. Then, water (10 mL) followed by EtOAc (15 mL) were added and the mixture was extracted. The organic layer was washed again with brine (15 mL), then it was dried over anh. $Na_2SO_4$ and filtered. Solvents were concentrated *in vacuo* and the resulting crude was purified by column chromatography in silica gel (using as eluent mixtures of EtOAc in hexane from 0% to 15%). Fractions containing the desired product were collected and concentrated *in vacuo* to afford a colorless oil (88 mg, 97% yield). Spectroscopic data matched with those reported in SciFinder® (taken from Enamine Ltd., spectrum ID: EN300-55412).

#### NMR spectra of key intermediates

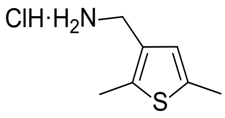


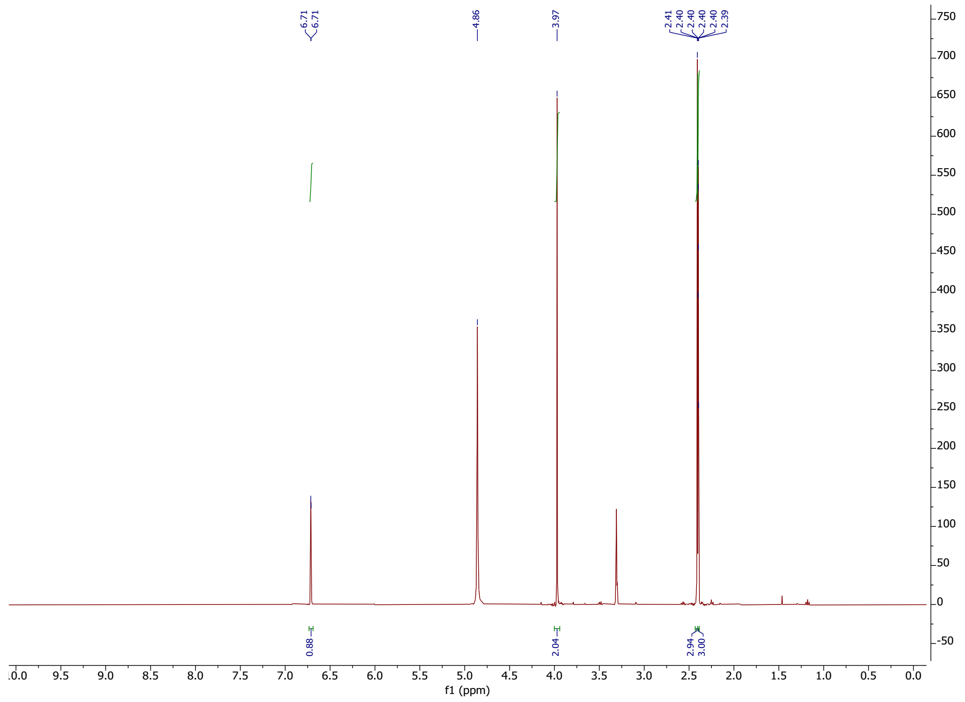


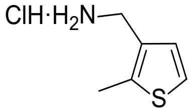


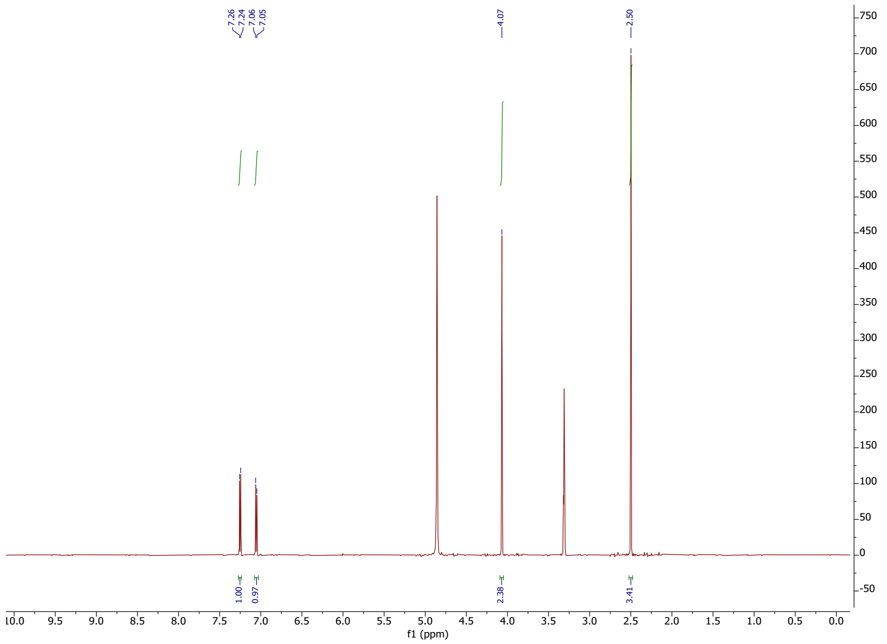

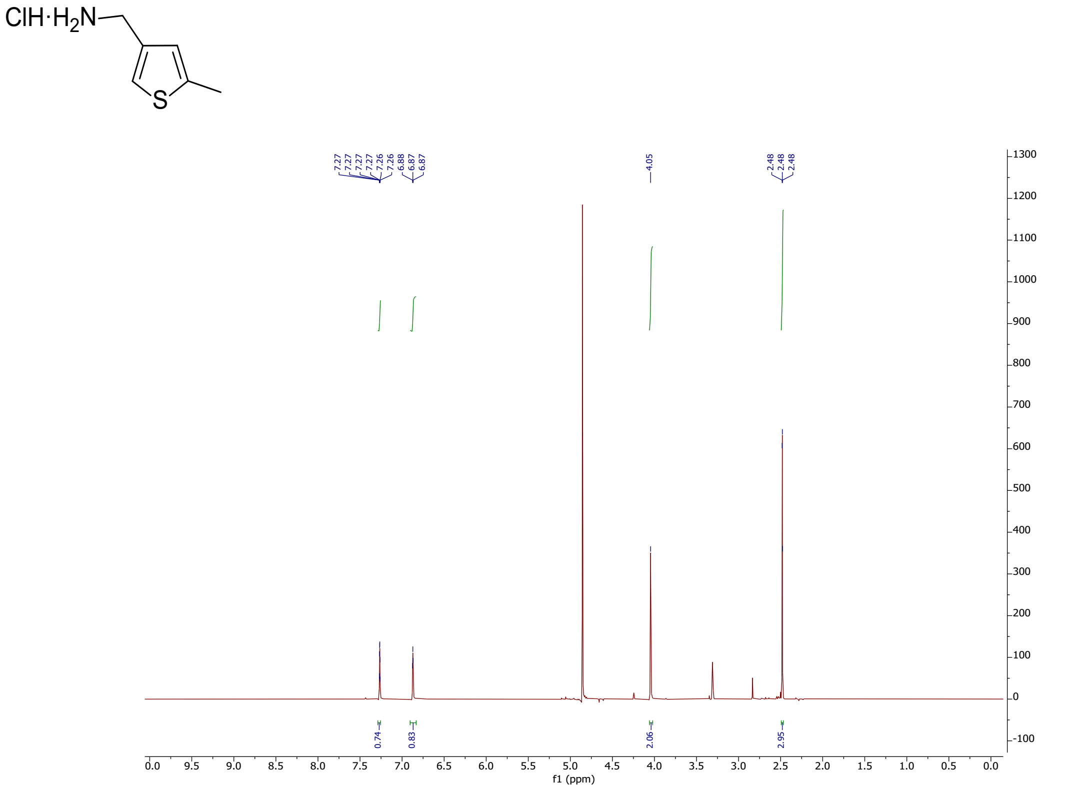
ClH·H2N
S
7.27
7.27
7.27
7.27
7.26
7.26
6.88
6.87
6.87
4.05
2.48
2.48
2.48
0.74
0.83
2.06
2.95
f1 (ppm)

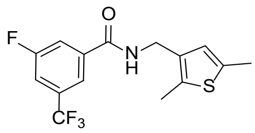

**22**

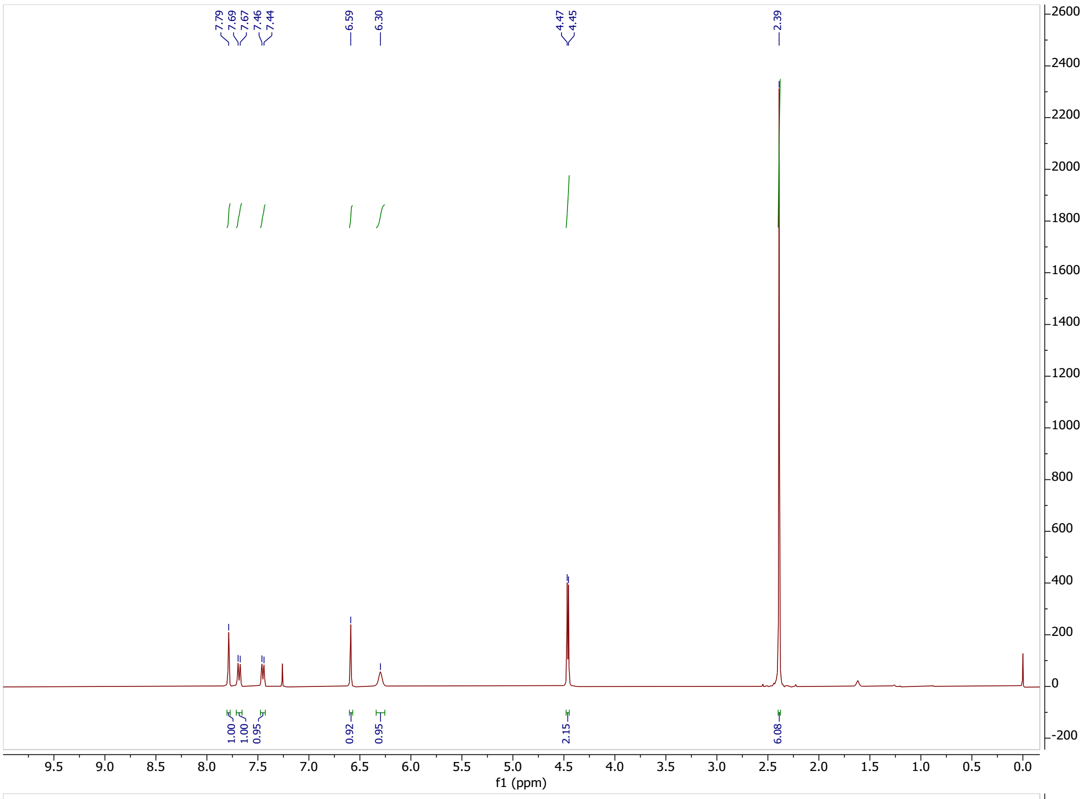

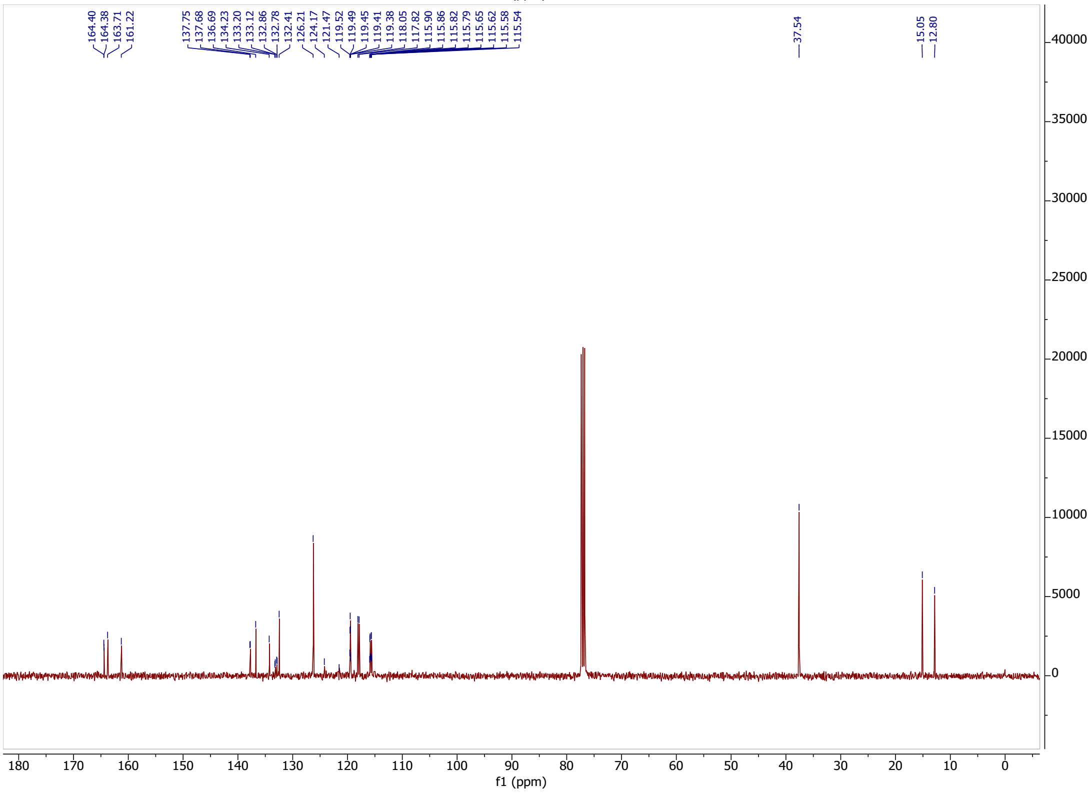

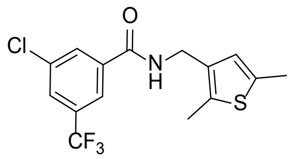
O
Cl
N
H
S
CF3

**23**

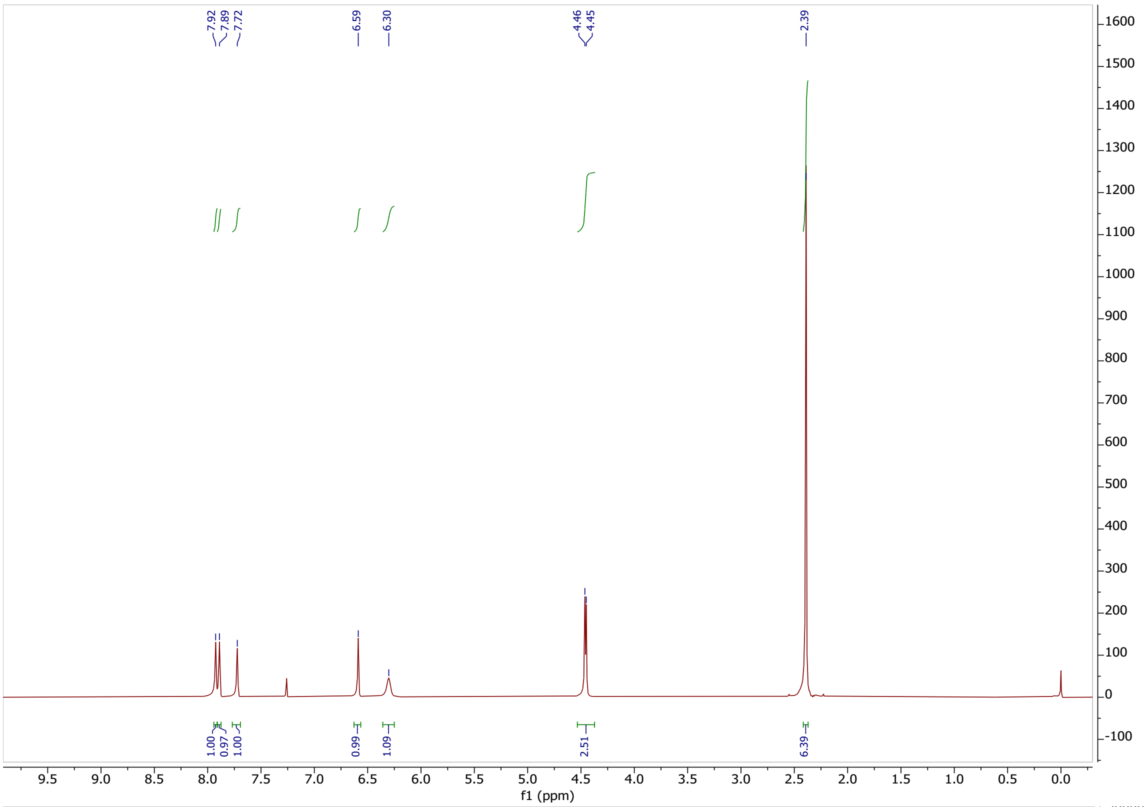
7.92
7.89
7.72
6.59
6.30
4.46
4.45
2.39
1.00
0.97
1.00
0.99
1.09
2.51
6.39
f1 (ppm)

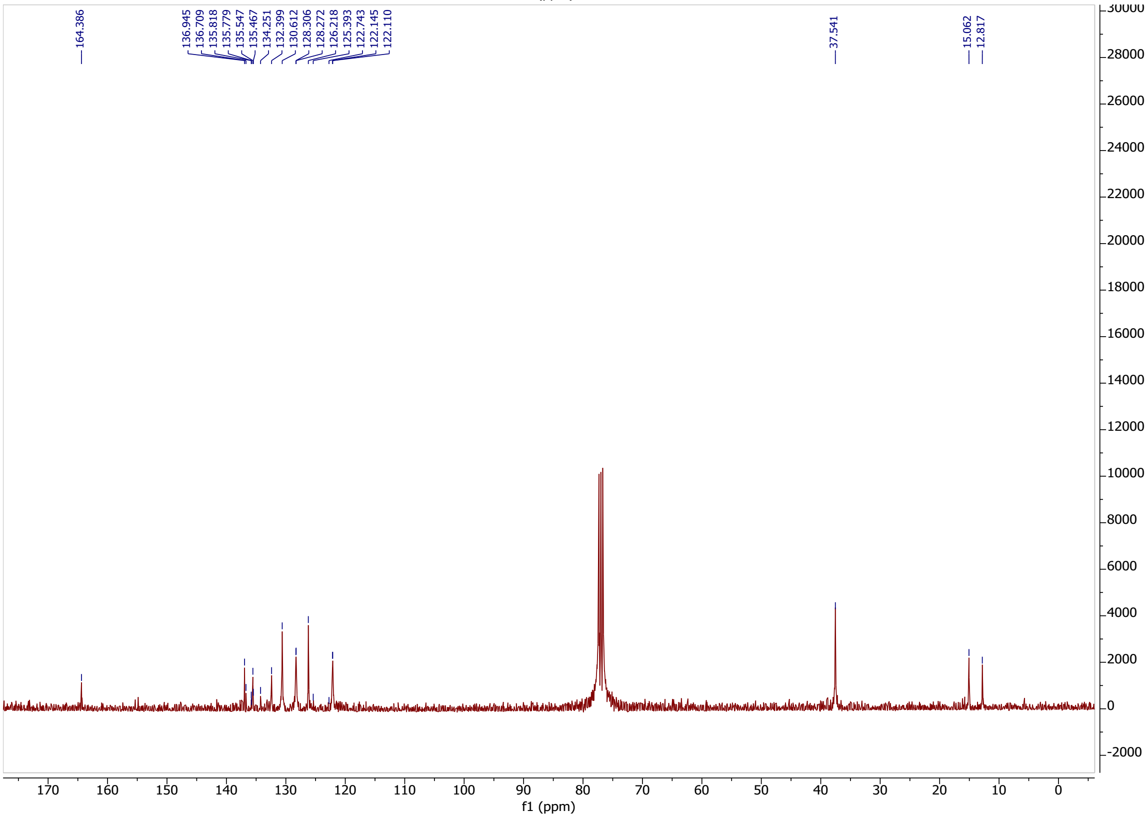
164.386
136.945
136.709
135.818
135.779
135.547
135.467
134.251
132.399
130.612
128.306
128.272
126.218
125.393
122.743
122.145
122.110
37.541
15.062
12.817
f1 (ppm)

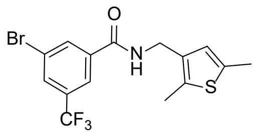
Br
O
N
H
S
CF3

**24**

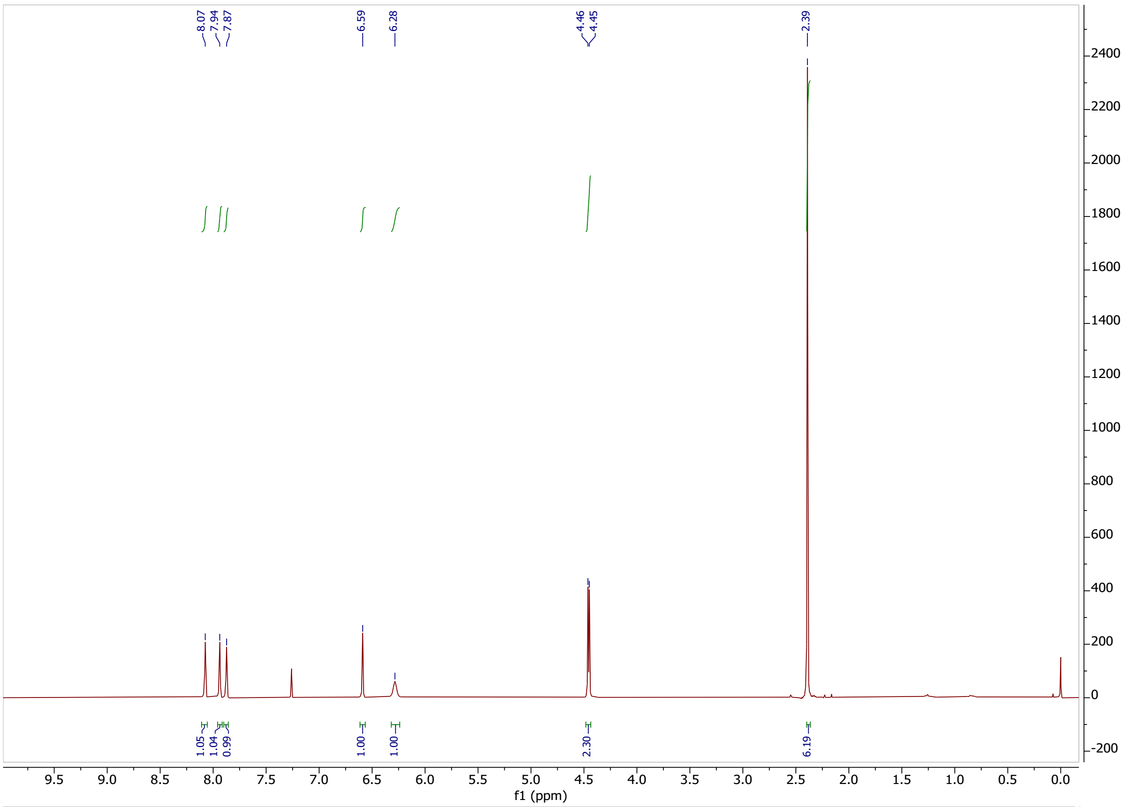
8.07
7.94
7.87
6.59
6.28
4.46
4.45
2.39
1.05
1.04
0.99
1.00
1.00
2.30
6.19
f1 (ppm)

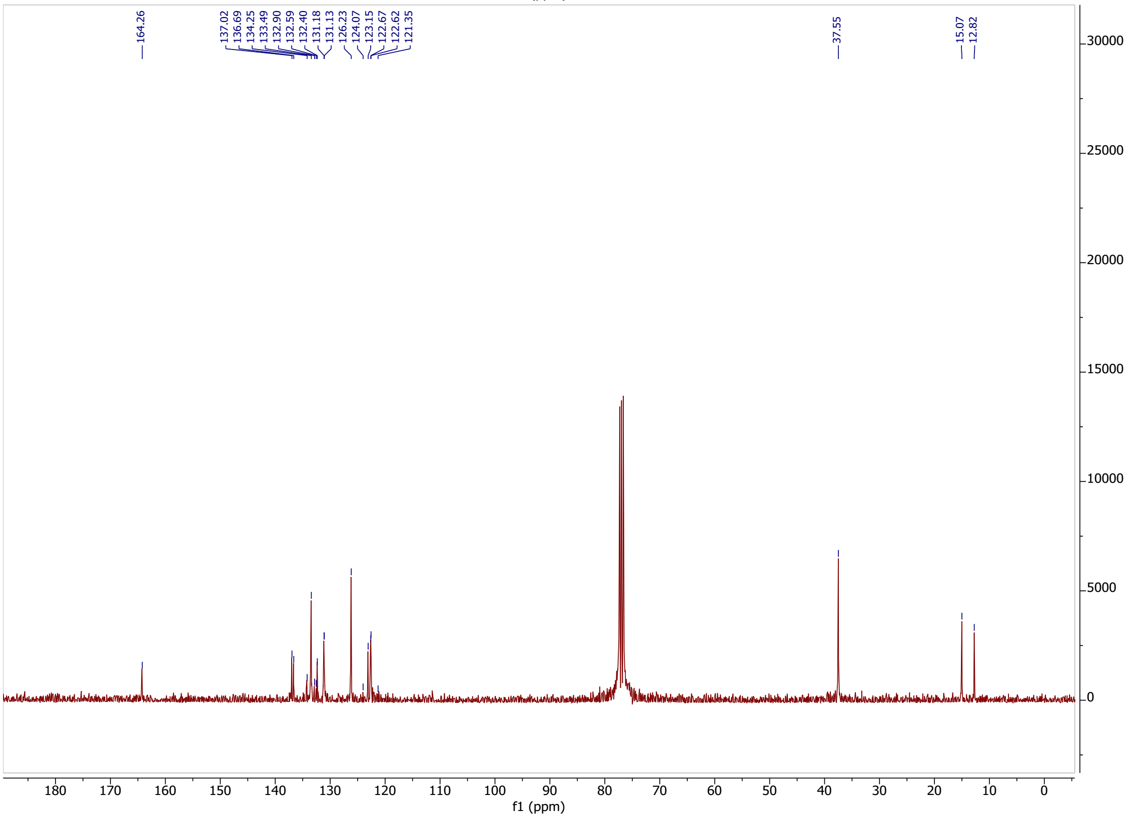
164.26
137.02
136.69
134.25
133.49
132.90
132.59
132.40
131.18
131.13
126.23
124.07
123.15
122.67
122.62
121.35
37.55
15.07
12.82
f1 (ppm)

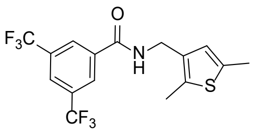
O
F3C
N
H
S
CF3

**25**

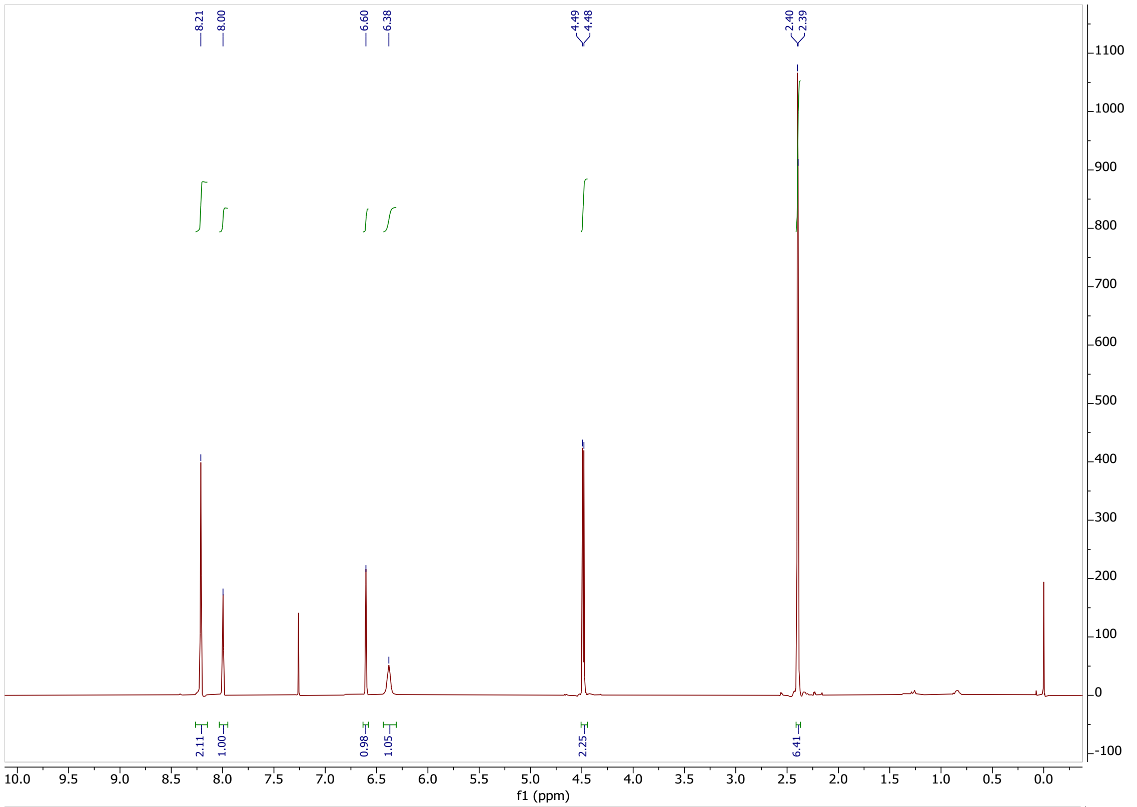
8.21
8.00
6.60
6.38
4.49
4.48
2.40
2.39
2.11
1.00
0.98
1.05
2.25
6.41
f1 (ppm)

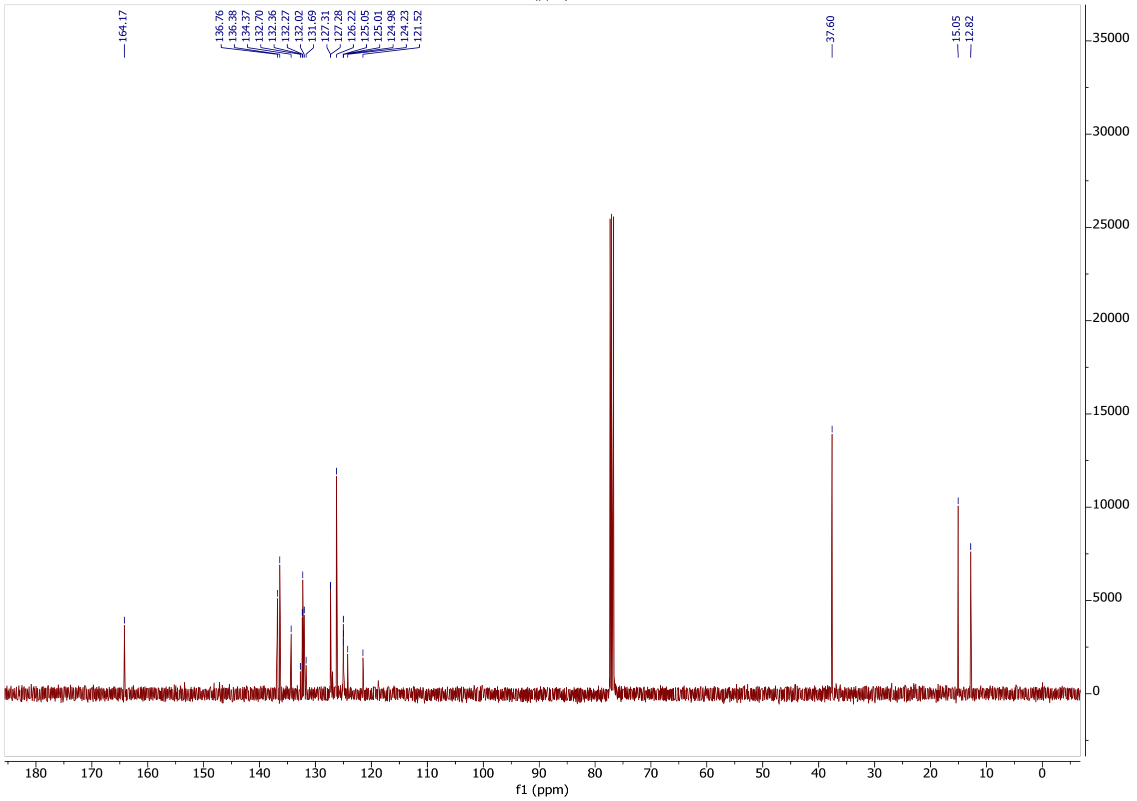
164.17
136.76
136.38
134.37
132.70
132.36
132.27
132.02
131.69
127.31
127.28
126.22
125.05
125.01
124.98
124.23
121.52
37.60
15.05
12.82
f1 (ppm)

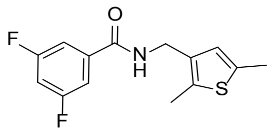

**26**

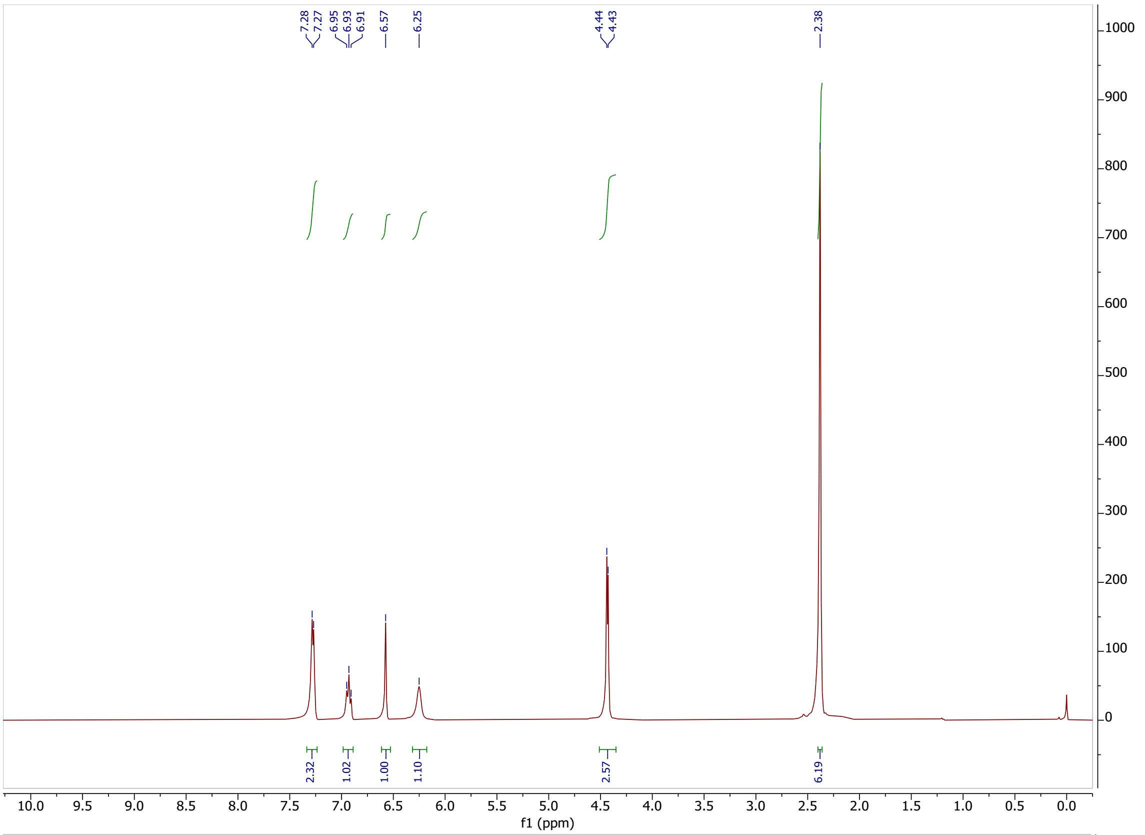

7.28
7.27
6.95
6.93
6.91
6.57
6.25
4.44
4.43
2.38
2.32
1.02
1.00
1.10
2.57
6.19
f1 (ppm)


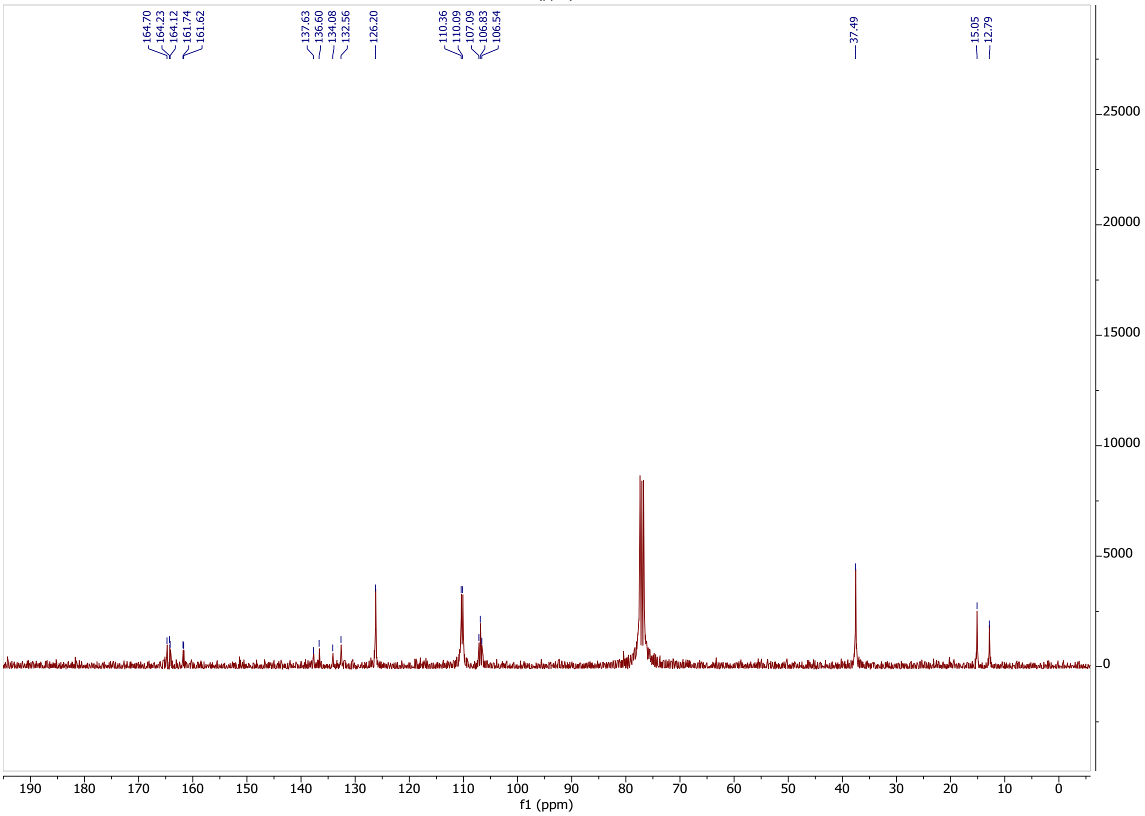

164.70
164.23
164.12
161.74
161.62
137.63
136.60
134.08
132.56
126.20
110.36
110.09
107.09
106.83
106.54
37.49
15.05
12.79
f1 (ppm)

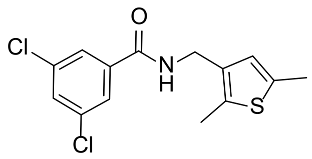

O
Cl
N
H
S
Cl


**27**

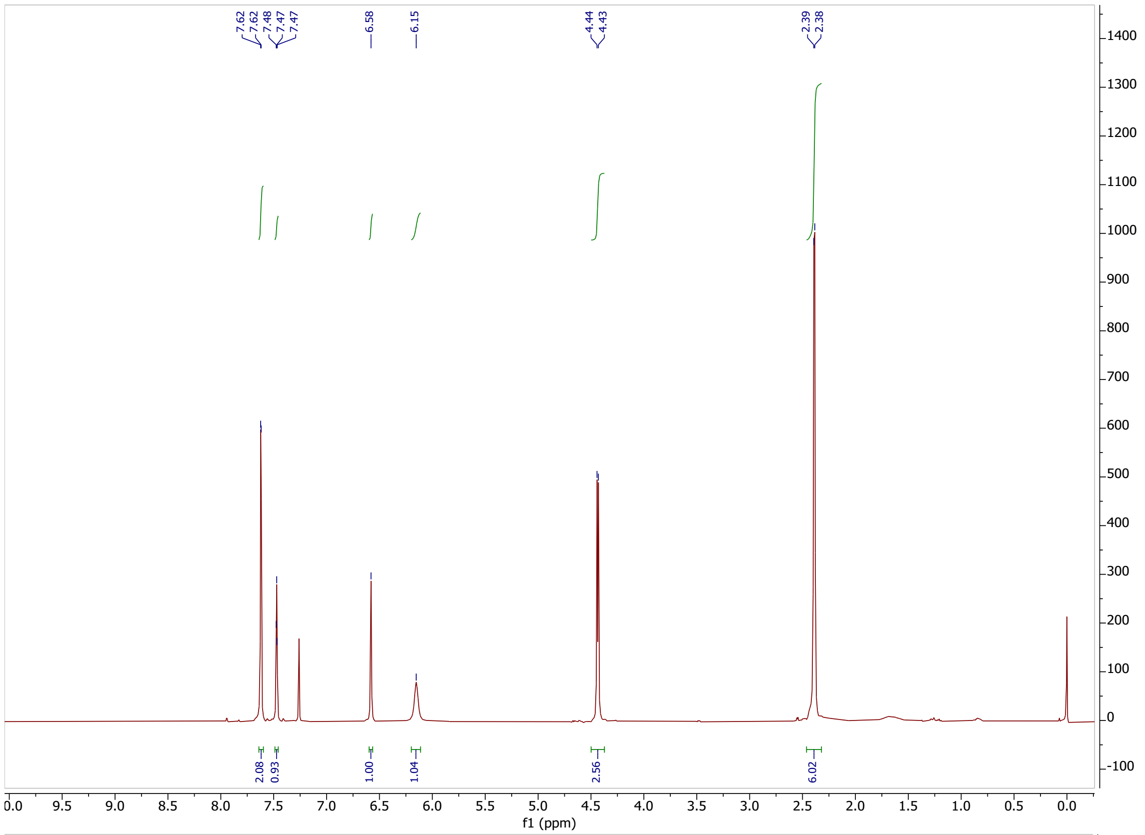

7.62
7.62
7.48
7.47
7.47
6.58
6.15
4.44
4.43
2.39
2.38
2.08
0.93
1.00
1.04
2.56
6.02
f1 (ppm)


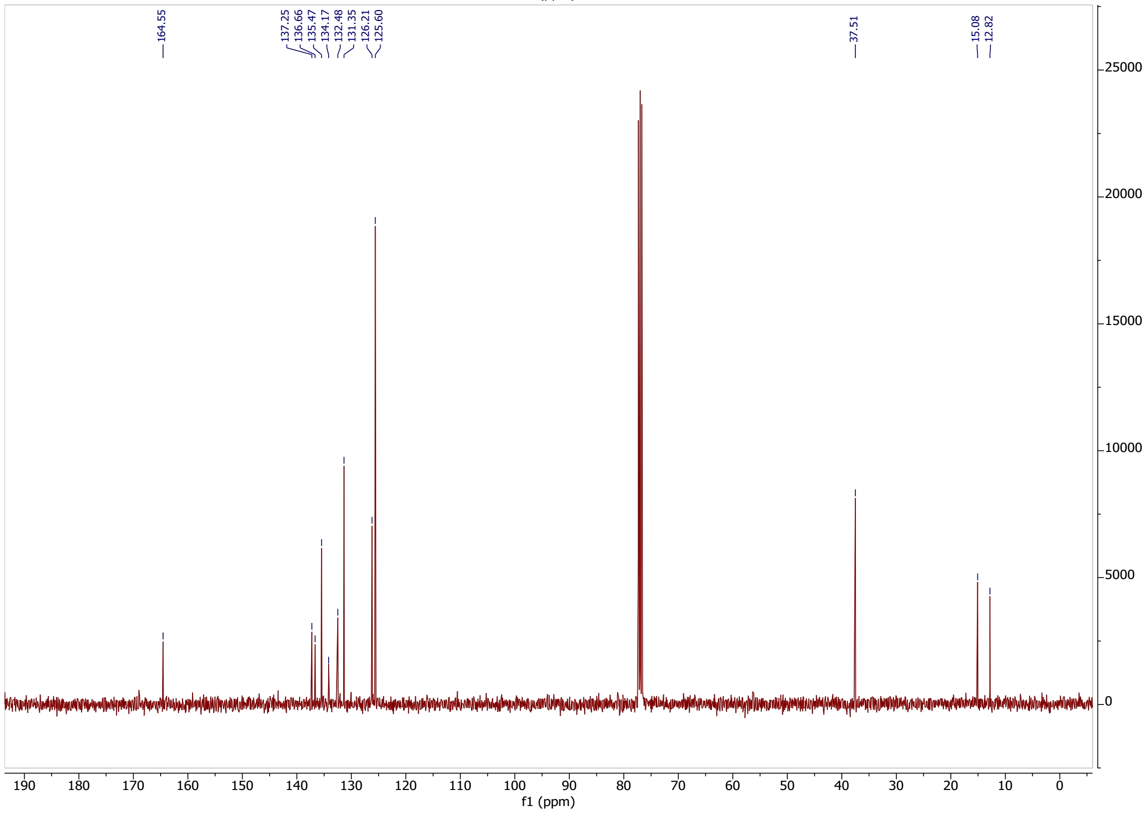

164.55
137.25
136.66
135.47
134.17
132.48
131.35
126.21
125.60
37.51
15.08
12.82
f1 (ppm)

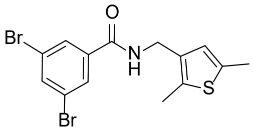

**28**

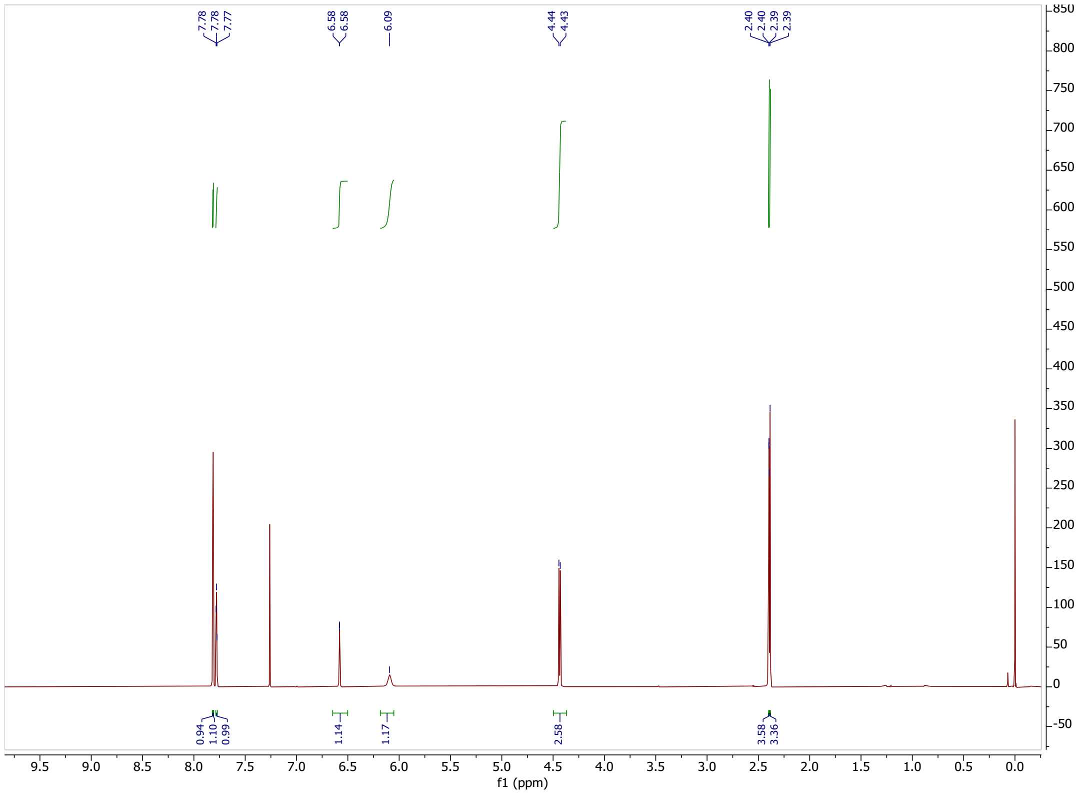

7.78
7.78
7.77
6.58
6.58
6.09
4.44
4.43
2.40
2.40
2.39
2.39
0.94
1.10
0.99
1.14
1.17
2.58
3.58
3.36
f1 (ppm)

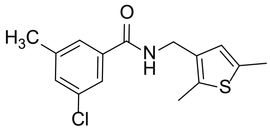

**29**

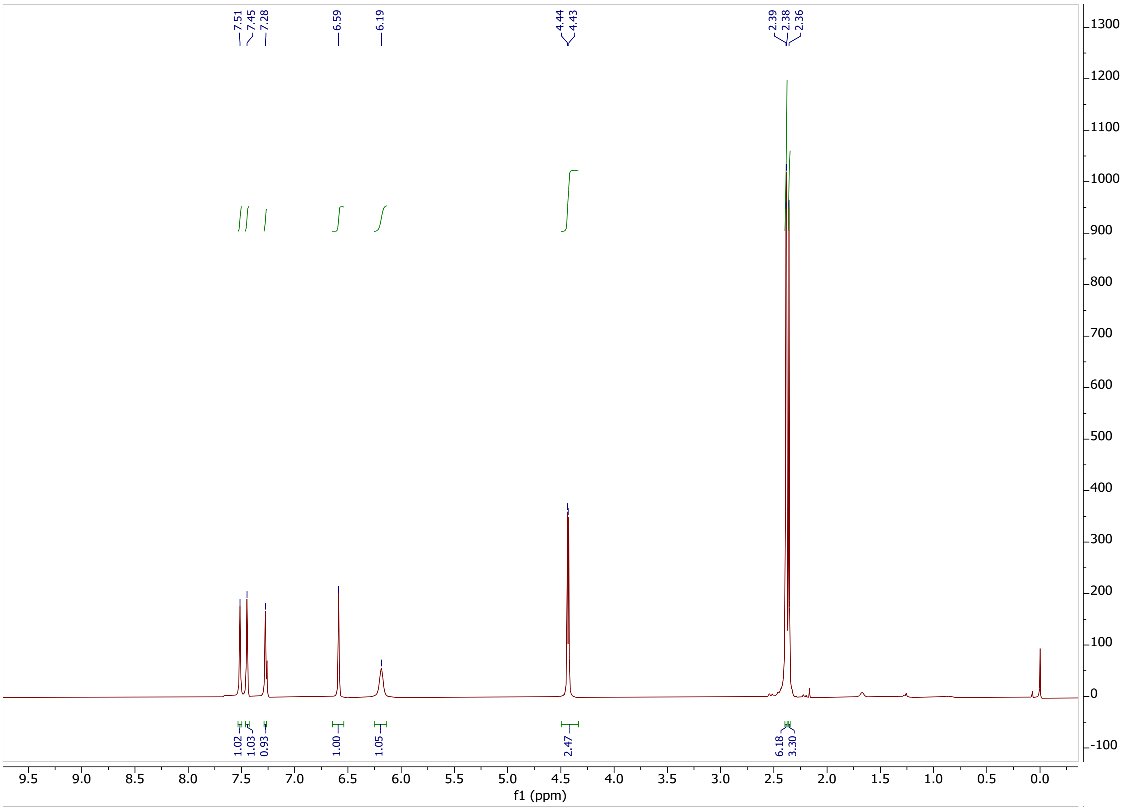

7.51
7.45
7.28
6.59
6.19
4.44
4.43
2.39
2.38
2.36
1.02
1.03
0.93
1.00
1.05
2.47
6.18
3.30
f1 (ppm)


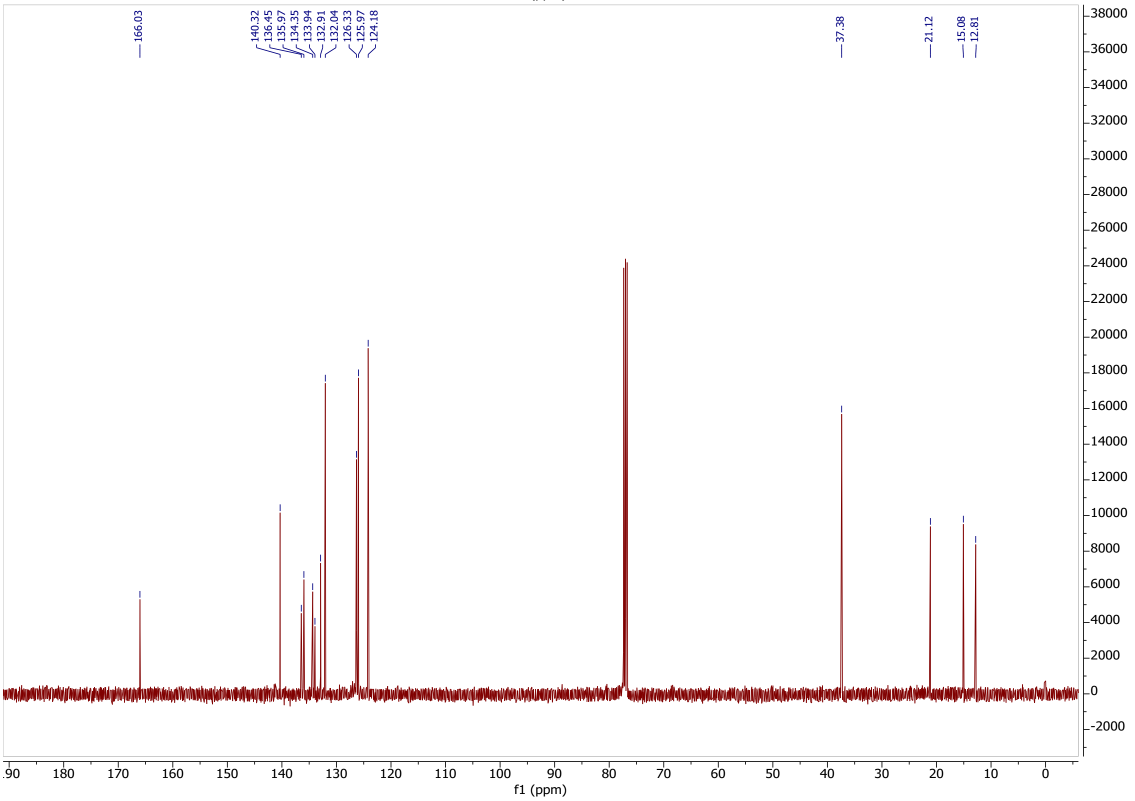

166.03
140.32
136.45
135.97
134.35
133.94
132.91
132.04
126.33
125.97
124.18
37.38
21.12
15.08
12.81
f1 (ppm)

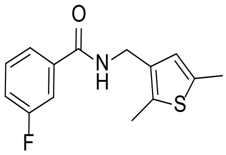

**30**

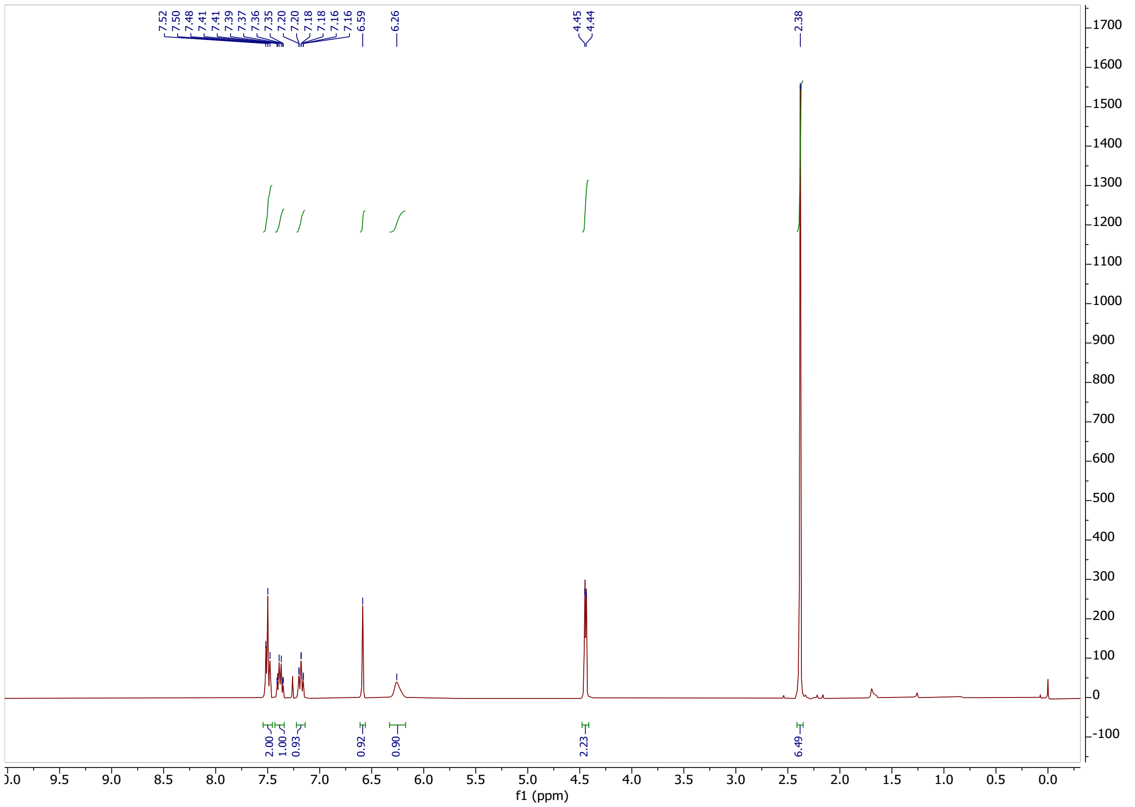

7.52 7.50 7.48 7.41 7.41 7.39 7.37 7.36 7.35 7.20 7.20 7.18 7.18 7.16 7.16 6.59
6.26
4.45 4.44
2.38
2.00 1.00 0.93 0.92 0.90 2.23 6.49
f1 (ppm)


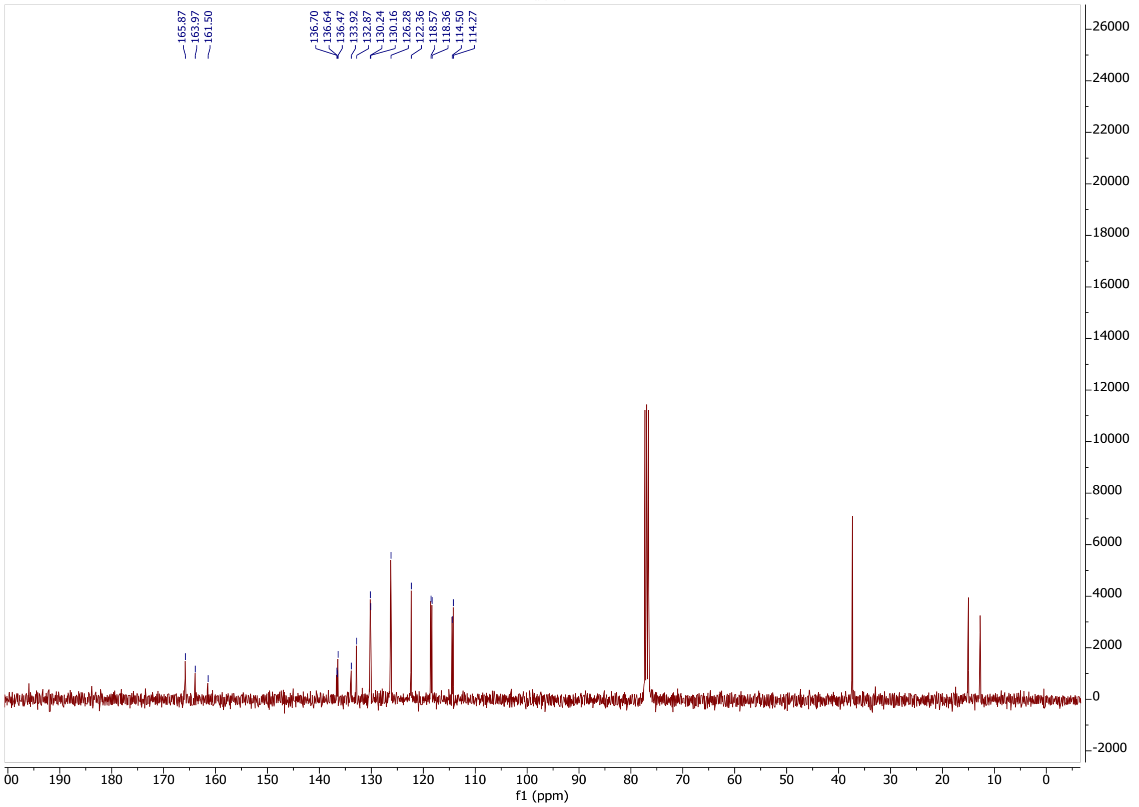

165.87 163.97 161.50
136.70 136.64 136.47 133.92 132.87 130.24 130.16 126.28 122.36 118.57 118.36 114.50 114.27
f1 (ppm)

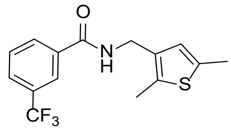
O
N
H
S
CF3

**31**

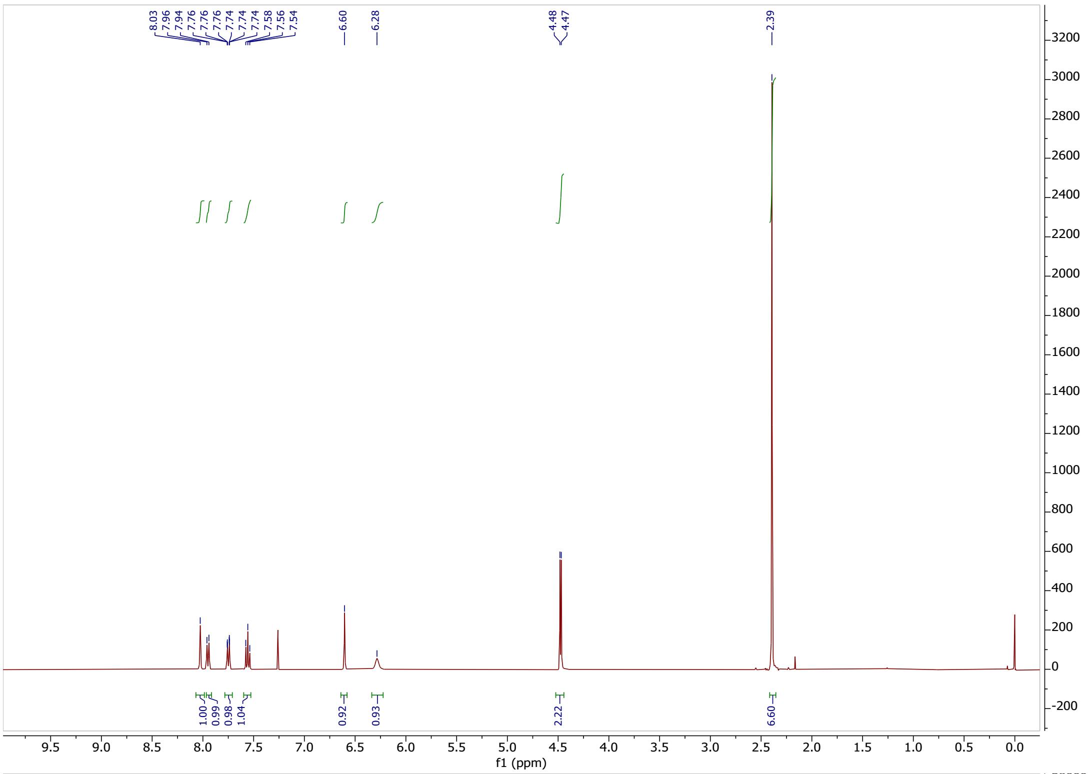
8.03
7.96
7.94
7.76
7.76
7.76
7.74
7.74
7.74
7.58
7.56
7.54
6.60
6.28
4.48
4.47
2.39
1.00
0.99
0.98
1.04
0.92
0.93
2.22
6.60
f1 (ppm)

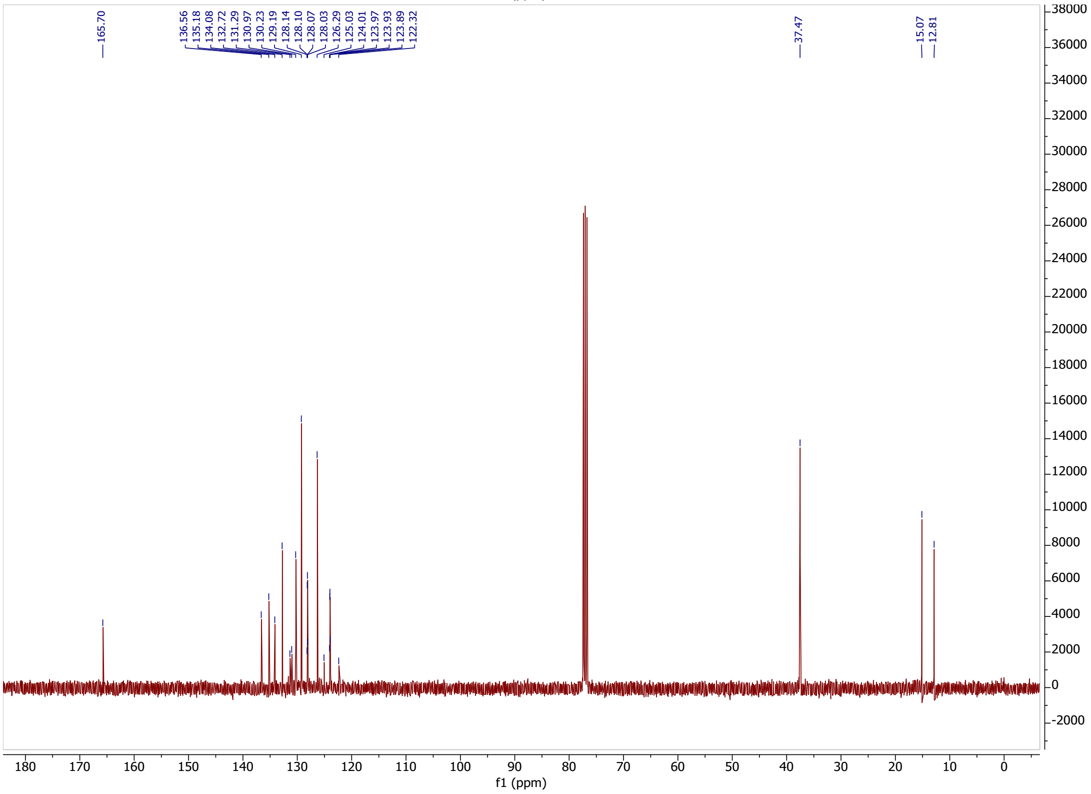
165.70
136.56
135.18
134.08
132.72
131.29
130.97
130.23
129.19
128.14
128.10
128.07
128.03
126.29
125.03
124.01
123.97
123.93
123.89
122.32
37.47
15.07
12.81
f1 (ppm)

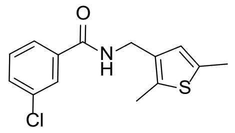
O
N
H
S
Cl

**32**

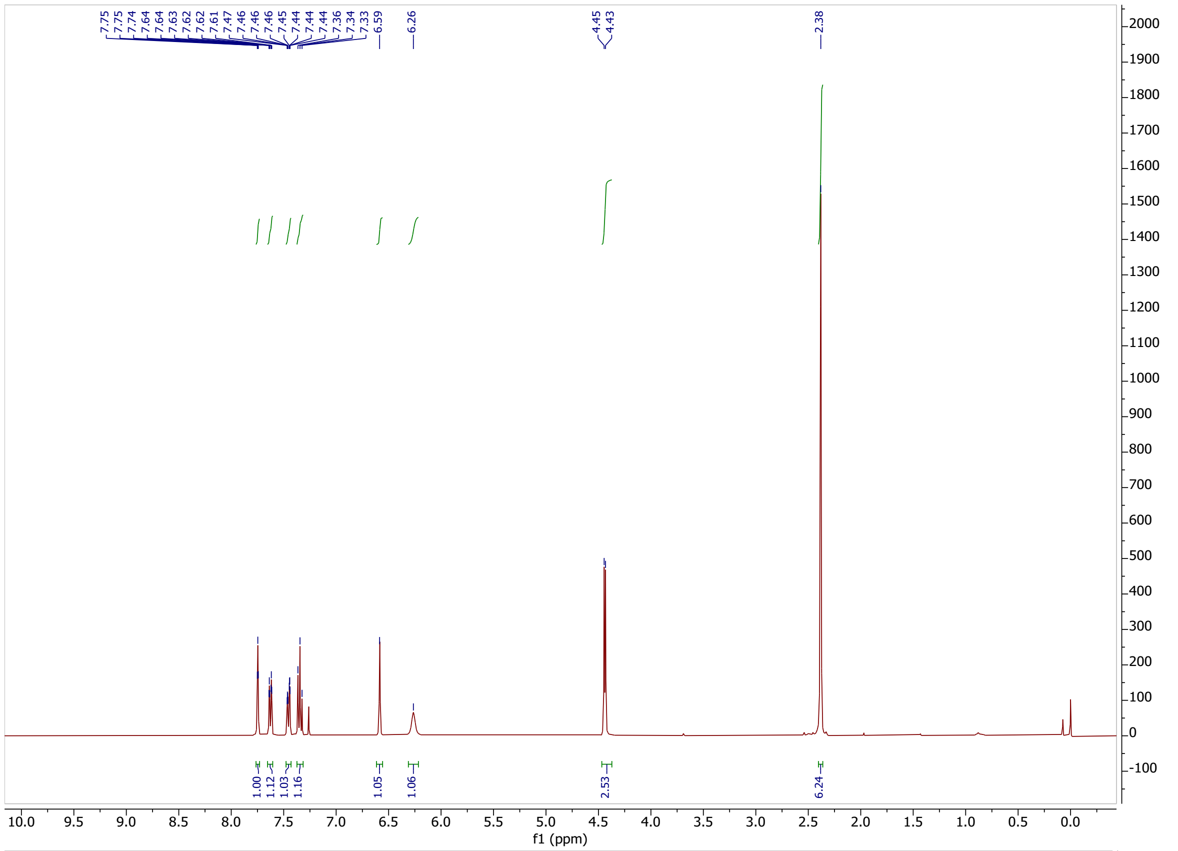
7.75
7.75
7.74
7.64
7.64
7.63
7.62
7.62
7.61
7.47
7.46
7.46
7.45
7.44
7.44
7.44
7.36
7.34
7.33
6.59
6.26
4.45
4.43
2.38
1.00
1.12
1.03
1.16
1.05
1.06
2.53
6.24
f1 (ppm)

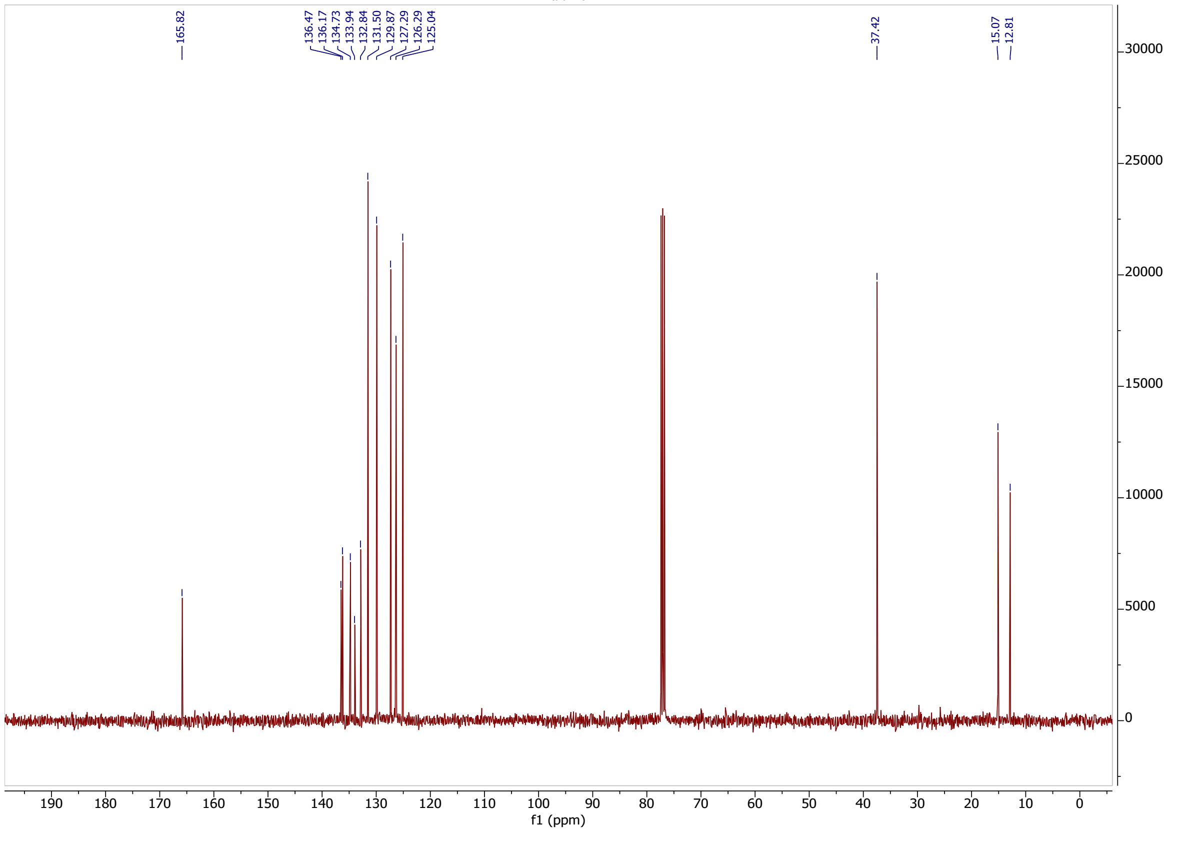
165.82
136.47
136.17
134.73
133.94
132.84
131.50
129.87
127.29
126.29
125.04
37.42
15.07
12.81
f1 (ppm)

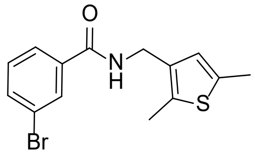
O
N
H
S
Br

**33**

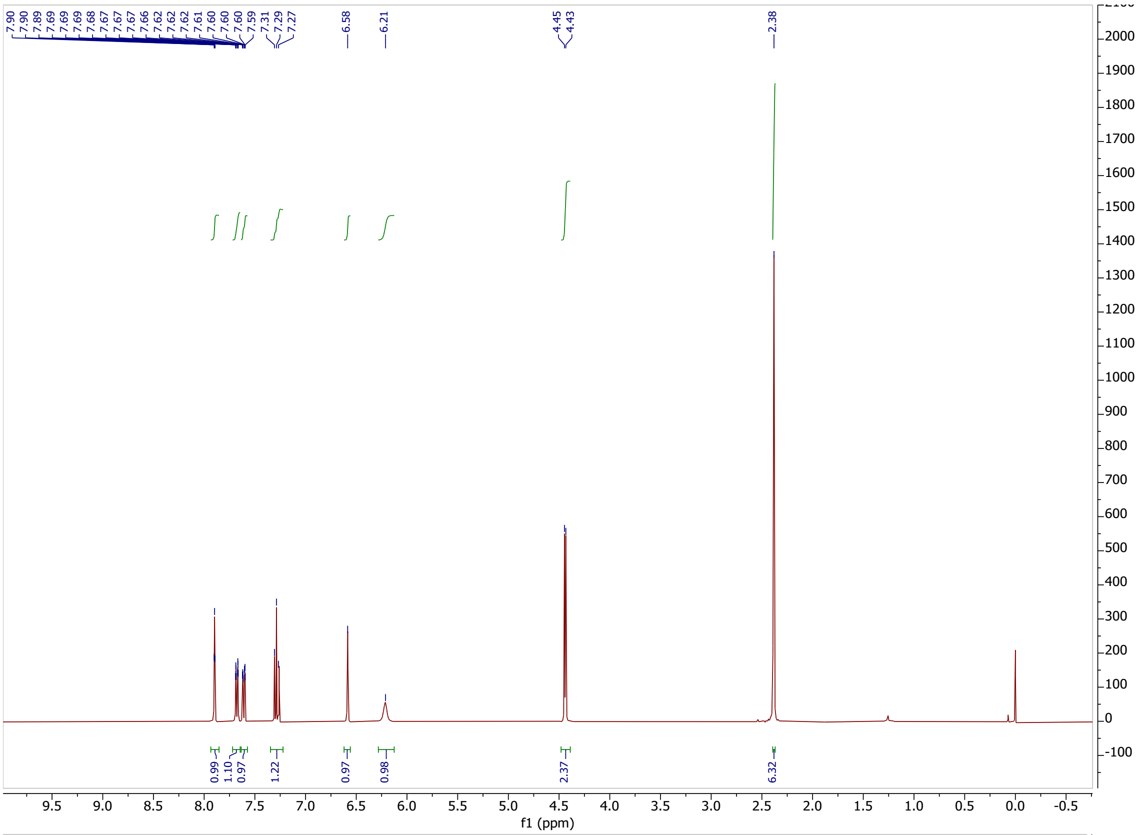
7.90
7.90
7.89
7.69
7.69
7.69
7.68
7.67
7.67
7.66
7.62
7.62
7.62
7.61
7.60
7.60
7.60
7.59
7.31
7.29
7.27
6.58
6.21
4.45
4.43
2.38
0.99
1.10
0.97
1.22
0.97
0.98
2.37
6.32
f1 (ppm)

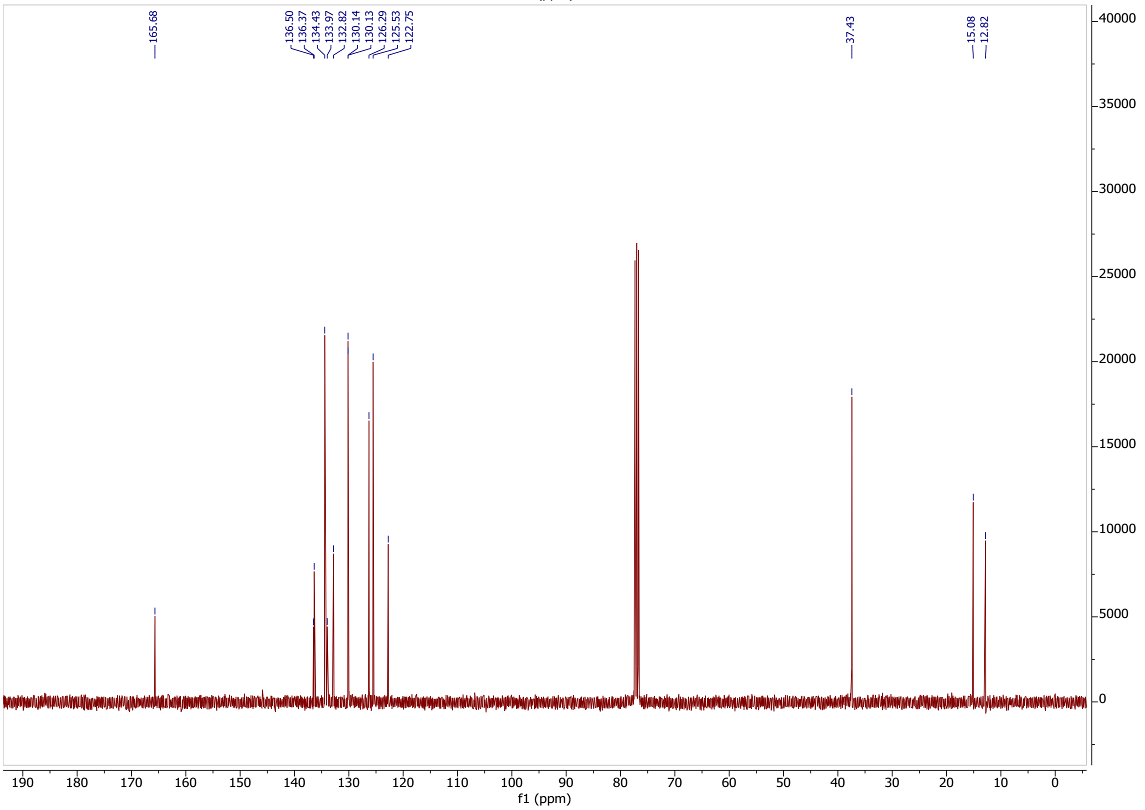
165.68
136.50
136.37
134.43
133.97
132.82
130.14
130.13
126.29
125.53
122.75
37.43
15.08
12.82
f1 (ppm)

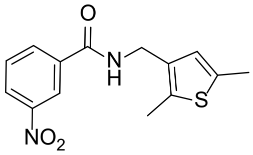


**34**

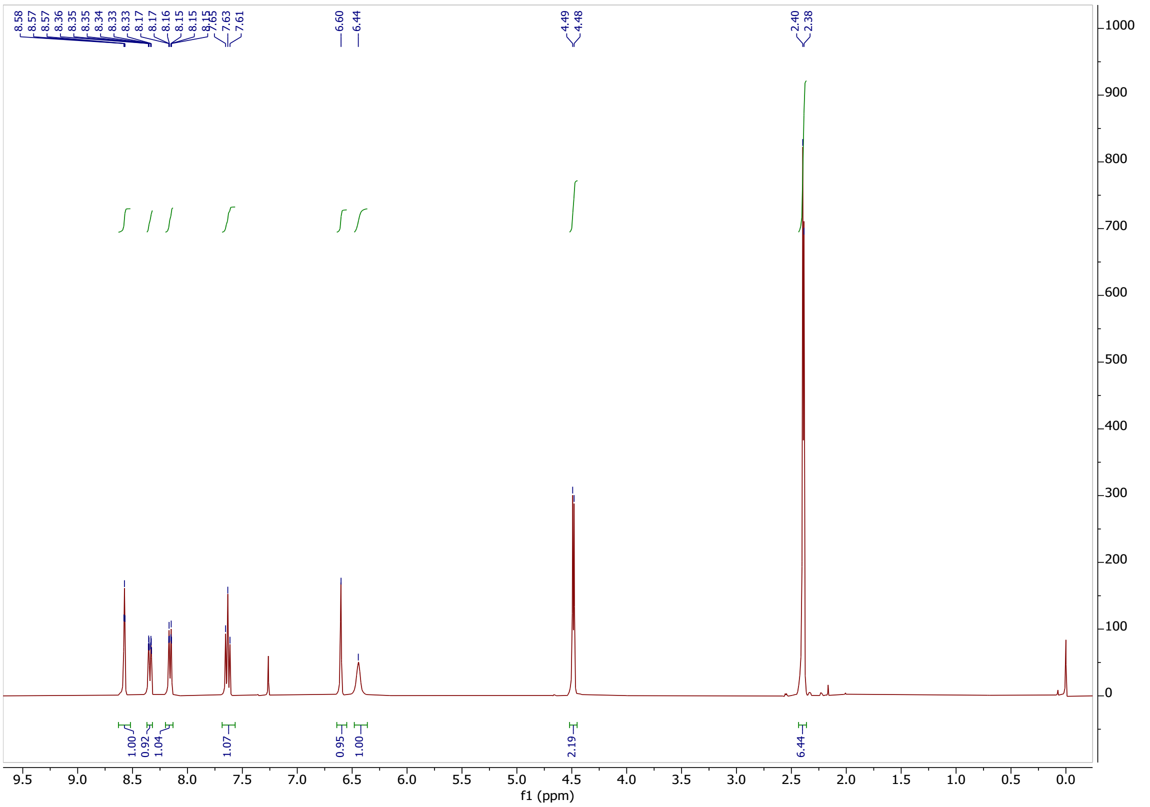

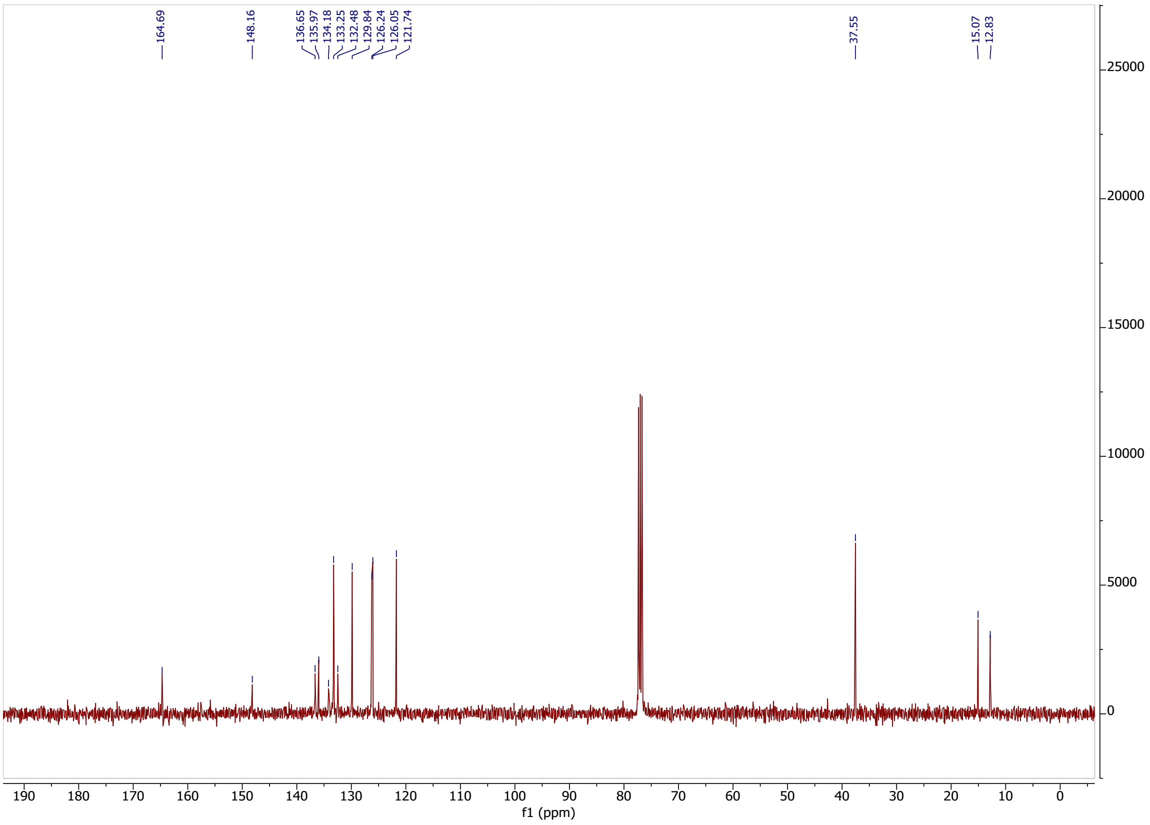

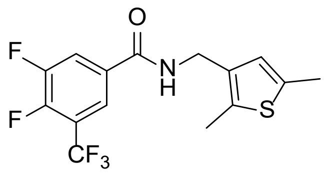

**35**

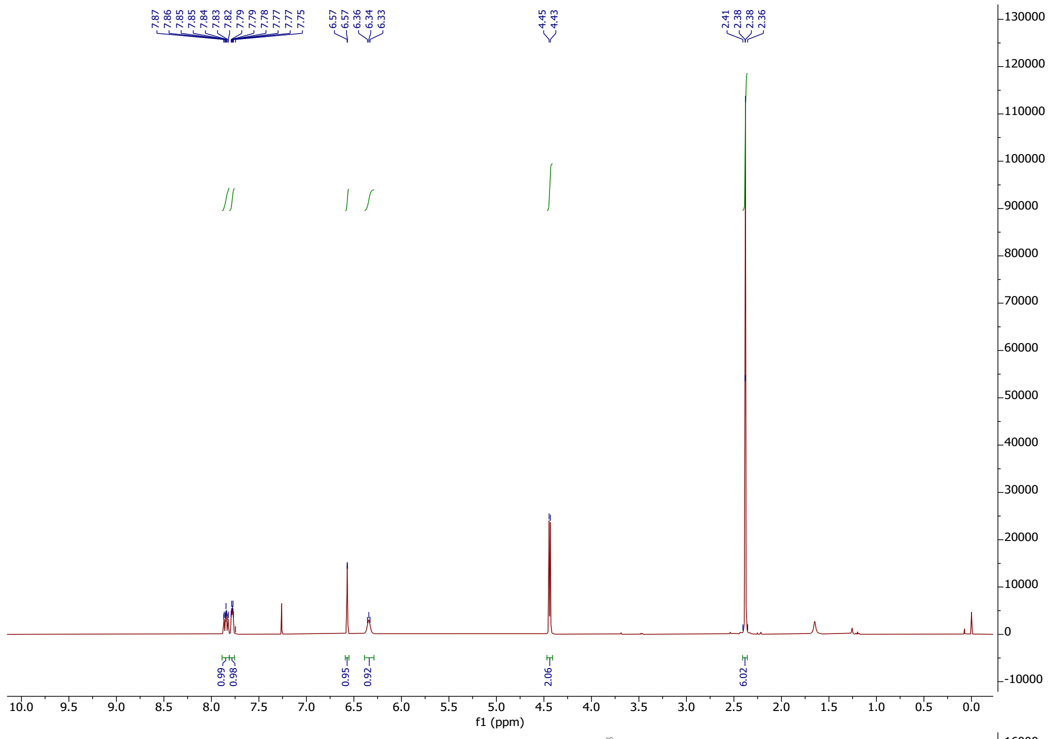

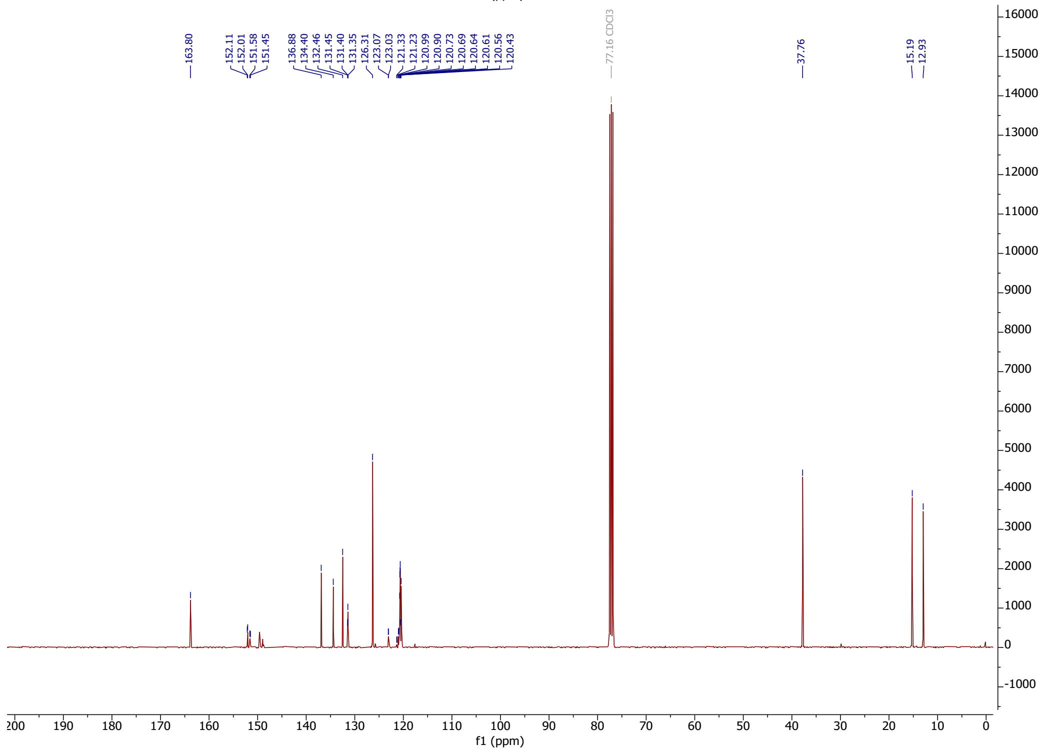

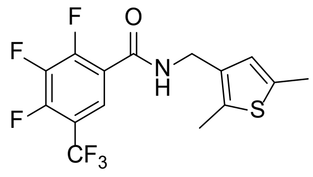
F
O
F
N
H
F
S
CF3

**36**

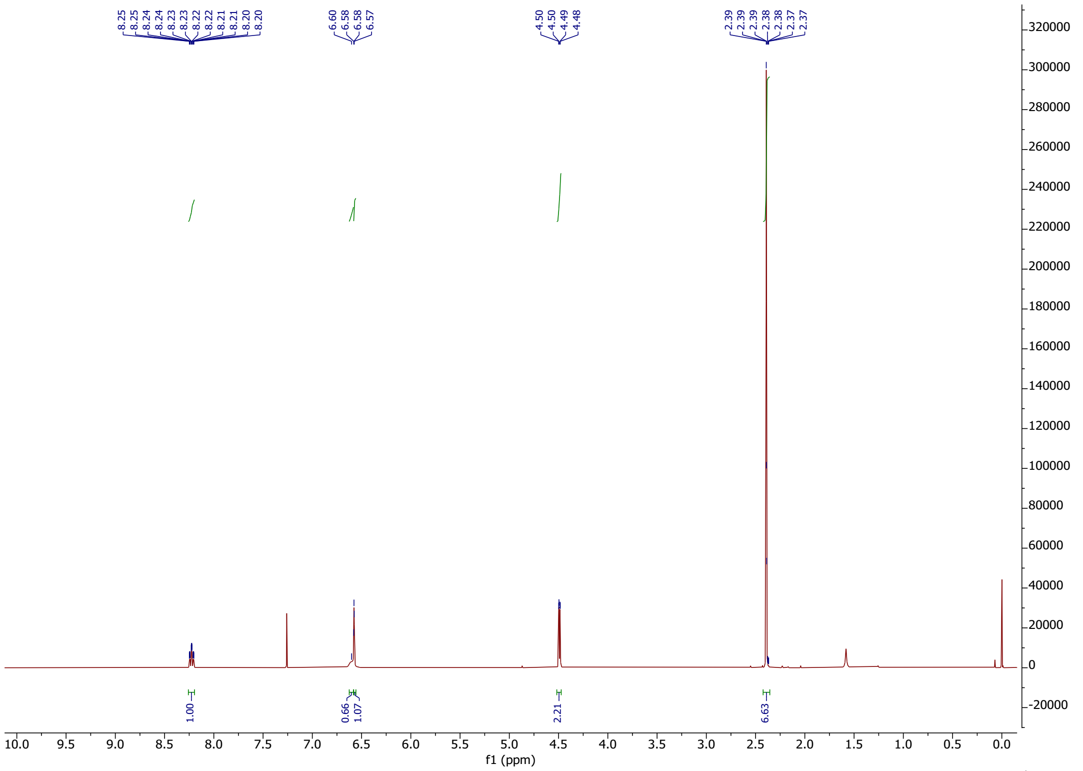
8.25
8.25
8.24
8.23
8.23
8.22
8.22
8.21
8.21
8.20
8.20
6.60
6.58
6.58
6.57
4.50
4.50
4.49
4.48
2.39
2.39
2.39
2.38
2.38
2.37
2.37
1.00
0.66
1.07
2.21
6.63
f1 (ppm)

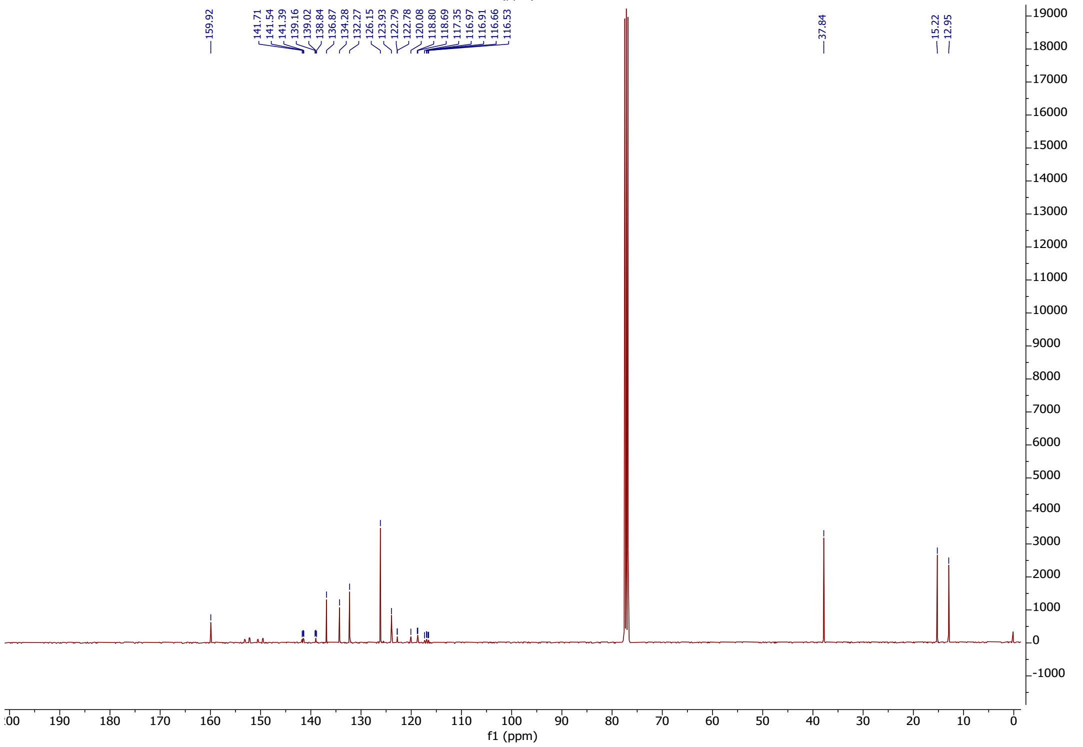
159.92
141.71
141.54
141.39
139.16
139.02
138.84
136.87
134.28
132.27
126.15
123.93
122.79
122.78
120.08
118.80
118.69
117.35
116.97
116.91
116.66
116.53
37.84
15.22
12.95
f1 (ppm)

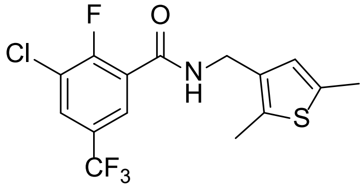

**37**

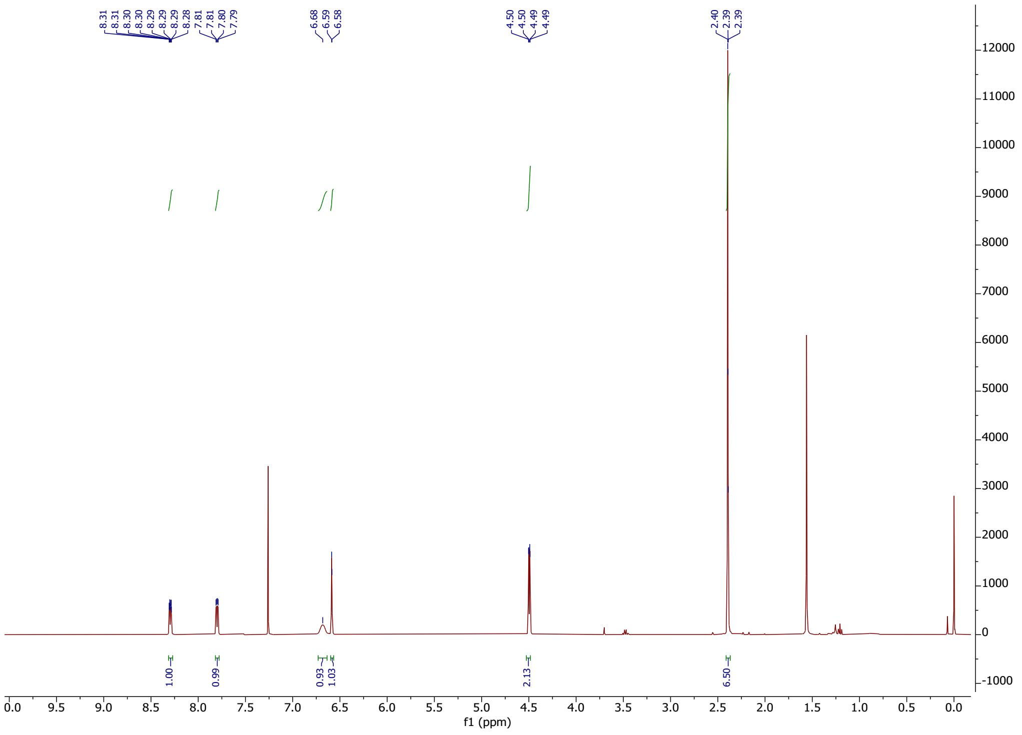


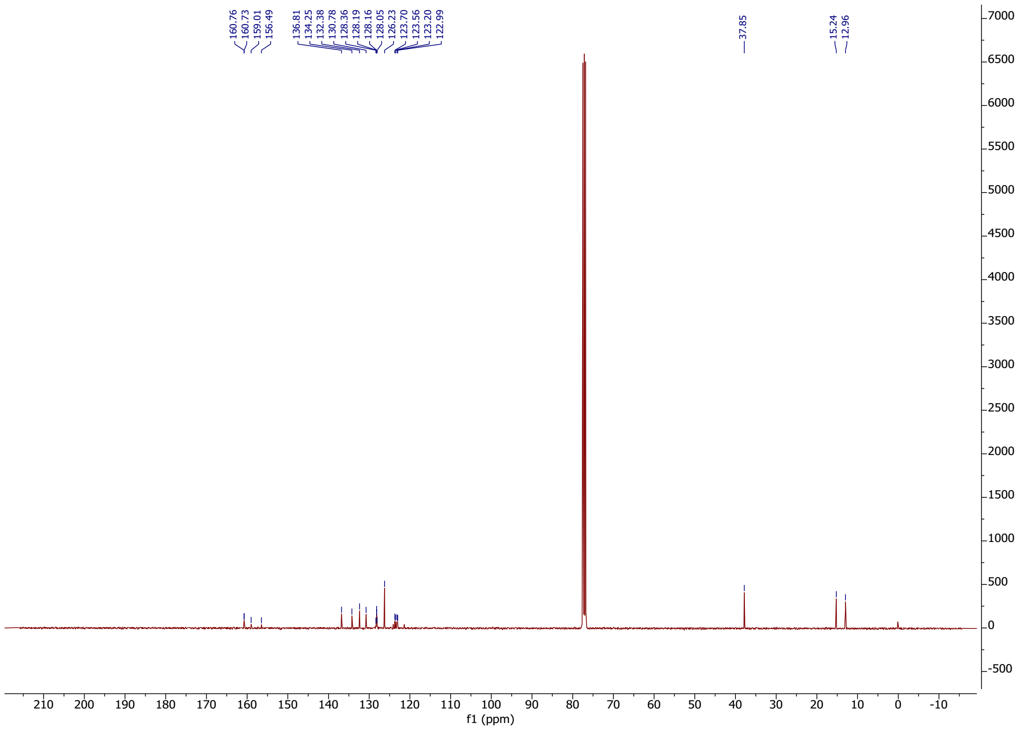

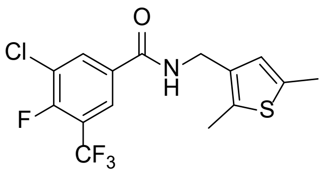

**38**

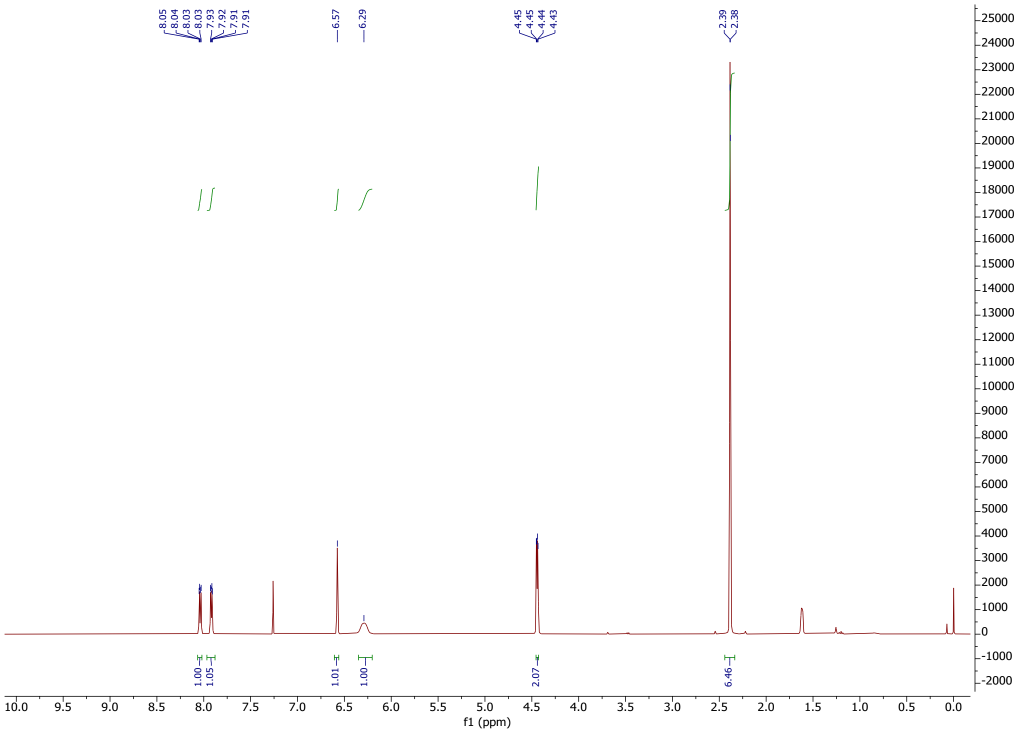

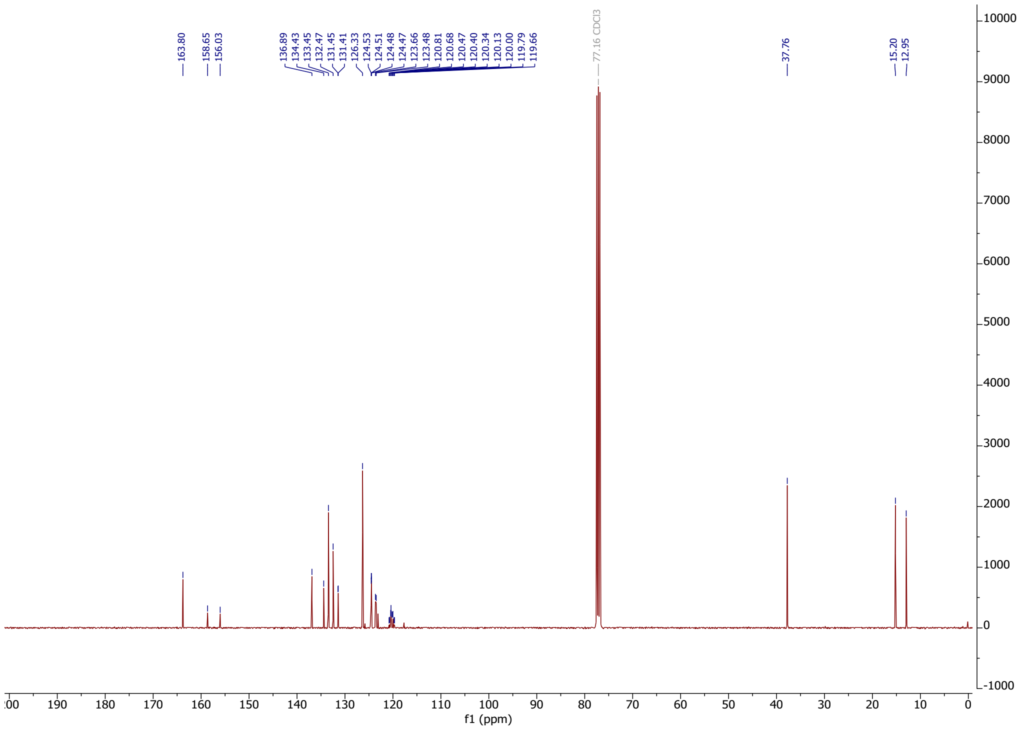

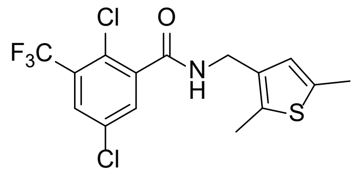
Cl
O
F3C
N
H
S
Cl

**39**

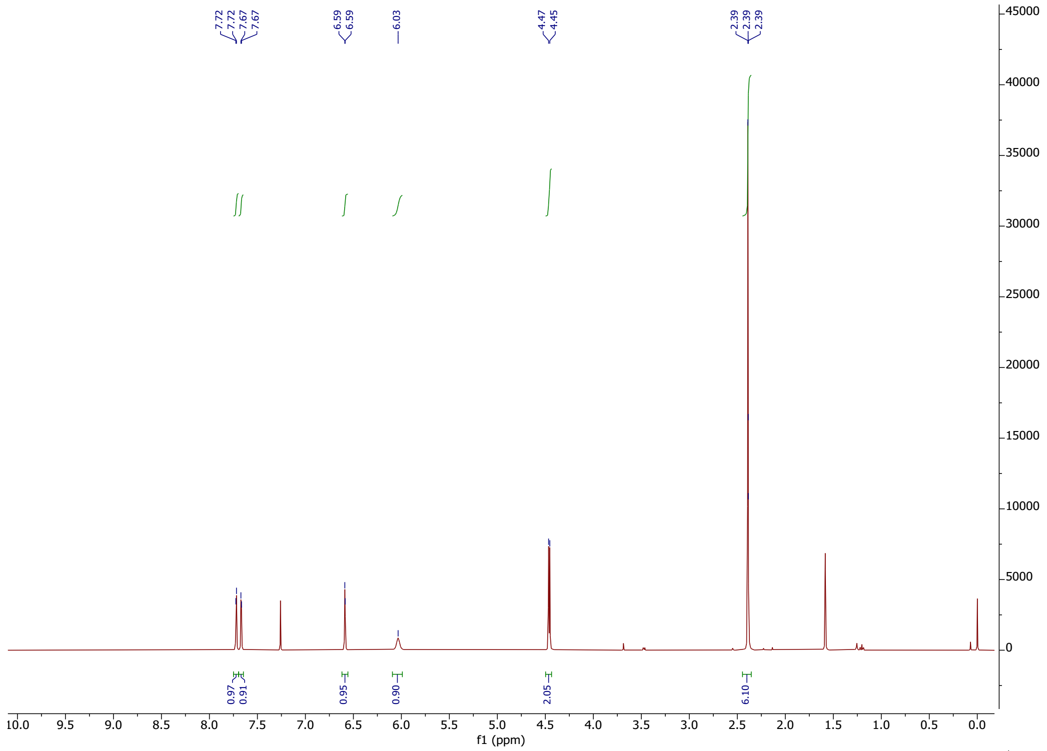
7.72
7.72
7.67
7.67
6.59
6.59
6.03
4.47
4.45
2.39
2.39
2.39
0.97
0.91
0.95
0.90
2.05
6.10
f1 (ppm)

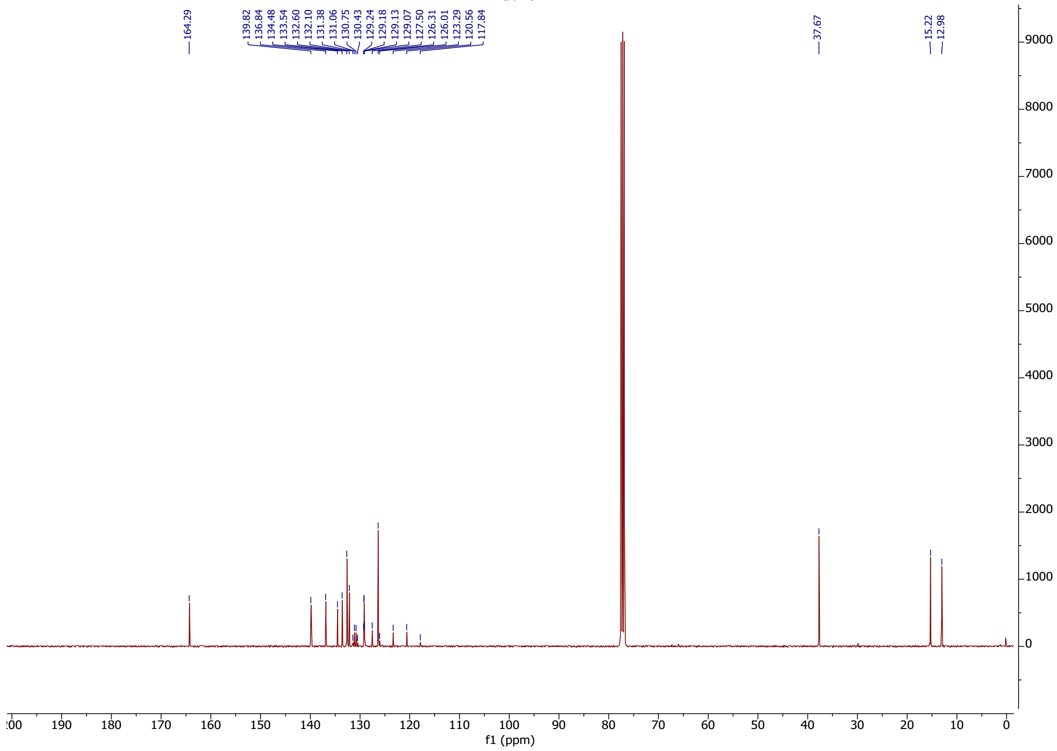
164.29
139.82
136.84
134.48
133.54
132.60
132.10
131.38
131.06
130.75
130.43
129.24
129.18
129.13
129.07
127.50
126.31
126.01
123.29
120.56
117.84
37.67
15.22
12.98
f1 (ppm)

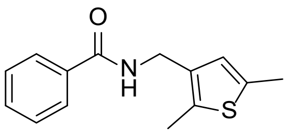

**40**

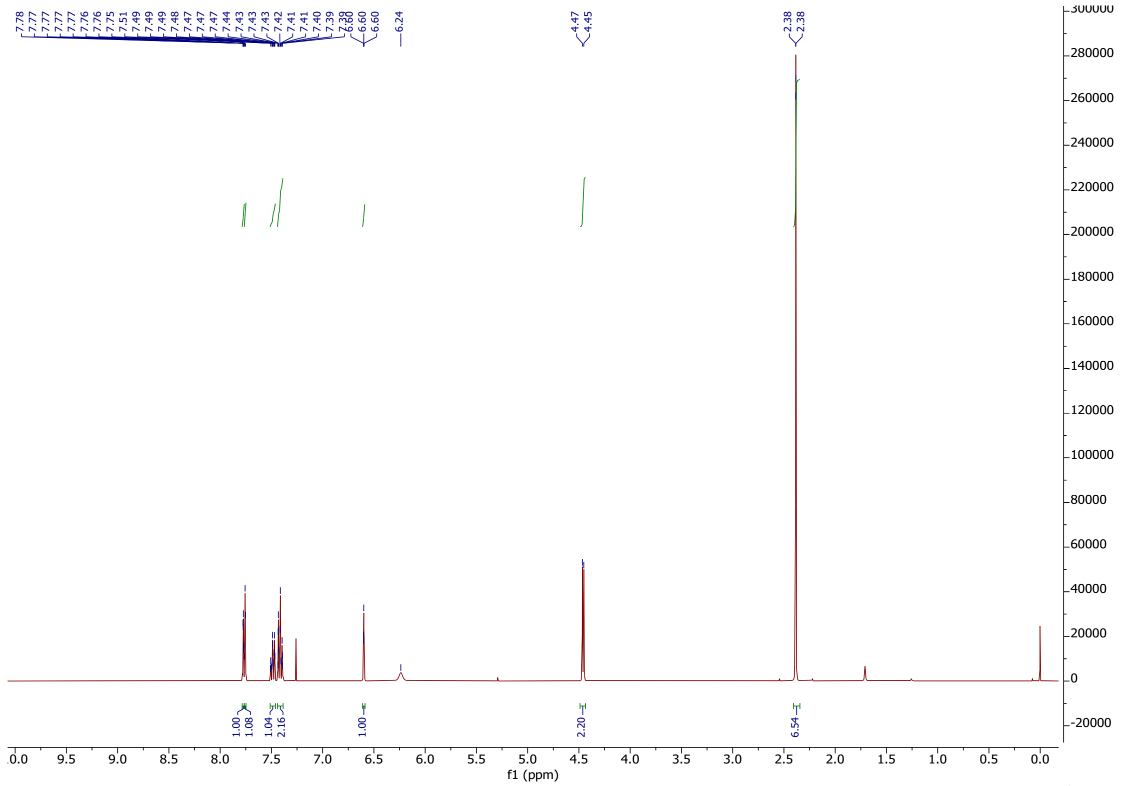

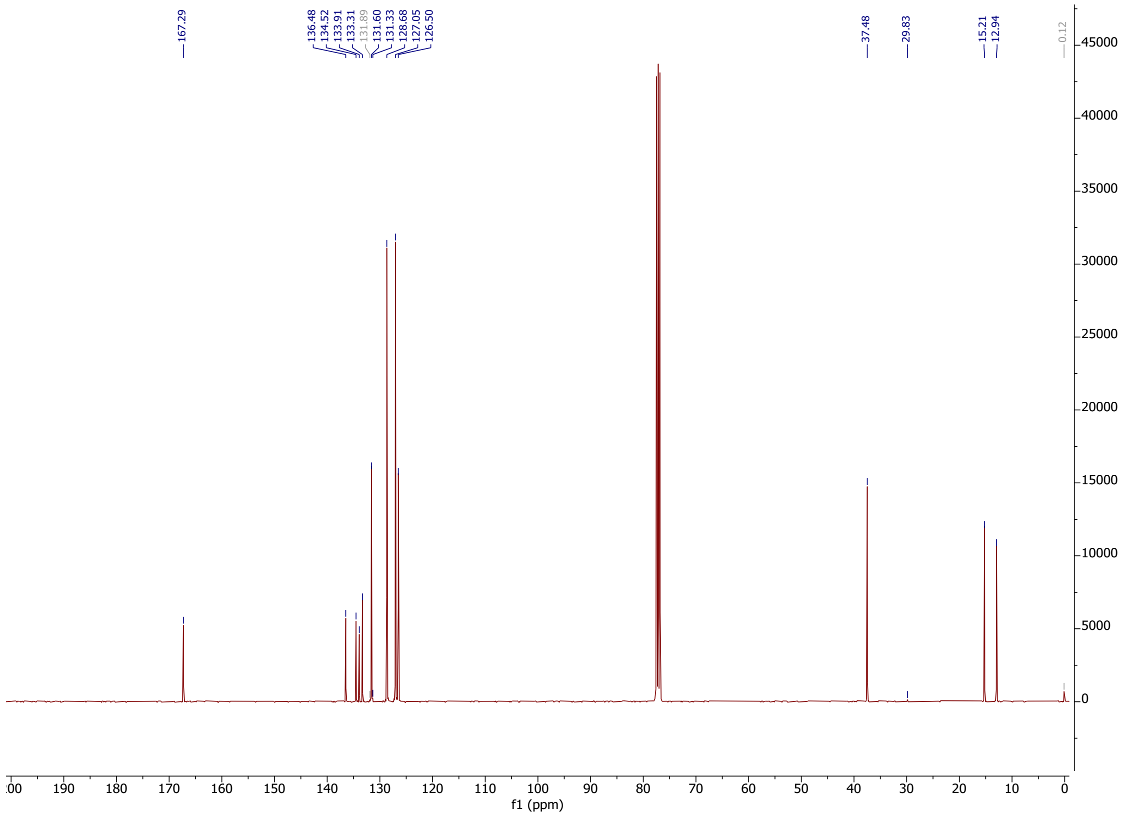

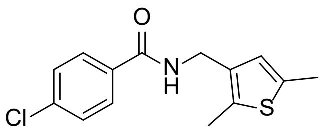

**41**

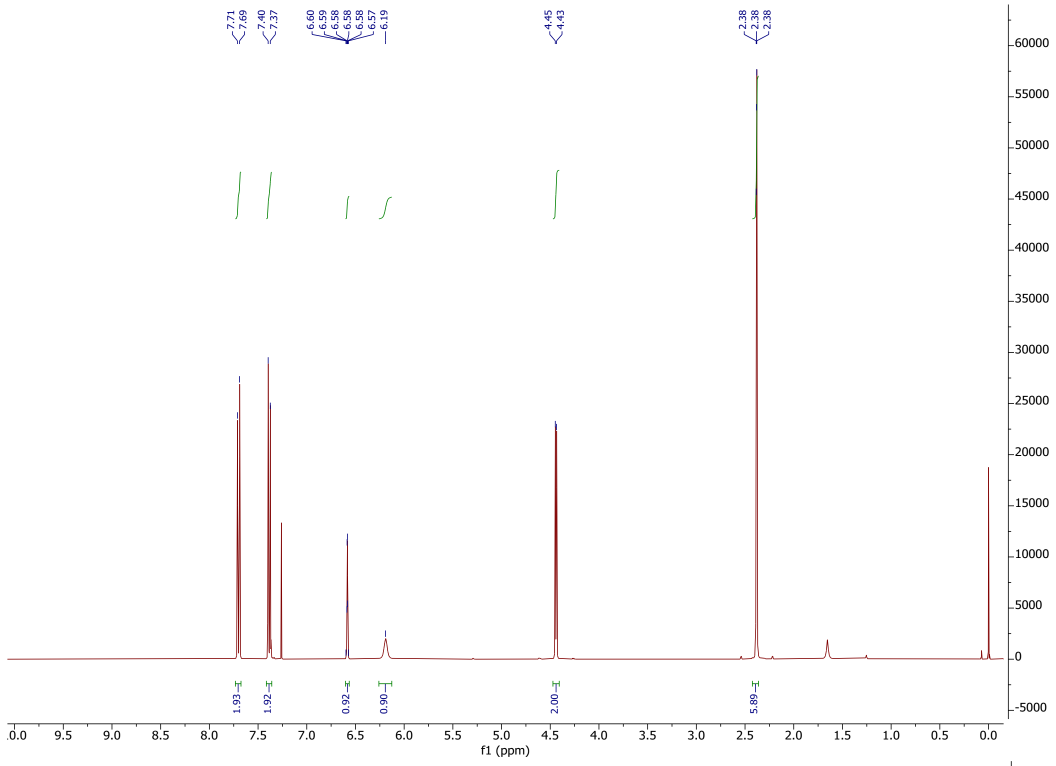


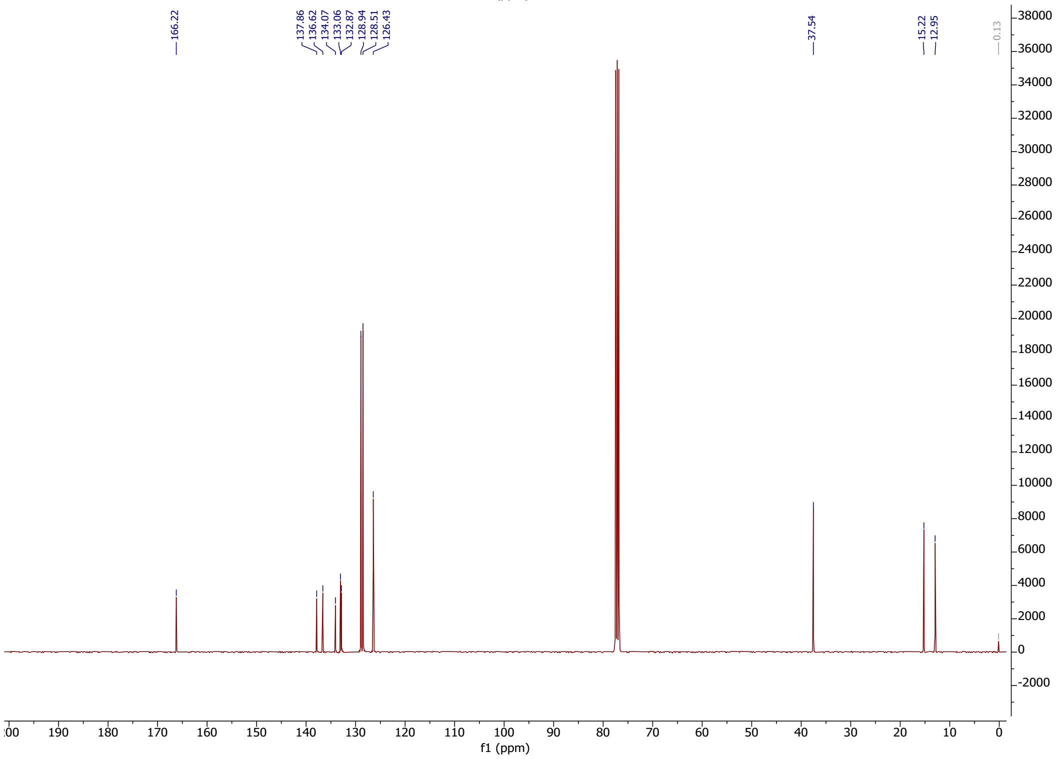

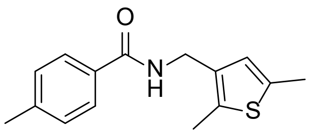

**42**

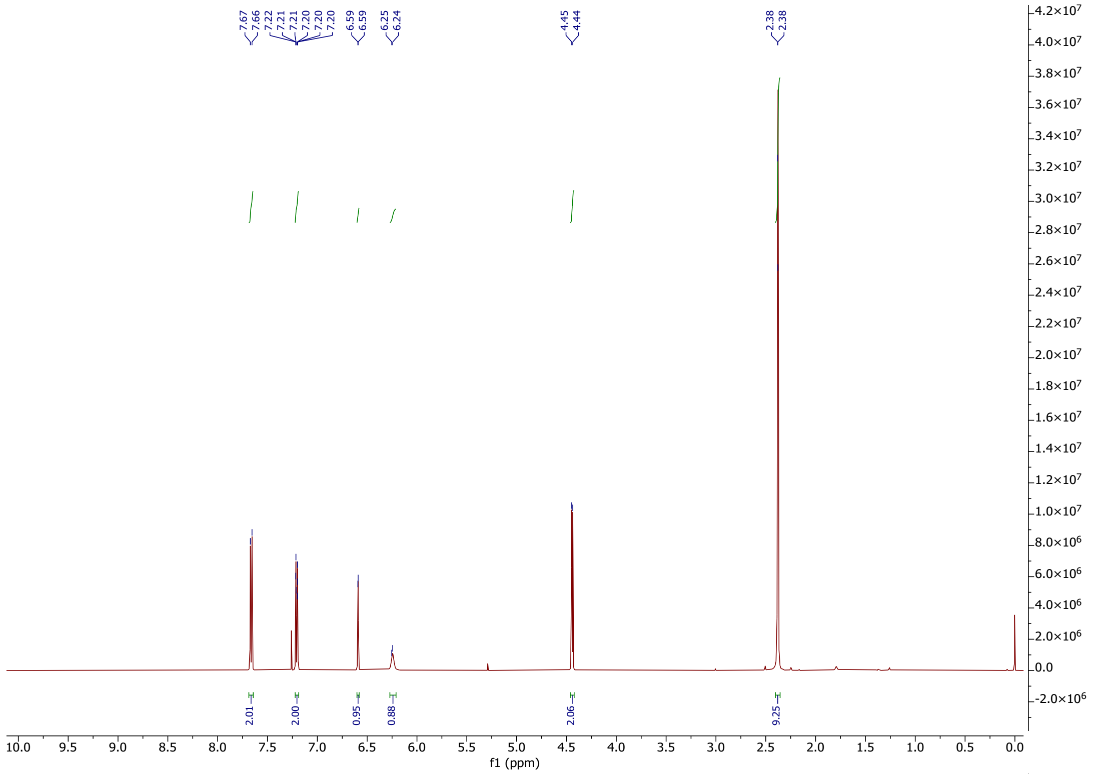

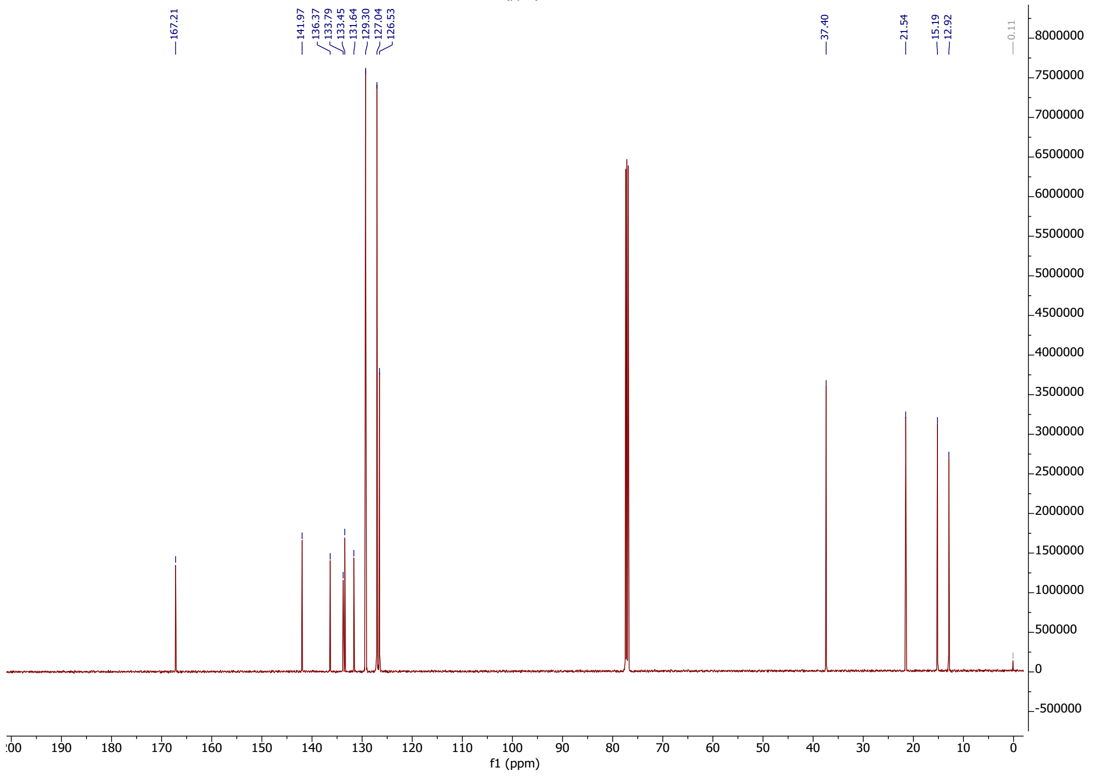

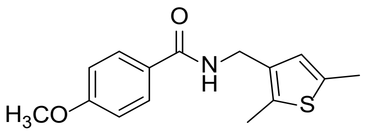
O
N
H
H3CO
S

**43**

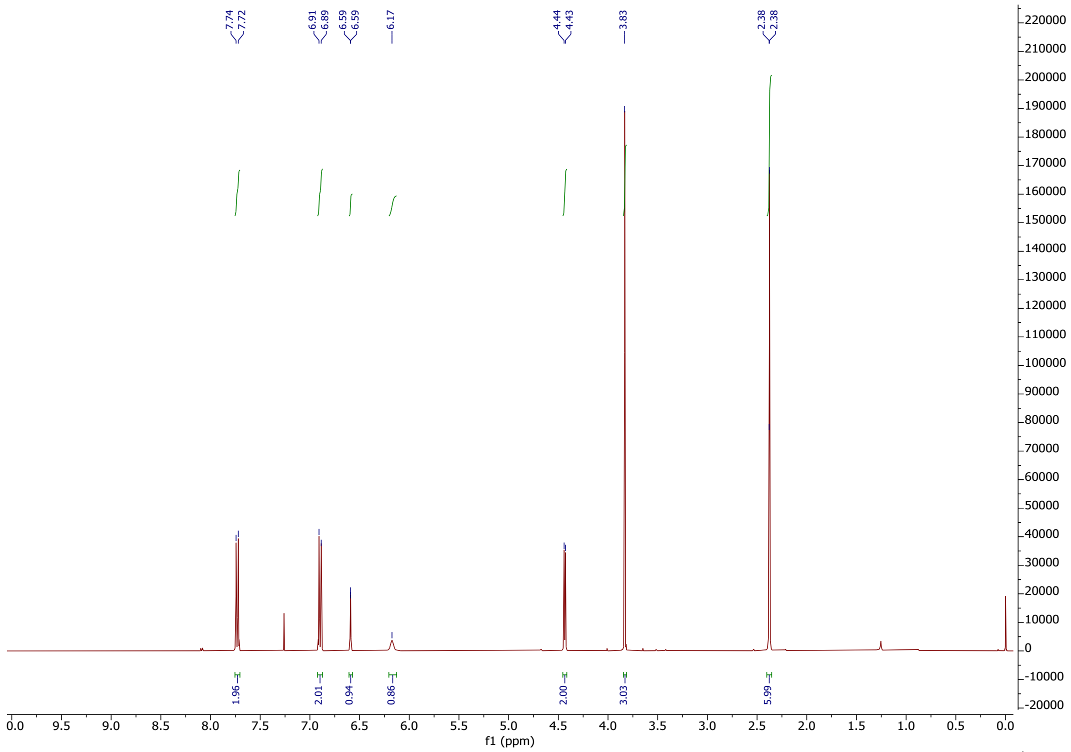
7.74
7.72
6.91
6.89
6.59
6.59
6.17
4.44
4.43
3.83
2.38
2.38
1.96
2.01
0.94
0.86
2.00
3.03
5.99
f1 (ppm)

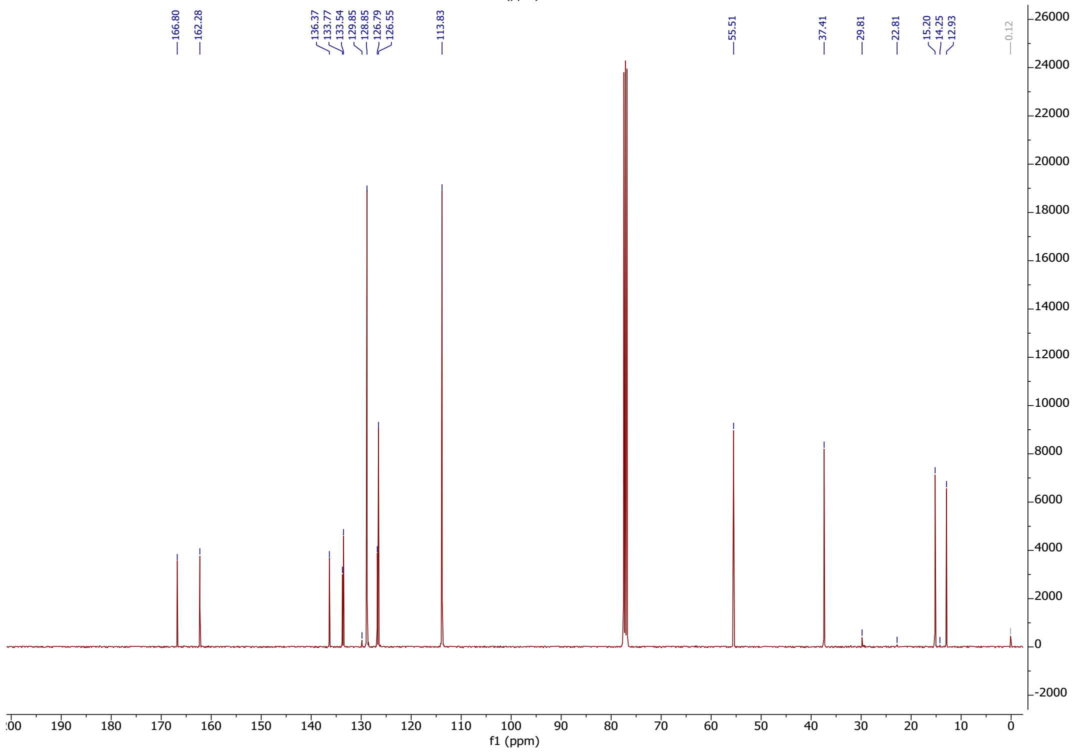
166.80
162.28
136.37
133.77
133.54
129.85
128.85
126.79
126.55
113.83
55.51
37.41
29.81
22.81
15.20
14.25
12.93
0.12
f1 (ppm)

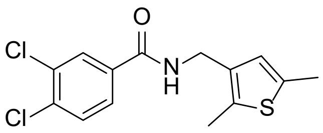
O
Cl
Cl
N
H
S

**44**

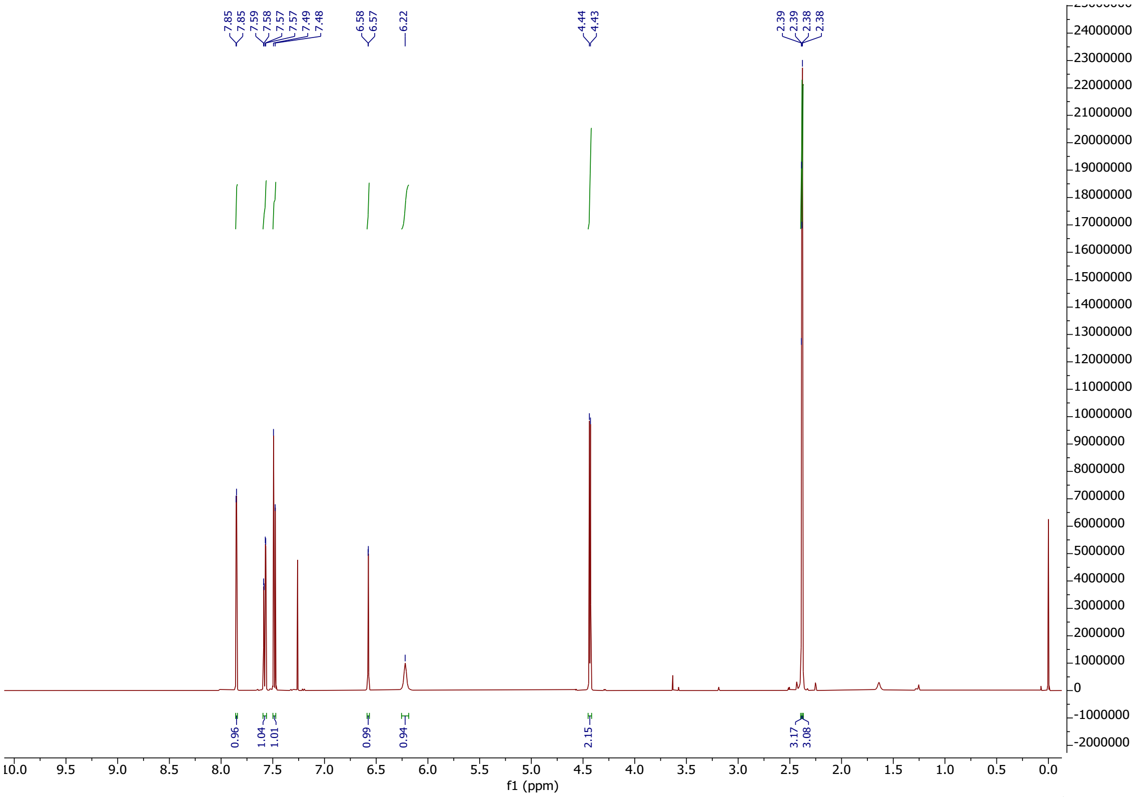
7.85
7.85
7.59
7.58
7.57
7.57
7.49
7.48
6.58
6.57
6.22
4.44
4.43
2.39
2.39
2.38
2.38
0.96
1.04
1.01
0.99
0.94
2.15
3.17
3.08
f1 (ppm)

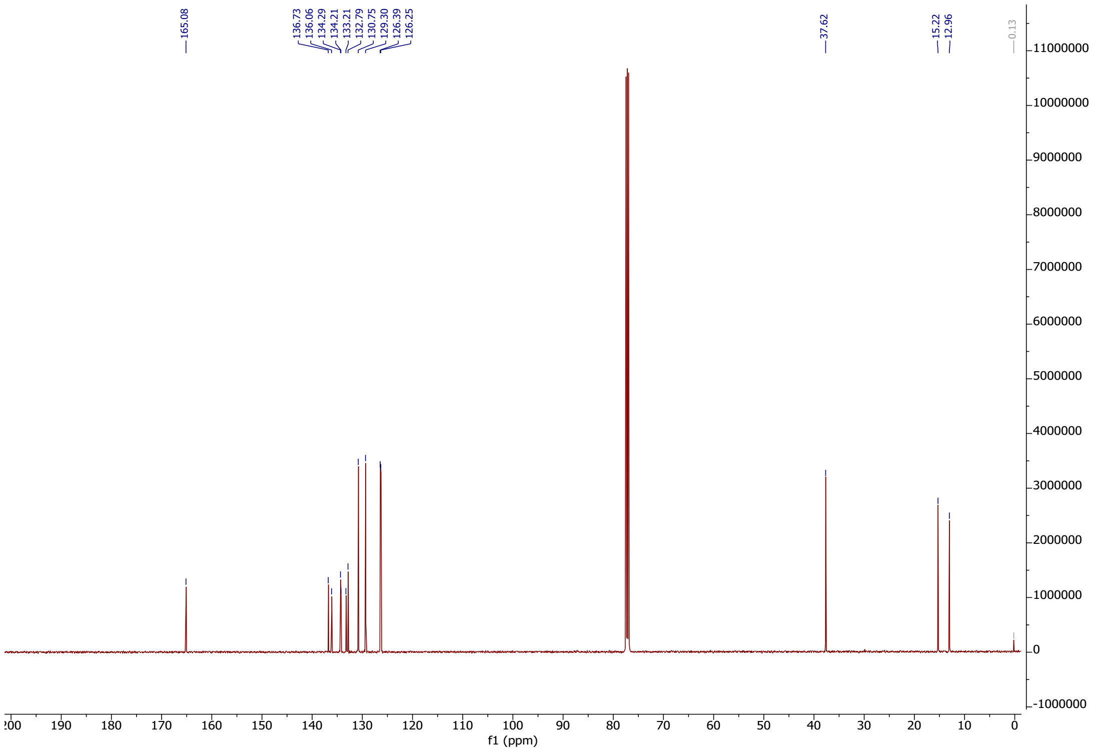
165.08
136.73
136.06
134.29
134.21
133.21
132.79
130.75
129.30
126.39
126.25
37.62
15.22
12.96
0.13
f1 (ppm)

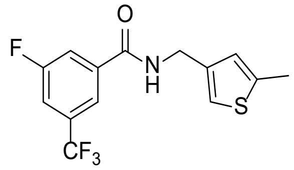

**45**

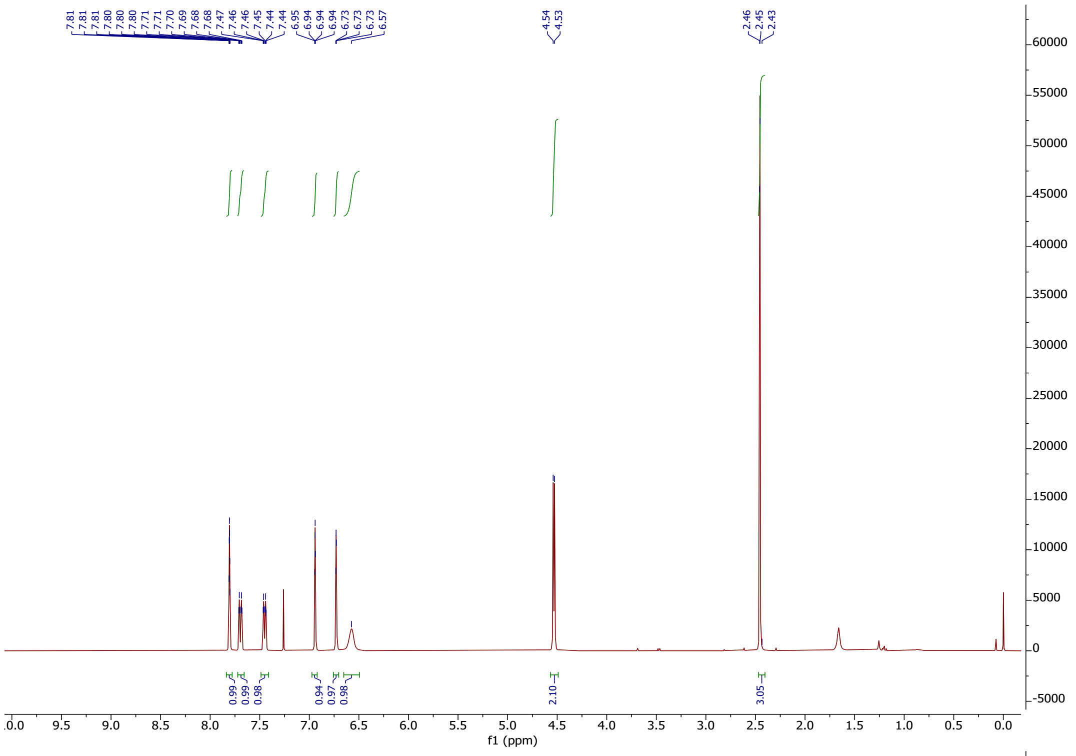


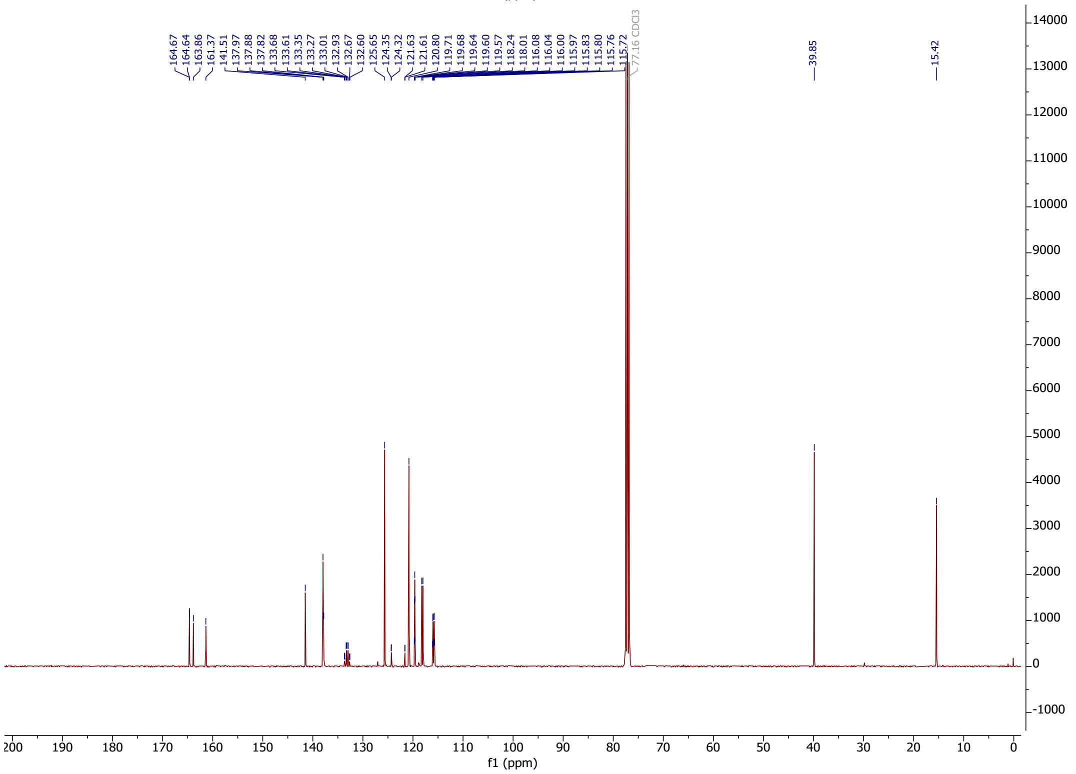

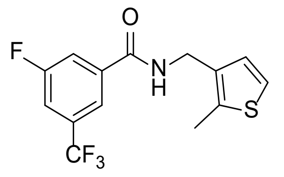

**46**

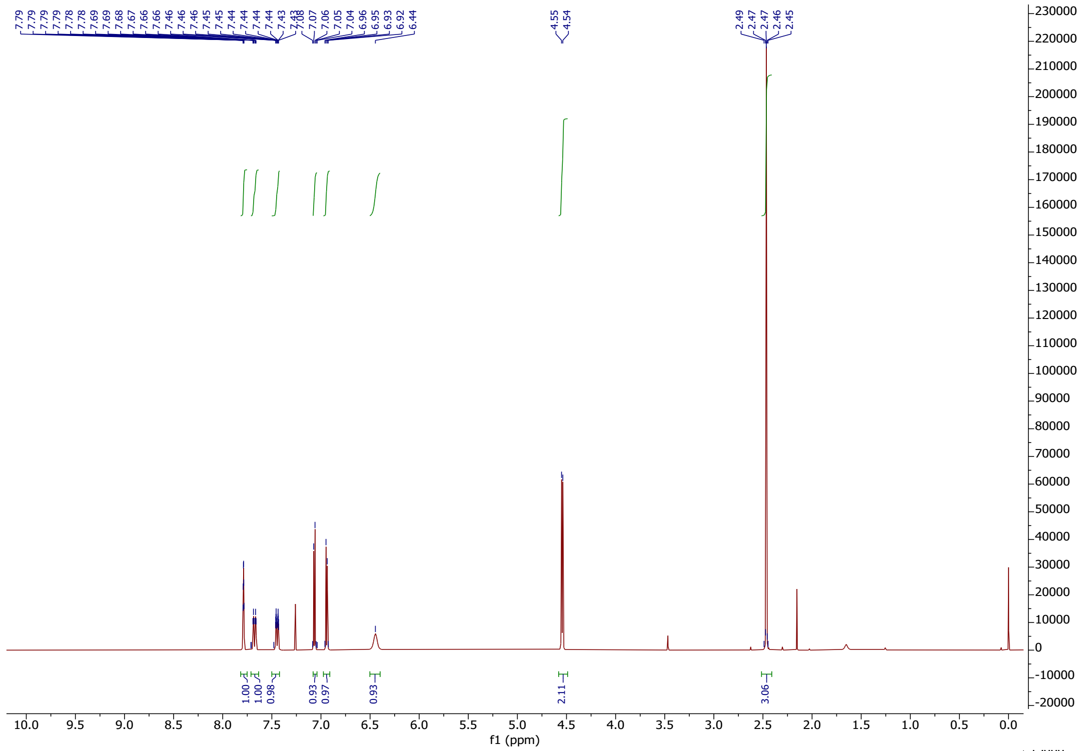

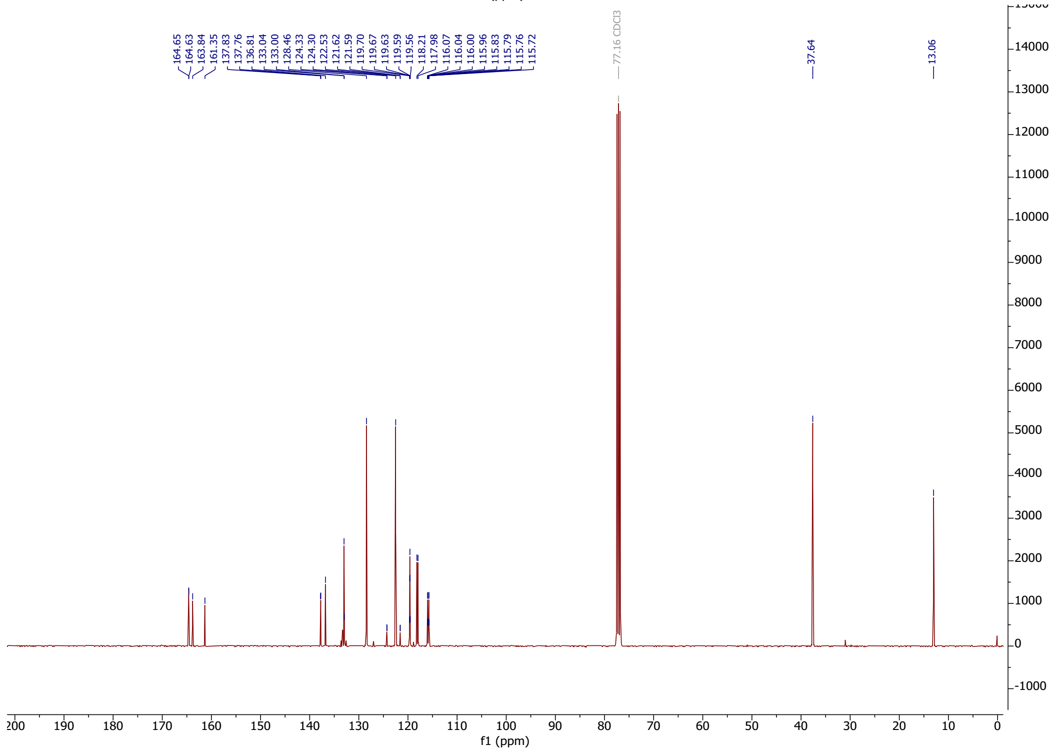

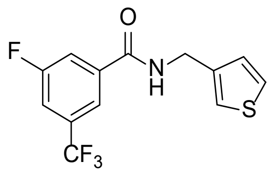


**47**

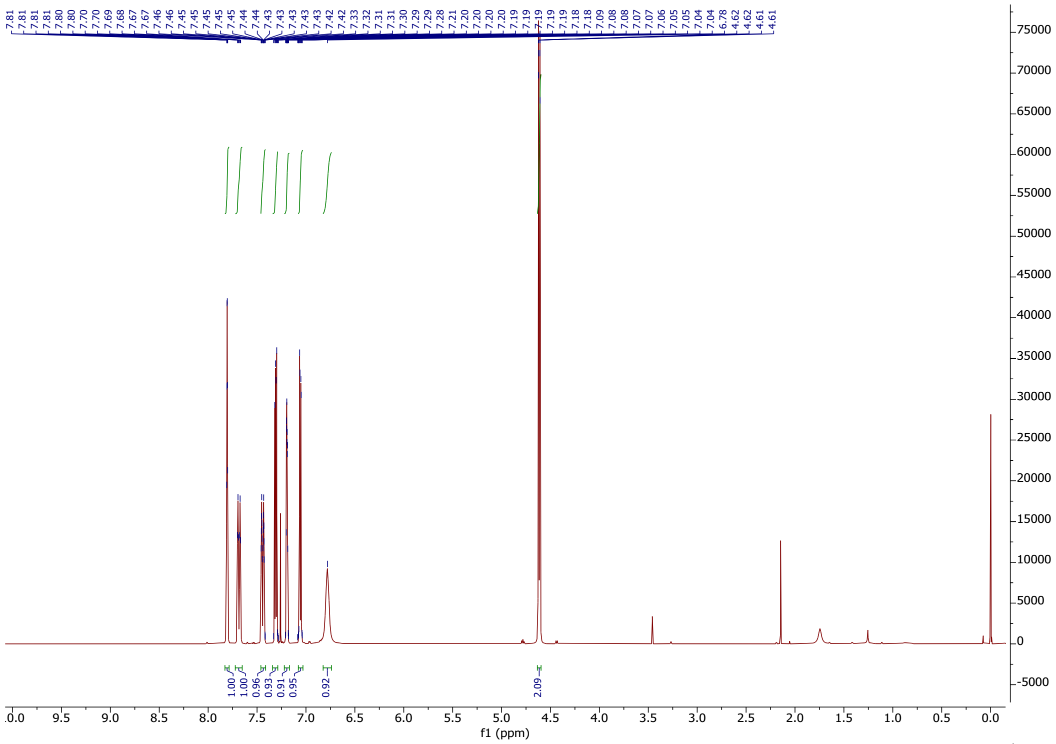


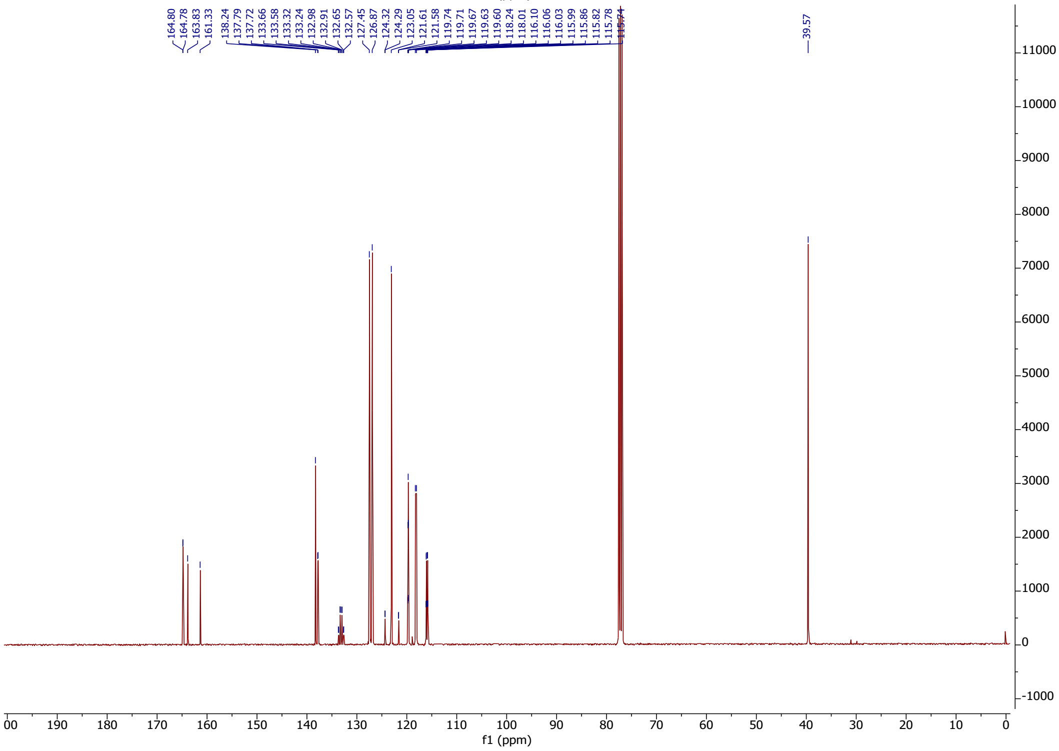

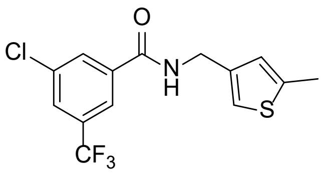

**48**

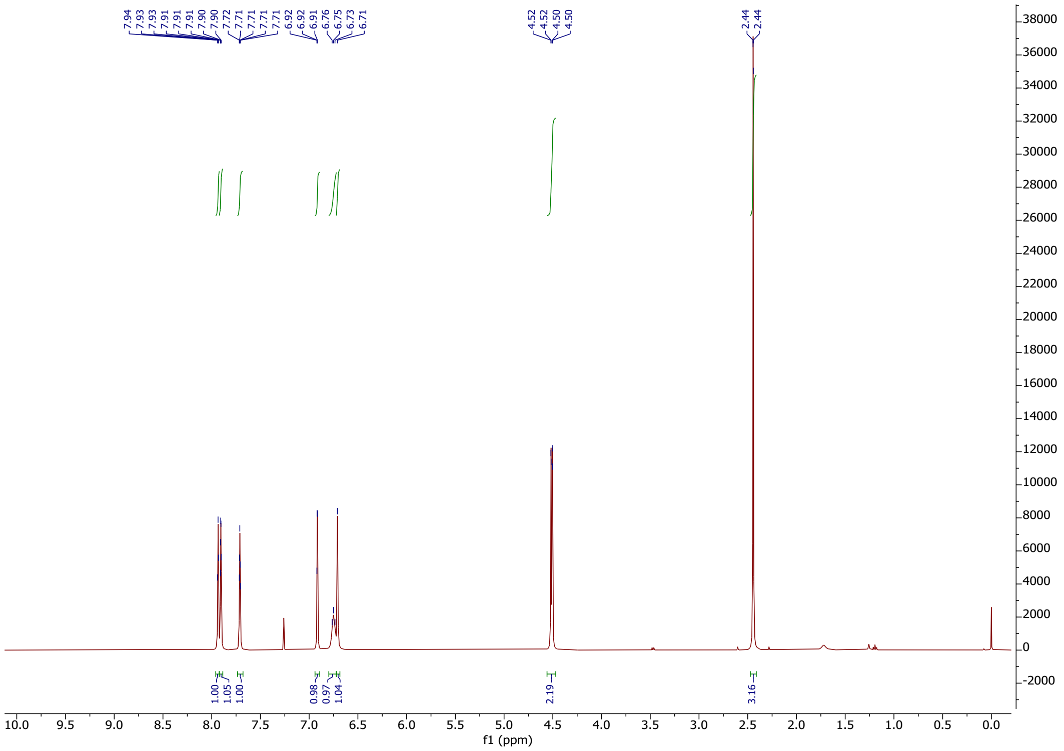

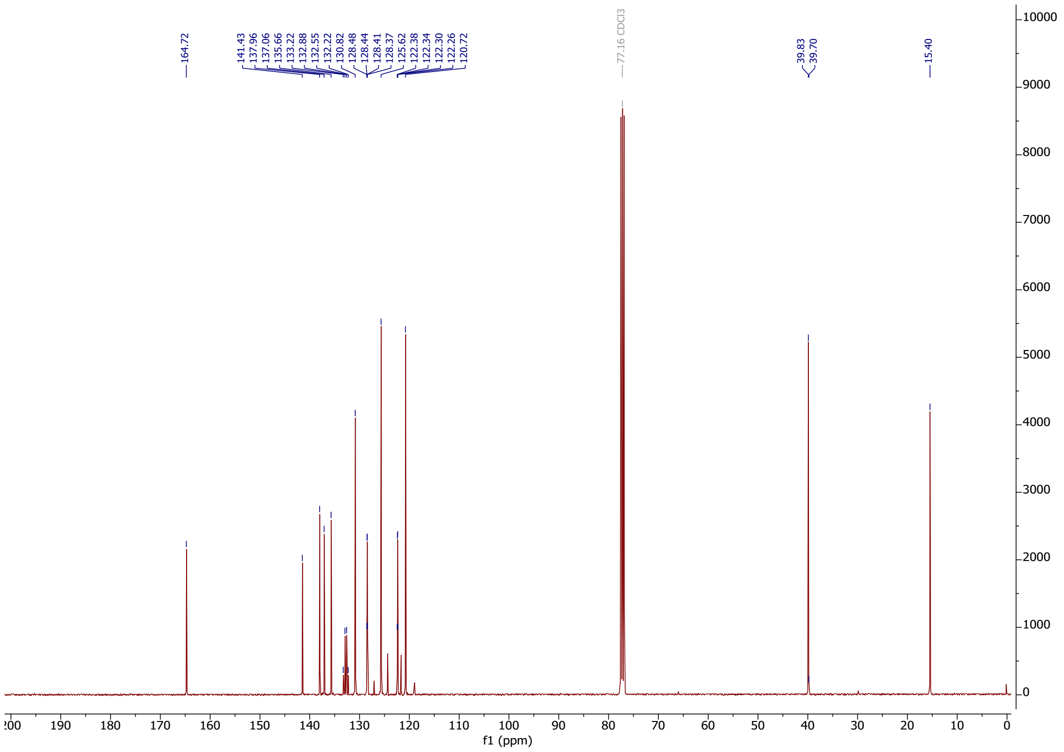

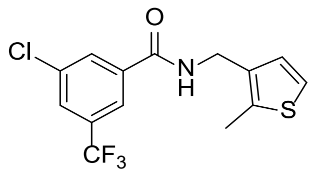

**49**

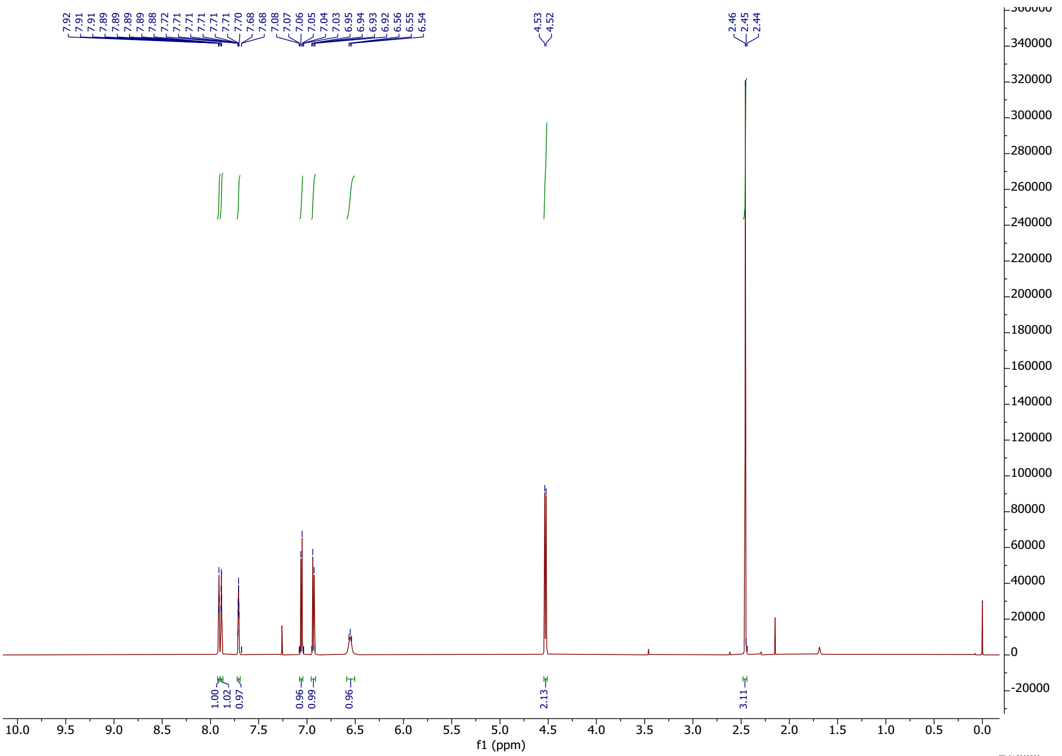

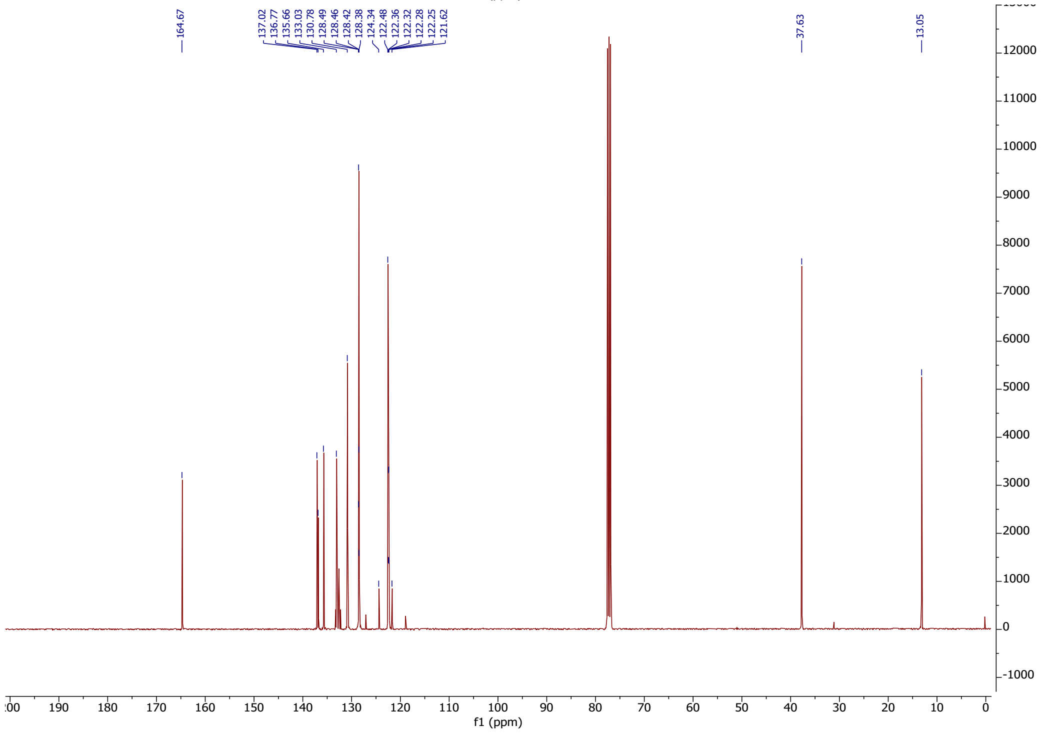

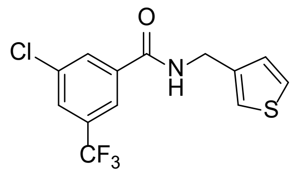

**50**

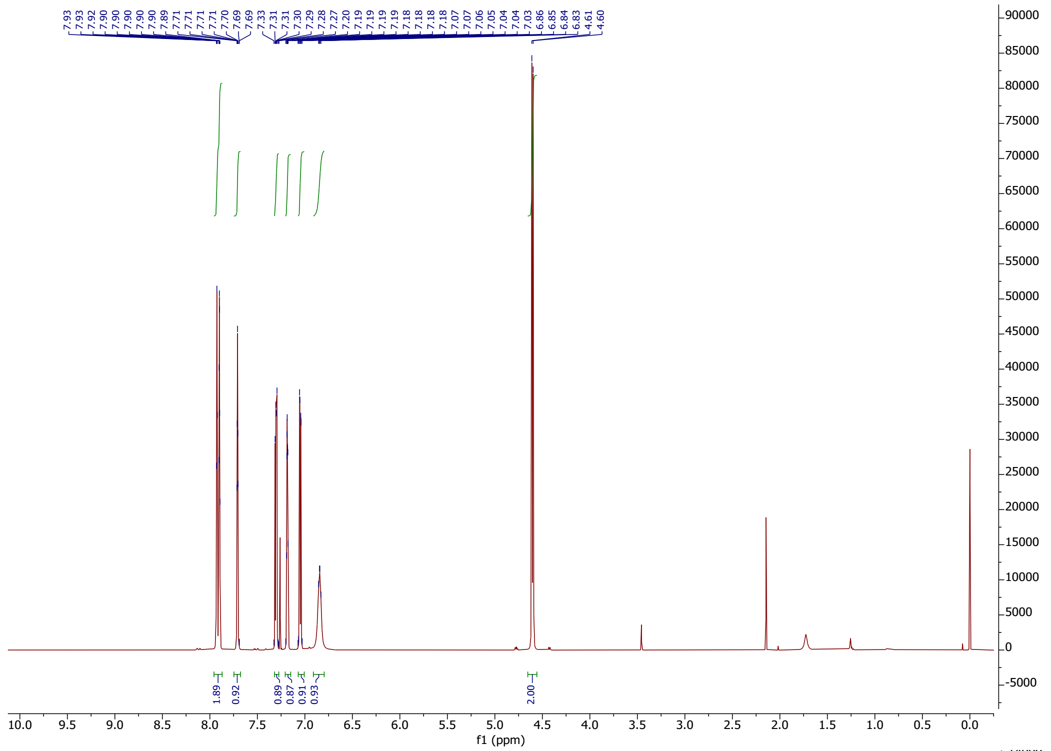

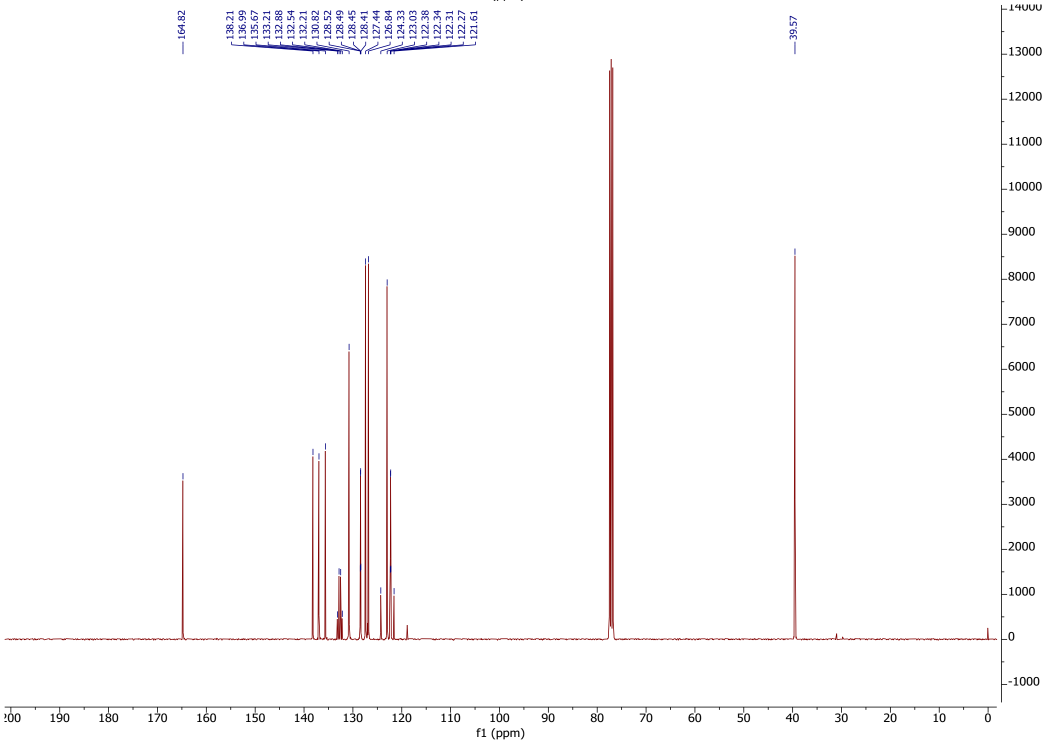

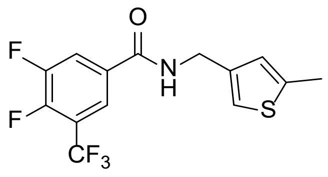

**51**

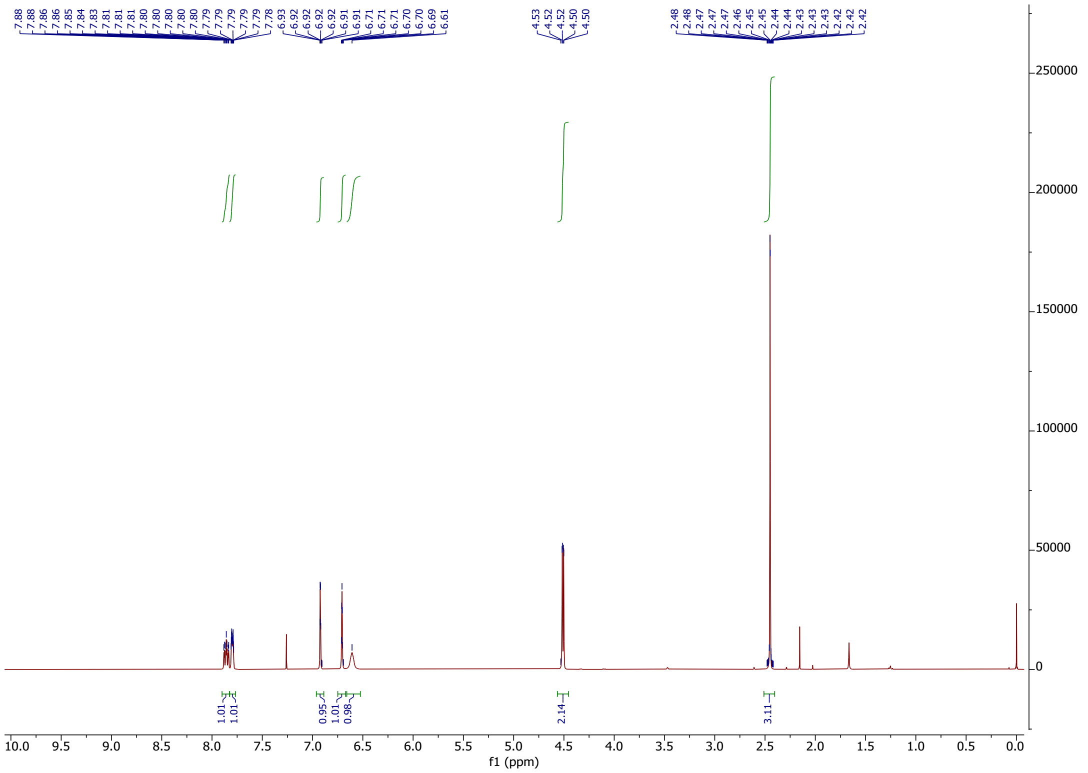

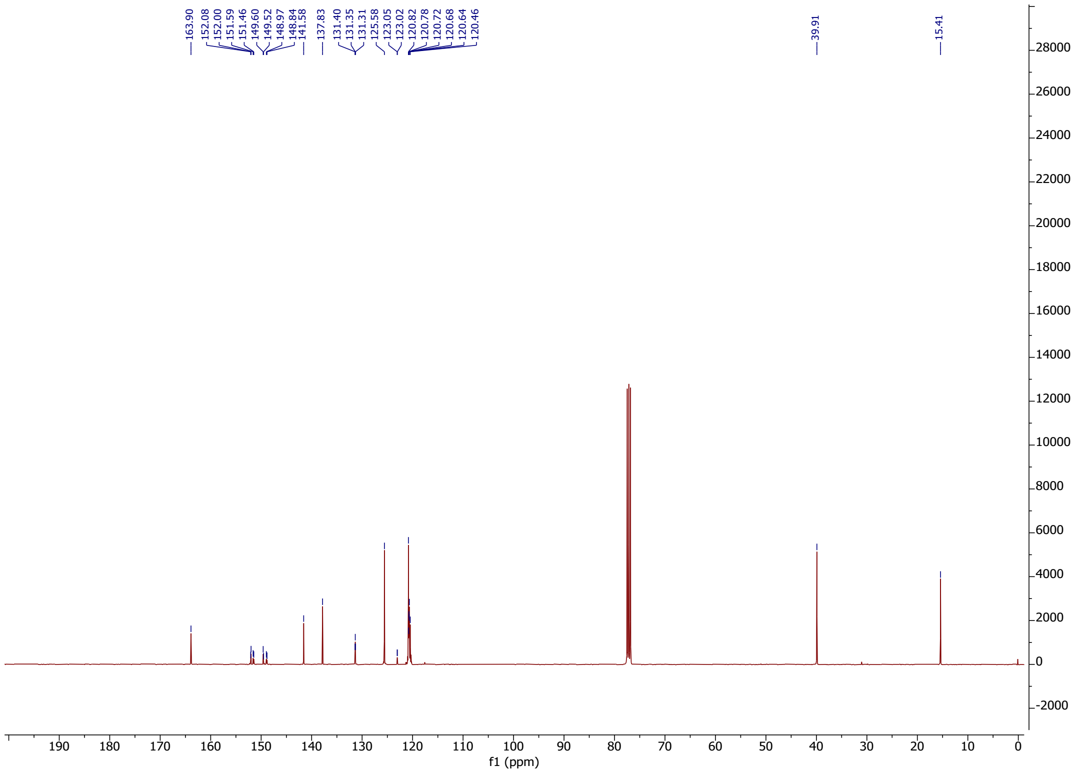

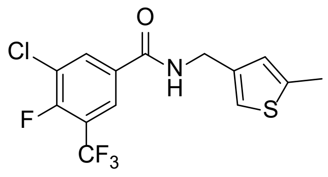

O
Cl
N
H
S
F
CF3


**52**

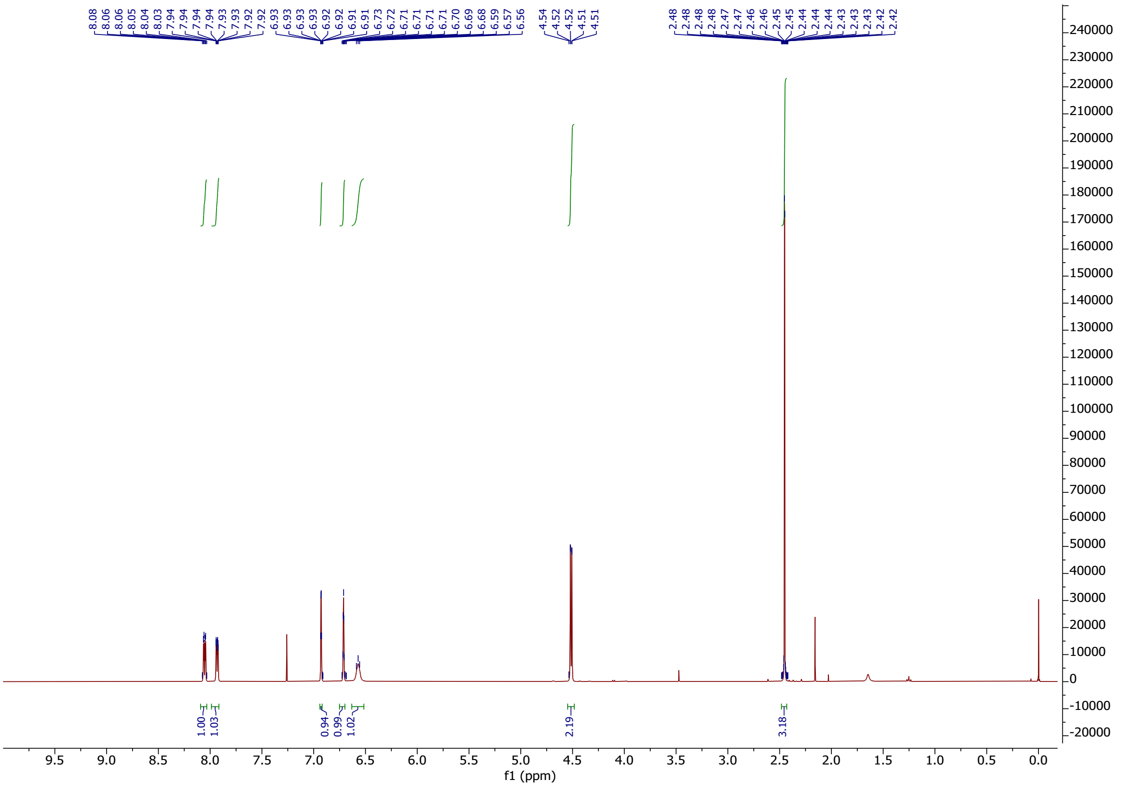

8.08 8.06 8.06 8.05 8.04 8.03 7.94 7.94 7.94 7.94 7.93 7.93 7.92 7.92 6.93 6.93 6.93 6.93 6.92 6.92 6.91 6.91 6.73 6.72 6.71 6.71 6.71 6.71 6.70 6.69 6.68 6.59 6.57 6.56 4.54 4.52 4.52 4.51 4.51 2.48 2.48 2.48 2.48 2.47 2.46 2.46 2.45 2.45 2.44 2.44 2.43 2.43 2.43 2.42 2.42
1.00 1.03 0.94 0.99 1.02 2.19 3.18
f1 (ppm)


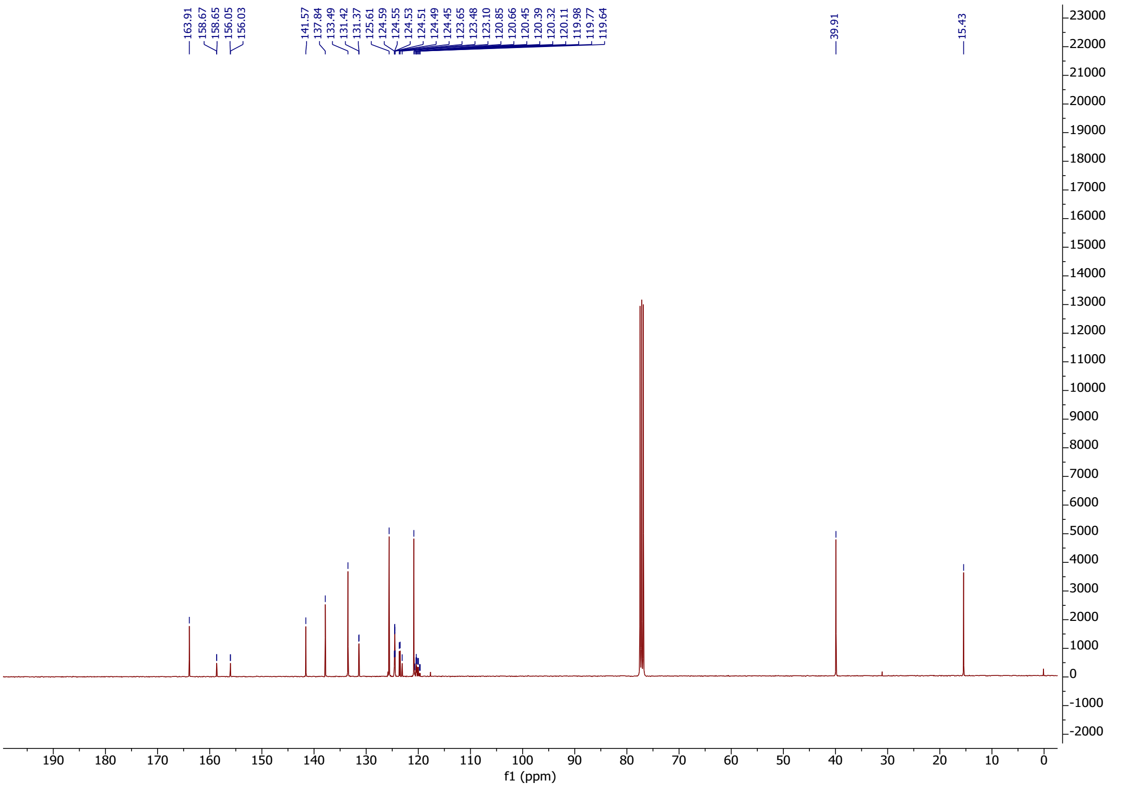

163.91 158.67 158.65 156.05 156.03 141.57 137.84 133.49 131.42 131.37 125.61 124.59 124.55 124.53 124.51 124.49 124.45 123.65 123.48 123.10 120.85 120.66 120.45 120.39 120.32 120.11 119.98 119.77 119.64 39.91 15.43
f1 (ppm)

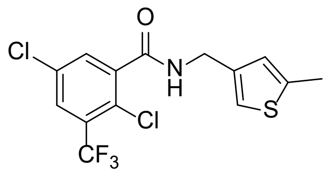

**53**

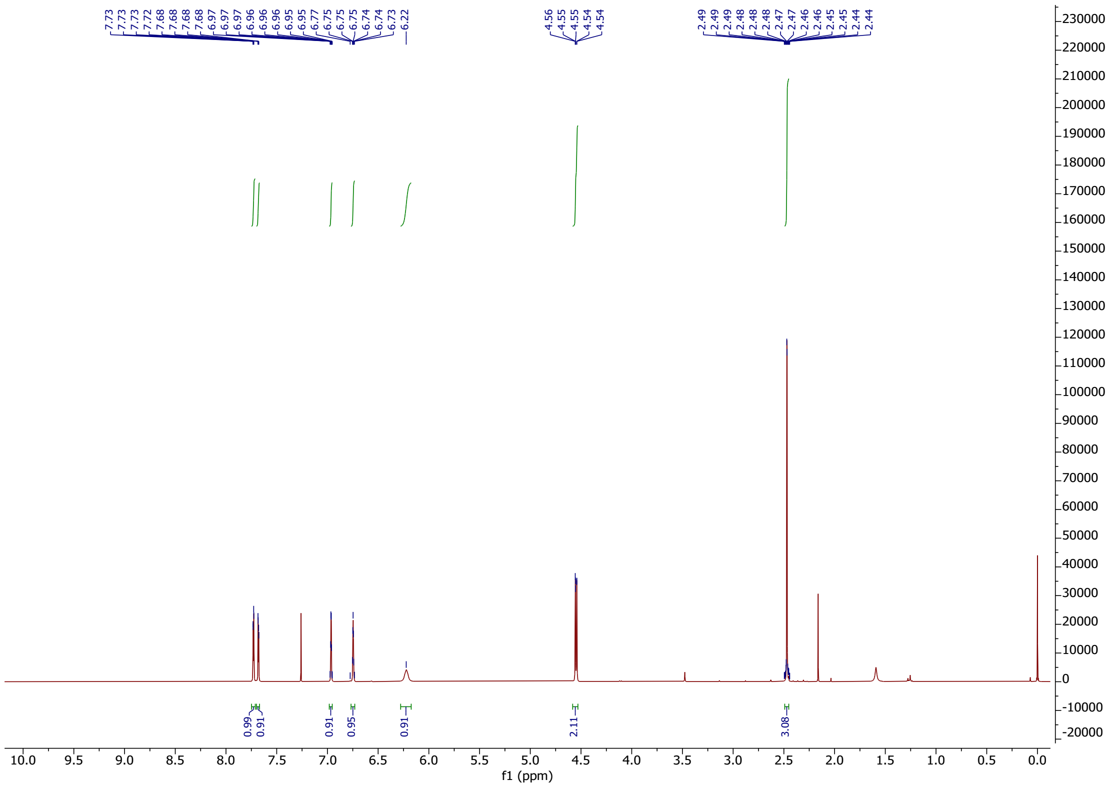

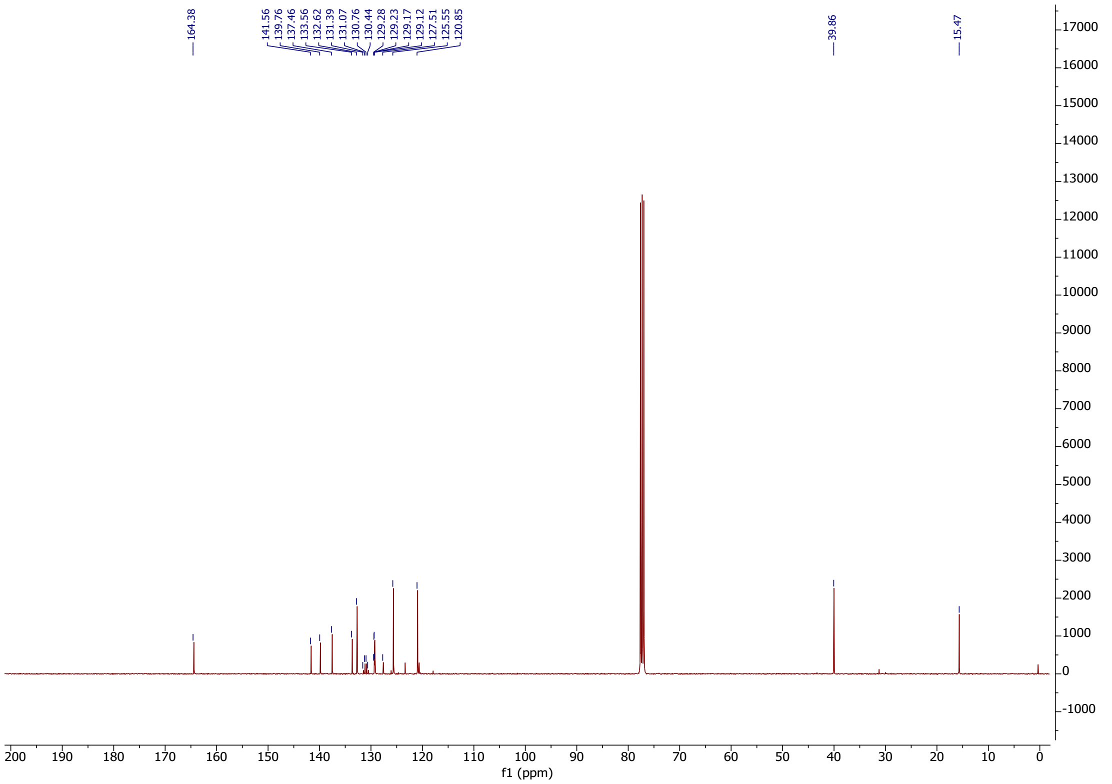

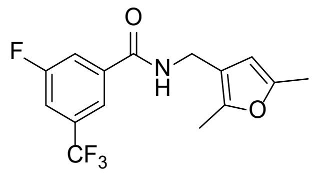

**54**

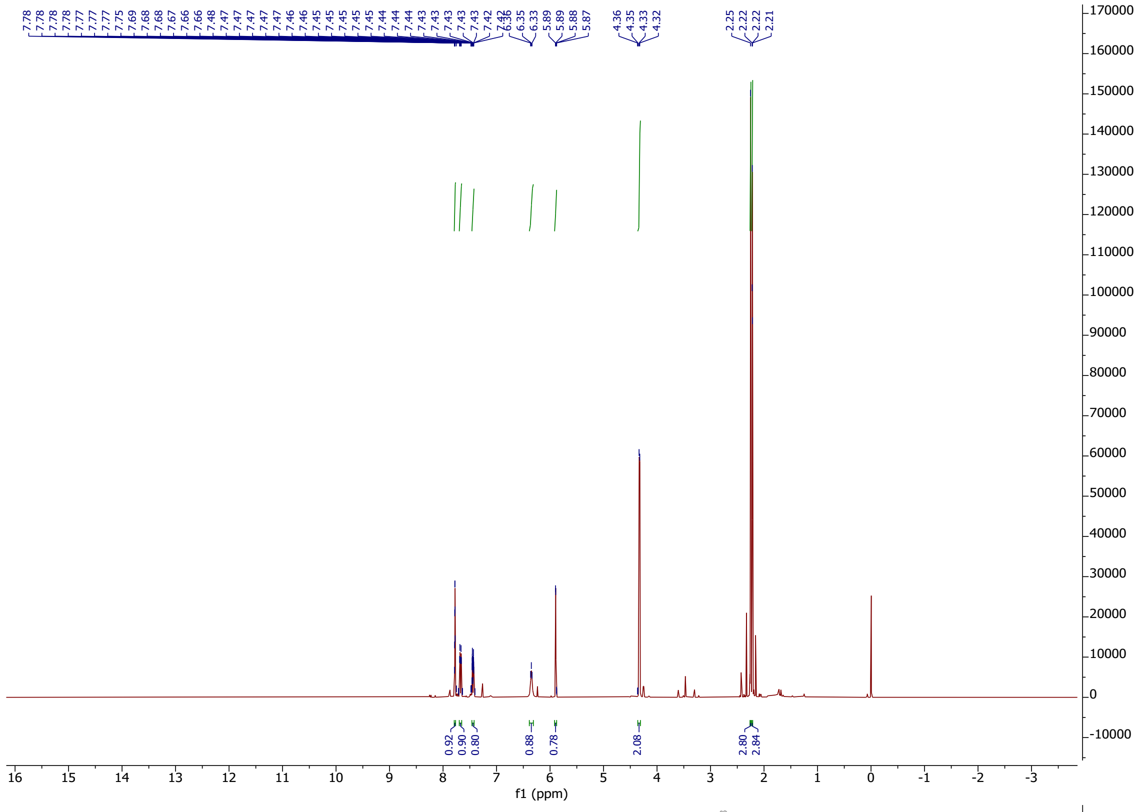

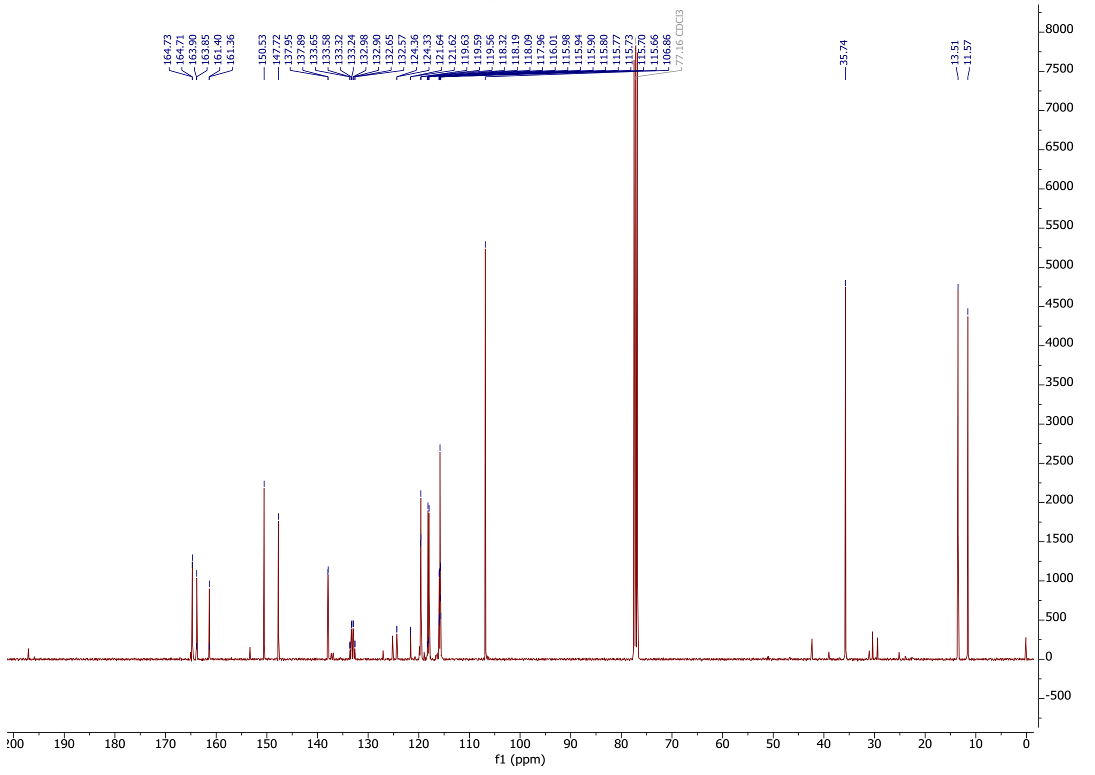

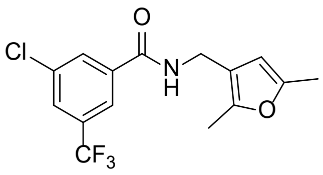

**55**

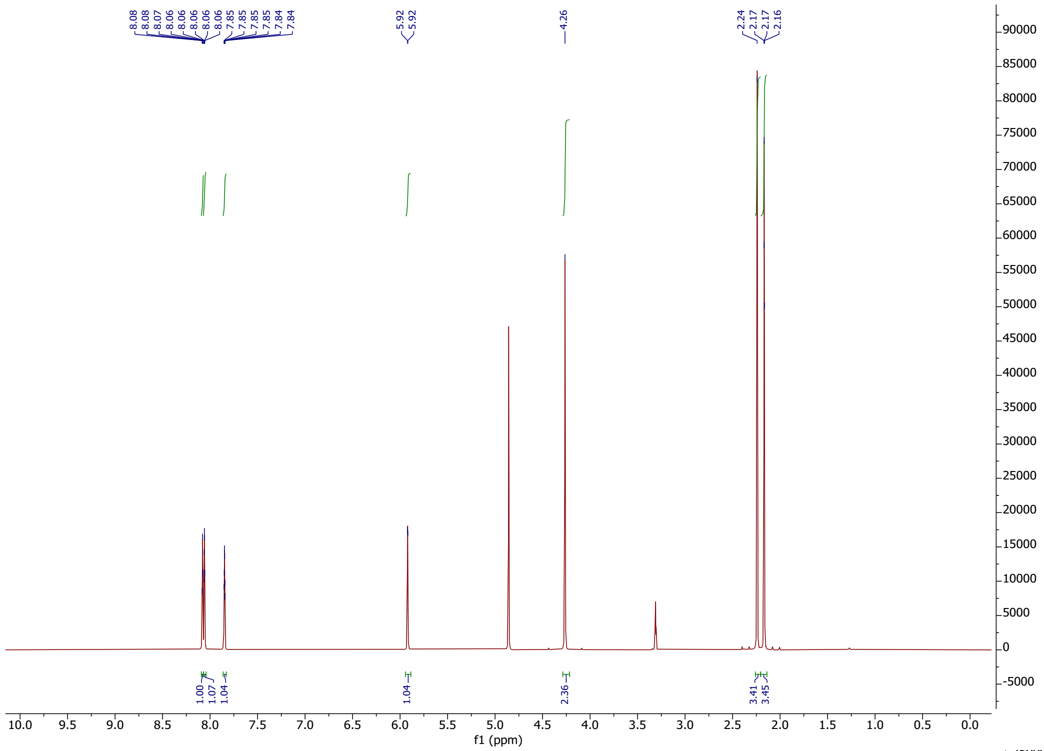

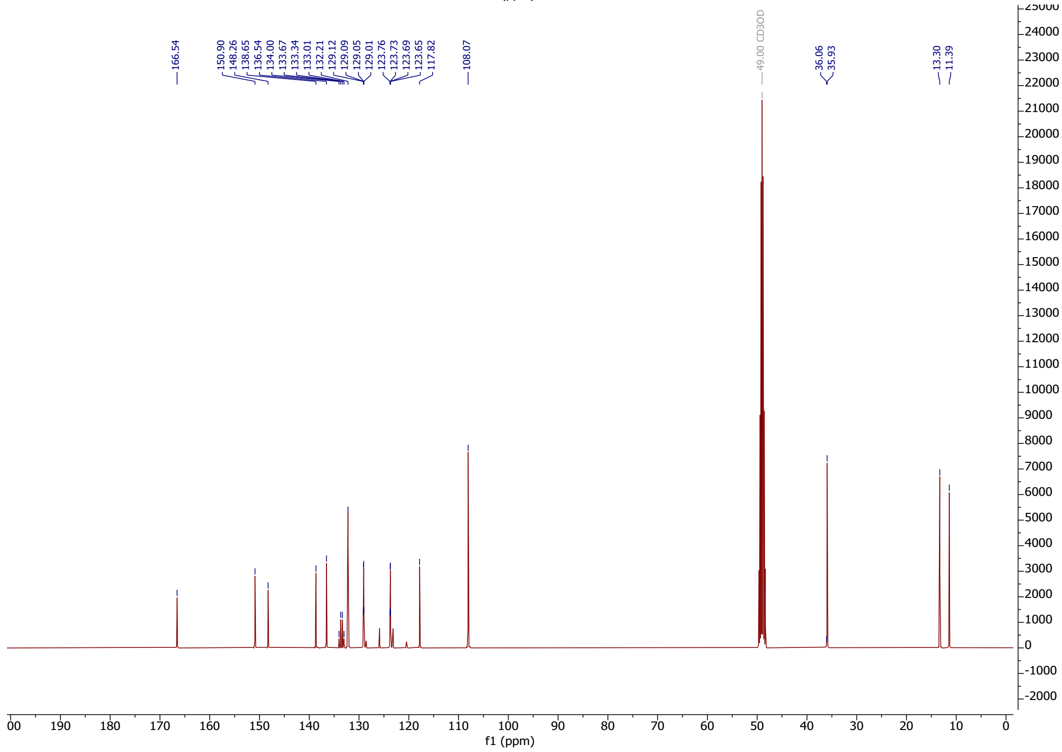

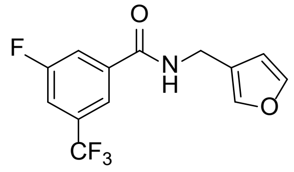

**56**

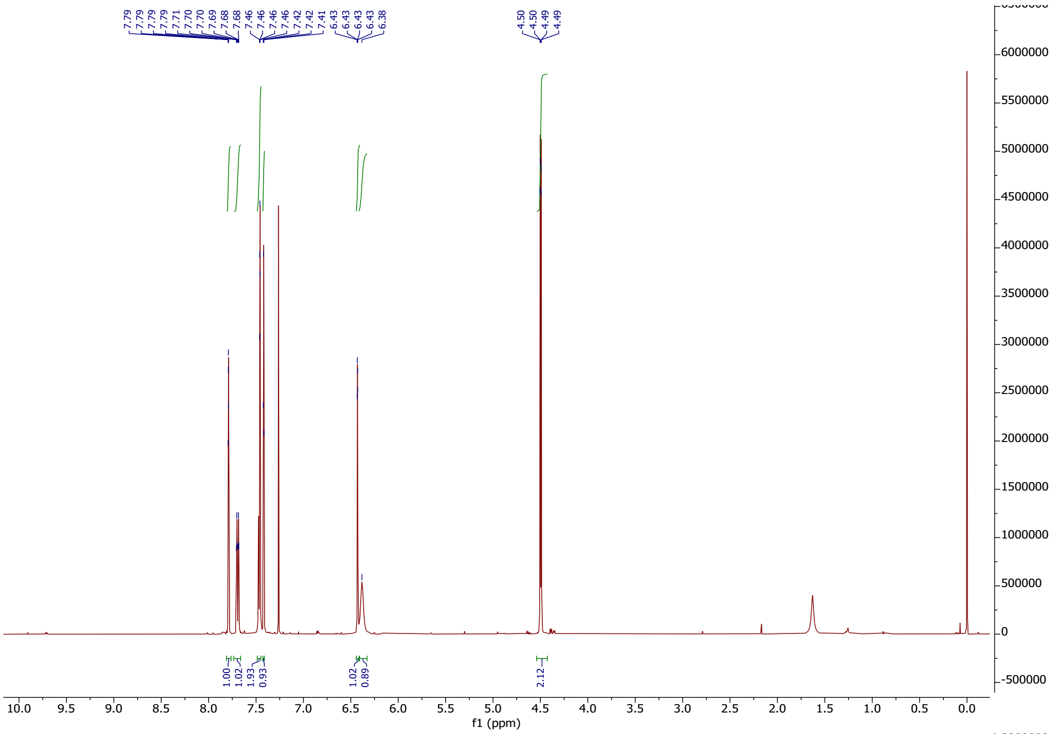

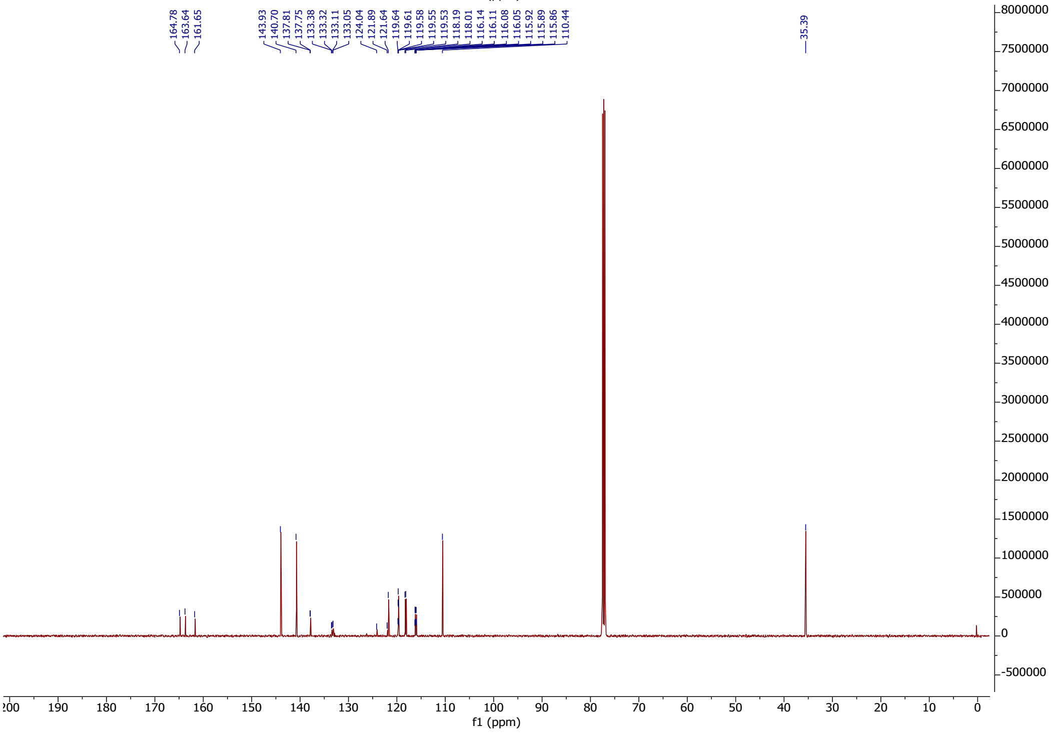

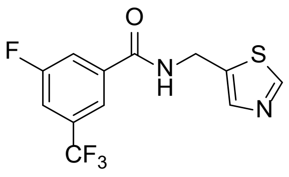

**57**

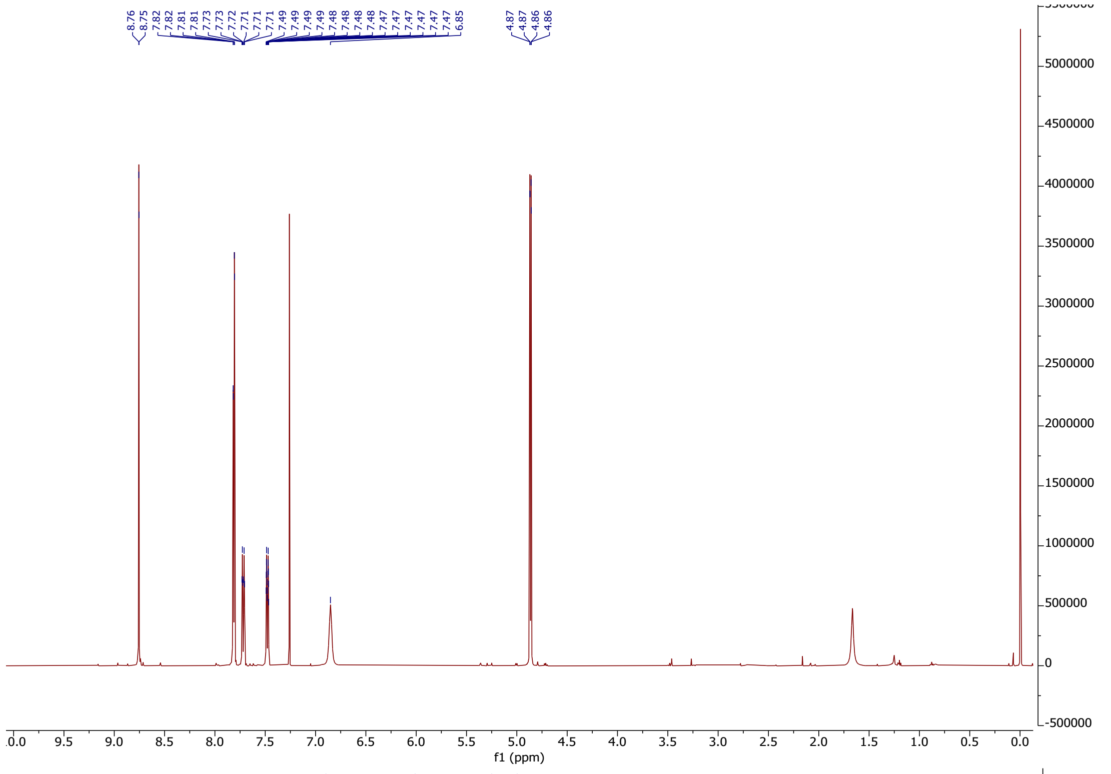

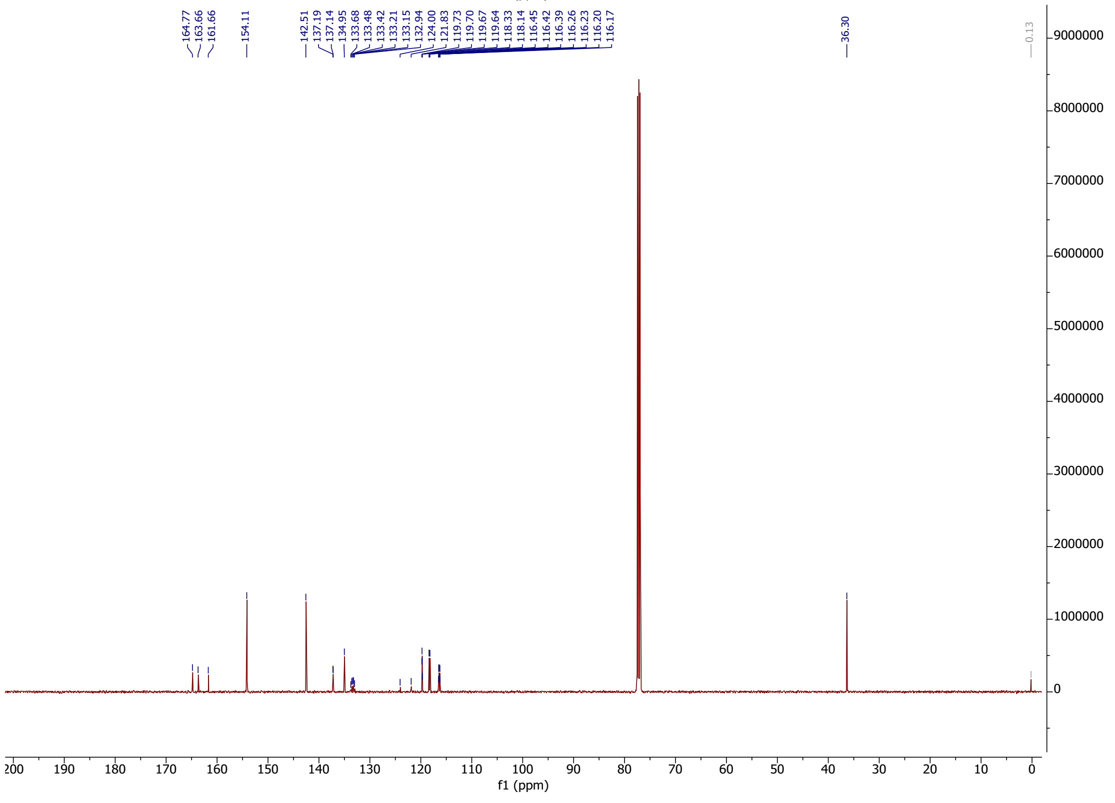

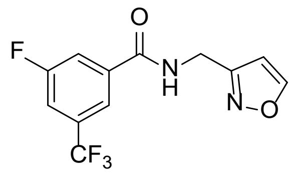

**58**

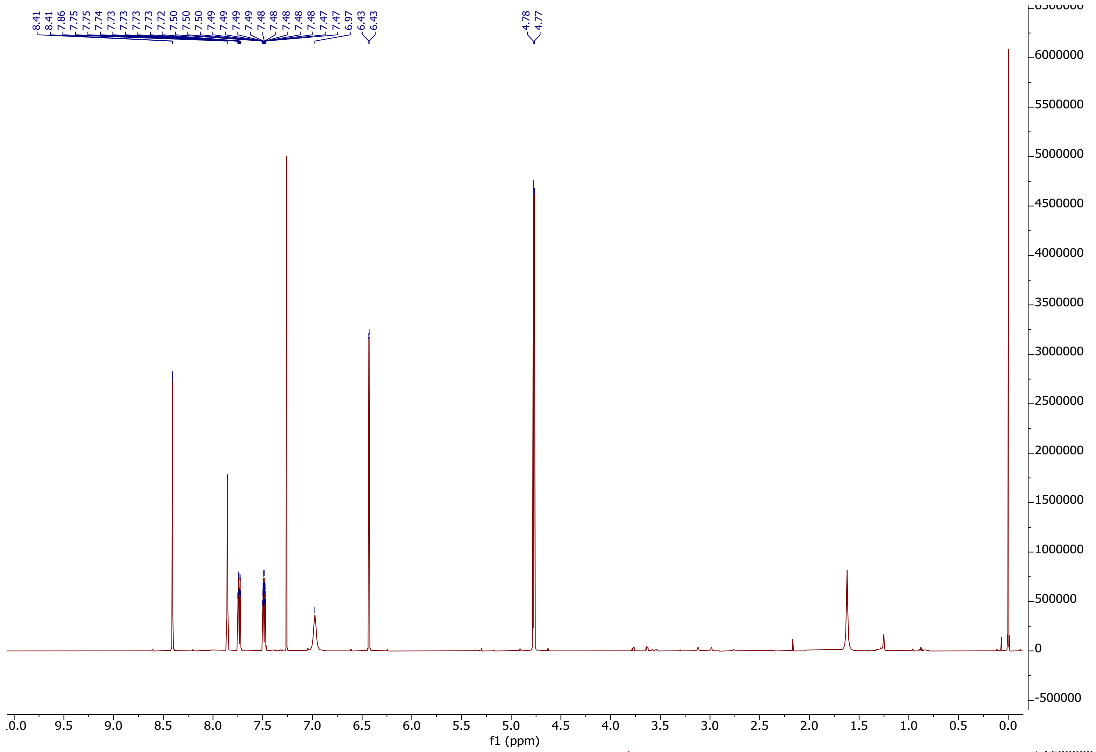

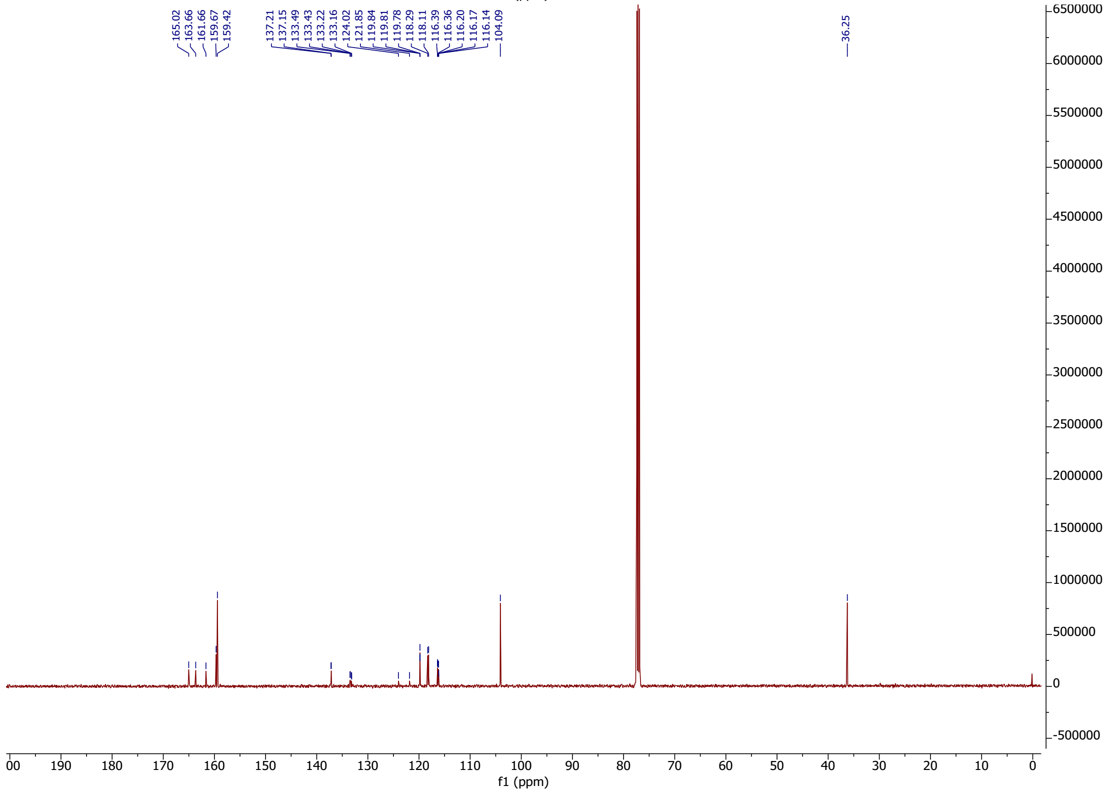

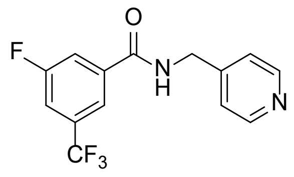

**59**

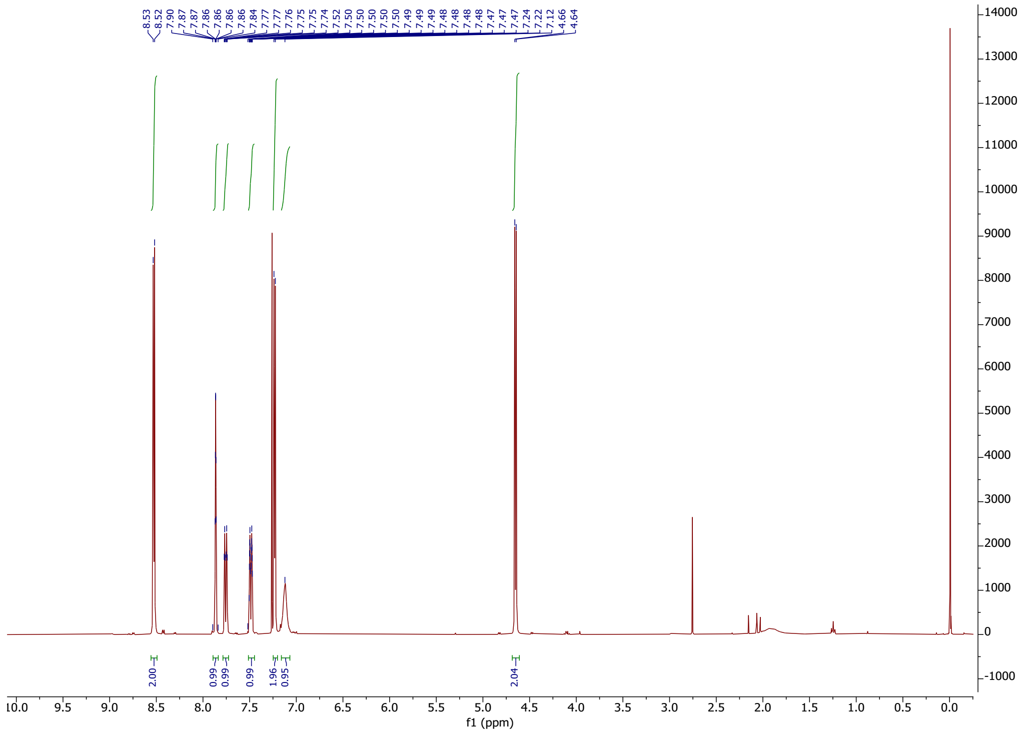

8.53
8.52
7.90
7.87
7.86
7.86
7.86
7.86
7.84
7.77
7.77
7.76
7.75
7.75
7.74
7.52
7.50
7.50
7.50
7.50
7.50
7.49
7.49
7.49
7.48
7.48
7.48
7.47
7.47
7.47
7.24
7.22
7.12
4.66
4.64
2.00
0.99
0.99
0.99
1.96
0.95
2.04
f1 (ppm)


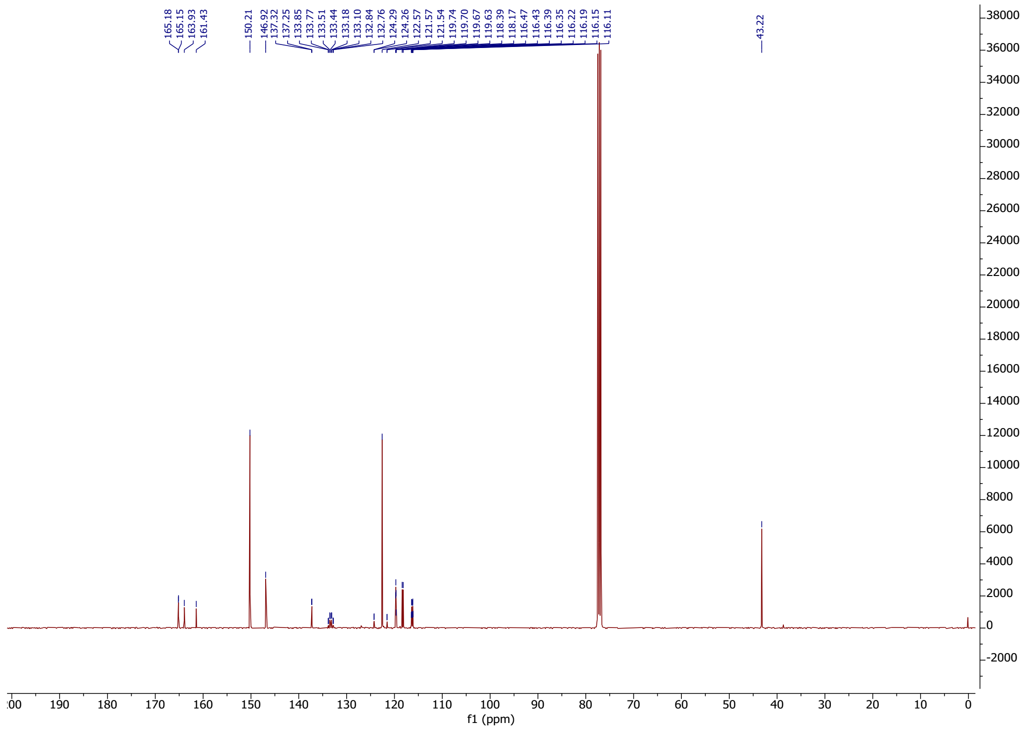

165.18
165.15
163.93
161.43
150.21
146.92
137.32
137.25
133.85
133.77
133.51
133.44
133.18
133.10
132.84
132.76
124.29
124.26
122.57
121.54
119.74
119.70
119.67
119.63
118.39
118.17
116.47
116.43
116.39
116.35
116.22
116.19
116.15
116.11
43.22
f1 (ppm)

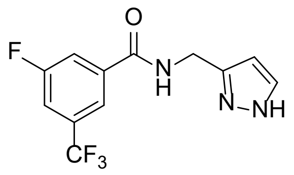

O
F
N
H
N
NH
CF3


**60**

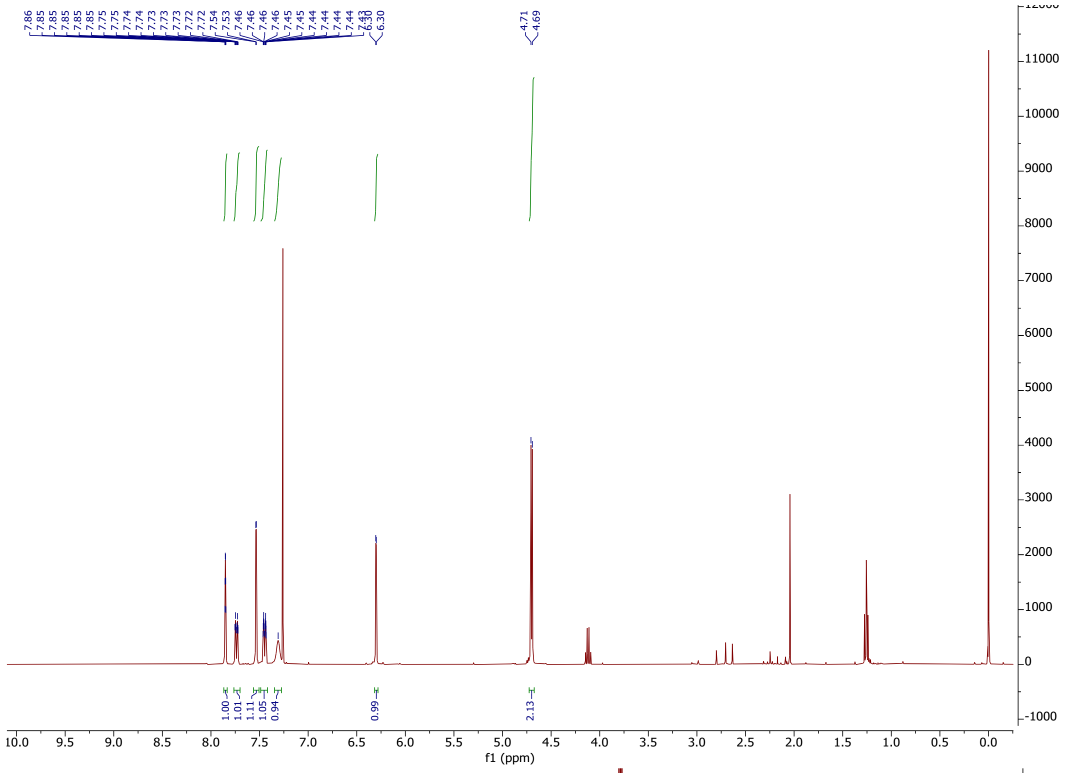

7.86
7.85
7.85
7.85
7.85
7.75
7.75
7.74
7.74
7.73
7.73
7.73
7.73
7.72
7.72
7.54
7.53
7.46
7.46
7.46
7.46
7.45
7.45
7.44
7.44
7.44
7.44
7.43
6.30
6.30
4.71
4.69
1.00
1.01
1.11
1.05
0.94
0.99
2.13
f1 (ppm)


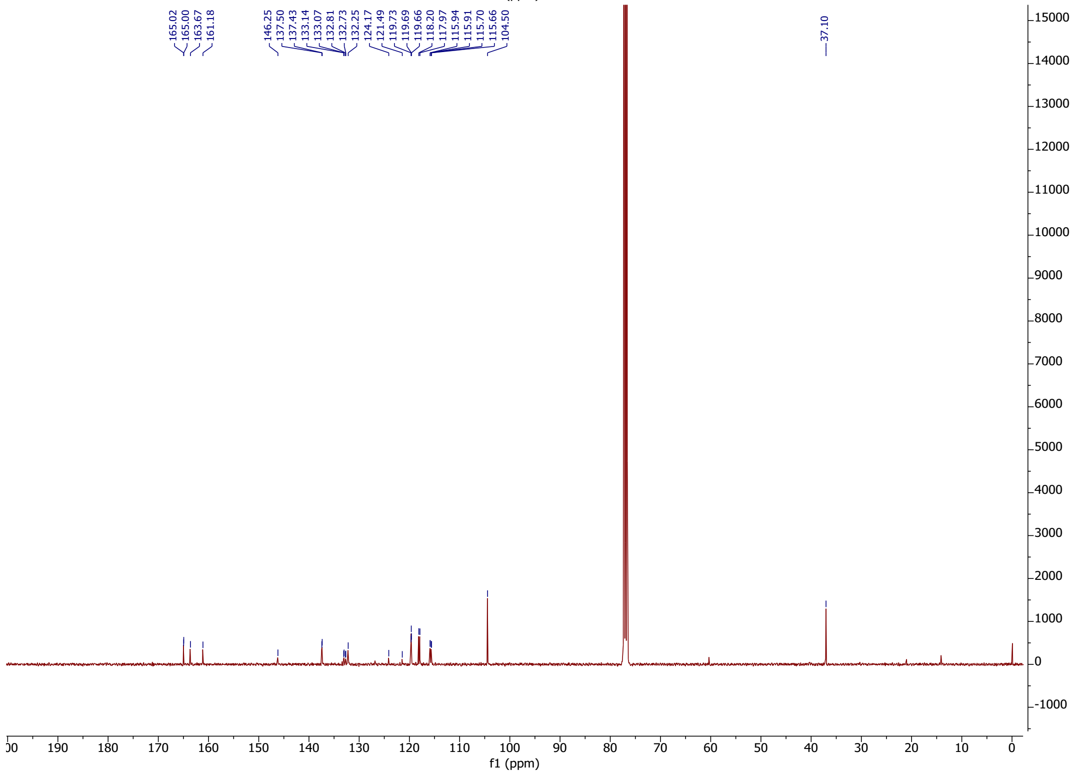

165.02
165.00
163.67
161.18
146.25
137.50
137.43
133.14
133.07
132.81
132.73
132.25
124.17
121.49
119.73
119.69
119.66
118.20
117.97
115.94
115.91
115.70
115.66
104.50
37.10
f1 (ppm)

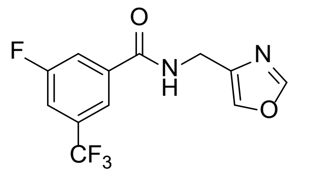
F
O
N
H
N
O
CF3

**61**

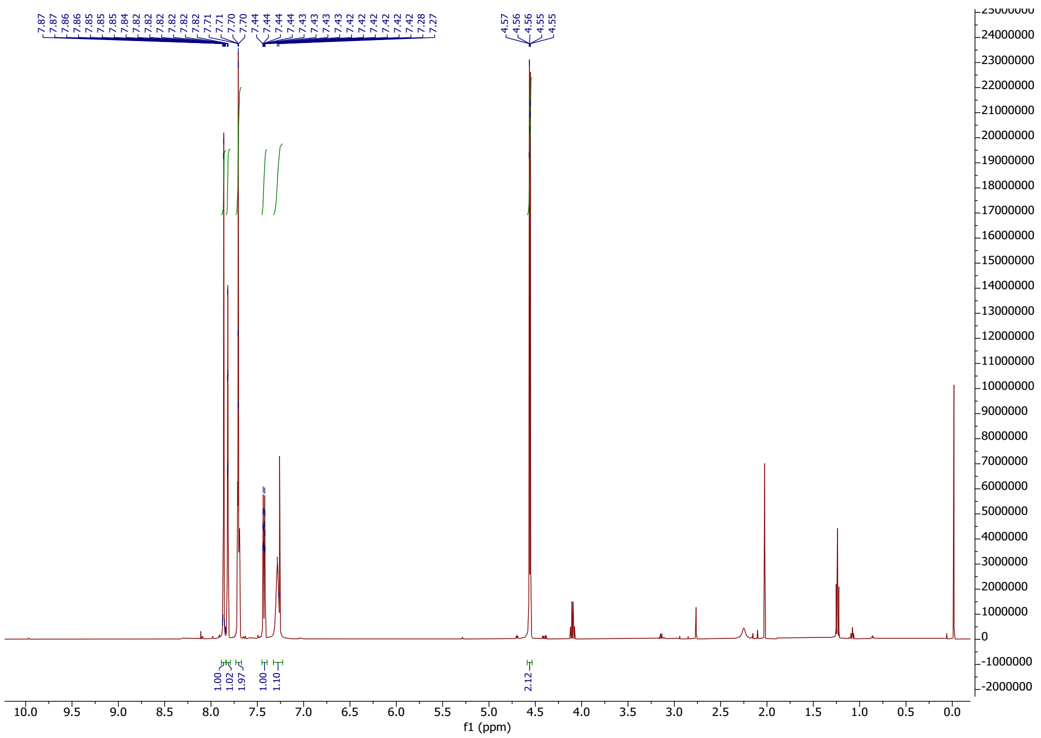
7.87 7.87 7.86 7.86 7.85 7.85 7.85 7.84 7.82 7.82 7.82 7.82 7.82 7.82 7.71 7.71 7.70 7.70 7.44 7.44 7.44 7.44 7.43 7.43 7.43 7.43 7.42 7.42 7.42 7.42 7.42 7.42 7.28 7.27
4.57 4.56 4.56 4.55 4.55
1.00 1.02 1.97 1.00 1.10
2.12
f1 (ppm)

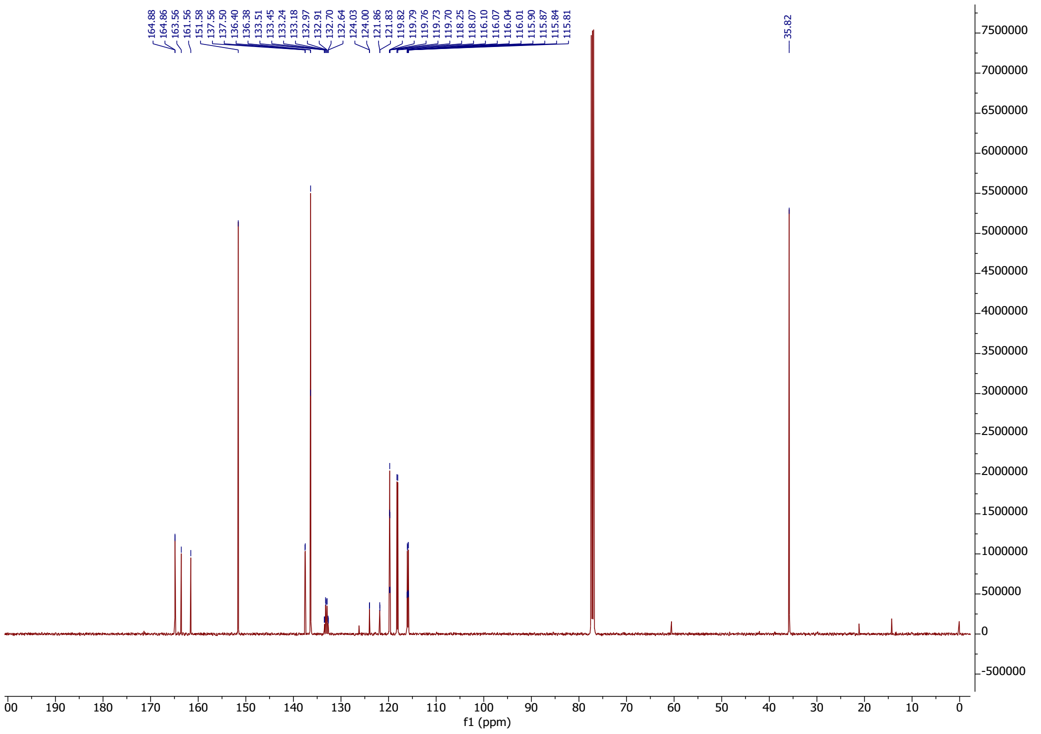
164.88 164.86 163.56 161.56 151.58 137.56 137.50 136.40 136.38 133.51 133.45 133.24 133.18 132.97 132.91 132.70 132.64 124.03 124.00 121.86 121.83 119.82 119.79 119.76 119.73 119.70 118.25 118.07 116.10 116.07 116.04 116.01 115.90 115.87 115.84 115.81
35.82
f1 (ppm)

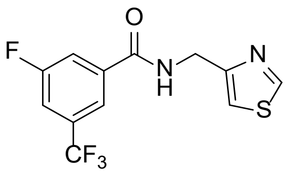

F
O
N
H
N
S
CF3


**62**

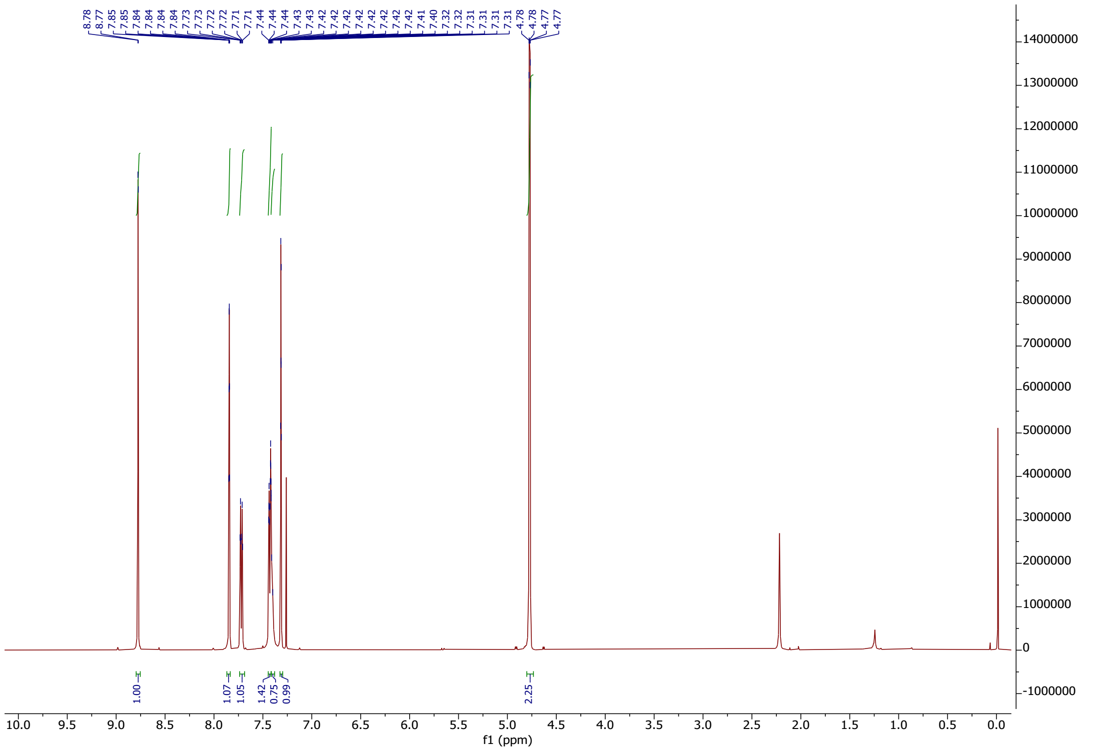

8.78
8.77
7.85
7.85
7.84
7.84
7.84
7.84
7.73
7.73
7.72
7.72
7.71
7.71
7.44
7.44
7.44
7.43
7.43
7.42
7.42
7.42
7.42
7.42
7.42
7.42
7.41
7.40
7.32
7.32
7.31
7.31
7.31
7.31
4.78
4.78
4.77
4.77
1.00
1.07
1.05
1.42
0.75
0.99
2.25
f1 (ppm)


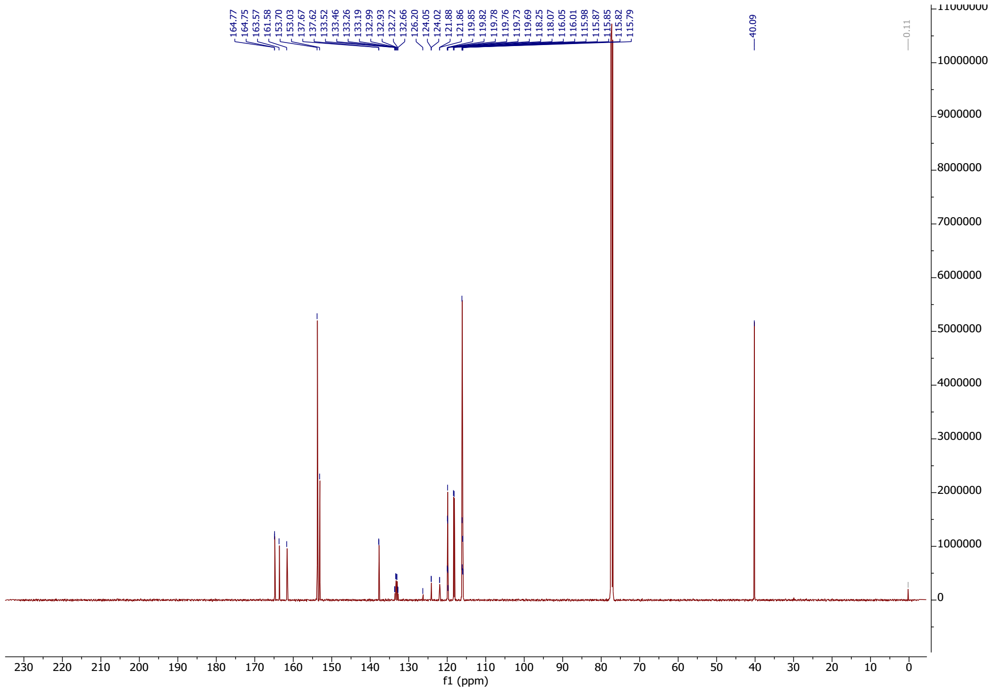

164.77
164.75
163.57
161.58
153.70
153.03
137.67
137.62
133.52
133.46
133.26
133.19
132.99
132.93
132.72
132.66
126.20
124.05
124.02
121.88
121.86
119.85
119.82
119.78
119.76
119.73
119.69
118.25
118.07
116.05
116.01
115.98
115.87
115.85
115.82
115.79
40.09
0.11
f1 (ppm)

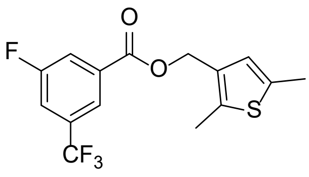

**63**

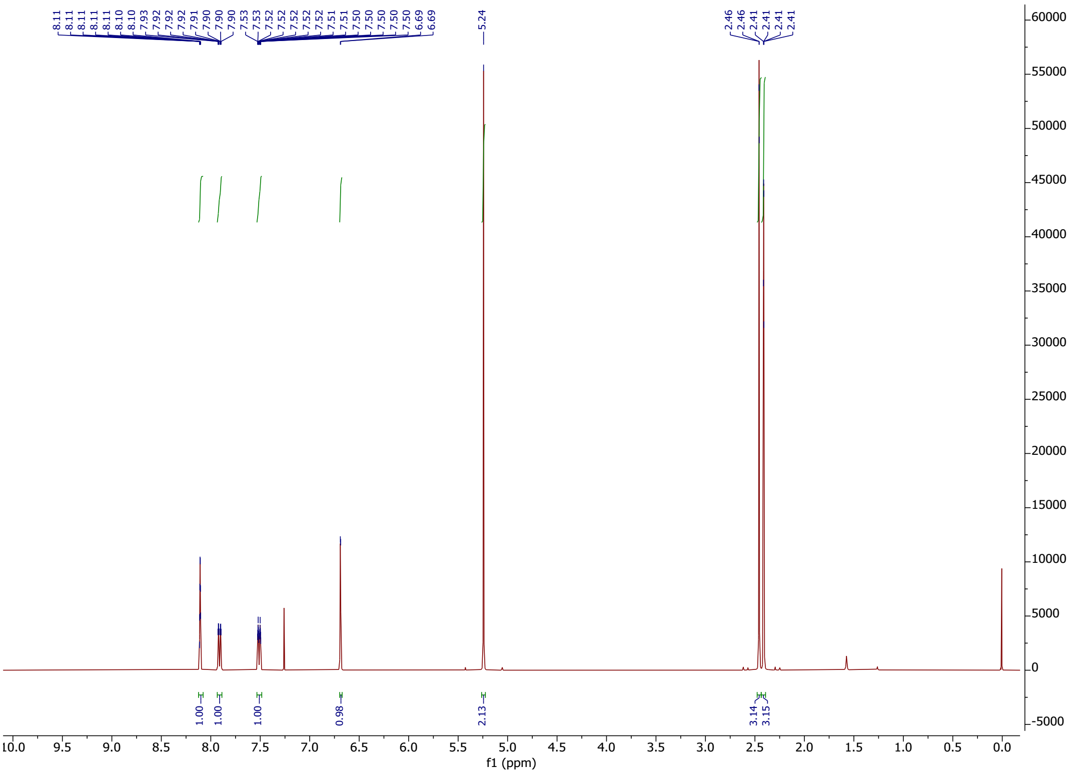

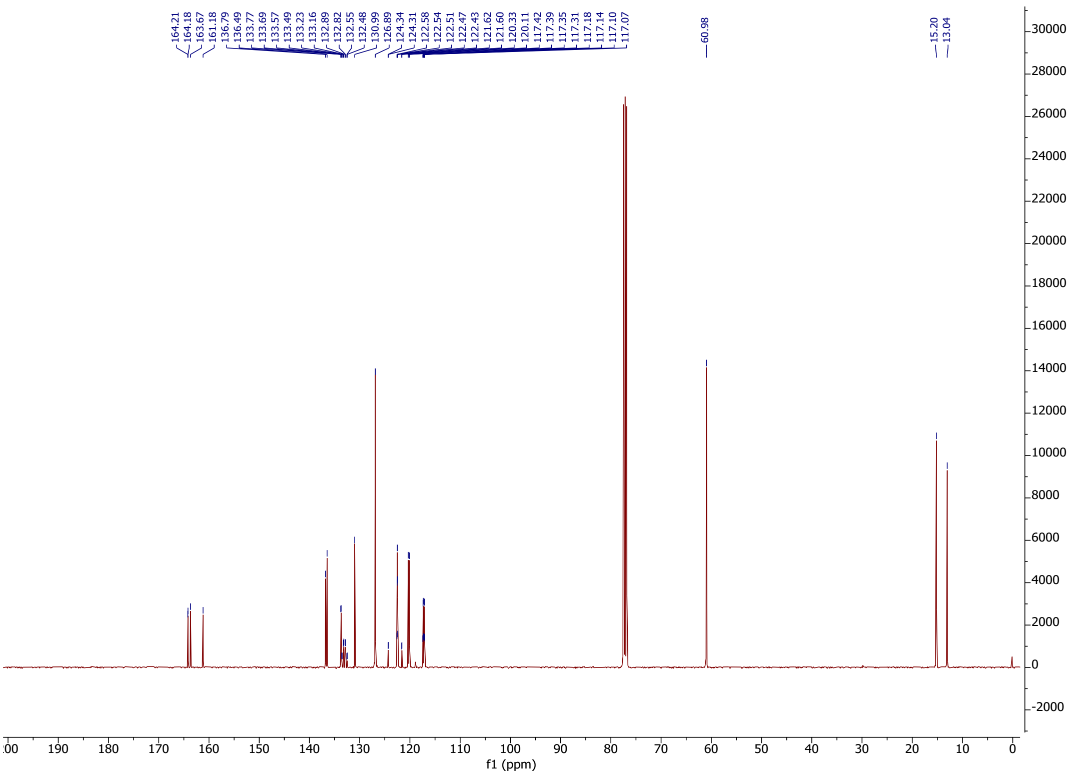

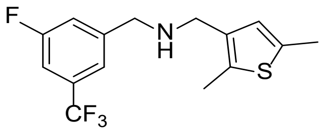

**64**

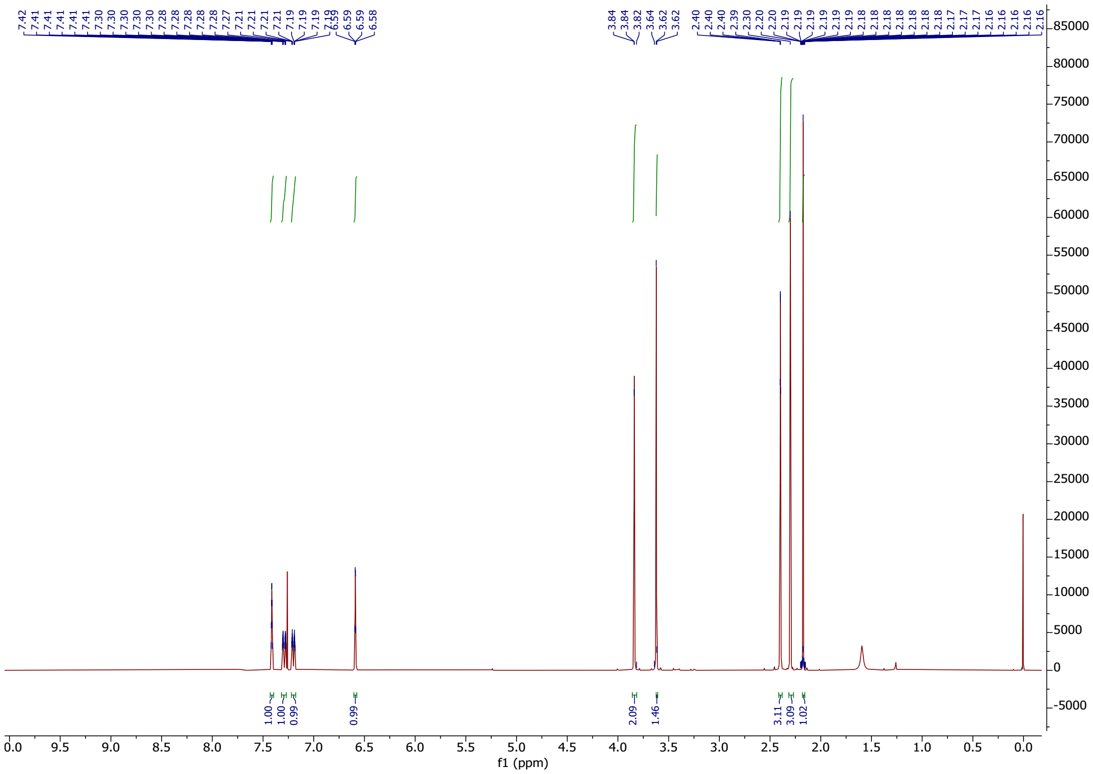

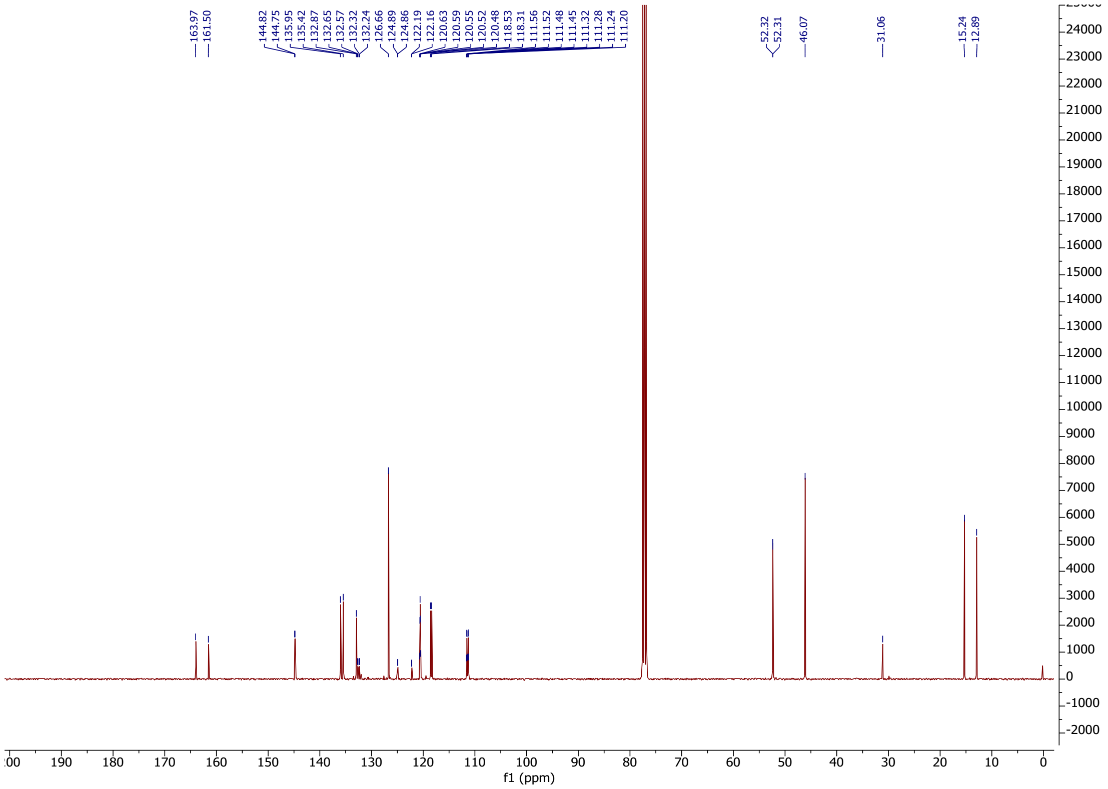

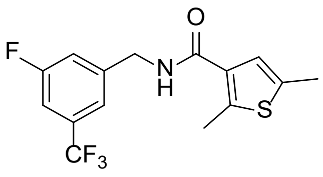

**65**

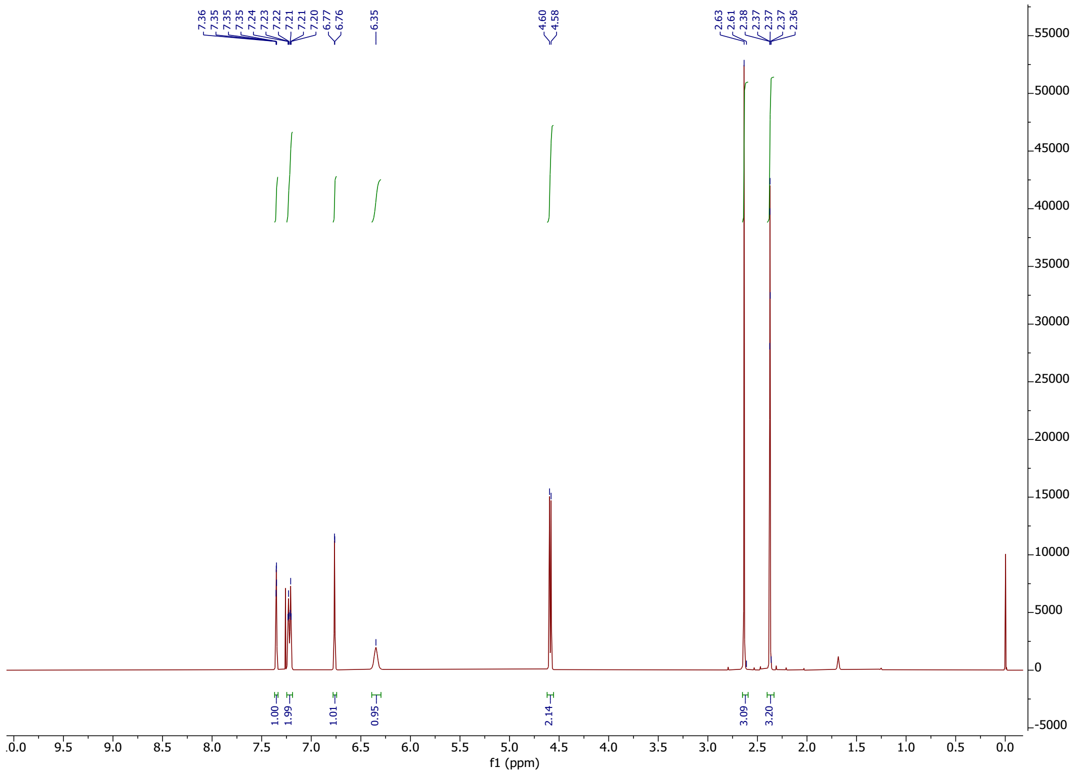

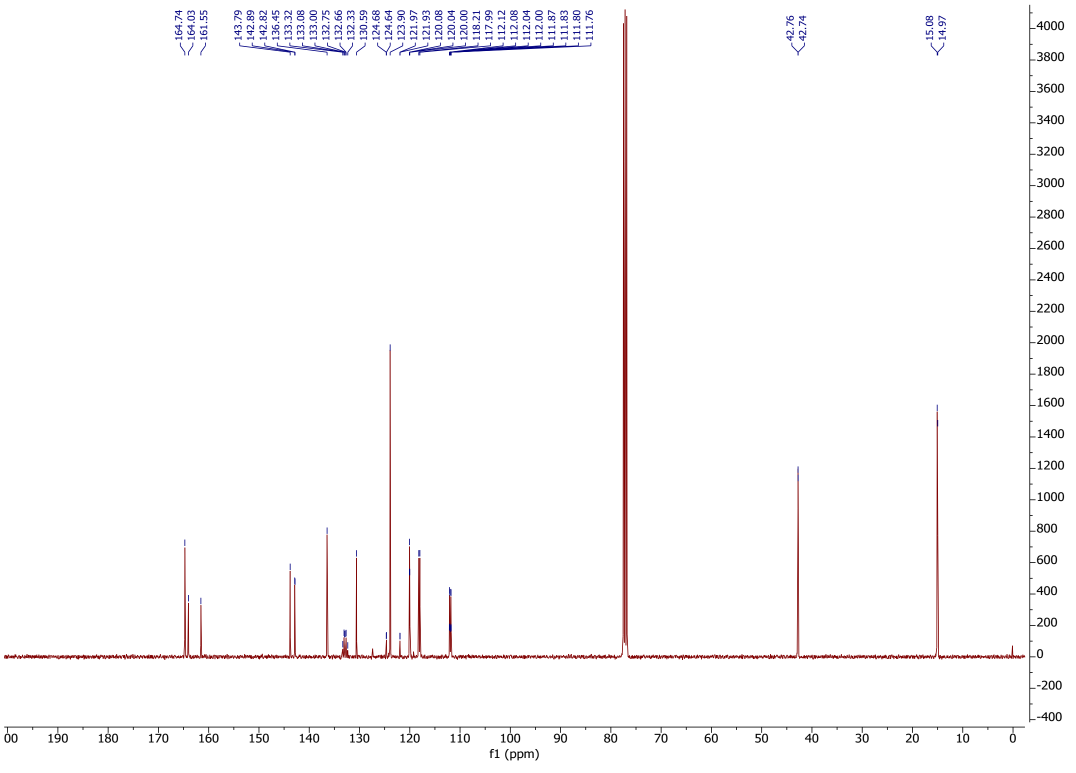

| Compound | Molecular Formula | Calculated | | | Found | | |
|---|---|---|---|---|---|---|---|
| | | C | H | N | C | H | N |
| **22** | $C_{15}H_{13}F_4NOS$ | 54.38 | 3.95 | 4.23 | 54.56 | 4.13 | 4.13 |
| **23 (VF-57a)** | $C_{15}H_{13}ClF_3NOS$ | 51.80 | 3.77 | 4.03 | 51.90 | 3.98 | 3.97 |
| **24** | $C_{15}H_{13}BrF_3NOS$ | 45.93 | 3.34 | 3.57 | 45.98 | 3.52 | 3.53 |
| **25** | $C_{16}H_{13}F_6NOS$ | 50.40 | 3.44 | 3.67 | 50.58 | 3.67 | 3.56 |
| **26** | $C_{14}H_{13}F_2NOS\cdot 1H_2O$ | 56.18 | 5.05 | 4.68 | 56.27 | 4.65 | 4.56 |
| **27** | $C_{14}H_{13}Cl_2NOS\cdot 1.25H_2O$ | 49.93 | 4.64 | 4.16 | 49.39 | 3.86 | 3.91 |
| **28** | $C_{14}H_{13}Br_2NOS$ | 41.71 | 3.25 | 3.47 | 42.10 | 3.17 | 3.37 |
| **29** | $C_{15}H_{16}ClNOS$ | 61.32 | 5.49 | 4.77 | 61.34 | 5.58 | 4.62 |
| **30** | $C_{14}H_{14}FNOS$ | 63.86 | 5.36 | 5.32 | 63.87 | 5.46 | 5.22 |
| **31** | $C_{15}H_{14}F_3NOS$ | 57.50 | 4.50 | 4.47 | 57.78 | 4.75 | 4.33 |
| **32** | $C_{14}H_{14}ClNOS$ | 60.10 | 5.04 | 5.01 | 60.32 | 4.77 | 4.90 |
| **33** | $C_{14}H_{14}BrNOS\cdot 2H_2O$ | 46.68 | 5.04 | 3.89 | 47.10 | 3.88 | 3.77 |
| **34** | $C_{14}H_{14}N_2O_3S$ | 57.92 | 4.86 | 9.65 | 58.20 | 5.08 | 9.48 |
| **35** | $C_{15}H_{12}F_5NOS\cdot 0.2\ C_3H_7NO$ | 51.57 | 3.46 | 4.01 | 51.60 | 3.42 | 4.85 |
| **36** | $C_{15}H_{11}F_6NOS$ | 49.05 | 3.02 | 3.81 | 49.21 | 3.14 | 3.74 |
| **37** | $C_{15}H_{12}ClF_4NOS$ | 49.25 | 3.31 | 3.83 | 49.40 | 3.21 | 3.66 |
| **38** | $C_{15}H_{12}ClF_4NOS$ | 49.25 | 3.31 | 3.83 | 49.22 | 3.29 | 3.72 |
| **39** | $C_{15}H_{12}Cl_2F_3NOS$ | 47.13 | 3.16 | 3.66 | 47.01 | 3.10 | 3.57 |
| **40** | $C_{14}H_{15}NOS$ | 68.54 | 6.16 | 5.71 | 68.52 | 6.10 | 5.60 |
| **41** | $C_{14}H_{14}ClNOS$ | 60.10 | 5.04 | 5.01 | 60.40 | 5.00 | 4.96 |
| **42** | $C_{15}H_{17}NOS$ | 69.46 | 6.61 | 5.40 | 69.24 | 6.55 | 5.28 |
| **43** | $C_{15}H_{17}NO_2S$ | 65.43 | 6.22 | 5.09 | 65.27 | 6.25 | 4.93 |
| **44** | $C_{14}H_{13}Cl_2NOS$ | 53.51 | 4.17 | 4.46 | 53.61 | 4.08 | 4.35 |
| **45** | $C_{14}H_{11}F_4NOS$ | 52.99 | 3.49 | 4.41 | 53.14 | 3.58 | 4.33 |
| **46** | $C_{14}H_{11}F_4NOS$ | 53.00 | 3.49 | 4.41 | 53.05 | 3.41 | 4.34 |
| **47** | $C_{13}H_9F_4NOS$ | 51.49 | 2.99 | 4.62 | 51.69 | 2.86 | 4.50 |
| **48** | $C_{14}H_{11}ClF_3NOS$ | 50.38 | 3.32 | 4.20 | 50.50 | 3.26 | 4.03 |
| **49** | $C_{14}H_{11}ClF_3NOS$ | 50.38 | 3.32 | 4.20 | 50.36 | 3.20 | 4.17 |
| **50** | $C_{13}H_9ClF_3NOS$ | 48.84 | 2.84 | 4.38 | 49.08 | 2.83 | 4.16 |
| **51** | $C_{14}H_{10}F_5NOS$ | 50.15 | 3.01 | 4.18 | 50.06 | 3.00 | 4.05 |
| **52** | $C_{14}H_{10}ClF_4NOS$ | 47.81 | 2.87 | 3.98 | 47.81 | 2.86 | 3.85 |
| **53** | $C_{14}H_{10}Cl_2F_3NOS$ | 45.67 | 2.74 | 3.80 | 45.68 | 2.67 | 3.74 |
| **54** | $C_{15}H_{13}F_4NO_2$ | 57.15 | 4.16 | 4.44 | 57.06 | 4.30 | 4.31 |
| **55** | $C_{15}H_{13}ClF_3NO_2$ | 54.31 | 3.95 | 4.22 | 54.29 | 4.09 | 4.03 |
| **56** | $C_{13}H_9F_4NO_2\cdot 0.35\ H_2O$ | 53.20 | 3.33 | 4.77 | 53.06 | 3.15 | 4.67 |
| **57** | $C_{12}H_8F_4N_2OS$ | 47.37 | 2.65 | 9.21 | 47.36 | 2.67 | 9.05 |
| **58** | $C_{12}H_8F_4N_2O_2$ | 50.01 | 2.80 | 9.72 | 50.16 | 2.64 | 9.55 |
| **59** | $C_{14}H_{10}F_4N_2O$ | 56.38 | 3.38 | 9.39 | 55.35 | 3.41 | 9.08 |
| **60** | $C_{12}H_9F_4N_3O$ | 50.18 | 3.16 | 14.63 | 50.26 | 3.23 | 14.32 |
| **61** | $C_{12}H_8F_4N_2O_2$ | 50.01 | 2.80 | 9.72 | 50.24 | 2.67 | 9.50 |
| **62** | $C_{12}H_8F_4N_2OS\cdot 0.3\ C_3H_6O$ | 47.65 | 2.80 | 9.03 | 47.85 | 2.54 | 9.05 |
| **63** | $C_{15}H_{12}F_4O_2S$ | 54.22 | - | 3.64 | 54.44 | - | 3.54 |
| **64** | $C_{15}H_{15}F_4NS$ | 56.77 | 4.76 | 4.41 | 56.85 | 4.82 | 4.63 |
| **65** | $C_{15}H_{13}F_4NOS$ | 54.38 | 3.95 | 4.23 | 54.46 | 3.83 | 4.19 |

**Table S1.** Elemental analysis data.

```
                        1        10         20         30         40         50
A/Victoria/361/2011     QKLPGNDNSTATLCLGHHAVPNGTIVKTITNDQIEVTNATELVQNSSIGEICDSPH-QIL
A/bald-eagle/FL/2022    -----SLVKSDQICIGYHANNSTEQVDTIMEKNVTVTHAQDILEKTHNGKLCDLNGVKPL
A/Virginia/ATCC3/2009   -----ATANADTLCIGYHANNSTDTVDTVLEKNVTVTHSVNLLEDKHNGKLCKLRGVAPL
A/Puerto-Rico/8/34      -----AAADADTICIGYHANNSTDTVDTVLEKNVTVTHSVNLLEDSHNGKLCRLKGIAPL
                                .:  :*:*:**  .    *.*: :.:: **:: ::::..  *::*        *

                       60         70         80          90        100        110
A/Victoria/361/2011     DGENCTLIDALLGDPQCDGFQN-KKWDLFVERSK-AYSNCYPYDVPDYASLRSLVASSGT
A/bald-eagle/FL/2022    ILKDCSVAGWLLGNPMCDEFTRVPEWSYIVERANPANDLCYPGSLNDYEELKHMLSRINH
A/Virginia/ATCC3/2009   HLGKCNIAGWILGNPECESLSTASSWSYIVETSSSDNGTCYPGDFIDYEELREQLSSVSS
A/Puerto-Rico/8/34      QLGKCNIAGWLLGNPECDPLLPVRSWSYIVETPNSENGICYPGDFIDYEELREQLSSVSS
                            .*.: . :**:* *: :     .*. :**   .    .  *** .. ** .*:   ::  .

                           120        130        140        150        160        170
A/Victoria/361/2011     LE---FNNESFNWTG-VTQNGTSSACIRRSNNSFFSRLNWLTRLNFKYPALNVTMPNNEQ
A/bald-eagle/FL/2022    FEKILIIPK-SSWPNHETSLGVSAACPYQGAPSFFRNVVWLIKKNDAYPTIKISYNNTNR
A/Virginia/ATCC3/2009   FERFEIFPKTSSWPNHDSNKGVTAACPHAGAKSFYKNLIWLVKKENSYPKLSKSYINDKG
A/Puerto-Rico/8/34      FERFEIFPKESSWPNHNT-NGVTAACSHEGKSSFYRNLLWLTEKEGSYPKLKNSYVNKKG
                        :*   :  :  .*  .  :   *.::**    .   **: .: **  . :   ** :. :  * :

                            180        190        200        210        220        230
A/Victoria/361/2011     FDKLYIWGVHHPVTDKDQIFLYAQSSGRITVSTKRSQQAVIPNIGYRPRIRNIPSRISIY
A/bald-eagle/FL/2022    EDLLILWGIHHSNNAEEQTNLYKNPTTYISVGTSTLNQRLAPKIATRSQVNGQRGRMDFF
A/Virginia/ATCC3/2009   KEVLVLWGIHHPSTSADQQSLYQNADAYVFVGTSRYSKKFKPEIAIRPKVRNQEGRMNYY
A/Puerto-Rico/8/34      KEVLVLWGIHHPPNSKEQQNLYQNENAYVSVVTSNYNRRFTPEIAERPKVRDQAGRMNYY
                         : * :**:**   .  :*  ** :     : * *.   .: . *:*. * ::..   .*:. :

                            240        250        260         270        280        290
A/Victoria/361/2011     WTIVKPGDILLINSTGNLIAPRGYFKI-RSGKSSIMRSDAPIGKCNSECITPNGSIPNDK
A/bald-eagle/FL/2022    WTILKPDDAIHFESNGNFIAPEYAYKIVKKGDSTIMKSGVEYGHCNTKCQTPVGAINSSM
A/Virginia/ATCC3/2009   WTLVEPGDKITFEATGNLVVPRYAFAMERNAGSGIIISDTPVHDCNTTCQTPKGAINTSL
A/Puerto-Rico/8/34      WTLLKPGDTIIFEANGNLIAPMYAFALSRGFGSGIITSNASMHECNTKCQTPLGAINSSL
                        **:::*.* : :::.**::.*   : : :    * *: *..   .**: * ** *:* ..

                             300        310        320
A/Victoria/361/2011     PFQNVNRITYGACPRYVKQSTLKLATGMRNVPEKQTR---
A/bald-eagle/FL/2022    PFHNIHPLTIGECPKYVKSNKLVLATGLRNSPLREKRRKR
A/Virginia/ATCC3/2009   PFQNIHPITIGKCPKYVKSTKLRLATGLRNVPSIQSR---
A/Puerto-Rico/8/34      PYQNIHPVTIGECPKYVRSAKLRMVTGLRNIPSIQSR---
                        *::*:: :* * **:**:. .* :.**:** *  :.*

                        1        10         20         30         40         50         60
A/Victoria/361/2011     GIFGAIAGFIENGWEGMVDGWYGFRHQNSEGRGQAADLKSTQAAIDQINGKLNRLIGKTN
A/bald-eagle/FL/2022    GLFGAIAGFIEGGWQGMVDGWYGYHHSNEQGSGYAADKESTQKAIDGVTNKVNSIIDKMN
A/Virginia/ATCC3/2009   GLFGAIAGFIEGGWTGMVDGWYGYHHQNEQGSGYAADLKSTQNAIDEITNKVNSVIEKMN
A/Puerto-Rico/8/34      GLFGAIAGFIEGGWTGMIDGWYGYHHQNEQGSGYAADQKSTQNAINGITNKVNTVIEKMN
                        *:*********.** **:*****::*.*.:* * *** :*** **: :..*:*:* * *

                                 70         80         90        100        110        120
A/Victoria/361/2011     EKFHQIEKEFSEVEGRIQDLEKYVEDTKIDLWSYNAELLVALENQHTIDLTDSEMNKLFE
A/bald-eagle/FL/2022    TQFEAVGREFNNLERRIENLNKKMEDGFLDVWTYNAELLVLMENERTLDFHDSNVKNLYD
A/Virginia/ATCC3/2009   TQFTAVGKEFNHLEKRIENLNKKVDDGFLDIWTYNAELLVLLENERTLDYHDSNVKNLYE
A/Puerto-Rico/8/34      IQFTAVGKEFNKLEKRMENLNKKVDDGFLDIWTYNAELLVLLENERTLDFHDSNVKNLYE
                         :*  : :**..:* *:::*:* ::*   :*:*:******* :**::*:*  **::::*::

                                130        140        150        160        170        180
A/Victoria/361/2011     KTKKQLRENAEDMGNGCFKIYHKCDNACIGSIRNGTYDHDVYRDEALNNRFQIKGVELKS
A/bald-eagle/FL/2022    KVRLQLRDNAKELGNGCFEFYHKCDNECMESVRNGTYDYPQYSEEARLKREEISGVKLES
A/Virginia/ATCC3/2009   KVRSQLKNNAKEIGNGCFEFYHKCDNTCMESVKNGTYDYPKYSEEAKLNREEIDGVKLES
A/Puerto-Rico/8/34      KVKSQLKNNAKEIGNGCFEFYHKCDNECMESVRNGTYDYPKYSEESKLNREKVDGVKLES
                        *.: **::**:::*****::****** *: *::*****:  * :*:  :* ::.**:*:*

                                 190        200        210        220
A/Victoria/361/2011     GYKDWILW-ISFAISCFLLCVALLGFIMWACQKGNIRCNICI
A/bald-eagle/FL/2022    VGTYQILSIYSTAASSLALAIMMAGLSLWMCSNGSLQCRICI
A/Virginia/ATCC3/2009   TRIYQILAIYSTVASSLVLVVSLGAISFWMCSNGSLQCRICI
A/Puerto-Rico/8/34      MGIYQILAIYSTVASSLVLLVSLGAISFWMCSNGSLQCRICI
                             **    * . *.: * : : .: :* *.:*.::*.***
```

**Figure S1.** Sequence alignment for the studied Influenza A viral strains. The green/blue lines denote the region of the sequence corresponding to the HA1/HA2 subunits of HA. The cleavage point is highlighted in bold (black). The T residue involved in site B of (*S*)-F0045 is highlighted in violet. Resistance mutations emerging in A/Virginia/ATCC3/2009 from exposure to RL-007 and **VF-57a** are highlighted in red.

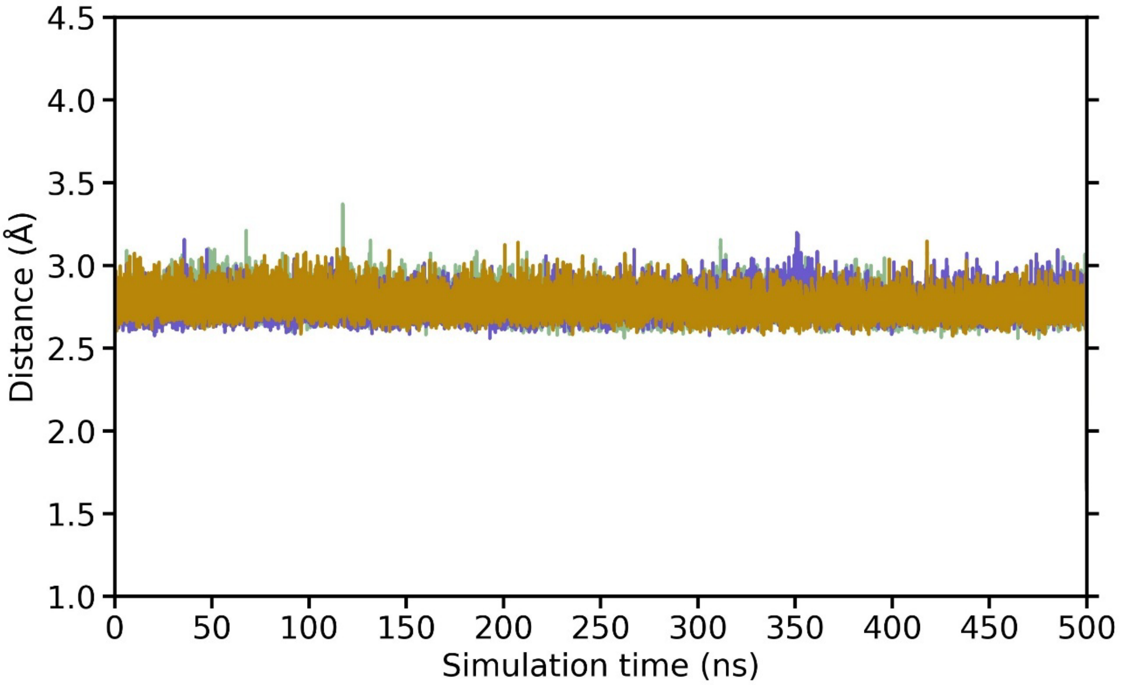


**Figure S2.** Distance between the carbonyl oxygen of (*S*)-F0045 and the hydroxyl oxygen of T318$_1$ (shown in green, lavender and beige) for the simulation of (*S*)-F0045 in site B binding pocket of the H1N1 A/Puerto Rico/8/1934 viral strain.

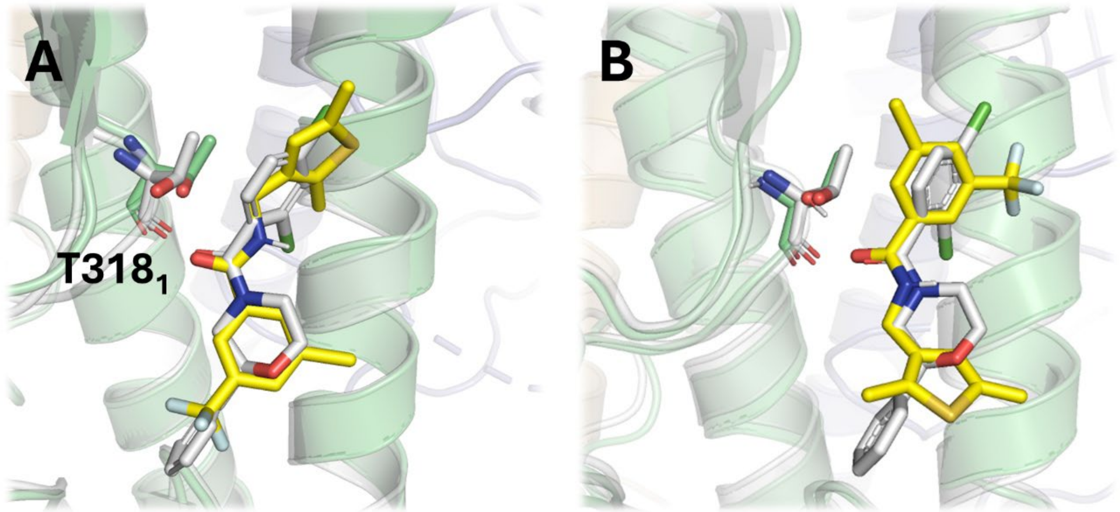


**Figure S3.** Representation of the two starting binding modes examined for **VF-57a** (yellow sticks) upon binding to site B in H1 HA (A/Virginia/ATCC3/2009). The two binding modes differ in the relative position of the thiophene and phenyl rings, allowing the formation of the hydrogen bond with T318$_1$. The X-ray binding mode of (*S*)-F0045 bound to H1 HA is shown as grey sticks.

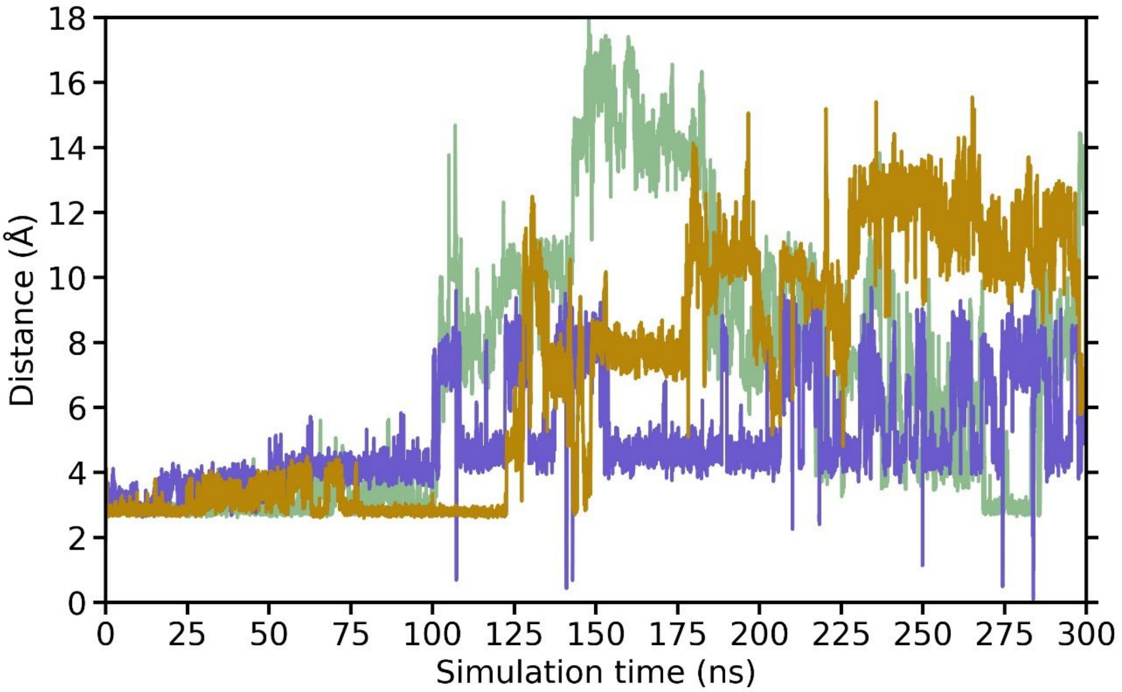


**Figure S4.** Distance between the carbonyl oxygen of **VF-57a** and the hydroxyl oxygen of T318$_1$ (shown in green, lavender and beige) for the simulation of **VF-57a** in site B binding pocket of the H1N1 A/Virginia/ATCC3/2009 viral strain.

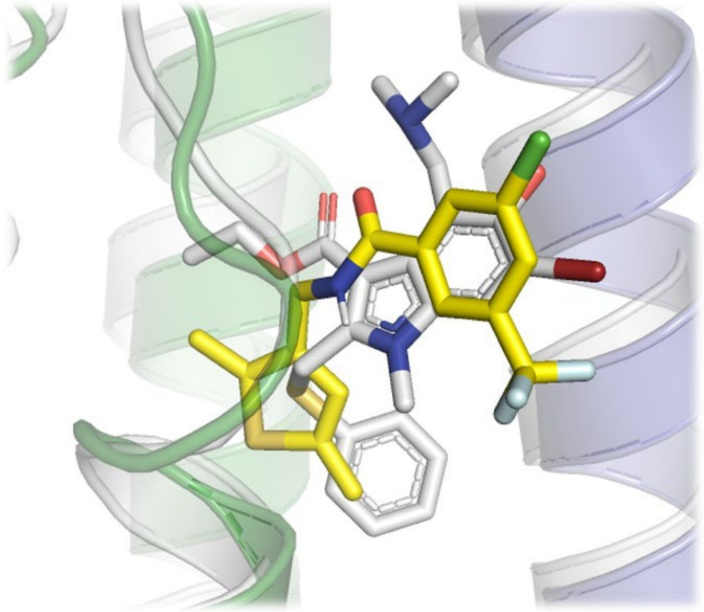

**Figure S5.** Representation of the starting binding mode examined for **VF-57a** ) (yellow sticks) upon binding to site A in H1 HA (A/Virginia/ATCC3/2009). Overlap of the X-ray pose of arbidol bound to the H3 HA (A/Hong Kong/1/1968) (grey sticks) and the binding pose of **VF-57a** in the H1 HA model (A/Virginia/ATCC3/2009.

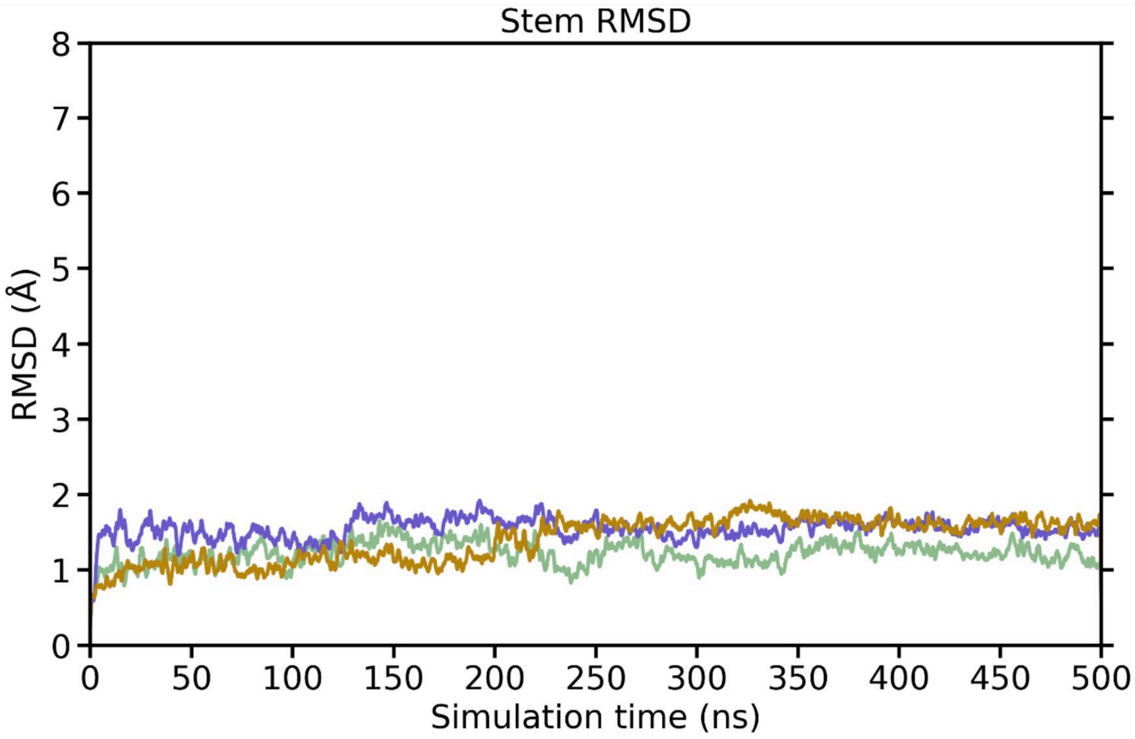


**Figure S6.** RMSD plot for the HA-stem backbone of each protomer (shown in green, lavender and beige) for the simulation of **VF-57a** in site A binding pocket of the H1N1 A/Virginia/ATCC3/2009 viral strain.

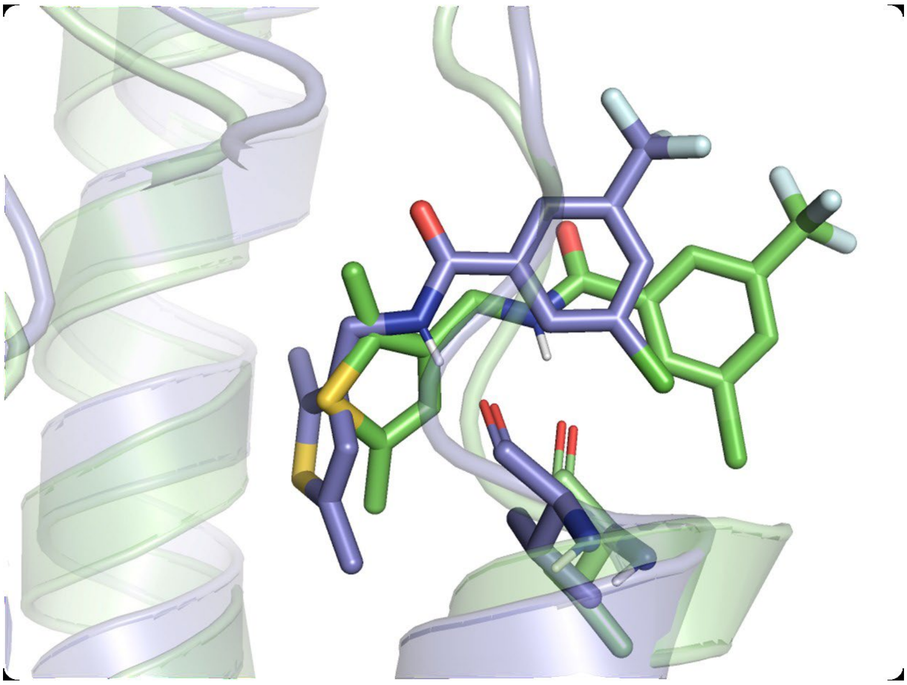

**Figure S7.** Representation of the different arrangement of the thiophene ring in the docked poses of **VF-57a** (green and lavender sticks) at the arbidol binding site upon overlap of two protomers of HA (A/Virginia/ATCC3/2009).

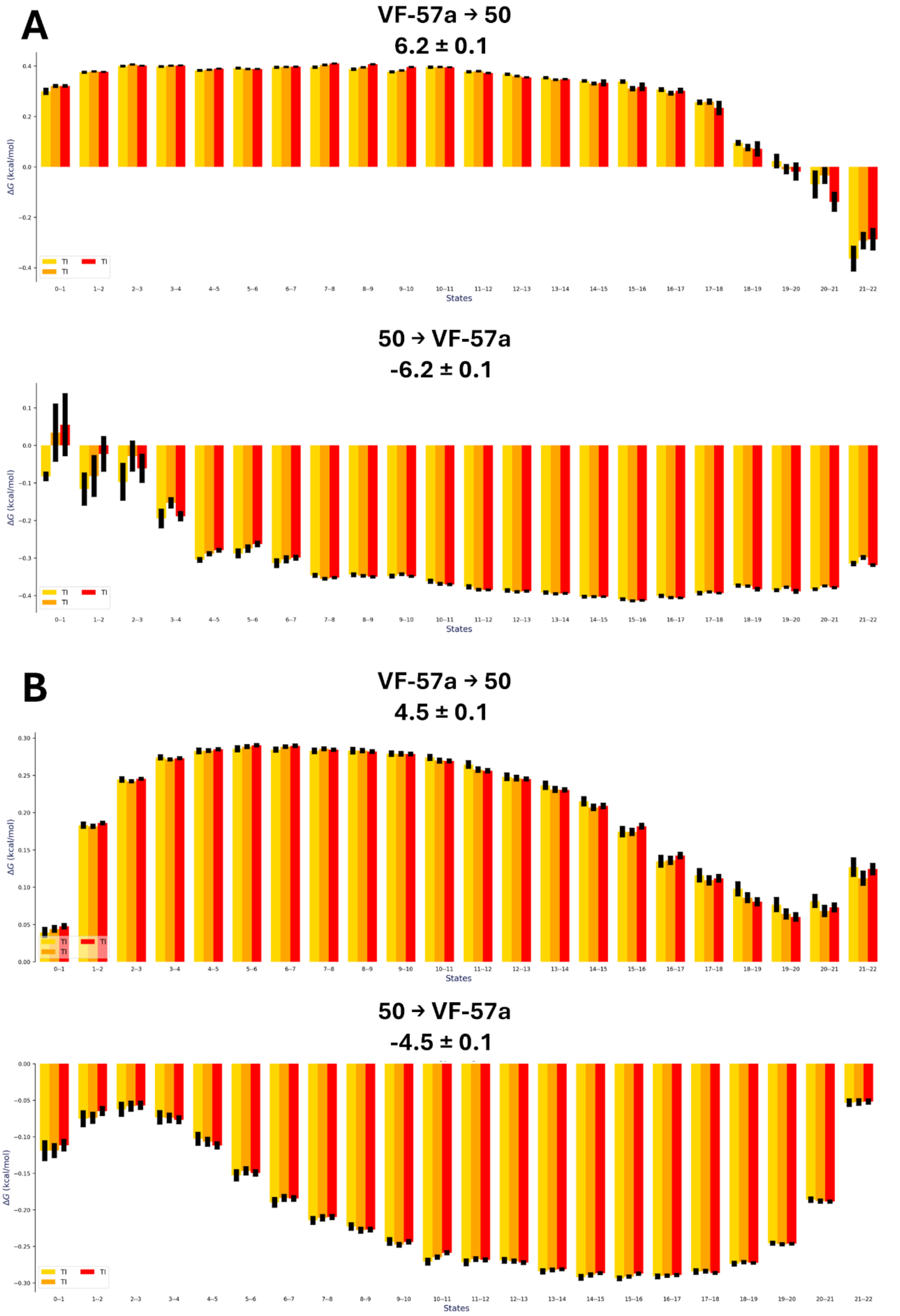


**Figure S8**. Representation of the free energy change determined from the last 6ns of each λ divided in steps of 2 ns (each in yellow, orange and red) for A) **VF-57a**↔50 (top: forward; bottom: backward) transformation in the ligand-protein complex (values in kcal·mol$^{-1}$) and B) TI analysis for the **VF-57a**↔50 (top: forward; bottom: backward) transformation in aqueous solution (values in kcal·mol$^{-1}$).

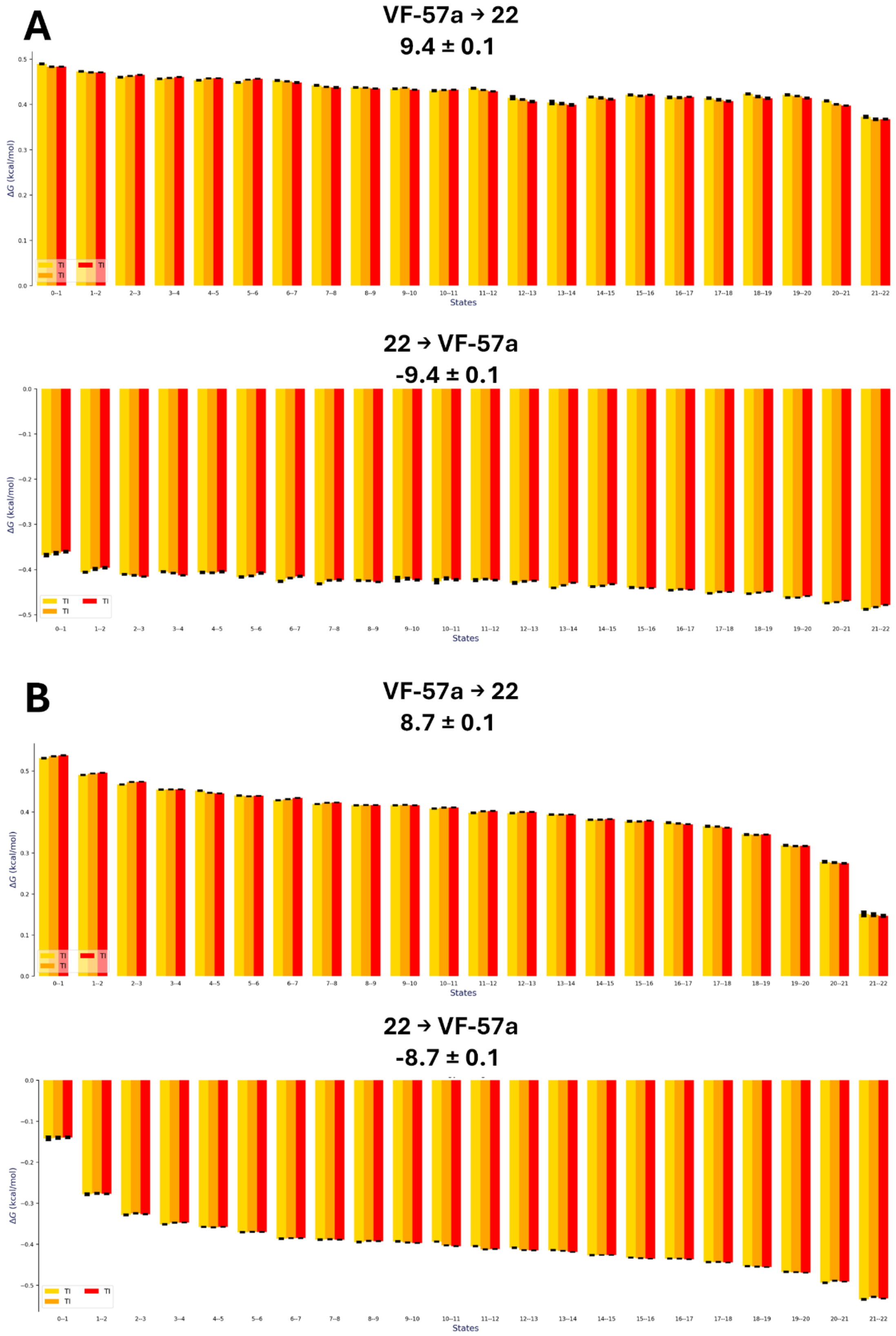


**Figure S9**. Representation of the free energy change determined from the last 6ns of each λ divided in steps of 2 ns (each in yellow, orange and red) for A) **VF-57a**↔22 (top: forward; bottom: backward) transformation in the ligand-protein complex (values in kcal·mol$^{-1}$) and B) TI analysis for the **VF-57a**↔22 (top: forward; bottom: backward) transformation in aqueous solution (values in kcal·mol$^{-1}$).

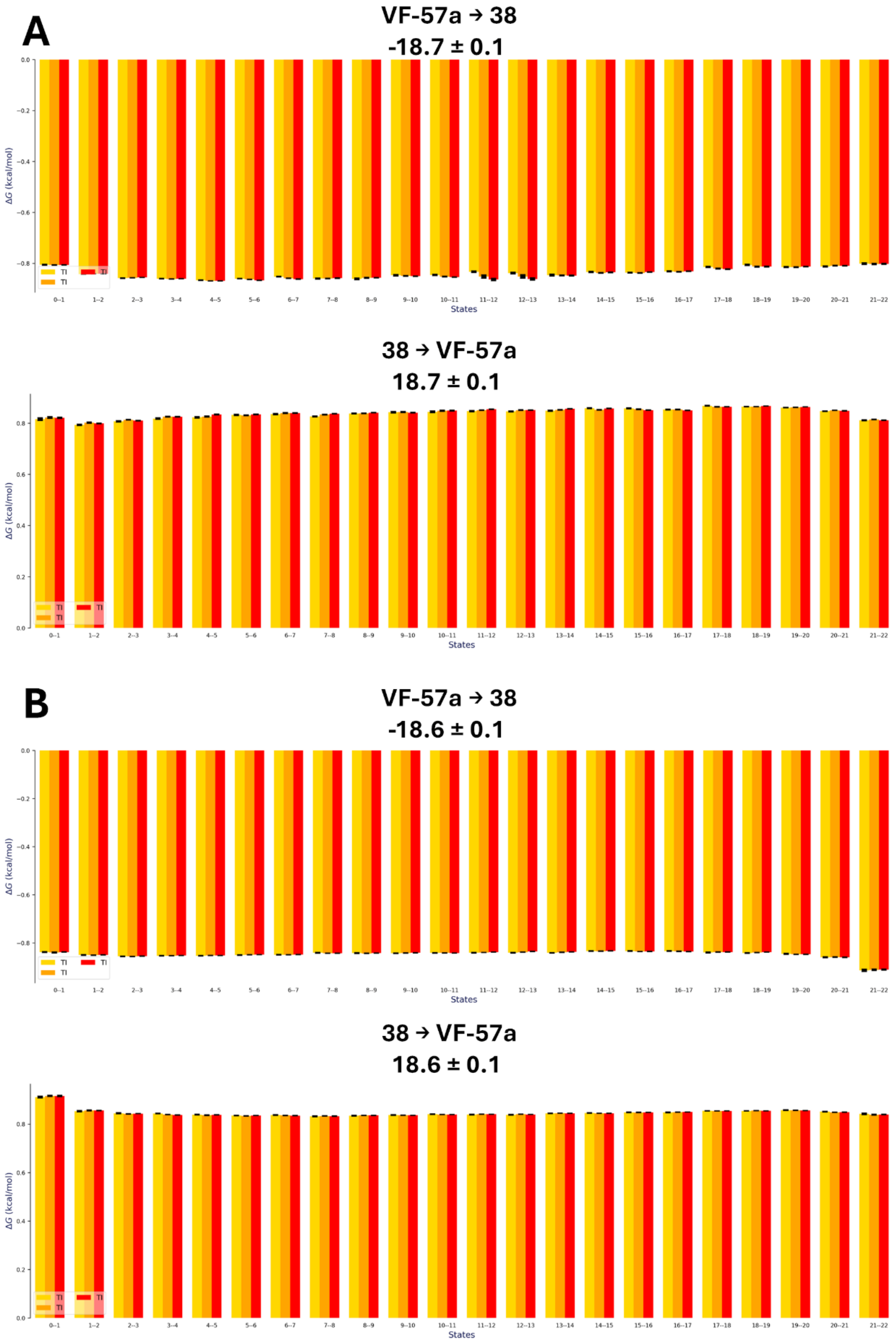


**Figure S10**. Representation of the free energy change determined from the last 6ns of each λ divided in steps of 2 ns (each in yellow, orange and red) for A) **VF-57a**↔38 (top: forward; bottom: backward) transformation in the ligand-protein complex (values in kcal·mol$^{-1}$) and B) TI analysis for the **VF-57a**↔38 (top: forward; bottom: backward) transformation in aqueous solution (values in kcal·mol$^{-1}$).

#### Area Percent Report

UNIVERSITAT DE BARCELONA

**Data file:** D:\Chemstation\1\Data\SVC\2025-06-10_SVC_CEM-VF-57a 2025-06-10 09-36-12\002-P1-A1-BLANCO.D

**Sample name:** BLANCO

| | | | |
|---|---|---|---|
| **Instrument:** | HPLCMS | **Location:** | P1-A1 |
| **Injection date:** | Tuesday, June 10, 2025 | **Injection:** | 1 of 1 |
| **Acq. method:** | Pepito 220-254nm.M | **Injection volume:** | 5.000 |

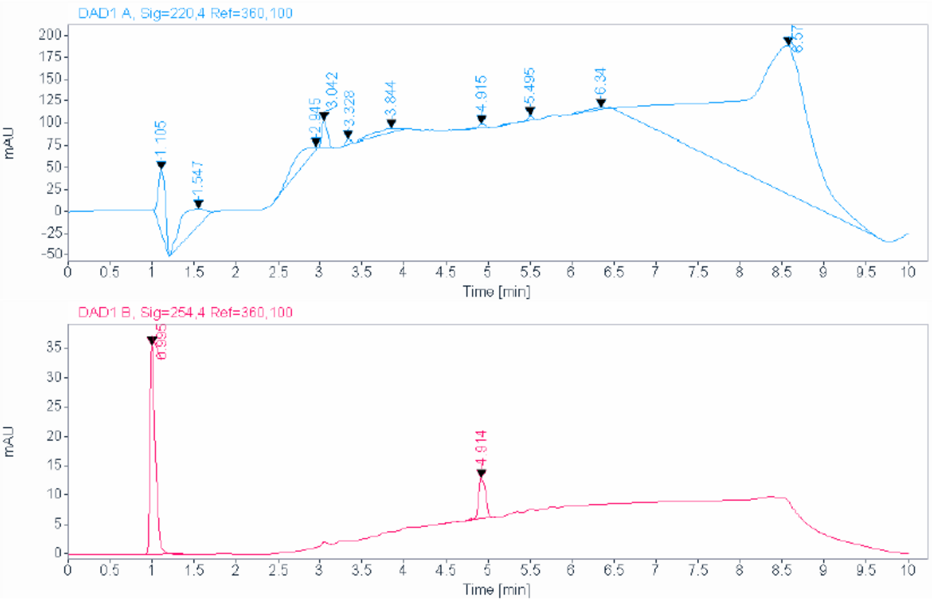


**Signal:** DAD1 A, Sig=220,4 Ref=360,100

| RT [min] | Type | Width [min] | Area | Height | Area% | Name |
|---|---|---|---|---|---|---|
| 1.105 | BB | 0.0988 | 420.2101 | 71.2057 | 3.2990 | |
| 1.547 | BB | 0.3577 | 549.3692 | 20.5612 | 4.3130 | |
| 2.945 | BB | 1.0895 | 342.2197 | 3.7054 | 2.6867 | |
| 3.042 | BB | 0.0759 | 137.4787 | 29.7655 | 1.0793 | |
| 3.328 | BB | 0.0787 | 24.7010 | 5.4697 | 0.1939 | |
| 3.844 | BB | 0.3324 | 130.6983 | 5.1451 | 1.0261 | |
| 4.915 | BB | 0.0746 | 25.2246 | 4.8542 | 0.1980 | |
| 5.495 | BB | 0.0975 | 35.4780 | 5.6336 | 0.2785 | |
| 6.340 | BB | 0.3043 | 51.7367 | 2.2498 | 0.4062 | |
| 8.570 | BB | 0.7761 | 11020.3027 | 169.1470 | 86.5191 | |
| | | Sum | 12737.4193 | | | |

**Signal:** DAD1 B, Sig=254,4 Ref=360,100

| RT [min] | Type | Width [min] | Area | Height | Area% | Name |
|---|---|---|---|---|---|---|
| 0.995 | BB | 0.0658 | 159.2350 | 35.8747 | 78.8943 | |
| 4.914 | BB | 0.0848 | 42.5983 | 7.0180 | 21.1057 | |
| | | Sum | 201.8333 | | | |

Figure S11. HPLC/UV (blank)

### Area Percent Report

| | | | |
|---|---|---|---|
| **Data file:** | D:\Chemstation\1\Data\SVC\2025-06-10_SVC_CEM-VF-57a 2025-06-10 09-36-12\001-P1-B8-CEM-VF-57a.D | | |
| **Sample name:** | CEM-VF-57a | | |
| **Instrument:** | HPLCMS | **Location:** | P1-B8 |
| **Injection date:** | Tuesday, June 10, 2025 | **Injection:** | 1 of 1 |
| **Acq. method:** | Pepito 220-254nm.M | **Injection volume:** | 15.000 |

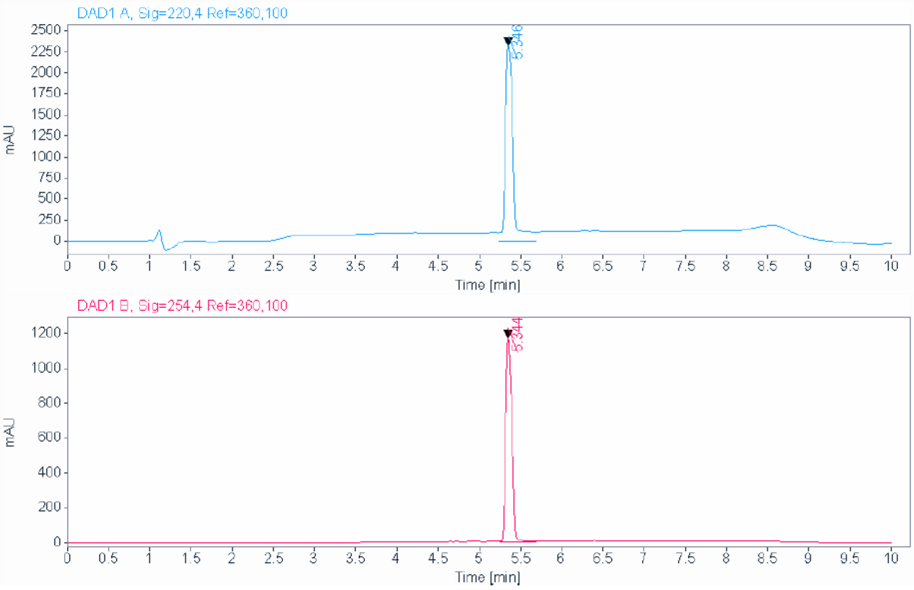


**Signal:** DAD1 A, Sig=220,4 Ref=360,100

| RT [min] | Type | Width [min] | Area | Height | Area% | Name |
|---|---|---|---|---|---|---|
| 5.346 | BB | 0.0902 | 12289.9238 | 2234.2600 | 100.0000 | |
| | | Sum | 12289.9238 | | | |

**Signal:** DAD1 B, Sig=254,4 Ref=360,100

| RT [min] | Type | Width [min] | Area | Height | Area% | Name |
|---|---|---|---|---|---|---|
| 5.344 | BB | 0.0846 | 5883.4629 | 1171.2074 | 100.0000 | |
| | | Sum | 5883.4629 | | | |

Figure S12. HPLC/UV (compound **VF-57a**)
